# Overlapping Neural Correlates Underpin Theory of Mind and Semantic Cognition: Evidence from a Meta-Analysis of 344 Functional Neuroimaging Studies

**DOI:** 10.1101/2023.08.16.553506

**Authors:** Eva Balgova, Veronica Diveica, Rebecca L. Jackson, Richard J. Binney

**Affiliations:** Cognitive Neuroscience Institute, Department of Psychology, Bangor University, UK; Department of Psychology, Aberystwyth University, UK; Montreal Neurological Institute, Department of Neurology and Neurosurgery, McGill University, Canada; Department of Psychology & York Biomedical Research Institute, University of York, UK

**Keywords:** semantic memory, social cognition, mentalizing, meta-analysis, default mode network

## Abstract

Key unanswered questions for cognitive neuroscience include whether social cognition is underpinned by specialised brain regions and to what extent it simultaneously depends on more domain-general systems. Until we glean a better understanding of the full set of contributions made by various systems, theories of social cognition will remain fundamentally limited. In the present study, we evaluate a recent proposal that semantic cognition plays a crucial role in supporting social cognition. While previous brain-based investigations have focused on dissociating these two systems, our primary aim was to assess the degree to which the neural correlates are overlapping, particularly within two key regions, the anterior temporal lobe (ATL) and the temporoparietal junction (TPJ). We focus on activation associated with theory of mind (ToM) and adopt a meta-analytic activation likelihood approach to synthesise a large set of functional neuroimaging studies and compare their results with studies of semantic cognition. As a key consideration, we sought to account for methodological differences across the two sets of studies, including the fact that ToM studies tend to use nonverbal stimuli while the semantics literature is dominated by language-based tasks. Overall, we observed consistent overlap between the two sets of brain regions, especially in the ATL and TPJ. This supports the claim that tasks involving ToM draw upon more general semantic retrieval processes. We also identified activation specific to ToM in the right TPJ, bilateral anterior mPFC, and right precuneus. This is consistent with the view that, nested amongst more domain-general systems, there is specialised circuitry that is tuned to social processes.

## 1. Introduction

The capacity to understand and respond appropriately to the thoughts and actions of others is of vital importance to our daily lives. When this ability breaks down, there are profound consequences for an individual’s ability to thrive in society (Frith, 2007; Frith & Frith, 2007). Therefore, a key challenge for neuroscience is to develop a full account of the cognitive and brain basis of social interaction.

The dominant mode within social neuroscience has been to seek out specialised neural subsystems dedicated to processing social (as opposed to more general kinds of) information (Apperly et al., 2005; Happé et al., 2017; Saxe & Powell, 2006; Spunt & Adolphs, 2017). This approach has uncovered evidence for the existence of category-sensitive cortex; regions that preferentially activate during the perception of certain social stimuli, such as faces (Kanwisher & Yovel, 2006), bodies (Downing & Kanwisher, 2010), and dyadic social interactions (Landsiedel et al., 2022). It has been argued that more complex inferential processes such as mental state attribution, or Theory of Mind, also engage highly specialised social brain areas (Apperly et al., 2005; Brüne & Brüne-Cohrs, 2006; Dodell-Feder et al., 2011; Gweon et al., 2012; Jacoby et al., 2016; Jenkins et al., 2014; Koster-Hale & Saxe, 2013; Richardson & Saxe, 2020; Ross & Olson, 2010; Saxe & Baron-Cohen, 2006; Saxe & Kanwisher, 2003b; Saxe & Wexler, 2005; Scholz et al., 2009; Simmons et al., 2010). However, the extent to which ‘higher-order’ systems (e.g., declarative memory; cognitive control) exhibit domain-specificity of this kind is hotly debated (e.g., Apperly et al., 2005; Binney & Ramsey, 2020; Ramsey & Ward, 2020). One factor keeping this debate from being resolved is that, to date, the role of domain-general systems in social cognition has received comparatively little attention and is not well understood. Consequently, neurobiological accounts of human social behaviour fall short of being comprehensive.

Recently, however, there has been increased interest in the involvement of a set of distributed domain-general networks in social processing. This includes the ‘multiple-demand network’ (MDN), a set of brain areas engaged by cognitively challenging tasks that span numerous cognitive domains (Assem et al., 2020; Duncan, 2010; Fedorenko et al., 2013; Hugdahl et al., 2015). MDN activity increases with working memory load and task switching demands, for example, and it has been suggested that this reflects the implementation of top-down attentional control to meet immediate task goals (Duncan, 2010, 2013). MDN regions have been implicated in social processes, including working memory for social content (Meyer et al., 2012), social conflict resolution (Zaki et al., 2010) and mental state attribution (e.g. Rothmayr et al., 2011; Samson et al., 2005; Van der Meer et al., 2011). A further set of co-activated brain regions, referred to collectively as the ‘default mode network’ (DMN), has also garnered a widely appreciated role in social cognition (Darda & Ramsey, 2019; Diveica et al., 2021; Zaki et al., 2010; Duncan, 2010; Fedorenko, 2014; Fedorenko et al., 2013; Hughes et al., 2019; Jackson et al., 2022; Mars et al., 2012; Schilbach et al., 2006; Spreng & Grady, 2010). The DMN is a large-scale functional network that tends to activate in the absence of an explicit task, and it has been proposed that it is ideally suited for supporting self-generated internally-orientated, as opposed to externally-orientated cognition (Margulies et al., 2016a; Smallwood et al., 2013). The DMN appears comprised of as many as three subsystems, and it is well accepted that at least one of these (which includes dorsomedial and ventrolateral prefrontal cortex, inferior parietal and lateral temporal regions) consistently activates during social processes like mental state attribution, which may in part relate to access to social knowledge (Spreng & Andrews-Hanna, 2015).

Indeed, it has recently been argued that a network known as the semantic cognition network (SCN; Humphreys et al., 2015; Jackson et al., 2019), has a crucial role in supporting social cognition (Balgova et al., 2022; Binney & Ramsey, 2020; Diveica et al., 2021). Semantic cognition (supported by the SCN) refers to neurocognitive systems involved in the acquisition and flexible retrieval of conceptual-level knowledge that exists to transform sensory inputs into meaningful, multimodal experiences. Conceptual knowledge critically underpins our capacity to recognise and interact with objects, words, people, and events in our environment (Patterson et al., 2007; Lambon Ralph et al., 2017), and Binney and Ramsey (2020) have argued that it should play a pivotal role in social cognition given that social interaction is, at its core, a process of *meaningful* exchange between persons. Support for this hypothesis has long existed within neuropsychological and comparative neuroscience literature, where there appears to be a tight coupling of general semantic deficits and social impairments (Bertoux et al., 2020; Irish et al., 2014; Klüver & Bucy, 1937; Miller et al., 2012; Souter et al., 2021; for a review see Olson et al., 2013 and Rouse et al., 2024). Evidence at the level of whole-brain networks has yet to be conclusively obtained.

The SCN is comprised of the IFG and posterolateral temporal cortex (inclusive of the posterior MTG and posterior ITG), which play a particular role in control-related processes, as well as the anterior temporal lobes (ATL) which underpin semantic representation processes (Jackson, 2021; Jefferies, 2013; Noonan et al., 2013; Lambon Ralph et al., 2017). There is a generally accepted notion that there is some degree of overlap between SCN regions and those brain regions involved in social cognition (Binney & Ramsey, 2020; Spreng & Andrews-Hanna, 2015), yet only very recently have there been direct explorations of this relationship. Moreover, most of these studies have focused on the differences, and divergence (Baetens et al., 2013; Hyatt et al., 2015), and relatively little discussion was given to the implications of any overlap. The matter of convergence has been elucidated in a series of recent targeted studies (Balgova et al., 2022; Binney & Ramsey, 2020; Diveica et al., 2021; Hodgson et al., 2022) although each of these are limited in particular ways. For instance, Balgova et al. (2022) was an fMRI study which employed a limited task set and thus could lack in generalisability.

Hodgson et al. (2002) and Diveica et al., (2021) used a meta-analytic approach to extract reliable findings from across large numbers of functional neuroimaging studies, and thereby circumvent the limitations of individual studies (Cumming, 2014; Eickhoff et al., 2012) which include low statistical power (Button et al., 2013) and vulnerability to idiosyncratic design/analysis choices (Botvinik-Nezer et al., 2020; Carp, 2012). Nonetheless, Hodgson et al. (2002) restricted their analyses to a limited brain volume. Diveica et al. (2021) on the other hand, while taking a whole brain approach, was focused on regions of the SCN that respond to increased cognitive control demands and they did not formally compare or contrast the whole SCN with activation maps from social cognitive tasks. These limitations (and others; see below) were addressed in the present study.

The present study addressed the overlap between brain networks involved in social and semantic cognition. It focused on one key aspect of social cognition, namely mental state attribution or ‘theory of mind’ (ToM), for three reasons. First, ToM is considered fundamental to successful social interactions (Apperly, 2012; Brüne & Brüne-Cohrs, 2006; Frith & Frith, 2005; Heleven & van Overwalle, 2018; van Hoeck et al., 2014). Second, there is a large body of literature, as is requisite for meta-analytic investigation. Third, ToM abilities enable one to describe, explain, predict, and infer the intentions, beliefs, and affective states of others (Adolphs, 2009; Brüne & Brüne-Cohrs, 2006; Frith & Frith, 2007, 2012; Frith & Frith, 2010; Happé et al., 2017; Premack & Woodruff, 1978). As such, ToM includes inferential processes that allow one to go beyond what is directly observable through the senses, thus appearing to be comparable to, and perhaps explained by, more general semantic processes that are specialized for the extraction of all types of meaning (Binney & Ramsey, 2020).

Most neural accounts of ToM implicate the temporoparietal junction (TPJ) alongside medial prefrontal cortex (mPFC) and the precuneus. Some accounts also include the posterior superior temporal sulcus (pSTS) and the ATL (Amodio & Frith, 2006; Mar, 2011; Molenberghs et al., 2016; Saxe & Kanwisher, 2003; Saxe & Wexler, 2005; Saxe, 2006; Saxe & Powell, 2006; Schurz et al., 2014, 2017). It is key to note that the term ‘TPJ’ is less frequently used in the semantic cognition literature than in social neuroscience, and the corresponding definition can be vague and heterogeneous. For present purposes, we interpret the label TPJ to refer to a large area that includes the posterolateral temporal cortex and the inferior parietal lobe, including the angular gyrus (AG) (Hodgson et al., 2022; Seghier, 2013, 2022; Seghier et al., 2010). Some accounts of semantic cognition include the AG and argue either that it is involved in the integration and storage of conceptual knowledge (Kuhnke et al., 2020), or as a temporary buffer (Humphreys & Tibon, 2022). However, the AG has also been attributed to other domain-general processes that extend beyond semantic processing (Cabeza et al., 2012; Geng & Vossel, 2013; Humphreys, Lambon Ralph, et al., 2021; Humphreys & Tibon, 2022). In the present study, we specifically anticipated overlap between the SCN and ToM network in the ATL and the TPJ as both regions are frequently implicated in putatively domain-specific social processes as well as semantic cognition (Balgova et al., 2022; Diveica et al., 2021; Humphreys, Lambon Ralph et al., 2021; Olson et al., 2013; Seghier et al., 2010).

We also aimed to investigate a potential hemispheric dissociation between social and semantic cognition at these sites. In semantic cognition, the role of the ATL is viewed as bilateral (albeit with a leftwards asymmetry when probed with verbal semantic information; Lambon Ralph et al., 2001; Rice, Hoffman, et al., 2015), whereas the role of the ATL in social cognition has been ascribed right lateralisation (Younes et al., 2022; Zahn et al., 2009). Evidence for this distinction is limited, however, because claims that the right, but not the left ATL, is key for social processing are based chiefly upon patient studies (Borghesani et al., 2019; Gainotti, 2015; Gorno-Tempini et al., 2003; Irish et al., 2014). Individual fMRI studies, on the other hand, typically indicate bilateral involvement or possibly a leftward asymmetry (Balgova et al., 2022; Binney, Hoffman, et al., 2016; Rice et al., 2018; Ross & Olson, 2010 but see Zahn et al., 2002; also see Arioli et al., 2021; Catricalà et al., 2020; Lin, Yang, et al., 2018; Pobric et al., 2016; Rice, Lambon Ralph, et al., 2015). The laterality of TPJ involvement in social cognition is unclear. In neuroimaging studies, it is often observed bilaterally (Molenberghs et al., 2016; Schurz et al., 2014), but selectivity of this region for ToM is argued to be limited to the right hemisphere by some authors (Perner et al., 2006; Saxe & Wexler, 2005) while others have reported greater selectivity in the left (Aichhorn et al., 2006, 2009). In semantic cognition, activation of regions within the TPJ tends to be left lateralised (Handjaras et al., 2017; Kuhnke et al., 2022; Seghier, 2013; Seghier et al., 2010). Collectively, these findings paint a complex picture regarding how the ToM and semantic networks converge and diverge at these ATL and TPJ sites.

Laterality differences may be of critical importance to differentiating semantic and social cognition networks. Alternatively, they could reflect a methodological confound which is that their typical neuroimaging assessments tend to use different types of stimuli and, for example, language-based tasks tend to drive greater left-hemisphere activation (Binder et al., 2009; Rice, Lambon Ralph, et al., 2015). A key aim of this study, therefore, was to investigate whether methodological factors give rise to a skewed pattern of activity in each domain. Most fMRI studies probing semantic cognition have used verbal stimuli (e.g., words/sentences) (Rice, Lambon Ralph, et al., 2015; Visser, Jefferies, et al., 2010). In contrast, nonverbal stimuli such as animations, vignettes, or free-viewing movie paradigms are popular in the ToM literature (Diveica et al., 2021; Molenberghs et al., 2016). Although both semantic cognition and ToM are typically viewed as modality-independent processes (Gallagher et al., 2000), these prevalent methodological differences could mar between-domain comparisons because activation patterns within each domain shift according to the stimulus presentation format. For example, a meta-analysis of fMRI studies found that non-verbal compared to verbal ToM tasks, evoke greater activation in the left precentral gyrus and left and right IFG, and lower activation in the mPFC, precuneus, and bilateral TPJ (Molenberghs et al., 2016). Similarly, Visser et al.’s (2010) meta-analysis of semantic cognition found that the laterality of ATL activation depends on whether stimuli were presented in the auditory versus visual modality (also see Krieger-Redwood et al., 2015; Rice, Lambon Ralph, et al., 2015). Thus, left unaccounted for, these kinds of systematic methodological differences could create the appearance of divergence between the two task-associated networks when there is, in fact, a common system with meaningful covariation driven by properties of the stimuli. In the present study, we controlled for stimulus format (verbal, non-verbal) and input modality (visual, auditory) to disentangle pervasive differences between networks from context dependent differences. In the same vein, we controlled for inter-domain differences in the types of baseline/control tasks used (e.g., active versus passive) and screened for the presence of social stimuli in the studies of semantics.

In summary, to determine the degree to which ToM and semantic cognition share an underlying neural basis, we performed a large-scale neuroimaging meta-analysis to systematically compare the ToM-related brain network with the SCN and with a primary focus on the ATL and TPJ. Moreover, we assessed the effect of stimulus format and sensory input modality on network overlap. To our knowledge, this is the first direct comparison of these two large-scale networks via these means (see Hodgson et al., 2022 for a region-specific analysis, and Diveica et al., 2021 for data specific to semantic control).

## 2. Materials and Methods

### 2.1. Literature selection and inclusion criteria

We leveraged a Theory of Mind (ToM) dataset curated by (Diveica et al., 2021), and a Semantic Cognition (SCN) dataset compiled by (Jackson, 2021). Both these studies performed a comprehensive and up-to-date literature review and followed best practice guidance for conducting meta-analyses (Müller et al., 2018). Below, we provide a brief description of each of these original datasets.

The general semantics analysis (257 studies, 415 contrasts, 3606 peaks) reported by Jackson (2021) was designed to capture all aspects of semantic cognition, including activation of conceptual level knowledge, as well as engagement of control processes that guide context- or task-appropriate retrieval of concepts. Studies were included if they compared a (more) semantic with a non- (or less-) semantic task or meaningful (or known) with meaningless (or unknown) stimuli. It included studies published between 2008 and 2019. The ToM analysis (136 experiments, 2158 peaks, 3452 participants) reported by Diveica et al. (2021) included studies published between 2014 and 2020 that employed a primary task involving inferences about the mental states of others, including their beliefs, intentions, and desires (but not sensory or emotional states). These studies were also required to compare the ToM task to a non-ToM task. Studies that looked at the passive observation of actions, social understanding, mimicry or imitation were not included unless the primary task included a clear ToM component. Studies investigating irony comprehension, those that employed trait inference tasks, and those that employed interactive games were also excluded. Both Jackson and Diveica et al. excluded contrasts that made comparisons between sub-components of the process of interest (but see the final paragraph in this Section). For example, Diveica et al. excluded affective ToM > cognitive ToM contrasts from the semantic cognition studies, and Jackson excluded abstract semantics > concrete semantics contrasts. This was critical for the present study because we were interested in common, core semantic/ToM processes that are subtracted out by these contrasts.

For these two datasets to be compared, it was essential to ensure that a similar, if not identical set of general exclusion criteria (i.e., those pertaining to the sample demographics, the imaging method, etc.) were applied. To this end, we initially planned to use the general inclusion/exclusion criteria described by Diveica et al. (2021) and reapply them to both the ToM and SCN datasets. In practice, we needed to implement a few minor modifications to these criteria. Below we summarise the final set of general criteria that we applied in the present study and highlight discrepancies from the approaches of Diveica et al. (2021) and Jackson (2021):

1. We included only peer-reviewed articles in English, and studies that employed task-based fMRI or PET, and only those that report whole-brain activation coordinates localised in one of two standardised stereotactic spaces (Talairach (TAL) or Montreal Neurological Institute (MNI)). Coordinates reported in TAL space were converted into MNI space using the Lancaster transform (tal2icbm transform (Lancaster et al., 2007) embedded within the GingerALE software version 3.0.2; http://brainmap.org/ale). Results from region-of-interest or small-volume correction analyses were excluded.
2. We included only studies that tested healthy adults to control for age-related changes in neural networks supporting cognition (e.g., see Hoffman & Morcom, 2018). A deviation from Diveica et al. (2021) was that we only considered studies reporting data from participants aged 18-40 years. If the age range of participants in a given study was not stated, we included the results in our datasets as long as the mean age of the participants was less than 40 years (if stated) and there was no clear indication that adults outside the range of 18-40 were included in the sample. This was a similar criterion to that used by Jackson (2021).
3. Diveica et al. (2021) included contrasts between the experimental task (i.e., ToM processing) and either an active control condition or rest/passive fixation. Jackson (2020) only included contrasts against active baselines. Therefore, we added additional contrasts involving rest/passive fixation into the SCN dataset. In the present study, active control conditions were characterised as either a high-level or low-level baseline; thus, over and above Diveica et al. (2021) and Jackson (2020), the present study differentiated low-level active baselines (e.g., visual stimulation with a string of hashmarks as a control for sentence reading) from rest/passive fixation. With these extra steps, we aimed to better account for methodological differences across domains (see more detail in **Section 2.3.1**).
4. Where present, multiple contrasts from the same group of participants were included if they met all the other inclusion criteria. We controlled for within-group effects by pooling contrasts into a single experiment (Müller et al., 2018; Turkeltaub et al., 2012) like Diveica et al. (2021) and Jackson (2020). This means that, when we refer to the numbers of experiments that constituted the units of input, we have counted contrasts from a single participant sample as one single experiment. In follow-up contrast analyses that compared different conditions (e.g., stimulus format or input modality), initially pooled contrasts related to these different conditions were separated (see more detail in **Section 2.2**). While Diveica and colleagues excluded the contrast with a smaller number of peaks after separating, we retained both of these contrasts to maximise the use of all available data.

Two further adjustments were made to the SCN dataset to make it optimally comparable to the ToM dataset. As discussed above, both Jackson and Diveica et al. excluded contrasts that made comparisons between sub-components of the process of interest and thus could subtract away core processes associated with ToM and semantic cognition. In the case of ToM, this left only those contrasts comparing ToM tasks with non-ToM tasks. Jackson, however, also included a small number of contrasts that compared more semantic tasks with less semantic tasks (e.g., an identity classification task using faces with varying degrees of familiarity used by Rotshtein et al., (2005) or a task contrasting personal familiar and famous familiar faces used by Sugiura et al., (2006). In the present study, we excluded these because they could subtract out some core processes or common regions. While this was likely of little consequence in Jackson’s (2021) study, the inclusion of these contrasts could, in principle, weaken the comparison of SCN data with the ToM data. An exception was applied to contrasts that pitted intelligible sentences against scrambled sentences because they were an important source of data in the verbal and auditory domain, and we reasoned that, while there is meaning present in both stimuli types at the single word level, the critical difference was meaningfulness at the sentence level. Finally, we identified and excluded a small number of experiments in Jackson’s SCN dataset (n=4) that used contrasts that could be viewed as probing ToM-related processing.

The final ToM dataset used in the present study comprised 114 experiments from 2800 participants, 159 contrasts, and 1893 peaks. The final SCN dataset used in the present study comprised 214 experiments, including data from 3934 participants, 410 contrasts, and 3803 peaks.

### 2.2. Categorising Contrasts by Stimulus Format and Sensory Input Modality

In line with our secondary aim of accounting for the effects of stimulus format and sensory input modality on network overlap, individual contrasts from both the ToM and SCN datasets were further categorised as being within the verbal domain or the non-verbal domain. Verbal paradigms used spoken or written language stimuli. Examples of non-verbal paradigms include those using pictures (e.g., of objects or actions), animations, videos, or environmental sounds (see Rice et al., 2015 for a similar approach). Moreover, contrasts were independently categorised according to whether stimuli were presented in the visual or auditory modality. In this case, pictorial stimuli, as well as written words and sentences were counted as visual stimuli (see Molenberghs et al., 2016; Visser et al., 2010 for similar approaches). In cases where both types of stimuli (e.g., verbal and non-verbal) were used in the same task, the contrast was excluded (e.g. Sommer et al., (2010).The reader is referred to **Table 1** and the **Supplementary Information** for the number of studies and a list of excluded contrasts in each of these categories.

**Table 1.** The number of experiments, contrasts, and peaks split according to the stimulus format, input modality, type of baseline, and presence of social content.

| Experiment Type | Theory of Mind |  |  |  | Semantic Cognition |  |  |  |
| --- | --- | --- | --- | --- | --- | --- | --- | --- |
|  | Peaks | Sample Size | Experiments | Contrasts (%) | Peaks | Sample Size | Experiments | Contrasts (%) |
| **Total** | **3803** | **2800** | **114** | **159 (100%)** | **1893** | **3934** | **214** | **410 (100%)** |
| **Baseline** |  |  |  |  |  |  |  |  |
| High | 2387 | 2749 | 111 | 151 (95%) | 1800 | 3064 | 170 | 283 (69%) |
| Low | 1026 | 33 | 2 | 2 (1%) | 19 | 880 | 47 | 88 (21%) |
| Rest | 387 | 98 | 5 | 6 (4%) | 74 | 388 | 24 | 38 (9%) |
| Both | 3 | NA | NA | NA | NA | 12 | 1 | 1 (0.2%) |
| **Social Content\*** |  |  |  |  |  |  |  |  |
| No | NA | NA | NA | NA | 2881 | 3610 | 193 | 323 (79%) |
| Yes | NA | NA | NA | NA | 527 | 323 | 20 | 48 (12%) |
| Mixed | NA | NA | NA | NA | 8 | 10 | 1 | 1 (0.2%) |
| **Stimulus\* format** |  |  |  |  |  |  |  |  |
| Verbal | 2684 | 988 | 46 | 59 (37%) | 542 | 3261 | 175 | 296 (72%) |
| Non-Verbal | 525 | 1858 | 71 | 89 (56%) | 1244 | 701 | 37 | 57 (14%) |
| Both/None | 207 | 53 | 4 | 5 (3.1%) | 33 | 341 | 15 | 19 (5%) |
| **Input Modality\*** |  |  |  |  |  |  |  |  |
| Visual | 2707 | 2664 | 106 | 142 (89%) | 1715 | 2864 | 152 | 285 (70%) |
| Auditory | 592 | 91 | 6 | 7 (4%) | 64 | 1023 | 60 | 76 (19%) |
| Both/None | 117 | 45 | 2 | 4 (3 %) | 40 | 180 | 8 | 11 (3%) |

\*The experiment counts exclude experiments using rest as the baseline conditions. See text for further details. Contrast % rounded up to the nearest 0.1%.

### 2.3. Further Methodological Considerations

Following the application of general inclusion/exclusion criteria and the categorization described in **Section 2.2**, we took additional steps to further characterize the two revised datasets and evaluate the potential for other confounds to influence their comparison. As we shall describe below, this led to further refinement which improved the suitability of the datasets for addressing our key research questions.

#### 2.3.1. Controlling for type of baseline

In semantic cognition research, it is widely accepted that the results of neuroimaging studies are affected, in important ways, by the choice of baseline task; a failure to perform adequate matching of baselines to experimental conditions in terms of perceptual input, response and attentional/executive demands, decreases sensitivity of subtractive designs to activation in brain areas associated with cross-modal integration, semantic processing and response selection (Price et al., 2005). Indeed, the use of passive rest or simple fixation as a baseline results in failure to reveal task-positive activation in anterior temporal areas (Binder et al., 2009; Price et al., 2005; Visser, Jefferies, et al., 2010), because minimal baseline task demands increase the opportunity for spontaneous semantic processing (associated with daydreaming and inner speech) to occur at an equal or greater depth/magnitude than that associated with more focused task-related semantic processing (Andrews-Hanna et al., 2014; Binder et al., 2009, 2016; Chiou et al., 2020; Humphreys et al., 2015; Visser, Jefferies, et al., 2010). While it is not typically discussed in the literature, this is also an important consideration for neuroimaging studies of social cognition because various forms of social inference are likely to occur during a state of mind-wandering (see, e.g., Diaz et al., 2013).

We observed that our SCN and ToM datasets differed considerably in the types of baselines used and that there was a higher degree of variability among semantic cognition studies (see **Table 1** and the **Supplementary Information**). This could have led to a confound in the inter-domain comparisons, namely a difference in the sensitivity to activation associated with cross-modal processing. To explore these issues, we (a) quantified these differences using three categories of baseline and (b) mapped the effect of including/excluding contrasts that used these baselines on the outcomes of ALE analysis within each domain. The results of this preliminary analysis informed our final approach to defining the datasets used for the inter-domain comparisons (see below). Previous attempts to deal with this issue have only distinguished between two types of baselines (e.g., Visser et al., 2010), but with a view to capturing greater specificity in these effects, we operationalized three, as follows:

1. High-level baselines were defined as those including an active task designed to approximate the demands of the main/experimental task without engaging the process of interest (ToM or semantic processing). This includes being generally well-matched to the experimental task in terms of perceptual (visual, auditory) properties, and means of behavioural output (overt/covert).
2. Low-level baselines were defined as having a task that required active engagement but one that differed from the main task in numerous ways, including perceptual properties, means of behavioural output, or difficulty.
3. Finally, the third category of baselines were those which required only passively watching a blank screen or maintaining visual fixation.

Our chief motivation for this finer differentiation of baseline types was to arrive at an optimal scenario in which we could remove cross-domain confounds while retaining as many data points, and therefore as much power, as possible. We decided on a stepwise approach in which we would compute the ALE map for each domain (i) with all contrasts included, then (ii) without contrasts involving rest/fixation, and finally, (iii) with neither the rest/fixation nor low-level baseline contrasts included. We visually compared the ALE maps generated at each step, as well as the associated output tables, paying attention to the gain or loss of suprathreshold clusters. We decided a priori that if inclusion/exclusion resulted in minimal change to the activation maps, then we would opt to retain contrasts in the sample. The authors acknowledge the arbitrariness of this criterion, but setting a more specific rule a priori was impractical due to the fact that the quality and quantity of these changes were likely to vary across different comparisons (due to sample size, etc). Therefore, to mitigate against this and ensure transparency, we (i) opted to fully report the results both prior to and following exclusions, and (ii) ensure all datasets and subsets were publicly available so that the community can reproduce our results and explore the consequences of certain methodological decisions (all data can be found at https://osf.io/ydnxh/).

We found that, in the case of the SCN data set, excluding passive/resting baselines resulted in additional activation in the left inferior temporal lobe and in right medial temporal areas (see **Supplementary Figure R2b and Supplementary Table R2**). The exclusion of contrasts utilising low-level baselines did not lead to any appreciable differences in the distribution of activations, but the size of clusters was reduced owing to the reduced sample and power. In the case of the ToM data, the impact of these exclusions was negligible due to a very low number of experiments with low-level and passive baselines (**Supplementary Figure R2a and Supplementary Table R1**). Overall, these outcomes are consistent with an expectation that the inclusion of passive baselines would occlude activation within parts of the SCN (Binder et al., 2009, 2016; Humphreys et al., 2015; Visser, Jefferies, et al., 2010). Exclusion of lower-level baselines, on the other hand, might be an overly conservative approach that prohibits the detection of activation that is common across domains. We, therefore, excluded only contrasts involving rest/passive baselines and from both datasets.

#### 2.3.2. Controlling for the ‘socialness’ of semantic stimuli

20 studies (48 contrasts) in Jackson’s (2021) original SCN data set, having otherwise met our revised exclusion/inclusion criteria, involved a task or stimuli that were, to some degree, social in nature. For example, some studies used social or emotion concepts, and others probed person knowledge through famous faces (e.g., Elfgren et al., 2006; Grabowski et al., 2001; Leveroni et al., 2000). We identified contrasts as being social if they used stimuli that consistently referred to (i.e., this was a defining feature for the stimuli) social characteristics of persons or group of people, a social behaviour or interaction, or any other socially-relevant concept (Diveica, Pexman & Binney, 2021). These studies required further consideration, particularly because of an ongoing debate concerning whether social semantics and general semantics depend upon independent or overlapping representational systems (Arioli et al., 2021; Binney et al., 2016; Binney & Ramsey, 2020; Olson et al., 2013; Pexman et al., 2023). It is possible that ToM tasks engage social concepts and therefore the same regions engaged by social semantic processing (e.g., the dorsal ATL; Binney & Ramsey, 2020; Ross & Olson, 2010; Zahn et al., 2007) without relying on general semantic areas. In this case, if we were to pool social semantic contrasts and general semantic contrasts, then we might obtain an exaggerated picture of the extent to which the ToM network overlaps with the general processing semantic network. However, there was also a pragmatic reason for including these studies: they are a key source of data related to non-verbal semantic processing (see **Table 1**) and excluding them could compromise our ability to remove the confounding effect of stimuli type. To account for this, we examined the effect of including/excluding these studies in our general semantic dataset. These results are fully reported in the **Supplementary Information No 2.** Briefly, the overall pattern remained almost the same when the social contrasts were excluded, apart from losing a small cluster in the right IFG and slightly less extensive left temporopolar activation. These differences are likely to be due to the reduction in the number of studies included and concern brain regions that were not the focus of the present study, and thus are not central to the conclusions made. Therefore, we decided to retain the social contrasts as part of our SCN dataset and include them in the cross-domain comparisons reported in the **Results** section.

### 2.3. Data Analysis

We performed coordinate-based meta-analyses, using the revised activation likelihood estimation (ALE) algorithm as implemented in the GingerALE 3.02 software (http://brainmap.org/ale) (Eickhoff et al., 2009, 2012, 2017; Laird et al., 2005). To ensure sufficient statistical power, analyses were only performed on samples comprising a minimum of 17 experiments (Eickhoff et al., 2016). Nonetheless, meta-analyses performed on small sample sizes are susceptible to potential publication bias, and caution should be given to interpretation of results from samples with less than 30 studies (Acar et al., 2018). The minimum sample size in this report, however, is n= 37. Each analysis was comprised of two stages. The first stage consisted of independent analyses of the ToM and SCN datasets, which were used to identify areas of consistent activation within each domain. Here, the ALE meta-analytic method treats the activation coordinates reported by each experiment as the center points of three-dimensional Gaussian probability distributions which differ in width to account for the reliability of the peak estimate based on the size of the participant sample (Eickhoff et al., 2009). These spatial probability distributions are aggregated, creating a voxel-wise modelled activation (MA) map for each experiment in the sample. Then, the voxel-wise union across the MA maps of all experiments is computed, resulting in an ALE map that quantifies the convergence of results across experiments (Turkeltaub et al., 2012). GingerALE tests for above-chance convergence (Eickhoff et al., 2012), thus permitting random-effects inferences. Following the recommendations of Müller et al. (2018), ALE maps of both the ToM and SCN domains were thresholded using cluster-level family-wise error (FWE) correction of *p* < .05 with a prior cluster-forming threshold of *p* < .001 (uncorrected), which was estimated via 5000 permutations. Cluster-level FWE correction has been shown to offer the best compromise between sensitivity to detect true convergence and spatial specificity (Eickhoff et al., 2016).

The ALE maps generated in this first stage were used as inputs for the second stage of analysis, comprised of conjunction and contrast analyses. These analyses were aimed at identifying similarities and differences, respectively, in neural activation between the SCN and ToM sets of studies. Conjunction images were generated using the voxel-wise minimum value of the ALE maps (Nichols et al., 2005). Contrast images were created by directly subtracting one ALE map from the other (Eickhoff et al., 2011). Differences in ALE scores were compared to a null distribution that was estimated via a permutation approach with 5000 repetitions. Given that there are no established methods for multiple comparison correction of ALE contrast maps (see Eickhoff et al., 2011), the contrast maps were thresholded using a more conservative threshold of p < 0.001 (uncorrected) and a minimum cluster size of 100 mm3. Moreover, we masked the contrast maps with the cluster-level FWE-corrected ALE maps resulting from the independent ALE analysis of the respective cognitive domain. Thresholded ALE maps were plotted on a MNI152 template brain using MRICroGL (https://www.nitrc.org/projects/mricrogl). We used FSL maths commands and FSL VIEW (https://www.nitrc.org/projects/fsl) to binarise the ALE maps for better visual clarity when displaying the conjunction.

In a final step, we conducted post hoc cluster analyses that afforded a complementary approach to evaluating whether clusters of activation identified in the two independent ALE analyses of the SCN and ToM data were driven by certain methodological characteristics (i.e., input modality and stimulus format). We examined the list of experiments that contributed to each cluster by at least one peak and computed the likelihood of contribution of a given experiment type. For these purposes, we used Fisher’s exact tests of independence and post-hoc pairwise comparisons in R studio Version 1.4.1106 (https://www.rstudio.com).

In summary, our analysis pipeline proceeded as follows. To address our primary question about similarities in the brain networks underpinning semantic and social cognition, we conducted independent ALE analyses on the ToM and SCN datasets which generated whole-brain activation maps. These maps were then used to create conjunction and contrast analyses aimed at identifying overlap and differences in the topology of activation between the two domains. We repeated these analyses having divided the SCN and ToM datasets into subsets containing experiments that used VERBAL stimuli on one hand, and NON-VERBAL stimuli on the other. This allowed examination of the effect of stimulus format. Then we split the datasets into subsets containing experiments that used VISUAL and AUDITORY stimuli and repeated the analyses to investigate the impact of sensory input modality. Finally, we performed cluster analyses to check whether the likelihood of finding activation within each cluster identified in the primary ALE analyses of the ToM and SCN data depends on experiment type (VERBAL, NON-VERBAL, VISUAL, AUDITORY).

## 3. Results

### 3.1. General Overlap Between Networks Subserving Theory of Mind and Semantic Cognition

Our principal analyses explored the extent to which neural networks engaged by ToM and semantic cognition tasks overlap (and diverge). Overall, the results reveal extensive areas of overlap including at key areas of interest (see **Figure 1** and **Table 2**; also see the independent ALE analysis results for each separate domain in **Supplementary Information No. 2: Supplementary Figure R1** and **Supplementary Table R1**). Specifically, there was a conjunction of ToM and SCN activity within the bilateral ATL that covered the temporal pole (TP) and the banks of the anterior STS, the MTG and STG in both hemispheres. In the left but not the right hemisphere, the area of overlap extended along the whole length of the MTG/STG towards the lateral temporoparietal junction (including the AG) as well as medial portions of the IPL. There was also a conjunction of activation in the left posterior ventral temporal lobe (ITG/FG), and in the lateral frontal cortex including pars orbitalis, triangularis and opercularis of the left IFG and the ventral precentral gyrus. There were smaller clusters on the bank of the right inferior frontal sulcus (pars triangularis), the left dorsomedial frontal cortex and left inferior precuneus.

**Figure 1.**
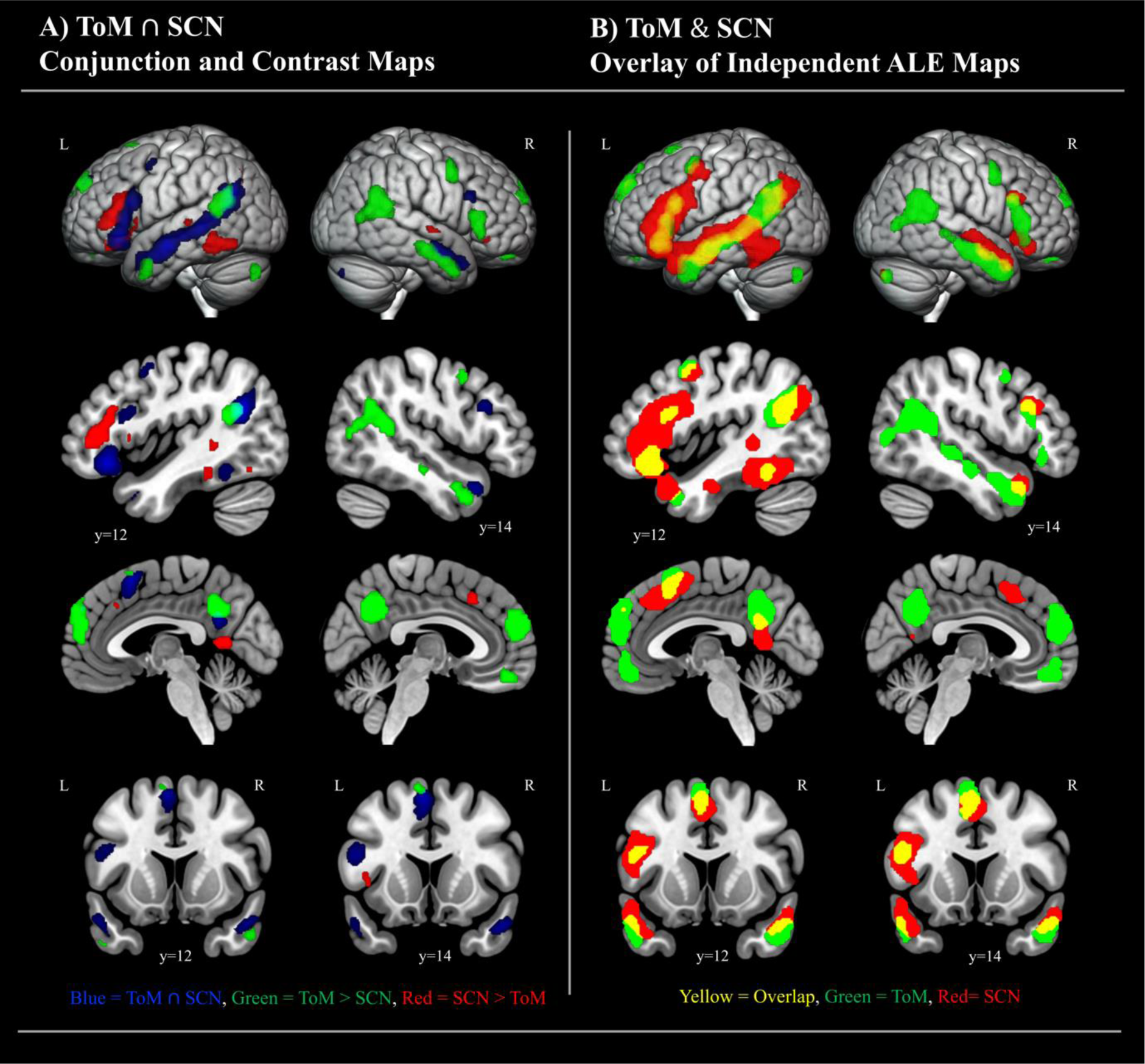
Common and differential activation for ToM (N=113) and SCN (N= 211). **Panel A** displays the conjunction alongside statistically significant differences revealed by the contrast analyses. The contrast maps in **Panel A** were thresholded with a cluster forming threshold at p<.001 and a minimum cluster size of 100mm^3^. In **Panel B,** we have overlaid the binarised versions of the complete ALE maps resulting from independent analysis of ToM and semantic cognition studies. This allows for full visualisation of the topography of the two networks and consideration of the relationship between them (also see Supplementary Figure R1 and Supplementary Table R1). The independent ALE maps were treated to a cluster-forming threshold at p<.001, and an FWE-corrected cluster-extent threshold at p<.05. The sagittal and coronal sections are chosen as representative slices positioned over peak coordinates at which there is the greatest conjunction in the bilateral anterior temporal lobes (left y= 12; right y= 14).

**Table 2.**
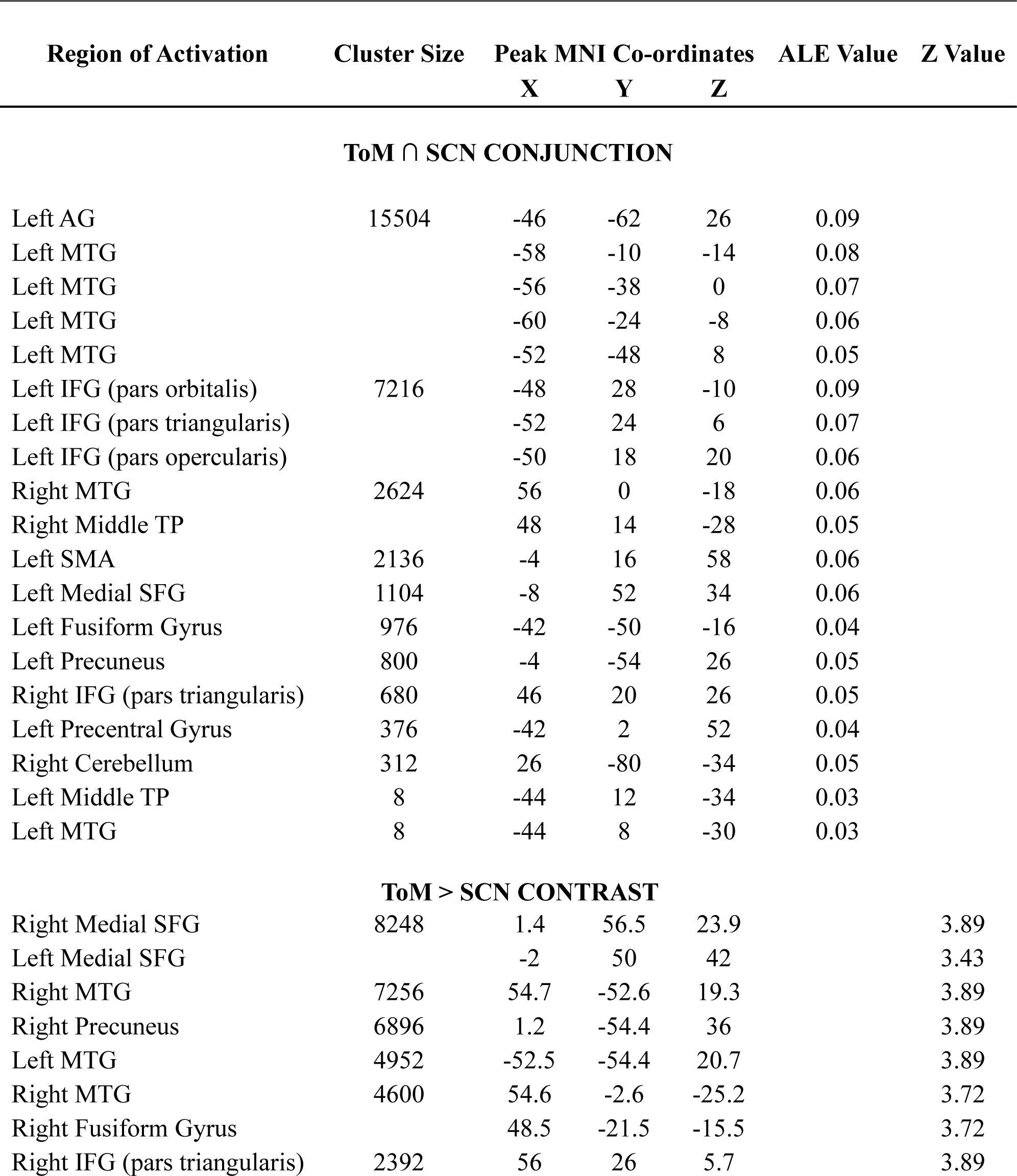
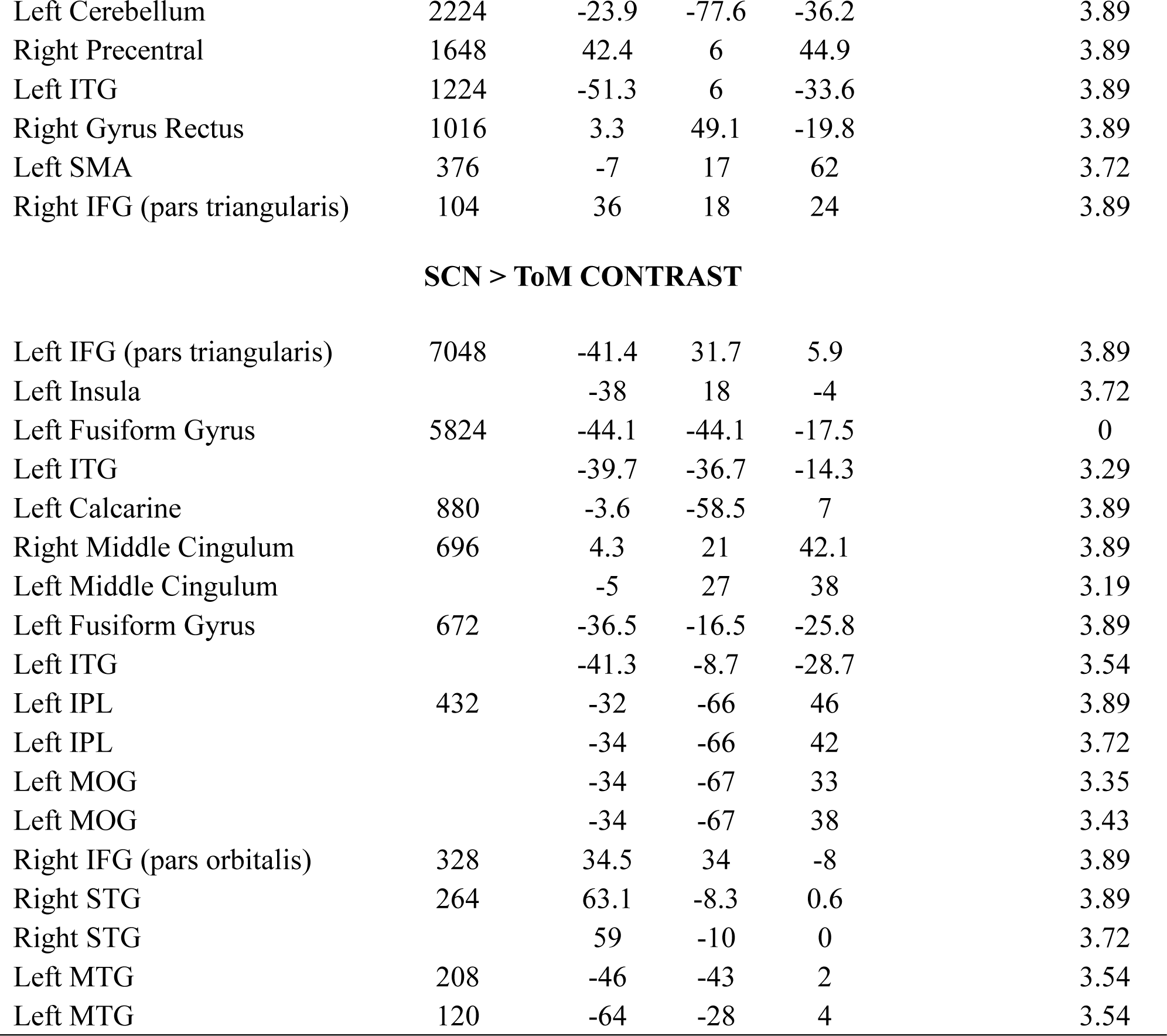
Conjunction and contrast analyses of the ToM (N= 113) and SCN (N= 211) experiments.

| Region of Activation | Cluster Size | Peak MNI Co-ordinates |  |  | ALE Value | Z Value |
| --- | --- | --- | --- | --- | --- | --- |
|  |  | X | Y | Z |  |  |
| **ToM ∩ SCN CONJUNCTION** |  |  |  |  |  |  |
| Left AG | 15504 | -46 | -62 | 26 | 0.09 |  |
| Left MTG |  | -58 | -10 | -14 | 0.08 |  |
| Left MTG |  | -56 | -38 | 0 | 0.07 |  |
| Left MTG |  | -60 | -24 | -8 | 0.06 |  |
| Left MTG |  | -52 | -48 | 8 | 0.05 |  |
| Left IFG (pars orbitalis) | 7216 | -48 | 28 | -10 | 0.09 |  |
| Left IFG (pars triangularis) |  | -52 | 24 | 6 | 0.07 |  |
| Left IFG (pars opercularis) |  | -50 | 18 | 20 | 0.06 |  |
| Right MTG | 2624 | 56 | 0 | -18 | 0.06 |  |
| Right Middle TP |  | 48 | 14 | -28 | 0.05 |  |
| Left SMA | 2136 | -4 | 16 | 58 | 0.06 |  |
| Left Medial SFG | 1104 | -8 | 52 | 34 | 0.06 |  |
| Left Fusiform Gyrus | 976 | -42 | -50 | -16 | 0.04 |  |
| Left Precuneus | 800 | -4 | -54 | 26 | 0.05 |  |
| Right IFG (pars triangularis) | 680 | 46 | 20 | 26 | 0.05 |  |
| Left Precentral Gyrus | 376 | -42 | 2 | 52 | 0.04 |  |
| Right Cerebellum | 312 | 26 | -80 | -34 | 0.05 |  |
| Left Middle TP | 8 | -44 | 12 | -34 | 0.03 |  |
| Left MTG | 8 | -44 | 8 | -30 | 0.03 |  |
| **ToM > SCN CONTRAST** |  |  |  |  |  |  |
| Right Medial SFG | 8248 | 1.4 | 56.5 | 23.9 |  | 3.89 |
| Left Medial SFG |  | -2 | 50 | 42 |  | 3.43 |
| Right MTG | 7256 | 54.7 | -52.6 | 19.3 |  | 3.89 |
| Right Precuneus | 6896 | 1.2 | -54.4 | 36 |  | 3.89 |
| Left MTG | 4952 | -52.5 | -54.4 | 20.7 |  | 3.89 |
| Right MTG | 4600 | 54.6 | -2.6 | -25.2 |  | 3.72 |
| Right Fusiform Gyrus |  | 48.5 | -21.5 | -15.5 |  | 3.72 |
| Right IFG (pars triangularis) | 2392 | 56 | 26 | 5.7 |  | 3.89 |

|  |  |  |  |  |  |
| --- | --- | --- | --- | --- | --- |
| Left Cerebellum | 2224 | -23.9 | -77.6 | -36.2 | 3.89 |
| Right Precentral | 1648 | 42.4 | 6 | 44.9 | 3.89 |
| Left ITG | 1224 | -51.3 | 6 | -33.6 | 3.89 |
| Right Gyrus Rectus | 1016 | 3.3 | 49.1 | -19.8 | 3.89 |
| Left SMA | 376 | -7 | 17 | 62 | 3.72 |
| Right IFG (pars triangularis) | 104 | 36 | 18 | 24 | 3.89 |

**SCN > ToM CONTRAST**
|  |  |  |  |  |  |
| --- | --- | --- | --- | --- | --- |
| Left IFG (pars triangularis) | 7048 | -41.4 | 31.7 | 5.9 | 3.89 |
| Left Insula |  | -38 | 18 | -4 | 3.72 |
| Left Fusiform Gyrus | 5824 | -44.1 | -44.1 | -17.5 | 0 |
| Left ITG |  | -39.7 | -36.7 | -14.3 | 3.29 |
| Left Calcarine | 880 | -3.6 | -58.5 | 7 | 3.89 |
| Right Middle Cingulum | 696 | 4.3 | 21 | 42.1 | 3.89 |
| Left Middle Cingulum |  | -5 | 27 | 38 | 3.19 |
| Left Fusiform Gyrus | 672 | -36.5 | -16.5 | -25.8 | 3.89 |
| Left ITG |  | -41.3 | -8.7 | -28.7 | 3.54 |
| Left IPL | 432 | -32 | -66 | 46 | 3.89 |
| Left IPL |  | -34 | -66 | 42 | 3.72 |
| Left MOG |  | -34 | -67 | 33 | 3.35 |
| Left MOG |  | -34 | -67 | 38 | 3.43 |
| Right IFG (pars orbitalis) | 328 | 34.5 | 34 | -8 | 3.89 |
| Right STG | 264 | 63.1 | -8.3 | 0.6 | 3.89 |
| Right STG |  | 59 | -10 | 0 | 3.72 |
| Left MTG | 208 | -46 | -43 | 2 | 3.54 |
| Left MTG | 120 | -64 | -28 | 4 | 3.54 |

In the context of this large overlap, the contrast analyses revealed key differences between ToM and SCN (**Figure 1**). On the lateral surface of the bilateral ATL the activation for ToM included an area of anterior MTG that the SCN did not. Moreover, in the right IPL/AG (within the TPJ), activation was only consistently identified for ToM. While both ToM and semantic cognition elicit reliable activation in the left TP, as well as the IPL/AG (TPJ), the contrast analyses revealed that voxels in this same areas had significantly higher ALE values for ToM compared to the SCN. Beyond our key areas of interest, compared to the semantic studies, ToM studies also showed higher convergence of activation in the right IFG, right precentral gyrus, bilateral anterior mPFC, left precuneus and left cerebellum. On the other hand, SCN experiments also showed increased convergence of activation in the ventral portion of the left pMTG stretching to the posterior ITG and FG, in the left MFG/IFG spreading towards the insula, and in the left inferior precuneus and right dorsal mPFC.

### 3.2. The Role of Stimulus Format (VERBAL versus NON-VERBAL)

In this next set of analyses, we explored the extent to which differences between the activation maps associated with ToM and semantic cognition could be explained by systematic differences in the types of tasks and stimuli used in each domain. We repeated the above comparisons, this time excluding contrasts involving nonverbal stimuli (i.e., only retaining those involving verbal stimuli). Both samples were large enough for the purposes of meta-analysis although there were many more experiments using verbal stimuli in the domain of semantic cognition than there were in the ToM dataset (VERBAL ToM: n= 46; VERBAL SCN: n= 175). Nonetheless, this analysis revealed a very similar pattern of conjunction to the principal set of comparisons reported in **Section 3.1** including the bilateral ATL (anterior MTG/STS) and the left TPJ (See **Figure 2** and **Supplementary Information No. 2: Supplementary Table R4**). However, there ceased to be any IFG activation for ToM tasks, and thus overlap between the two domains was absent in this region. A similar observation was made at the left posterior STS/MTG, and other small clusters of conjunction were no longer present. This could simply be due to the substantial reduction in size of the ToM experiment sample (from 113 to 46). However, we looked at the cluster analyses for the IFG and found that verbal ToM experiments were significantly less likely than nonverbal ToM experiments to contribute to the clusters in the bilateral IFG (see **Supplementary Information No.2: Supplementary Figure CA1** and **Supplementary Table CA1** for more detail). This is contrary to expectations given that the left IFG is strongly engaged in language processing (Friederici, 2011). It is, however, consistent with prior results from the false belief > false photograph contrast employed in many of these verbal ToM studies (see Diveica et al., 2021; Schurz et al., 2014), and it is possible this result reflects the large number of these contrasts included in this meta-analysis. One explanation for this observation could be differences between the ToM tasks (e.g., false belief) and the control/baseline (e.g., false photograph) task in terms of the semantic/syntactic operations that need to be performed. Should there be greater or equivalent difficulty in the control task, then IFG activation could be subtracted away (see Diveica et al., 2021 for analyses that address this issue).

**Figure 2.**
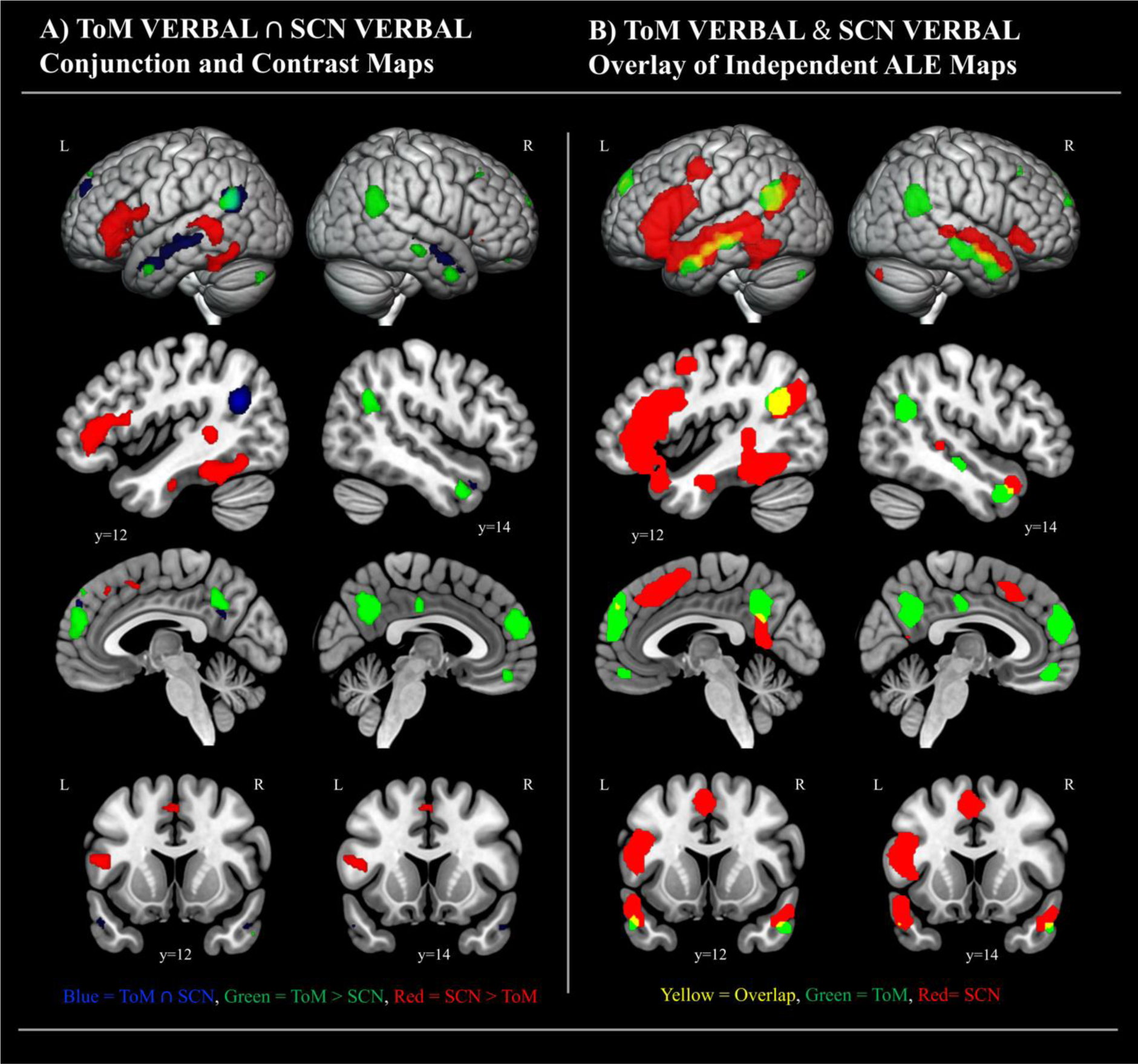
Common and differential activation for VERBAL ToM (N=46) and VERBAL SCN (N= 175). The initial ALE maps were treated to a cluster-forming threshold at p<.001, and an FWE-corrected cluster-extent threshold at p<.05 prior to the conjunction and contrast analyses. The contrast maps in **Panel A** were additionally thresholded with a cluster-forming threshold at p<.001 and a minimum cluster size of 100mm^3^. **Panel A** displays the conjunction alongside side statistically significant differences. In **Panel B,** we have overlaid the binarised versions of the complete ALE maps resulting from independent analysis of VERBAL ToM and VERBAL SCN studies. This allows for full visualisation of the topography of the associated networks (also see Supplementary Figures R4 & R5 and Supplementary Table R6 & R7). The sagittal and coronal sections are chosen as representative slices positioned over peak coordinates at which there is the greatest conjunction in the bilateral anterior temporal lobes (left y= 12; right y= 14).

In the corresponding contrast analyses, the differences between ToM and the SCN in the bilateral ATL and left TPJ were less pronounced, yet they remained. There also continued to be more consistent activation of the right TPJ for ToM. This was also true of the left anterior mPFC, and left precuneus. Indeed, while the extent of the clusters changed because of the reduced sample size in the ToM dataset, we continued to find more consistent involvement of the right TPJ, mPFC and left precuneus in ToM as compared to semantic cognition. Given that there were less studies in the ToM than SCN dataset, it is unlikely that these cross-domain differences could be attributed to lower statistical power in the case of SCN. For more detail see **Supplementary Information No. 2: Supplementary Figures R4 & R5 Panel A** and **Supplementary Table R6 & R7**).

When we limited the datasets to experiments utilising nonverbal stimuli, the results of the ALE analysis for ToM remained mostly unchanged from that seen in **Section 3.1**. In the case of semantic cognition, the number and extent of clusters was greatly diminished which reflects the reduced sample size (see **Figure 3**). Indeed, there were more experiments using nonverbal stimuli in the domain of ToM than there were exploring semantic cognition (ToM: n= 71; SCN: n= 37) and, as a consequence of the reduced sample size in the SCN domain and consequent lack of initial convergent ALE activation, there was no conjunction between the domains in the left TPJ. Overlap was still present in key regions of interest including the left pMTG, left ITG and some small aspects of the left IFG. Notably, even though visual inspection of the independent ALE maps for each domain suggests a large difference in terms of bilateral ATL activation, there were no significant differences revealed by the contrast analysis. The bilateral TPJ responded selectively to ToM in this analysis, while the posterior ITG was only present in the SCN. The ALE maps for each domain can be found in **Supplementary Information No. 2: Supplementary Figures R4 & R5 Panel B and Supplementary Tables R6 & R7**. In the cluster analysis of either the ToM or semantic domain, we found that the likelihood of finding activation in the respective ATL or TPJ areas did not depend on the verbal/non-verbal nature of the stimuli. This finding suggests that the inability to identify convergent left TPJ activation in the non-verbal SCN sample, and, consequently, overlap with non-verbal ToM, is indeed due to reduced statistical power. The cluster analysis showed that non-verbal experiments did however contribute more to the bilateral IFG and SFG in the ToM domain (See more detail in **Supplementary Information No.2: Supplementary Figure CA1** and **Supplementary Table CA1**).

**Figure 3.**
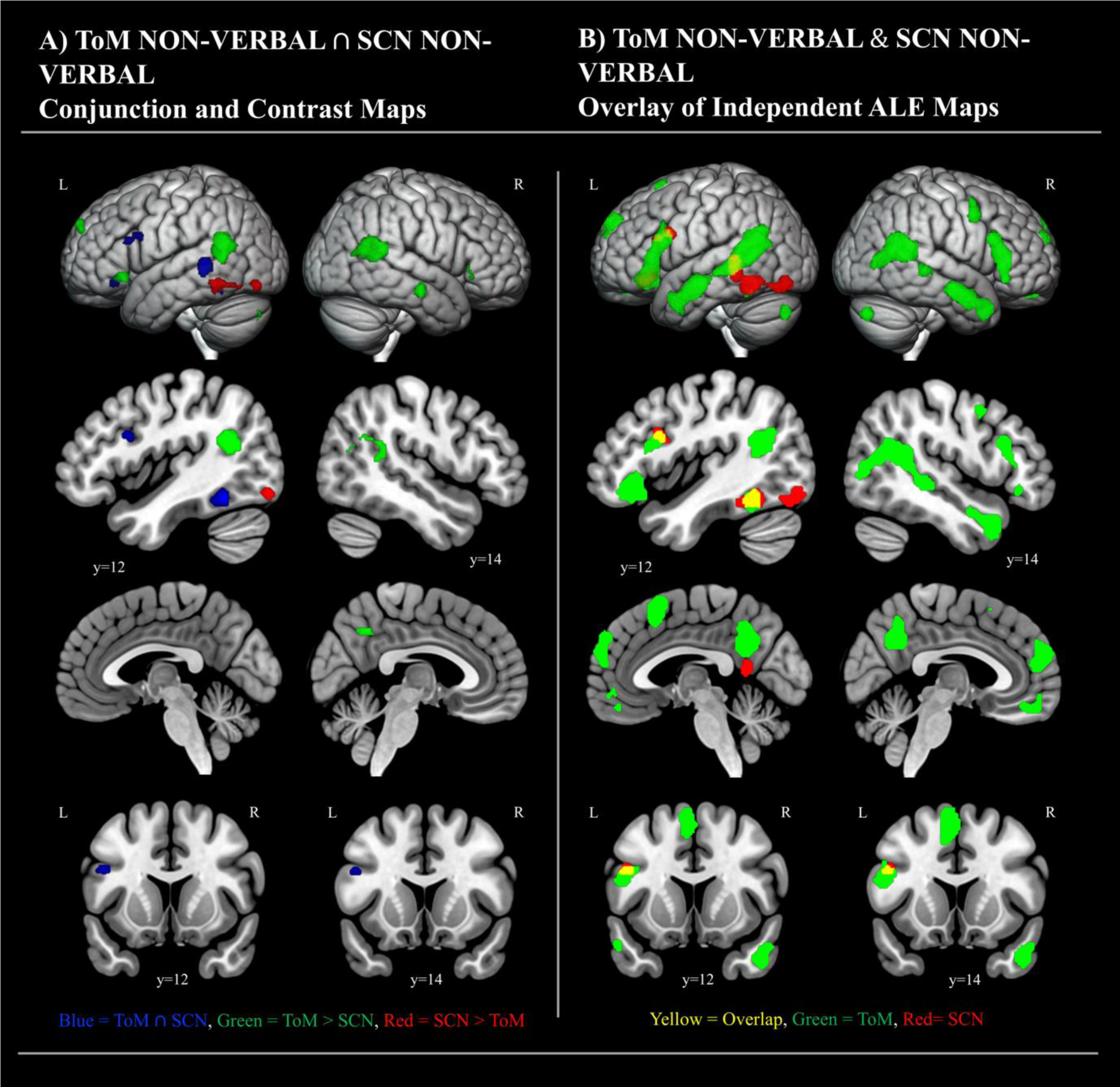
Common and differential activation for NON-VERBAL ToM (N=71) and NON-VERBAL SCN (N= 37). The initial ALE maps were treated to a cluster forming threshold at p<.001 and an FWE corrected cluster-extent threshold at p<.05 prior to the conjunction and contrast analyses. The contrast maps in **Panel A** were additionally thresholded with a cluster forming threshold at p<.001 and a minimum cluster size of 100mm^3^. **Panel A** displays the conjunction alongside side statistically significant differences. In **Panel B,** we have overlaid the binarised versions of the complete ALE maps resulting from independent analysis of VERBAL ToM and VERBAL SCN studies. This allows for full visualisation of the topography of the associated networks (also see Supplementary Figure R4 & R5 and Supplementary Table R6 & R7). The sagittal and coronal sections are chosen as representative slices positioned over peak coordinates at which there is the greatest conjunction in the bilateral anterior temporal lobes (left y= 12; right y= 14).

### 3.3. The Role of Sensory Input Modality (VISUAL versus AUDITORY)

We also investigated the impact of sensory input modality. Importantly, both domains were dominated by experiments using visually presented stimuli. Comparisons limited to the auditory experiments were not possible due to a very small sample of ToM data. Overall, the pattern and extent of the common activation for VISUAL experiments (ToM: n= 106; SCN: n= 152) remained highly similar to our original analysis (**Section 3.1**), with common clusters of activation in key semantic areas (See **Figure 4** and **Supplementary Information No. 2: Supplementary Table R10**), including the left ATL, left IFG, left pMTG and ITG/FG and the left IPL/AG. There were also clusters of conjunction in the left medial SFG and precuneus. However, unlike in the initial analysis, there was no right ATL activation for the visual SCN experiments, and therefore no overlap between domains in the right ATL. Indeed, the cluster analyses revealed that visual relative to auditory SCN contrasts were less likely to contribute to the right ATL cluster, suggesting that it is unlikely that the absence of right ATL activation for visual SCN can be explained by reduced power per se. Instead, it seems more likely that the auditory contrasts were driving this cluster in the case of semantic cognition. One possibility is that this reflects increased effort in studies that use auditory stimuli (see **Discussion**). Other minor differences to the initial analyses are a diminished area of conjunction in the left middle STG and an absence of a conjunction in the right IFG (see **Supplementary Information No.2: Supplementary Figure CA1** and **Supplementary Table CA1** for more detail).

**Figure 4.**
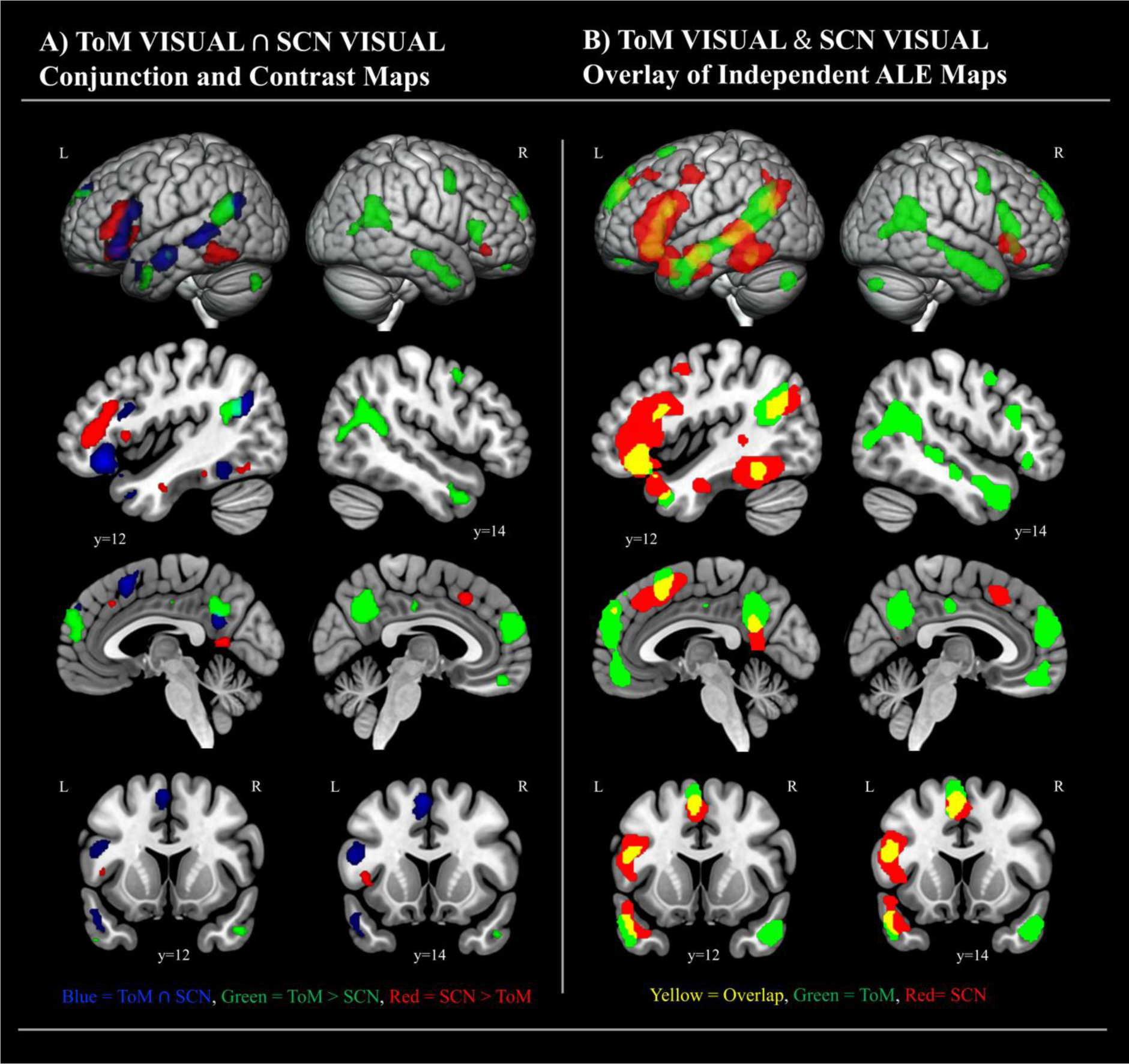
Common and differential activation for VISUAL ToM (N=106) and VISUAL SCN (N= 152). The initial ALE maps were treated to a cluster-forming threshold at p<.001, and an FWE-corrected cluster-extent threshold at p<.05 prior to the conjunction and contrast analyses. The contrast maps in **Panel A** were additionally thresholded with a cluster-forming threshold at p<.001 and a minimum cluster size of 100mm^3^. **Panel A** displays the conjunction alongside side statistically significant differences. In **Panel B,** we have overlaid the binarised versions of the complete ALE maps resulting from independent analysis of VERBAL ToM and VERBAL SCN studies. This allows for full visualisation of the topography of the associated networks (also see Supplementary Figures R8 & R9 and Supplementary Table R11 & R12). The sagittal and coronal sections are chosen as representative slices positioned over peak coordinates at which there is the greatest conjunction in the bilateral anterior temporal lobes (left y= 12; right y= 14).

As in our full analysis, the contrast analysis found more consistently identified activation in visual ToM than visual semantic cognition studies in the right TPJ, right IFG, precentral gyrus, anterior mPFC and precuneus. A small portion of the bilateral IFG remained more reliably engaged across SCN studies, as did the MFG, anterior mPFC and left precuneus. For more detail see the VISUAL and AUDITORY ToM and VISUAL and AUDITORY SCN ALE maps in **Supplementary Information No. 2: Supplementary Figures R8 & R9** and **Supplementary Tables R11 & R12**.

Although they do not directly relate to the study’s main questions, for sake of completeness and to allow for comparisons with prior meta-analyses (Molenberghs et al., 2016; Rice, Lambon Ralph, et al., 2015; Visser, Jefferies, et al., 2010) we also performed conjunctive and contrastive analyses within each domain which compare each stimulus format and sensory modality (e.g., comparisons of the VERBAL SCN and NON-VERBAL SCN data sets, the VISUAL SCN and AUDITORY SCN data). The results of these analyses can be found in the supplementary information (see **Supplementary Information No. 2: Supplementary Figures R6, R7 and R10 and Supplementary Tables R8, R9, R13 and R14)**.

## 4. Discussion

The present study aimed to glean a clearer understanding of the contribution of the general semantic system to social cognition. To this end, we took a neuroimaging meta-analytic approach to assess the degree to which engagement in ToM tasks shares neural correlates with semantic processes. The key findings were as follows:

1. Overall, there was a strikingly large degree of overlap between the activation likelihood maps for ToM and the SCN. This was most evident in the bilateral ATL, the left STS, left MTG, left TPJ, and left IFG, which are all key regions for semantic processing (Binder et al., 2009). This suggests that semantic processes are integral to performing theory of mind tasks.
2. Most differences that emerged were mainly a matter of the extent of regional activation, which is likely driven by discrepancies in the sample size contributing to each ALE map. Nonetheless, there were a few notable exceptions.
3. The right TPJ, anterior aspects of the bilateral MTG, bilateral mPFC, and the bilateral precuneus, were consistently identified in ToM but not SCN studies. Significant differences remained even after controlling for methodological factors, including the type of experimental stimuli, input modality and baseline condition used to probe each domain. This is consistent with claims that the function of these regions (e.g., the right TPJ) are tuned towards processing social stimuli.
4. The posterior ITG and dorsal IFG (both in the left hemisphere) were consistently identified in SCN studies but not in ToM studies. This difference was even more pronounced after controlling for stimulus format and modality. One possibility is that this reflects differences in task difficulty, which we did not account for (see Diveica et al., 2021).
5. Activation in bilateral IFG and SFG, irrespective of domain, appears to be driven by stimulus format. Right ATL activation could be driven by input modality. However, there are other uncontrolled methodological confounds that may have also played a role (e.g., task difficulty, processing effort, experiment number differences across domains). These findings highlight the need for future studies, whose aim is to contrast different cognitive domains, to systematically control for these types of methodological factors.

We interpret these results as generally supporting a recent proposal that social cognition draws upon a set of domain-general systems and processes dedicated to semantic cognition (Binney & Ramsey, 2020). We elaborate on these arguments and discuss each of the key findings in the following paragraphs.

### 4.1. Two sides of the same coin? The relationship between semantic cognition and theory of mind

It is argued that progress in social neuroscience theory will rapidly accelerate if the field embraces established models of other, more general domains of cognition (Amodio, 2019; Binney & Ramsey, 2020; Spunt & Adolphs, 2017). Theoretical advances in, for example, the domain of human learning and memory, are not always (immediately) incorporated within the social neuroscience literature, yet they are valuable opportunities to generate new hypotheses and more detailed models of social cognition, both in terms of mechanisms and neural bases (Amodio, 2019). Binney and Ramsey (2020) argue that reflections on theories of semantic cognition could prove particularly fruitful in this regard. They also highlight the striking similarities between the topologies of brain regions activated during neuroimaging studies of social cognition and semantic cognition, drawing particular attention to the ATL, the TPJ (including the angular gyrus and posterolateral temporal lobe), and the inferior frontal cortex. Prior to the present study, however, these activation maps had not been formally compared at the level of the whole brain (see Hodgson et al., 2022 for a region-specific analysis). Our results confirm a large degree of overlap, which raises questions about the nature of the various processes that afford the theory of mind ability (for related discussion, see (Arioli & Canessa, 2021; Deschrijver & Palmer, 2020). We specifically argue that it suggests that theory of mind processes involve cognitive mechanisms related to conceptual retrieval and semantic inference.

What does semantic processing contribute to theory of mind? Semantic memory or, conceptual knowledge, refers to a database of the meaning of words, objects, events and behaviours (Lambon Ralph et al., 2017). Thus, it is essential for recognising social signals, both verbal and nonverbal, that provide clues to someone’s cognitive or affective state. Moreover, it provides a means of cognitive abstraction that enables inference and representations of complex beliefs and intentions that we cannot directly observe (Adolphs, 2010; Binney & Ramsey, 2020). Finally, it guides the generation of responses that are appropriate to the observed behaviour, having considered the identity and social roles of the other agent or agents, as well as the wider social context. For example, should one see someone appear to laugh at a funeral, they must interpret the audiovisual signals and resolve any potential ambiguities (e.g., could it, in fact, be crying?). Then one must infer their likely mental state, particularly given their identity/role (e.g., the bereaved next of kin), and generate a context-appropriate social response (e.g., in this case, suppression of smiling or laughter). Now imagine the possible consequences of having impaired semantic knowledge (e.g., in semantic dementia, Rouse et al., 2024). Failure to correctly recognise the identity and/or the actions of the agent could lead to a misattribution of mental state, and/or socially inappropriate behaviour.

How tightly coupled are theory of mind and semantic processes? We argue our findings, together with prior patient (Binney, Henry, et al., 2016; Ding et al., 2020; Edwards-Lee et al., 1997; Snowden et al., 2018), animal (Klüver & Bucy, 1937) and neuroscientific studies involving healthy populations (Balgova et al., 2022; Diveica et al., 2021), suggest the underlying systems are closely linked (also see Binney & Ramsey, 2020; Olson et al., 2013; Rouse et al., 2024). One possibility is that theory of mind can be considered a case of semantic processes, rather than something distinct, and this means it would operate upon the same basic principles (and neural underpinnings; Binney & Ramsey, 2020). An alternative possibility is that theory of mind draws partly on general semantic processes (e.g., in the act of representing another’s cognitive/affective states), but also on distinct, more specialised processes which are supported by regions outside the SCN (e.g., systems involved in detecting the extent to which there is a mismatch between those states and one’s own; (Arioli & Canessa, 2021; Deschrijver & Palmer, 2020). Akin to this, is the finding that the cortical network supporting language processes (which involve activating links between words and meaning) is partially overlapping but also separable from the theory of mind network (Fedorenko & Varley, 2016; Paunov et al., 2022). Our results align more straightforwardly with the second possibility, although they do not rule out the first (see Section 4.2). At the very least, as we shall elaborate in the following paragraphs, our study advances understanding of the precise functional contribution of different brain regions to theory of mind. This level of specificity is sometimes missing from, and is important for development of, existing neurobiological accounts of theory of mind (Saxe & Kanwisher, 2003; Saxe & Wexler, 2005).

### 4.2 Functional fractionation of the ‘social brain’

We observed pervasive differences between the activation likelihood maps for ToM and SCN. Specifically, activation of the right TPJ, anterior aspects of the bilateral MTG, bilateral mPFC, and the bilateral precuneus appear more attuned to ToM tasks. All these regions are included in descriptions of putative brain regions specialized for theory of mind (Saxe, 2006; Saxe & Powell, 2006; Schurz et al., 2014, 2017). However, they are also considered part of the default-mode network (DMN) (Andrews-Hanna et al., 2014; Buckner et al., 2008; Spreng et al., 2009; Spreng & Grady, 2010), a resting-state network proposed to support various forms of internally orientated cognition (i.e., cognition that is decoupled from sensory processing (Margulies et al., 2016; Smallwood et al., 2013), including memory-driven cognition (Murphy et al., 2018). The DMN has been explicitly linked to social cognition (Mars et al., 2012; Spreng et al. 2009; Shillbach et al. 2008) although it has also been shown that regions activated by social tasks are, to some degree, distinct from what are considered ‘core’ regions of the DMN (Jackson, 2021; Jackson et al., 2016; Mars et al., 2012). In the present study, however, it was core DMN regions (especially those around the sagittal midline) that showed differences between semantic cognition and ToM. ‘Core’ regions have been argued to represent information related to the self and to allow for integration of Self and Other information via interaction with other DMN subsystems (Spreng & Andrews-Hanna, 2015).

Our results shed new light on the relationship between the ‘social brain’ and domain-general networks by highlighting significant overlap with the SCN. Important clues might also be gleaned from the way in which activation patterns diverge, and the fact that this occurs most notably within the right hemisphere homologues of left-lateralised SCN regions (e.g., the TPJ). One possible account of these observation is that engaging in ToM recruits the SCN plus additional regions that are more tuned to social processes. Alternatively, these regions may all comprise one widely distributed but nonetheless functionally integrated network, that exhibits systematic variation in the involvement of some of its nodes (particularly across hemispheres) owed to task-related or stimulus-related factors (e.g., input modality). ‘Socialness’ of a task (or perhaps the degree of involvement of Self- and Other-related processes (Chiou et al., 2022; Platek et al., 2004; Quesque & Brass, 2019) could be one such task-related factor (Binney & Ramsey, 2020; Pexman et al., 2023). Further research is needed to directly probe these factors and how they drive network involvement within and across domains. In the remainder of this discussion, we expand on debates surrounding the ATL and the TPJ because they are ascribed key roles in both ToM and in semantic cognition.

### 4.3. The role of anterior temporal lobes in theory of mind

Convergent neuropsychological and neuroimaging evidence strongly implicates the ATL in semantic knowledge representations. Semantic knowledge underpins a wide range of meaning-imbued behaviours, including language use, action understanding and interactions with objects (Patterson & Lambon Ralph, 2016; Lambon Ralph et al., 2017). By extension, we argue that the contribution of the ATL to ToM, and to social cognition more generally, is the supply of conceptual level information which constrains inferences about the intentions and actions of other agents (Binney & Ramsey, 2020). The current study revealed reliable overlap between ToM and semantic processing in the ATLs, which supports this hypothesis. The present findings also complement those of prior fMRI studies that directly explored the relationship between social and general semantic processing in the ATL. Across all these studies, two consistent findings have emerged. First, a ventrolateral portion of the left ATL responds equally to socially relevant concepts and more general concepts (both concrete and abstract), and this is irrespective of whether concepts are probed via verbal or pictorial stimuli (Binney, Hoffman, et al., 2016; Rice et al., 2018). The same ventrolateral region also activates during three different verbal and nonverbal ToM tasks, which suggests that conceptual information is accessed during ToM (Balgova et al., 2022). Second, there are some differences between social and general semantic tasks within the dorsolateral ATL (Binney et al., 2016; Rice et al., 2018; also see Arioli et al., 2021; Lin, Wang, et al., 2018; Lin, Yang, et al., 2018; Mellem et al., 2016; Ross & Olson, 2010; Zahn et al., 2007) although the location of this difference moves around across studies. Importantly, the differences are small compared to the amount of overlap. Indeed, the ATL subregion differentiating between ToM and SCN in the present study was abutting a much larger left ATL cluster which was activated consistently across both domains (also see Beauchamp, 2015; Deen et al., 2015 for comparisons of social perception with language and voice perception).

This overall pattern is consistent with the *graded semantic hub* account (Bajada et al., 2019; Binney et al., 2012; Rice, Hoffman, et al., 2015), which characterises the bilateral ATL as a unified representational space, all of which is engaged by the encoding and retrieval of semantic information of any kind. The centre of this hub exists over the ventrolateral ATL and its engagement in semantic processing is largely invariant to stimulus factors (e.g., modality). Towards the edges of this space, however, there are gradual shifts in semantic function such that regions on the periphery are more sensitive to certain types of semantic features (for a computational exploration of this general hypothesis, see Plaut, 2002). Why exactly ToM tasks would engage the dorsolateral ATL more than general semantic tasks is unclear. One possibility is that the meaning conveyed by typical ToM stimuli (i.e., the state of mind of an actor in absence of explicit descriptors) is not directly observable and, therefore must be inferred to a greater extent than in a typical semantic task. This may rely heavily on verbally-mediated semantic information, which has been shown to engage the dorsolateral ATL more (Binder et al., 2009; Rice, Hoffman, et al., 2015; Visser & Lambon Ralph, 2011). Another possibility is that it reflects a proximity to and strong connectivity with the limbic system (via the uncinate fasciculus; (Bajada et al., 2017; Binney et al., 2012; Papinutto et al., 2016) and a role of this ATL region in processing semantic features related to emotion (Olson et al., 2007; Rice, Hoffman, et al., 2015; Vigliocco et al., 2014).

The ventrolateral areas of the ATL implicated in recent studies of semantic processing (Binney et al., 2016; Rice et al., 2018) and theory of mind (Balgova et al., 2022) sit posterior to Brodmann’s area 38, and include the anterior ITG (including its basal surface) as well as the anterior fusiform. In the present study, the ATL subregions implicated were limited to the MTG and STG, and there was no evidence of more ventral involvement. This can be accounted for by signal distortion and signal loss that is typically observed with conventional forms of the fMRI technique. ATL-optimised distortion-corrected fMRI studies, on the other hand, detect robust ventral ATL activation during both semantic and ToM tasks (Balgova et al., 2022; Binney et al., 2010; Castelli et al., 2000; Devlin et al., 2000; Sharp et al., 2004). This methodological factor may also be particularly important for understanding the lack of left ATL activation for nonverbal stimuli. Prior distortion-corrected fMRI studies have shown that activation to nonverbal stimuli is almost entirely limited to ventral and ventromedial ATL structures which are regions that suffer the most from signal dropout (Rice, Lambon Ralph, et al., 2015; Visser, Embleton, et al., 2010).

There were also differences in the extent to which the right ATL was engaged, with a greater proportion of the right anterior MTG involved in ToM. Moreover, the involvement of the right ATL in semantic processing was dependent on including studies using auditory verbal stimuli. This confirms prior studies which also found that auditory verbal (or ‘spoken’) stimuli activate the ATL bilaterally, whereas written stimuli which show a left bias (Marinkovic et al., 2003; Rice, Lambon Ralph, et al., 2015). Thus, while ATL involvement in ToM appears always to be bilateral, right-sided involvement in semantic processing appears to be related to stimulus factors. This could be understood more broadly in terms of processing effort. Indeed, auditory semantic stimuli are typically sentences which require both rapid processing of individual tokens, as well as processing of combinatorial meaning, and which could work the semantic system more vigorously than other types of stimuli (see Visser, Jefferies, et al., 2010 for similar arguments). In a similar vein, the bilateral ATL activation during ToM tasks could reflect the semantic richness of stimuli. These observations are, however, not consistent with the right ATL having a distinctly social function (Bonnì et al., 2015; Gainotti, 2015; Gainotti et al., 2003; Gainotti & Marra, 2011; Pobric et al., 2016).

### 4.4. The Temporo-Parietal Junction

The TPJ has been associated with a variety of cognitive domains, including attention, language, and episodic memory, and many of them bilaterally (Binder et al., 2009; Humphreys & Lambon Ralph, 2015; Igelström & Graziano, 2017; Özdem et al., 2017). It is also now becoming clear that these functions fractionate along an anterior-posterior, as well as a dorsal-ventral axis (Bzdok et al., 2013; Hodgson & Lambon Ralph, 2008; Humphreys & Lambon Ralph, 2015). The present study shows that STS/STG and inferior parietal involvement in ToM is bilateral (Bzdok et al., 2012; Hodgson et al., 2022; Molenberghs et al., 2016; Schurz et al., 2014, 2020). The inferior parietal lobe (including the angular gyrus) is involved in semantic processing bilaterally (Binder et al., 2009; see also Bonner et al., 2013; Kuhnke et al., 2022), whereas posterior MTG/STS involvement is left-lateralised (Jackson, 2021). Taken together, these results suggest that parts of the left TPJ serve a function common to ToM and SCN (Numssen et al., 2020). For example, the left angular gyrus has been implicated in integration and storage of conceptual knowledge by some authors (Binder et al., 2009; Kuhnke et al., 2020) and attributed with a more domain-general role by others (e.g., the multi-sensory buffering of spatio-temporally extended representations; Humphreys, Lambon Ralph, et al., (2021; Humphreys & Tibon, 2022)). The left MTG/STS appears to be involved in processes that constrain semantic retrieval and which could also be engaged during ToM (Diveica et al., 2021). The right TPJ does not appear to be engaged by semantic processing, which is consistent with claims that it has a selective role in social and moral processing (Numssen et al., 2020; Saxe & Kanwisher, 2003; Saxe & Wexler, 2005; Young et al., 2010). However, the present study cannot rule out involvement in other cognitive domains.

### 4.5. Concluding remarks and future directions

In conclusion, we observed considerable overlap between the cortical networks engaged by semantic tasks and theory of mind tasks. These observations add to growing set of convergent findings from across neuropsychology, comparative and cognitive neuroscience which suggest this overlap reflects shared underlying processes and, further, that ToM relies in part on processes related to semantic cognition (Binney & Ramsey, 2020). Alternatively, this overlap could, on closer inspection, turn out to reflect tightly yet separately packed cognitive functions that only dissociate when investigated at higher spatial resolutions or at the level of individual participants (Lee & McCarthy, 2016). Further research is needed to explore these alternatives. Furthermore, inferences afforded by functional neuroimaging data are merely correlational and, therefore, the field needs to increasingly turn to patient models such as stroke, temporal lobe epilepsy, and frontotemporal dementia (Kumfor, Hazelton, et al., 2017; Kumfor, Honan, et al., 2017; Rankin, 2020, 2021), and non-invasive techniques like transcranial magnetic stimulation, to directly probe whether certain brain regions are necessary for both social *and* semantic cognition.

## Supporting information

Supplementary_Information_No.2_Results

## Acknowledgements and Funding

This work was supported by the Economic and Social Research Council (ESRC) Wales Doctoral Training Partnership in the form of a PhD studentship [ES/P00069X/1] (awarded to VD and RJB; PhD student: VD) and a British Academy Postdoctoral Fellowship [pf170068] (awarded to RLJ). The funding sources were not involved in the study design, in the collection, analysis or interpretation of data, in the report writing or decision to submit this work for publication. For the purpose of open access, the author has applied a Creative Commons Attribution (CC BY) licence to any Author Accepted Manuscript version arising.

## CRediT Author Statement

Eva Balgova: conceptualisation, methodology, data curation, investigation, formal analysis, visualisation, writing-original draft, writing-review and editing.

Veronica Diveica: conceptualisation, methodology, data curation, investigation, funding acquisition, writing-review and editing

Rebecca L. Jackson: conceptualisation, methodology, data curation, investigation, funding acquisition, writing-review and editing

Richard J. Binney: conceptualisation, methodology, funding acquisition, supervision, project administration, writing-original draft, writing-review and editing

## Data Availability Statement

Following open science initiatives (e.g., Munafò et al., 2017), the raw data sets, including study characteristics and the input and output files of all analyses, are openly available on the Open Science Framework (OSF) project page (https://osf.io/ydnxh/).

## References

Acar, F., Seurinck, R., Eickhoff, S. B., & Moerkerke, B. (2018). Assessing robustness against potential publication bias in Activation Likelihood Estimation (ALE) meta-analyses for fMRI. PLoS ONE, 13(11), 1–23. 10.1371/journal.pone.0208177

Adolphs, R. (2009). The Social Brain: Neural Basis of Social Knowledge. Annual Review of Psychology, 60(1), 693–716. 10.1146/annurev.psych.60.110707.163514

Adolphs, R. (2010). Conceptual Challenges and Directions for Social Neuroscience. Neuron, 65(6), 752–767. 10.1016/j.neuron.2010.03.006

Aichhorn, M., Perner, J., Kronbichler, M., Staffen, W., & Ladurner, G. (2006). Do visual perspective tasks need theory of mind? NeuroImage, 30(3), 1059–1068. 10.1016/j.neuroimage.2005.10.026

Aichhorn, M., Perner, J., Weiss, B., Kronbichler, M., Staffen, W., & Ladurner, G. (2009). Temporo-parietal Junction Activity in Theory-of-Mind Tasks: Falseness, Beliefs, or Attention. Journal of Cognitive Neuroscience, 21(6), 1179–1192. 10.1162/jocn.2009.21082

Alexander Diaz, B., van der Sluis, S., Moens, S., Benjamins, J. S., Migliorati, F., Stoffers, D., den Braber, A., Poil, S. S., Hardstone, R., Van’t Ent, D. V., Boomsma, D. I., de Geus, E., Mansvelder, H. D., Van Someren, E. J. W., & Linkenkaer-Hansen, K. (2013). The Amsterdam Resting-state Questionnaire reveals multiple phenotypes of resting-state cognition. Frontiers in Human Neuroscience, 7(JUL), 1–15. 10.3389/fnhum.2013.00446

Amodio, D. M. (2019). Social Cognition 2.0: An Interactive Memory Systems Account. Trends in Cognitive Sciences, 23(1), 21–33. 10.1016/j.tics.2018.10.002

Amodio, D. M., & Frith, C. D. (2006). Meeting of minds: The medial frontal cortex and social cognition. Nature Reviews Neuroscience, 7(4), 268–277. 10.1038/nrn1884

Andrews-Hanna, J. R., Smallwood, J., & Spreng, R. N. (2014). The default network and self-generated thought: component processes, dynamic control, and clinical relevance. Annals of the New York Academy of Sciences, 1316(1), 29–52. 10.1111/nyas.12360

Apperly, I. A. (2012). What is “theory of mind”? Concepts, cognitive processes and individual differences. Quarterly Journal of Experimental Psychology, 65(5), 825–839. 10.1080/17470218.2012.676055

Apperly, I. A., Samson, D., & Humphreys, G. W. (2005). Domain-specificity and theory of mind: evaluating neuropsychological evidence. Trends in Cognitive Sciences, 9(12), 572–577.

Arioli, M., & Canessa, N. (2021). Overlapping and specific neural correlates for empathizing, affective mentalizing, and cognitive mentalizing : A coordinate-based meta-analytic study. March, 1–28. 10.1002/hbm.25570

Arioli, M., Gianelli, C., & Canessa, N. (2020). Neural representation of social concepts: a coordinate-based meta-analysis of fMRI studies. Brain Imaging and Behavior. 10.1007/s11682-020-00384-6

Arioli, M., Gianelli, C., & Canessa, N. (2021). Neural representation of social concepts: a coordinate-based meta-analysis of fMRI studies. Brain Imaging and Behavior, 15(4), 1912–1921. 10.1007/s11682-020-00384-6

Assem, M., Glasser, M. F., Van Essen, D. C., & Duncan, J. (2020). A Domain-General Cognitive Core Defined in Multimodally Parcellated Human Cortex. Cerebral Cortex (New York, N.Y. : 1991), 30(8), 4361–4380. 10.1093/cercor/bhaa023

Baetens, K., Ma, N., Steen, J., & Van Overwalle, F. (2013). Involvement of the mentalizing network in social and non-social high construal. Social Cognitive and Affective Neuroscience, 9(6), 817–824. 10.1093/scan/nst048

Bajada, C. J., Haroon, H. A., Azadbakht, H., Parker, G. J. M., Lambon Ralph, M. A., & Cloutman, L. L. (2017). The tract terminations in the temporal lobe: Their location and associated functions. Cortex, 97, 277–290. 10.1016/j.cortex.2016.03.013

Bajada, C. J., Trujillo-Barreto, N. J., Parker, G. J. M., Cloutman, L. L., & Lambon Ralph, M. A. (2019). A structural connectivity convergence zone in the ventral and anterior temporal lobes: Data-driven evidence from structural imaging. Cortex, 120, 298–307. 10.1016/j.cortex.2019.06.014

Balgova, E., Diveica, V., Walbrin, J., & Binney, R. J. (2022). The role of the ventrolateral anterior temporal lobes in social cognition. Human Brain Mapping, May, 1–20. 10.1002/hbm.25976

Beauchamp, M. S. (2015). The social mysteries of the superior temporal sulcus. Trends in Cognitive Sciences, 19(9), 489–490. 10.1016/j.tics.2015.07.002

Bertoux, M., Duclos, H., Caillaud, M., Segobin, S., Merck, C., de La Sayette, V., Belliard, S., Desgranges, B., Eustache, F., & Laisney, M. (2020). When affect overlaps with concept: emotion recognition in semantic variant of primary progressive aphasia. Brain, 3850–3864. 10.1093/brain/awaa313

Binder, J. R., Conant, L. L., Humphries, C. J., Fernandino, L., Simons, S. B., Aguilar, M., Desai, R. H., Binder, J. R., Conant, L. L., Humphries, C. J., Simons, S. B., Aguilar, M., & Desai, R. H. (2016). Toward a brain-based componential semantic representation. Cognitive Neuropsychology, 33(3–4), 130–174. 10.1080/02643294.2016.1147426

Binder, J. R., Desai, R. H., Graves, W. W., & Conant, L. L. (2009). Where is the semantic system? A critical review and meta-analysis of 120 functional neuroimaging studies. Cerebral Cortex, 19(12), 2767–2796. 10.1093/cercor/bhp055

Binney, R. J., Embleton, K. V., Jefferies, E., Parker, G. J. M. M., & Lambon Ralph, M. A. (2010). The Ventral and Inferolateral Aspects of the Anterior Temporal Lobe Are Crucial in Semantic Memory: Evidence from a Novel Direct Comparison of Distortion-Corrected fMRI, rTMS, and Semantic Dementia. Cerebral Cortex, 20(11), 2728–2738. 10.1093/cercor/bhq019

Binney, R. J., Henry, M. L., Babiak, M., Pressman, P. S., Santos-Santos, M. A., Narvid, J., Mandelli, M. L., Strain, P. J., Miller, B. L., Rankin, K. P., Rosen, H. J., & Gorno-Tempini, M. L. (2016). Reading words and other people: A comparison of exception word, familiar face and affect processing in the left and right temporal variants of primary progressive aphasia. Cortex, 82, 147–163. 10.1016/j.cortex.2016.05.014

Binney, R. J., Hoffman, P., & Lambon Ralph, M. A. (2016). Mapping the Multiple Graded Contributions of the Anterior Temporal Lobe Representational Hub to Abstract and Social Concepts: Evidence from Distortion-corrected fMRI. Cerebral Cortex, 26(11), 4227–4241. 10.1093/cercor/bhw260

Binney, R. J., Parker, G. J. M., & Lambon Ralph, M. A. (2012). Convergent Connectivity and Graded Specialization in the Rostral Human Temporal Lobe as Revealed by Diffusion-Weighted Imaging Probabilistic Tractography. Journal of Cognitive Neuroscience, 24(10), 1998–2014. 10.1162/jocn_a_00263

Binney, R. J., & Ramsey, R. (2020). Social Semantics: The role of conceptual knowledge and cognitive control in a neurobiological model of the social brain. In Neuroscience and Biobehavioral Reviews (Vol. 112, pp. 28–38). Elsevier Ltd. 10.1016/j.neubiorev.2020.01.030

Bonner, M. F., Peelle, J. E., Cook, P. A., & Grossman, M. (2013). Heteromodal conceptual processing in the angular gyrus. NeuroImage, 71, 175–186. 10.1016/j.neuroimage.2013.01.006

Bonnì, S., Koch, G., Miniussi, C., Bassi, M. S., Caltagirone, C., & Gainotti, G. (2015). Role of the anterior temporal lobes in semantic representations: Paradoxical results of a cTBS study. Neuropsychologia, 76, 163–169. 10.1016/j.neuropsychologia.2014.11.002

Borghesani, V., Narvid, J., Battistella, G., Shwe, W., Watson, C., Binney, R. J., Sturm, V., Miller, Z., Mandelli, M. L., Miller, B., & Gorno-Tempini, M. L. (2019). “Looks familiar, but I do not know who she is”: The role of the anterior right temporal lobe in famous face recognition. Cortex, 115, 72–85. 10.1016/j.cortex.2019.01.006

Botvinik-Nezer, R., Holzmeister, F., Camerer, C. F., Dreber, A., Huber, J., Johannesson, M., Kirchler, M., Iwanir, R., Mumford, J. A., Adcock, R. A., Avesani, P., Baczkowski, B. M., Bajracharya, A., Bakst, L., Ball, S., Barilari, M., Bault, N., Beaton, D., Beitner, J., … Schonberg, T. (2020). Variability in the analysis of a single neuroimaging dataset by many teams. Nature, 582(7810), 84–88. 10.1038/s41586-020-2314-9

Brüne, M., & Brüne-Cohrs, U. (2006). Theory of mind—evolution, ontogeny, brain mechanisms and psychopathology. Neuroscience & Biobehavioral Reviews, 30(4), 437–455. 10.1016/J.NEUBIOREV.2005.08.001

Buckner, R. L., Andrews-Hanna, J. R., & Schacter, D. L. (2008). The brain’s default network: Anatomy, function, and relevance to disease. In Annals of the New York Academy of Sciences (Vol. 1124, pp. 1–38). 10.1196/annals.1440.011

Button, K. S., Ioannidis, J. P. A., Mokrysz, C., Nosek, B. A., Flint, J., Robinson, E. S. J., & Munafò, M. R. (2013). Power failure: Why small sample size undermines the reliability of neuroscience. Nature Reviews Neuroscience, 14(5), 365–376. 10.1038/nrn3475

Bzdok, D., Langner, R., Schilbach, L., Jakobs, O., Roski, C., Caspers, S., Laird, A. R., Fox, P. T., Zilles, K., & Eickhoff, S. B. (2013). Characterization of the temporo-parietal junction by combining data-driven parcellation, complementary connectivity analyses, and functional decoding. NeuroImage, 81, 381–392. 10.1016/j.neuroimage.2013.05.046

Bzdok, D., Schilbach, L., Vogeley, K., Schneider, K., Laird, A. R., Langner, R., & Eickhoff, S. B. (2012). Parsing the neural correlates of moral cognition: ALE meta-analysis on morality, theory of mind, and empathy. Brain Structure and Function, 217(4), 783–796.

Cabeza, R., Ciaramelli, E., & Moscovitch, M. (2012). Cognitive contributions of the ventral parietal cortex: an integrative theoretical account. Trends in Cognitive Sciences, 16(6), 338–352. 10.1016/j.tics.2012.04.008

Carp, J. (2012). On the plurality of (methodological) worlds: Estimating the analytic flexibility of fmri experiments. Frontiers in Neuroscience, 6(OCT), 1–13. 10.3389/fnins.2012.00149

Castelli, F., Happé, F., Frith, U., & Frith, C. (2000). Movement and Mind: A Functional Imaging Study of Perception and Interpretation of Complex Intentional Movement Patterns. NeuroImage, 12(3), 314–325. 10.1006/nimg.2000.0612

Catricalà, E., Conca, F., Fertonani, A., Miniussi, C., & Cappa, S. F. (2020). State-dependent TMS reveals the differential contribution of ATL and IPS to the representation of abstract concepts related to social and quantity knowledge. Cortex, 123, 30–41. 10.1016/j.cortex.2019.09.018

Chiou, R., Cox, C. R., & Lambon Ralph, M. A. (2022). Bipartite functional fractionation within the neural system for social cognition supports the psychological continuity of self versus other. Cerebral Cortex, 1–23. 10.1093/cercor/bhac135

Chiou, R., Humphreys, G. F., & Lambon Ralph, M. A. (2020). Bipartite functional fractionation within the default network supports disparate forms of internally oriented cognition. Cerebral Cortex, 30(10), 5484–5501. 10.1093/cercor/bhaa130

Cumming, G. (2014). The New Statistics: Why and How. Psychological Science, 25(1), 7–29. 10.1177/0956797613504966

Deen, B., Koldewyn, K., Kanwisher, N., & Saxe, R. (2015). Functional Organization of Social Perception and Cognition in the Superior Temporal Sulcus. Cerebral Cortex, 25(11), 4596–4609. 10.1093/cercor/bhv111

Deschrijver, E., & Palmer, C. (2020). Reframing social cognition: Relational versus representational mentalizing. Psychological Bulletin, 146(11), 941–969. 10.1037/bul0000302

Devlin, J. T., Russell, R. P., Davis, M. H., Price, C. J., Wilson, J., Moss, H. E., Matthews, P. M., & Tyler, L. K. (2000). Susceptibility-Induced Loss of Signal: Comparing PET and fMRI on a Semantic Task. NeuroImage, 11(6), 589–600. 10.1006/nimg.2000.0595

Ding, J., Chen, K., Liu, H., Huang, L., Chen, Y., Lv, Y., Yang, Q., Guo, Q., Han, Z., & Lambon Ralph, M. A. (2020). A unified neurocognitive model of semantics language social behaviour and face recognition in semantic dementia. Nature Communications, 11(1). 10.1038/s41467-020-16089-9

Diveica, V., Koldewyn, K., & Binney, R. J. (2021). Establishing a role of the semantic control network in social cognitive processing: A meta-analysis of functional neuroimaging studies. NeuroImage, 245(November), 118702. 10.1016/j.neuroimage.2021.118702

Dodell-Feder, D., Koster-Hale, J., Bedny, M., & Saxe, R. (2011). FMRI item analysis in a theory of mind task. NeuroImage, 55(2), 705–712. 10.1016/j.neuroimage.2010.12.040

Downing, P., & Kanwisher, N. (2010). A cortical area specialized for visual processing of the human body. Journal of Vision, 1(3), 341–341. 10.1167/1.3.341

Duncan, J. (2010). The multiple-demand (MD) system of the primate brain: mental programs for intelligent behaviour. Trends in Cognitive Sciences, 14(4), 172–179. 10.1016/j.tics.2010.01.004

Duncan, J. (2013). The Structure of Cognition: Attentional Episodes in Mind and Brain. Neuron, 80(1), 35–50. 10.1016/j.neuron.2013.09.015

Edwards-Lee, T., Miller, B. L., Benson, D. F., Cummings, J. L., Russell, G. L., Boone, K., & Mena, I. (1997). The temporal variant of frontotemporal dementia. Brain, 120(6), 1027–1040. 10.1093/brain/120.6.1027

Eickhoff, S. B., Bzdok, D., Laird, A. R., Kurth, F., & Fox, P. T. (2012). Activation likelihood estimation meta-analysis revisited. NeuroImage, 59(3), 2349–2361. 10.1016/j.neuroimage.2011.09.017

Eickhoff, S. B., Bzdok, D., Laird, A. R., Roski, C., Caspers, S., Zilles, K., & Fox, P. T. (2011). Co-activation patterns distinguish cortical modules, their connectivity and functional differentiation. NeuroImage, 57(3), 938–949. 10.1016/j.neuroimage.2011.05.021

Eickhoff, S. B., Laird, A. R., Fox, P. M., Lancaster, J. L., & Fox, P. T. (2017). Implementation errors in the GingerALE Software: Description and recommendations. Human Brain Mapping, 38(1), 7–11. 10.1002/hbm.23342

Eickhoff, S. B., Laird, A. R., Grefkes, C., Wang, L. E., Zilles, K., & Fox, P. T. (2009). Coordinate-based activation likelihood estimation meta-analysis of neuroimaging data: A random-effects approach based on empirical estimates of spatial uncertainty. Human Brain Mapping, 30(9), 2907–2926. 10.1002/hbm.20718

Eickhoff, S. B., Nichols, T. E., Laird, A. R., Hoffstaedter, F., Amunts, K., Fox, P. T., Bzdok, D., & Eickhoff, C. R. (2016). Behavior, sensitivity, and power of activation likelihood estimation characterized by massive empirical simulation. NeuroImage, 137, 70–85. 10.1016/j.neuroimage.2016.04.072

Eickhoff, S., Bzdok, D., Laird, A., Roski, C., Zilles, K., & Fox, P. (2011). Connectivity and Functional Differentiation. NeuroImage, 57(3), 938–949. 10.1016/j.neuroimage.2011.05.021.Co-activation

Elfgren, C., van Westen, D., Passant, U., Larsson, E.-M., Mannfolk, P., & Fransson, P. (2006). fMRI activity in the medial temporal lobe during famous face processing. NeuroImage, 30(2), 609–616. 10.1016/j.neuroimage.2005.09.060

Fedorenko, E., Duncan, J., & Kanwisher, N. (2013). Broad domain generality in focal regions of frontal and parietal cortex. Proceedings of the National Academy of Sciences of the United States of America, 110(41), 16616–16621. 10.1073/pnas.1315235110

Fedorenko, E., & Varley, R. (2016). Language and thought are not the same thing: Evidence from neuroimaging and neurological patients. Annals of the New York Academy of Sciences, 1369(1), 132–153. 10.1111/nyas.13046

Friederici, A. D. (2011). The brain basis of language processing: From structure to function. Physiological Reviews, 91(4), 1357–1392. 10.1152/physrev.00006.2011

Frith, C. D. (2007). The social brain? Philosophical Transactions of the Royal Society B: Biological Sciences, 362(1480), 671–678. 10.1098/rstb.2006.2003

Frith, C. D., & Frith, U. (2007). Social Cognition in Humans. Current Biology, 17(16), R724–R732. 10.1016/j.cub.2007.05.068

Frith, C. D., & Frith, U. (2012). Mechanisms of Social Cognition. Annual Review of Psychology, 63(1), 287–313. 10.1146/annurev-psych-120710-100449

Frith, C., & Frith, U. (2005). Theory of mind. Current Biology, 15(17), R644–R645. 10.1016/j.cub.2005.08.041

Frith, U., & Frith, C. (2010). The social brain: allowing humans to boldly go where no other species has been. Philosophical Transactions of the Royal Society B: Biological Sciences, 365(1537), 165–176. 10.1098/rstb.2009.0160

Gainotti, G. (2015). Is the difference between right and left ATLs due to the distinction between general and social cognition or between verbal and non-verbal representations? Neuroscience and Biobehavioral Reviews, 51, 296–312. 10.1016/j.neubiorev.2015.02.004

Gainotti, G., Barbier, A., & Marra, C. (2003). Slowly progressive defect in recognition of familiar people in a patient with right anterior temporal atrophy. Brain, 126(4), 792–803. 10.1093/brain/awg092

Gainotti, G., & Marra, C. (2011). Differential contribution of right and left temporo-occipital and anterior temporal lesions to face recognition disorders. Frontiers in Human Neuroscience, 5(JUNE), 1–11. 10.3389/fnhum.2011.00055

Gallagher, H. L., Happé, F., Brunswick, N., Fletcher, P. C., Frith, U., & Frith, C. D. (2000). Reading the mind in cartoons and stories: an fMRI study of ‘theory of mind’ in verbal and nonverbal tasks. Neuropsychologia, 38(1), 11–21. 10.1016/S0028-3932(99)00053-6

Geng, J. J., & Vossel, S. (2013). Re-evaluating the role of TPJ in attentional control: Contextual updating? Neuroscience and Biobehavioral Reviews, 37(10), 2608–2620. 10.1016/j.neubiorev.2013.08.010

Gorno-tempini, M. L., Rankin, K. P., Woolley, J. D., Rosen, H. J., Phengrasamy, L., & Miller, B. L. (2003). Right Temporal Atrophy.

Grabowski, T. J., Damasio, H., Tranel, D., Ponto, L. L. B., Hichwa, R. D., & Damasio, A. R. (2001). A role for left temporal pole in the retrieval of words for unique entities. Human Brain Mapping, 13(4), 199–212. 10.1002/hbm.1033

Gweon, H., Dodell-Feder, D., Bedny, M., & Saxe, R. (2012). Theory of Mind Performance in Children Correlates With Functional Specialization of a Brain Region for Thinking About Thoughts. Child Development, 83(6), 1853–1868. 10.1111/j.1467-8624.2012.01829.x

Handjaras, G., Leo, A., Cecchetti, L., Papale, P., Lenci, A., Marotta, G., Pietrini, P., & Ricciardi, E. (2017). Modality-independent encoding of individual concepts in the left parietal cortex. Neuropsychologia. 10.1016/j.neuropsychologia.2017.05.001

Happé, F., Cook, J. L., & Bird, G. (2017). The Structure of Social Cognition: In(ter)dependence of Sociocognitive Processes. Annual Review of Psychology, 68(1), 243–267. 10.1146/annurev-psych-010416-044046

Heleven, E., & Van Overwalle, F. (2018). The neural basis of representing others’ inner states. In Current Opinion in Psychology (Vol. 23, pp. 98–103). Elsevier B.V. 10.1016/j.copsyc.2018.02.003

Hodgson, C., & Lambon Ralph, M. A. (2008). Mimicking aphasic semantic errors in normal speech production: Evidence from a novel experimental paradigm. Brain and Language, 104(1), 89–101. 10.1016/j.bandl.2007.03.007

Hodgson, V. J., Lambon Ralph, M. A., & Jackson, R. L. (2022). The cross-domain functional organization of posterior lateral temporal cortex: insights from ALE meta-analyses of 7 cognitive domains spanning 12,000 participants. Cerebral Cortex, 1–17. 10.1093/cercor/bhac394

Hoffman, P., & Morcom, A. M. (2018). Age-related changes in the neural networks supporting semantic cognition: A meta-analysis of 47 functional neuroimaging studies. Neuroscience and Biobehavioral Reviews, 84(November), 134–150. 10.1016/j.neubiorev.2017.11.010

Hugdahl, K., Raichle, M. E., Mitra, A., & Specht, K. (2015). On the existence of a generalized non-specific task-dependent network. Frontiers in Human Neuroscience, 9(AUGUST), 1–15. 10.3389/fnhum.2015.00430

Hughes, C., Cassidy, B. S., Faskowitz, J., Avena-Koenigsberger, A., Sporns, O., & Krendl, A. C. (2019). Age differences in specific neural connections within the Default Mode Network underlie theory of mind. NeuroImage, 191, 269–277. 10.1016/J.NEUROIMAGE.2019.02.024

Humphreys, G. F., Hoffman, P., Visser, M., Binney, R. J., & Lambon Ralph, M. A. (2015). Establishing task- and modality-dependent dissociations between the semantic and default mode networks. Proceedings of the National Academy of Sciences, 112(25), 7857–7862. 10.1073/pnas.1422760112

Humphreys, G. F., & Lambon Ralph, M. A. (2015). Fusion and Fission of Cognitive Functions in the Human Parietal Cortex. Cerebral Cortex, 25(10), 3547–3560. 10.1093/cercor/bhu198

Humphreys, G. F., Lambon Ralph, M. A., & Simons, J. S. (2021). A Unifying Account of Angular Gyrus Contributions to Episodic and Semantic Cognition. Trends in Neurosciences, 44(6), 452–463. 10.1016/j.tins.2021.01.006

Humphreys, G. F., Ralph, M. A. L., Simons, J. S., Lambon Ralph, M. A., & Simons, J. S. (2021). A Unifying Account of Angular Gyrus Contributions to Episodic and Semantic Cognition. Trends in Neurosciences, 44(6), 452–463. 10.1016/j.tins.2021.01.006

Humphreys, G. F., & Tibon, R. (2022). Dual-axes of functional organisation across lateral parietal cortex: the angular gyrus forms part of a multi-modal buffering system. Brain Structure and Function, 0123456789. 10.1007/s00429-022-02510-0

Hyatt, C. J., Calhoun, V. D., Pearlson, G. D., & Assaf, M. (2015). Specific default mode subnetworks support mentalizing as revealed through opposing network recruitment by social and semantic FMRI tasks. Human Brain Mapping, 36(8), 3047–3063. 10.1002/hbm.22827

Igelström, K. M., & Graziano, M. S. A. (2017). The inferior parietal lobule and temporoparietal junction: A network perspective. Neuropsychologia, 105(January), 70–83. 10.1016/j.neuropsychologia.2017.01.001

Irish, M., Hodges, J. R., & Piguet, O. (2014). Right anterior temporal lobe dysfunction underlies theory of mind impairments in semantic dementia. Brain, 137(4), 1241–1253. 10.1093/brain/awu003

Jackson, R. L. (2021). The neural correlates of semantic control revisited. NeuroImage, 224(September 2020), 117444. 10.1016/j.neuroimage.2020.117444

Jackson, R. L., Cloutman, L. L., & Lambon Ralph, M. A. (2019). Exploring distinct default mode and semantic networks using a systematic ICA approach. Cortex, 113, 279–297. 10.1016/j.cortex.2018.12.019

Jackson, R. L., Hoffman, P., Pobric, G., & Lambon Ralph, M. A. (2016). The semantic network at work and rest: Differential connectivity of anterior temporal lobe subregions. Journal of Neuroscience, 36(5), 1490–1501. 10.1523/JNEUROSCI.2999-15.2016

Jackson, R. L., Humphreys, G. F., Rice, G. E., Binney, R. J., Lambon Ralph, M. A., & Jackson or Matthew Lambon Ralph, R. A. (2022). A Network-level Test of the Role of the Co-activated Default Mode Network in Episodic Recall and Social Cognition. BioRxiv, 44(0), 2021.01.08.425921. https://www.biorxiv.org/content/10.1101/2021.01.08.425921v2%0Ahttps://www.biorxiv.org/content/10.1101/2021.01.08.425921v2.abstract

Jacoby, N., Bruneau, E., Koster-Hale, J., & Saxe, R. (2016). Localizing Pain Matrix and Theory of Mind networks with both verbal and non-verbal stimuli. NeuroImage, 126, 39–48. 10.1016/j.neuroimage.2015.11.025

Jefferies, E. (2013). The neural basis of semantic cognition: Converging evidence from neuropsychology, neuroimaging and TMS. In Cortex (Vol. 49, Issue 3, pp. 611–625). Masson SpA. 10.1016/j.cortex.2012.10.008

Jenkins, A. C., Dodell-Feder, D., Saxe, R., & Knobe, J. (2014). The neural bases of directed and spontaneous mental state attributions to group agents. PLoS ONE, 9(8). 10.1371/journal.pone.0105341

Kanwisher, N., & Yovel, G. (2006). The fusiform face area: A cortical region specialized for the perception of faces. Philosophical Transactions of the Royal Society B: Biological Sciences, 361(1476), 2109–2128. 10.1098/rstb.2006.1934

Klüver, H., & Bucy, P. C. (1937). “ Psychic blindness” and other symptoms following bilateral temporal lobectomy in Rhesus monkeys. American Journal of Physiology, 119, 3552–353.

Koster-Hale, J., & Saxe, R. (2013). Theory of Mind: A Neural Prediction Problem. Neuron, 79(5), 836–848. 10.1016/j.neuron.2013.08.020

Krieger-Redwood, K., Teige, C., Davey, J., Hymers, M., & Jefferies, E. (2015). Conceptual control across modalities: Graded specialisation for pictures and words in inferior frontal and posterior temporal cortex. Neuropsychologia, 76, 92–107. 10.1016/j.neuropsychologia.2015.02.030

Kuhnke, P., Chapman, C. A., Cheung, V. K. M., Turker, S., Graessner, A., Martin, S., Williams, K. A., & Hartwigsen, G. (2021). The role of the angular gyrus in semantic cognition - A synthesis of five functional neuroimaging studies (Issue December). 10.1101/2021.12.21.473704

Kuhnke, P., Chapman, C. A., Cheung, V. K. M., Turker, S., Graessner, A., Martin, S., Williams, K. A., & Hartwigsen, G. (2022). The role of the angular gyrus in semantic cognition: a synthesis of five functional neuroimaging studies. Brain Structure and Function, 0123456789. 10.1007/s00429-022-02493-y

Kuhnke, P., Kiefer, M., & Hartwigsen, G. (2020). Task-Dependent Recruitment of Modality-Specific and Multimodal Regions during Conceptual Processing. Cerebral Cortex (New York, N.Y. : 1991), 30(7), 3938–3959. 10.1093/cercor/bhaa010

Kumfor, F., Hazelton, J. L., De Winter, F.-L., de Langavant, L. C., & Van den Stock, J. (2017). Clinical Studies of Social Neuroscience: A Lesion Model Approach. In Neuroscience and Social Science (pp. 255–296). Springer International Publishing. 10.1007/978-3-319-68421-5_12

Kumfor, F., Honan, C., McDonald, S., Hazelton, J. L., Hodges, J. R., & Piguet, O. (2017). Assessing the “social brain” in dementia: Applying TASIT-S. Cortex, 93, 166–177. 10.1016/J.CORTEX.2017.05.022

Laird, A. R., Lancaster, J. L., & Fox, P. T. (2005). BrainMap: The social evolution of a human brain mapping database. Neuroinformatics, 3(1), 65–77. 10.1385/ni:3:1:065

Lambon Ralph, M. A., Mcclelland, J. L., Patterson, K., Galton, C. J., & Hodges, J. R. (2001). No right to speak? The relationship between object naming and semantic impairment: Neuropsychological evidence and a computational model. Journal of Cognitive Neuroscience, 13(3), 341–356. 10.1162/08989290151137395

Lambon Ralph, M. A. L., Jefferies, E., Patterson, K., & Rogers, T. T. (2017). The neural and computational bases of semantic cognition. Nature Reviews Neuroscience, 18(1), 42–55. 10.1038/nrn.2016.150

Landsiedel, J., Daughters, K., Downing, P. E., & Koldewyn, K. (2022). The role of motion in the neural representation of social interactions in the posterior temporal cortex. NeuroImage, 119533. 10.1016/j.neuroimage.2022.119533

Lee, S. M., & McCarthy, G. (2016). Functional Heterogeneity and Convergence in the Right Temporoparietal Junction. Cerebral Cortex, 26(3), 1108–1116. 10.1093/cercor/bhu292

Leveroni, C. L., Seidenberg, M., Mayer, A. R., Mead, L. A., Binder, J. R., & Rao, S. M. (2000). Neural Systems Underlying the Recognition of Familiar and Newly Learned Faces. The Journal of Neuroscience, 20(2), 878–886. 10.1523/JNEUROSCI.20-02-00878.2000

Lin, N., Wang, X., Xu, Y., Wang, X., Hua, H., Zhao, Y., & Li, X. (2018). Fine Subdivisions of the Semantic Network Supporting Social and Sensory–Motor Semantic Processing. Cerebral Cortex, 28(8), 2699–2710. 10.1093/cercor/bhx148

Lin, N., Yang, X., Li, J., Wang, S., Hua, H., Ma, Y., & Li, X. (2018). Neural correlates of three cognitive processes involved in theory of mind and discourse comprehension. Cognitive, Affective and Behavioral Neuroscience, 18(2), 273–283. 10.3758/s13415-018-0568-6

Mar, R. A. (2011). The neural bases of social cognition and story comprehension. Annual Review of Psychology, 62, 103–134. 10.1146/annurev-psych-120709-145406

Margulies, D. S., Ghosh, S. S., Goulas, A., Falkiewicz, M., Huntenburg, J. M., Langs, G., Bezgin, G., Eickhoff, S. B., Castellanos, F. X., Petrides, M., Jefferies, E., & Smallwood, J. (2016a). Situating the default-mode network along a principal gradient of macroscale cortical organization. Proceedings of the National Academy of Sciences of the United States of America, 113(44), 12574–12579. 10.1073/pnas.1608282113

Margulies, D. S., Ghosh, S. S., Goulas, A., Falkiewicz, M., Huntenburg, J. M., Langs, G., Bezgin, G., Eickhoff, S. B., Castellanos, F. X., Petrides, M., Jefferies, E., & Smallwood, J. (2016b). Situating the default-mode network along a principal gradient of macroscale cortical organization. Proceedings of the National Academy of Sciences of the United States of America, 113(44), 12574–12579. 10.1073/pnas.1608282113

Marinkovic, K., Dhond, R. P., Dale, A. M., Glessner, M., Carr, V., & Halgren, E. (2003). Spatiotemporal Dynamics of Modality-Specific and Supramodal Word Processing. Neuron, 38(3), 487–497. 10.1016/S0896-6273(03)00197-1

Mars, R. B., Neubert, F. X., Noonan, M. A. P., Sallet, J., Toni, I., & Rushworth, M. F. S. (2012). On the relationship between the “default mode network” and the “social brain.” Frontiers in Human Neuroscience, JUNE 2012, 1–9. 10.3389/fnhum.2012.00189

Mellem, M. S., Jasmin, K. M., Peng, C., & Martin, A. (2016). Sentence processing in anterior superior temporal cortex shows a social-emotional bias. Neuropsychologia, 89, 217–224. 10.1016/j.neuropsychologia.2016.06.019

Meyer, M. L., Spunt, R. P., Berkman, E. T., Taylor, S. E., & Lieberman, M. D. (2012). Evidence for social working memory from a parametric functional MRI study. Proceedings of the National Academy of Sciences of the United States of America, 109(6), 1883–1888. 10.1073/pnas.1121077109

Miller, L. A., Hsieh, S., Lah, S., Savage, S., Hodges, J. R., & Piguet, O. (2012). One size does not fit all: Face emotion processing impairments in semantic dementia, behavioural-variant frontotemporal dementia and Alzheimer’s disease are mediated by distinct cognitive deficits. Behavioural Neurology, 25(1), 53–60. 10.3233/BEN-2012-0349

Molenberghs, P., Johnson, H., Henry, J. D., & Mattingley, J. B. (2016a). Understanding the minds of others: A neuroimaging meta-analysis. Neuroscience and Biobehavioral Reviews, 65, 276–291. 10.1016/j.neubiorev.2016.03.020

Molenberghs, P., Johnson, H., Henry, J. D., & Mattingley, J. B. (2016b). Understanding the minds of others: A neuroimaging meta-analysis. Neuroscience and Biobehavioral Reviews, 65, 276–291. 10.1016/j.neubiorev.2016.03.020

Müller, V. I., Cieslik, E. C., Laird, A. R., Fox, P. T., Radua, J., Mataix-Cols, D., Tench, C. R., Yarkoni, T., Nichols, T. E., Turkeltaub, P. E., Wager, T. D., & Eickhoff, S. B. (2018). Ten simple rules for neuroimaging meta-analysis. Neuroscience & Biobehavioral Reviews, 84(April 2017), 151–161. 10.1016/j.neubiorev.2017.11.012

Munafò, M. R., Nosek, B. A., Bishop, D. V. M., Button, K. S., Chambers, C. D., Percie Du Sert, N., Simonsohn, U., Wagenmakers, E. J., Ware, J. J., & Ioannidis, J. P. A. (2017). A manifesto for reproducible science. In Nature Human Behaviour (Vol. 1, Issue 1, pp. 1–9). Nature Publishing Group. 10.1038/s41562-016-0021

Murphy, C., Jefferies, E., Rueschemeyer, S.-A., Sormaz, M., Wang, H., Margulies, D. S., & Smallwood, J. (2018). Distant from input: Evidence of regions within the default mode network supporting perceptually-decoupled and conceptually-guided cognition. NeuroImage, 171(January), 393–401. 10.1016/j.neuroimage.2018.01.017

Noonan, K. A., Jefferies, E., Visser, M., & Lambon Ralph, M. A. (2013). Going beyond Inferior Prefrontal Involvement in Semantic Control: Evidence for the Additional Contribution of Dorsal Angular Gyrus and Posterior Middle Temporal Cortex. Journal of Cognitive Neuroscience, 25(11), 1824–1850. 10.1162/jocn_a_00442

Numssen, O., Bzdok, D., & Hartwigsen, G. (2020). Hemispheric specialization within the inferior parietal lobe across cognitive domains. ELife, March, 1–25. 10.1101/2020.07.01.181602

Olson, I. R., McCoy, D., Klobusicky, E., & Ross, L. A. (2013). Social cognition and the anterior temporal lobes: A review and theoretical framework. Social Cognitive and Affective Neuroscience. 10.1093/scan/nss119

Olson, I. R., Plotzker, A., & Ezzyat, Y. (2007). The Enigmatic temporal pole: A review of findings on social and emotional processing. In Brain (Vol. 130, Issue 7, pp. 1718–1731). Oxford University Press. 10.1093/brain/awm052

Özdem, C., Brass, M., Van der Cruyssen, L., & Van Overwalle, F. (2017). The overlap between false belief and spatial reorientation in the temporo-parietal junction: The role of input modality and task. Social Neuroscience, 12(2), 207–217. 10.1080/17470919.2016.1143027

Papinutto, N., Galantucci, S., Mandelli, M. L., Gesierich, B., Jovicich, J., Caverzasi, E., Henry, R. G., Seeley, W. W., Miller, B. L., Shapiro, K. A., & Gorno-Tempini, M. L. (2016). Structural connectivity of the human anterior temporal lobe: A diffusion magnetic resonance imaging study. Human Brain Mapping, 37(6), 2210–2222. 10.1002/hbm.23167

Patterson, K., & Lambon Ralph, M. A. (2016). The Hub-and-Spoke Hypothesis of Semantic Memory. Neurobiology of Language, 765–775. 10.1016/B978-0-12-407794-2.00061-4

Patterson, K., Nestor, P. J., & Rogers, T. T. (2007). Where do you know what you know? The representation of semantic knowledge in the human brain. In Nature Reviews Neuroscience (Vol. 8, Issue 12, pp. 976–987). 10.1038/nrn2277

Paunov, A. M., Blank, I. A., Jouravlev, O., Mineroff, Z., Gallée, J., & Fedorenko, E. (2022). Differential tracking of linguistic vs. Mental state content in naturalistic stimuli by language and theory of mind (tom) brain networks. Neurobiology of Language, 3(3), 413–440. 10.1162/nol_a_00071

Perner, J., Aichhorn, M., Kronbichler, M., Staffen, W., & Ladurner, G. (2006). Thinking of mental and other representations: the roles of left and right temporo-parietal junction. Social Neuroscience, 1(3–4), 245–258. 10.1080/17470910600989896

Pexman, P., Diveica, V., & Binney, R. J. (2021). Social Semantics : The Organisation and Grounding of Abstract Concepts. November. 10.31234/osf.io/wrbgp

Pexman, P. M., Diveica, V., & Binney, R. J. (2023). Social semantics: the organization and grounding of abstract concepts. Philosophical Transactions of the Royal Society B: Biological Sciences, 378(1870). 10.1098/rstb.2021.0363

Platek, S. M., Keenan, J. P., Gallup, G. G., & Mohamed, F. B. (2004). Where am I? The neurological correlates of self and other. Cognitive Brain Research, 19(2), 114–122. 10.1016/j.cogbrainres.2003.11.014

Plaut, D. C. (2002). Graded modality-specific specialisation in semantics: A computational account of optic aphasia. Cognitive Neuropsychology, 19(7), 603–639. 10.1080/02643290244000112

Pobric, G., Ralph, M. A. L., & Zahn, R. (2016). Hemispheric specialization within the superior anterior temporal cortex for social and nonsocial concepts. Journal of Cognitive Neuroscience, 28(3), 351–360. 10.1162/jocn_a_00902

Premack, D., & Woodruff, G. (1978). Does the chimpanzee have a theory of mind? Behavioral and Brain Sciences, 1(4), 515–526. 10.1017/S0140525X00076512

Price, C. J., Devlin, J. T., Moore, C. J., Morton, C., & Laird, A. R. (2005). Meta-analyses of object naming: Effect of baseline. Human Brain Mapping, 25(1), 70–82. 10.1002/hbm.20132

Quesque, F., & Brass, M. (2019). The Role of the Temporoparietal Junction in Self-Other Distinction. Brain Topography, 32(6), 943–955. 10.1007/s10548-019-00737-5

Ramsey, R., & Ward, R. (2020). Putting the Nonsocial Into Social Neuroscience: A Role for Domain-General Priority Maps During Social Interactions. Perspectives on Psychological Science, 15(4), 1076–1094. 10.1177/1745691620904972

Rankin, K. P. (2020). Brain Networks Supporting Social Cognition in Dementia. Current Behavioral Neuroscience Reports, 203–211. 10.1007/s40473-020-00224-3

Rankin, K. P. (2021). Measuring Behavior and Social Cognition in FTLD. In Frontotemporal Dementias (pp. 51–65). Springer.

Rice, G. E., Hoffman, P., Binney, R. J., & Lambon Ralph, M. A. (2018). Concrete versus abstract forms of social concept: An fMRI comparison of knowledge about people versus social terms. Philosophical Transactions of the Royal Society B: Biological Sciences, 373(1752), 20170136. 10.1098/rstb.2017.0136

Rice, G. E., Hoffman, P., & Lambon Ralph, M. A. (2015). Graded specialization within and between the anterior temporal lobes. Annals of the New York Academy of Sciences, 1359(1), 84–97. 10.1111/nyas.12951

Rice, G. E., Ralph, M. A. L., & Hoffman, P. (2015). The roles of left versus right anterior temporal lobes in conceptual knowledge: An ALE meta-analysis of 97 functional neuroimaging studies. Cerebral Cortex, 25(11), 4374–4391. 10.1093/cercor/bhv024

Richardson, H., & Saxe, R. (2020). Early signatures of and developmental change in brain regions for theory of mind. Neural Circuit and Cognitive Development, 467–484. 10.1016/b978-0-12-814411-4.00021-4

Ross, L. A., & Olson, I. R. (2010). Social cognition and the anterior temporal lobes. NeuroImage, 49(4), 3452–3462. 10.1016/j.neuroimage.2009.11.012

Rothmayr, C., Sodian, B., Hajak, G., Döhnel, K., Meinhardt, J., & Sommer, M. (2011). Common and distinct neural networks for false-belief reasoning and inhibitory control. NeuroImage, 56(3), 1705–1713. 10.1016/j.neuroimage.2010.12.052

Rotshtein, P., Henson, R. N. A., Treves, A., Driver, J., & Dolan, R. J. (2005). Morphing Marilyn into Maggie dissociates physical and identity face representations in the brain. Nature Neuroscience, 8(1), 107–113. 10.1038/nn1370

Rouse, M. A., Binney, R. J., Patterson, K., Rowe, J. B., & Lambon Ralph, M. A. (2024). A neuroanatomical and cognitive model of impaired social behaviour in frontotemporal dementia. Brain. 10.1093/brain/awae040

Rouse, M. A., Binney, R. J., Patterson, K., Rowe, J. B., & Ralph, A. L. (2023). A neuroanatomical and cognitive model of impaired social behaviour in frontotemporal dementia. February, 1–26.

Samson, D., Apperly, I. A., Kathirgamanathan, U., & Humphreys, G. W. (2005). Seeing it my way: A case of a selective deficit in inhibiting self-perspective. Brain, 128(5), 1102–1111. 10.1093/brain/awh464

Saxe, R. (2006). Why and how to study Theory of Mind with fMRI. Brain Research, 1079(1), 57–65. 10.1016/j.brainres.2006.01.001

Saxe, R., & Baron-Cohen, S. (2006). The neuroscience of theory of mind. In Social neuroscience (Vol. 1, Issues 3–4). 10.1080/17470910601117463

Saxe, R., & Kanwisher, N. (2003). People thinking about thinking people. The role of the temporo-parietal junction in “theory of mind”. NeuroImage, 19(4), 1835–1842. http://www.ncbi.nlm.nih.gov/pubmed/12948738

Saxe, R., & Powell, L. J. (2006). It’s the Thought That Counts They Take the Prize. Psychological Science, 324(June), 2009.

Saxe, R., & Wexler, A. (2005). Making sense of another mind: the role of the right temporo-parietal junction. Neuropsychologia, 43(10), 1391–1399. 10.1016/j.neuropsychologia.2005.02.013

Schilbach, L., Wohlschlaeger, A. M., Kraemer, N. C., Newen, A., Shah, N. J., Fink, G. R., & Vogeley, K. (2006). Being with virtual others: Neural correlates of social interaction. Neuropsychologia, 44(5), 718–730. 10.1016/j.neuropsychologia.2005.07.017

Scholz, J., Triantafyllou, C., Whitfield-Gabrieli, S., Brown, E. N., & Saxe, R. (2009). Distinct Regions of Right Temporo-Parietal Junction Are Selective for Theory of Mind and Exogenous Attention. PLoS ONE, 4(3), e4869. 10.1371/journal.pone.0004869

Schurz, M., Radua, J., Aichhorn, M., Richlan, F., & Perner, J. (2014). Fractionating theory of mind: A meta-analysis of functional brain imaging studies. Neuroscience & Biobehavioral Reviews, 42, 9–34. 10.1016/j.neubiorev.2014.01.009

Schurz, M., Radua, J., Tholen, M. G., Maliske, L., Margulies, D. S., Mars, R. B., Sallet, J., & Kanske, P. (2020). Toward a hierarchical model of social cognition: A neuroimaging meta-analysis and integrative review of empathy and theory of mind. Psychological Bulletin, No Pagination Specified-No Pagination Specified.

Schurz, M., Tholen, M. G., Perner, J., Mars, R. B., & Sallet, J. (2017). Specifying the brain anatomy underlying temporo-parietal junction activations for theory of mind: A review using probabilistic atlases from different imaging modalities. Human Brain Mapping, 38(9), 4788–4805. 10.1002/hbm.23675

Seghier, M. L. (2013). The angular gyrus: Multiple functions and multiple subdivisions. Neuroscientist, 19(1), 43–61. 10.1177/1073858412440596

Seghier, M. L. (2022). Multiple functions of the angular gyrus at high temporal resolution. Brain Structure and Function, June. 10.1007/s00429-022-02512-y

Seghier, M. L., Fagan, E., & Price, C. J. (2010). Functional Subdivisions in the Left Angular Gyrus Where the Semantic System Meets and Diverges from the Default Network. Journal of Neuroscience, 30(50), 16809–16817. 10.1523/JNEUROSCI.3377-10.2010

Sharp, D. J., Scott, S. K., & Wise, Richard, J. S. (2004). Monitoring and the Controlled Processing of Meaning: Distinct Prefrontal Systems. Cerebral Cortex, 14(1), 1–10. 10.1093/cercor/bhg086

Simmons, W. K., Reddish, M., Bellgowan, P. S. F. F., & Martin, A. (2010). The selectivity and functional connectivity of the anterior temporal lobes. Cerebral Cortex, 20(4), 813–825. 10.1093/cercor/bhp149

Smallwood, J., Tipper, C., Brown, K., Baird, B., Engen, H., Michaels, J. R., Grafton, S., & Schooler, J. W. (2013). Escaping the here and now: Evidence for a role of the default mode network in perceptually decoupled thought. NeuroImage, 69, 120–125. 10.1016/j.neuroimage.2012.12.012

Snowden, J. S., Harris, J. M., Thompson, J. C., Kobylecki, C., Jones, M., Richardson, A. M., & Neary, D. (2018). Semantic dementia and the left and right temporal lobes. Cortex, 107, 188–203. 10.1016/j.cortex.2017.08.024

Sommer, M., Sodian, B., Döhnel, K., Schwerdtner, J., Meinhardt, J., & Hajak, G. (2010). In psychopathic patients emotion attribution modulates activity in outcome-related brain areas. Psychiatry Research - Neuroimaging, 182(2), 88–95. 10.1016/j.pscychresns.2010.01.007

Souter, N., Lindquist, K. A., & Jefferies, E. (2021). Impaired emotion perception and categorization in semantic aphasia. July. 10.31234/osf.io/cy37z

Spreng, R. N., & Andrews-Hanna, J. R. (2015). The Default Network and Social Cognition. Brain Mapping: An Encyclopedic Reference, 3(December 2015), 165–169. 10.1016/B978-0-12-397025-1.00173-1

Spreng, R. N., & Grady, C. L. (2010). Patterns of brain activity supporting autobiographical memory, prospection, and theory of mind, and their relationship to the default mode network. Journal of Cognitive Neuroscience, 22(6), 1112–1123. 10.1162/jocn.2009.21282

Spreng, R. N., Mar, R. A., & Kim, A. S. N. (2009). The common neural basis of autobiographical memory, prospection, navigation, theory of mind, and the default mode: A quantitative meta-analysis. Journal of Cognitive Neuroscience, 21(3), 489–510. 10.1162/jocn.2008.21029

Spunt, R. P., & Adolphs, R. (2017). A new look at domain specificity: Insights from social neuroscience. In Nature Reviews Neuroscience (Vol. 18, Issue 9, pp. 559–567). Nature Publishing Group. 10.1038/nrn.2017.76

Sugiura, M., Sassa, Y., Watanabe, J., Akitsuki, Y., Maeda, Y., Matsue, Y., Fukuda, H., & Kawashima, R. (2006). Cortical mechanisms of person representation: Recognition of famous and personally familiar names. NeuroImage, 31(2), 853–860. 10.1016/j.neuroimage.2006.01.002

Turkeltaub, P. E., Eickhoff, S. B., Laird, A. R., Fox, M., Wiener, M., & Fox, P. (2012). Minimizing within-experiment and within-group effects in activation likelihood estimation meta-analyses. Human Brain Mapping, 33(1), 1–13. 10.1002/hbm.21186

Van der Meer, L., Groenewold, N. A., Nolen, W. A., Pijnenborg, M., & Aleman, A. (2011). Inhibit yourself and understand the other: Neural basis of distinct processes underlying Theory of Mind. NeuroImage, 56(4), 2364–2374. 10.1016/j.neuroimage.2011.03.053

Van Hoeck, N., Begtas, E., Steen, J., Kestemont, J., Vandekerckhove, M., & Van Overwalle, F. (2014). False belief and counterfactual reasoning in a social environment. NeuroImage, 90, 315–325. 10.1016/j.neuroimage.2013.12.043

Vigliocco, G., Kousta, S.-T. T., Della Rosa, P. A., Vinson, D. P., Tettamanti, M., Devlin, J. T., & Cappa, S. F. (2014). The neural representation of abstract words: The role of emotion. Cerebral Cortex, 24(7), 1767–1777. 10.1093/cercor/bht025

Visser, M., Embleton, K. V., Jefferies, E., Parker, G. J., & Ralph, M. A. L. (2010). The inferior, anterior temporal lobes and semantic memory clarified: Novel evidence from distortion-corrected fMRI. Neuropsychologia, 48(6), 1689–1696. 10.1016/j.neuropsychologia.2010.02.016

Visser, M., Jefferies, E., Lambon Ralph, M. A., Ralph, L., Visser, M., Jefferies, E., & Lambon Ralph, M. A. (2010). Semantic Processing in the Anterior Temporal Lobes : A Meta-analysis of the Functional. Journal of Cognitive Neuroscience, 22(6), 1083–1094. 10.1162/jocn.2009.21309

Visser, M., & Lambon Ralph, M. A. (2011). Differential Contributions of Bilateral Ventral Anterior Temporal Lobe and Left Anterior Superior Temporal Gyrus to Semantic Processes. Journal of Cognitive Neuroscience, 23(10), 3121–3131. 10.1162/jocn_a_00007

Younes, K., Borghesani, V., Montembeault, M., Spina, S., Mandelli, M. L., Welch, A. E., Weis, E., Callahan, P., Elahi, F. M., Hua, A. Y., Perry, D. C., Karydas, A., Geschwind, D., Huang, E., Grinberg, L. T., Kramer, J. H., Boxer, A. L., Rabinovici, G. D., Rosen, H. J., … Gorno-Tempini, M. L. (2022). Right temporal degeneration and socioemotional semantics: semantic behavioural variant frontotemporal dementia. Brain : A Journal of Neurology, 145(11), 4080–4096. 10.1093/brain/awac217

Young, L., Camprodon, J. A., Hauser, M., Pascual-Leone, A., & Saxe, R. (2010). Disruption of the right temporoparietal junction with transcranial magnetic stimulation reduces the role of beliefs in moral judgments. Proceedings of the National Academy of Sciences of the United States of America, 107(15), 6753–6758. 10.1073/pnas.0914826107

Zahn, R., Moll, J., Iyengar, V., Huey, E. D., Tierney, M., Krueger, F., & Grafman, J. (2009). Social conceptual impairments in frontotemporal lobar degeneration with right anterior temporal hypometabolism. Brain, 132(3), 604–616. 10.1093/brain/awn343

Zahn, R., Moll, J., Krueger, F., Huey, E. D., Garrido, G., & Grafman, J. (2007). Social concepts are represented in the superior anterior temporal cortex. Proceedings of the National Academy of Sciences, 104(15), 6430–6435. 10.1073/pnas.0607061104

Zaki, J., Hennigan, K., Weber, J., & Ochsner, K. N. (2010). Social cognitive conflict resolution: Contributions of domain-general and domain-specific neural systems. Journal of Neuroscience, 30(25), 8481–8488. 10.1523/JNEUROSCI.0382-10.2010

