## Supplementary_Information_No.1_Methods for "Overlapping Neural Correlates Underpin Theory of Mind and Semantic Cognition: Evidence from a Meta-Analysis of 344 Functional Neuroimaging Studies"

for

### Investigating the Similarities in Neural Networks Underpinning Theory of Mind and Semantic Cognition: A Meta-Analysis

This document contains lists with all the experiments included in the meta-analysis and their characteristics and associated references. The complete raw datasets can be accessed via OSF (https://osf.io/ydnxh/).

#### *Semantic Cognition Data Sets and Subsets*

***Table M1 SC*** *List of studies included in the* ***semantic cognition all baselines*** *(N = 214) meta-analysis.* ***Note:*** *SC= Semantic Cognition, N= Sample Size*

| *Authors | Year | Imaging Method | Social Content | Instructional Cue | Stimulus Domain | Sensory Input Modality | Baseline Type | N | Contrast | Age Range |
|---|---|---|---|---|---|---|---|---|---|---|
| AbdulSabur et al. | 2014 | fMRI | NO | EXPLICIT | VERBAL | VISUAL | LOW | 18 | narrative production>recitation | |
| Abraham et al. | 2012 | fMRI | NO | EXPLICIT | VERBAL | VISUAL | HIGH | 19 | high&low divergent thinking (semantic)>1&2 back letter identity (working memory) | 22.42 (19-29) |
| Alain, He & Grady | 2008 | fMRI | NO | EXPLICIT | VERBAL | AUDITORY | HIGH/REST | 16 | sound category>sound location | 26 (21-31) |
| | | | | | | | | | sound category>rest | |
| Assadollahi, Meinzer, Flaisch, Obleser & Rockstroh | 2009 | fMRI | NO | IMPLICIT | VERBAL | VISUAL | HIGH/REST | 20 | nouns followed by 1&3 argument verbs>letter strings followed by 1&3argument verbs | 27.1 (?-?) |
| | | | | | | | | | nouns followed by 1 argument verbs>fixation cross (rest) | |
| | | | | | | | | | nouns followed by 3 argument verbs>fixation cross (rest) | |
| Axmacher, Bialleck, Weber, Helmstaedter, Elger & Fell | 2009 | fMRI | NO | EXPLICIT | VERBAL | VISUAL | HIGH | 32 | word decision>spatial decision | 27.7 (?-?) |
| Bagga et al. | 2013 | fMRI | NO | EXPLICIT | VERBAL | VISUAL | HIGH | 18 | semantic>case matching judgement | 35.25 (?-?) |
| Barros-Loscertales et al. | 2012 | fMRI | NO | EXPLICIT | VERBAL | VISUAL | LOW | 59 | control words>hashmarks baseline | 22.51 (17-37) |
| Baumgaertener, Weiller & Buchel | 2002 | fMRI | NO | EXPLICIT | VERBAL | VISUAL | HIGH | 9 | word>pseudoword in sentence | 29.4 (23-49) |
| Baumgaertner et al. | 2007 | fMRI | NO | IMPLICIT | VERBAL/NONVERBAL | AUDITORY/VISUAL | HIGH | 19 | sentences>reversed sentences | 27.3 (20-41) |
| | | | | | | | | | videos>scrambled videos | |
| Bautista & Wilson | 2016 | fMRI | NO | IMPLICIT | VERBAL | AUDITORY | HIGH | 12 | clear>scrambled rotated speech | 21.4 (20-28) |
| Bick, Goelman & Frost | 2008 | fMRI | NO | EXPLICIT | VERBAL | VISUAL | LOW | 14 | semantic>visual control | 28.3 (?-?) |
| | | | | | | | | | morphologiacl>visual control | |

| | | | | | | | | | | |
|---|---|---|---|---|---|---|---|---|---|---|
| | | | | | | | | | orthographic>visual control | |
| | | | | | | | | | phonological>visual control | |
| Binder, Frost, Hammeke, Bellgowan, Rao & Cox | 1999 | fMRI | NO | EXPLICIT | VERBAL | AUDITORY | HIGH | 30 | semantic>phonological decision | ? (18-35) |
| Binder et al. | 2003 | fMRI | NO | EXPLICIT | VERBAL | VISUAL | HIGH | 24 | word>nonword | 25.2 (18-49) |
| Birn et al. | 2010 | fMRI | NO | EXPLICIT | VERBAL | VISUAL | LOW/HIGH | 14 | category & letter fluency>months ( automatic speech) | 32.2 (22-48) |
| | | | | | | | | | category>letter fluency | |
| Bonhage, Fiebach, Bahlmann & Mueller | 2014 | fMRI | NO | EXPLICIT | VERBAL | VISUAL | HIGH | 18 | sentence fragments>ungrammatical word strings | 25 (20-31) |
| Bonhage, Mueller, Friederici & Fiebach | 2015 | fMRI | NO | EXPLICIT | VERBAL | VISUAL | HIGH | 18 | sentences>jabberwocky sentences | 26.6 (2.8) |
| Booth et al. | 2006 | fMRI | NO | EXPLICIT | VERBAL | VISUAL | HIGH/LOW | 13 | meaning (semantic) judgement>rhyming( phonological) judgement | 22.3 (?-?) |
| | | | | | | | | | meaning(semantic)>control(symbols) | |
| Boulenger, Hauk & Pulvermuller | 2009 | fMRI | NO | EXPLICIT | VERBAL | VISUAL | LOW | 18 | sentences> hashmarks | 24.3 (?-?) |
| | | | | | | | | | sentences> hashmarks | |
| Bozic & Marslen-Wilson | 2013 | fMRI | NO | IMPLICIT | VERBAL | AUDITORY | HIGH | 13 | speech>musical rain | ? (?-?) |
| Brambati, Benoit, Monetta, Belleville & Joubert | 2010 | fMRI | YES | EXPLICIT | BOTH | BOTH | HIGH | 12 | general & specific occupation judgement>baseline (scrambled face) | 23 (20-26) |
| Bruffaerts, Dupont, Peeters, De Deyne, Storms & Vandenberghe | 2013 | fMRI | NO | EXPLICIT | BOTH | VISUAL | HIGH | 19 | real > scrambled pictures & words | ? (19-26) |
| Bulut, Hung, Tzeng & Wu | 2017 | fMRI | NO | IMPLICIT | VERBAL | VISUAL | HIGH | 20 | sentences>unstructured word lists | 23 (19-29) |
| | | | | | | | | | sentences>unstructured character list | |
| Cai, Kochiyama, Osaka & Wu | 2007 | fMRI | NO | IMPLICIT | VERBAL | AUDITORY | HIGH/REST | 15 | words>nonsense words | ? (21-23) |
| | | | | | | | | | words>rest | |

| | | | | | | | | | |
|---|---|---|---|---|---|---|---|---|---|
| Cao, Peng, Liu, Jin, Fan, Deng, & Booth | 2009 | fMRI | NO | EXPLICIT | VERBAL | VISUAL | LOW | 13 | meaning>perceptual decision | 22.3 (20-30) |
| Cappa, Perani, Schnur, Tettamanti & Fazio | 1998 | PET | NO | EXPLICIT | VERBAL | VISUAL | HIGH | 13 | words>pseudowords | ? (22-26) |
| | | | | | | | | | animal visual knowledge>pseudowords | |
| | | | | | | | | | tool visual knowledge>pseudowords | |
| | | | | | | | | | animal associative knowledge>pseudowords | |
| | | | | | | | | | tools functional knowledge>pseudowords | |
| Carota, Kriegeskorte, Nili & Pulvermuller | 2017 | fMRI | NO | IMPLICIT | VERBAL | VISUAL | LOW | 23 | words>hashmarks | 29 (?-?) |
| Carota, Moseley & Pulvermueller | 2012 | fMRI | NO | IMPLICIT | VERBAL | VISUAL | LOW | 18 | all words>hashmarks | 29 (?-?) |
| | | | | | | | | | tool words>hashmarks | |
| | | | | | | | | | animal words>hashmarks | |
| | | | | | | | | | food words>hashmarks | |
| Chan, Tang, Tang, Lee, Lo & Kwong | 2009 | fMRI | NO | IMPLICIT | VERBAL | VISUAL | HIGH/REST | 22 | synonyms>pseudocharacters | ? (19-30) |
| | | | | | | | | | synonyms>Korean characters (unknown) | |
| | | | | | | | | | synonyms>rest | |
| Chiao, Harada, Oby, Li, Parrish & Bridge | 2009 | fMRI | YES | EXPLICIT | NONVERBAL | VISUAL | LOW | 12 | uniform status judgement>colour change detection | 20.7 (?-?) |
| | | | | | | | | | face status judgement>colour change detection | |
| | | | | | | | | | car status judgement>colour change detection | |
| Chou, Chen, Wu & Booth | 2009 | fMRI | NO | EXPLICIT | VERBAL | VISUAL | HIGH | 31 | related words>false font | 20.9 (?-?) |
| | | | | | | | | | unrelated words>false font | |
| Chou, Chen, Wu & Booth | 2009 | fMRI | NO | EXPLICIT | VERBAL | VISUAL | HIGH | 32 | related words>false font | |
| | | | | | | | | | unrelated words>false font | |

| | | | | | | | | | | |
|---|---|---|---|---|---|---|---|---|---|---|
| Chouinard, Morrissey, Kohler & Goodale | 2008 | fMRI | NO | EXPLICIT | NONVERBAL | VISUAL | HIGH | 14 | objects>scrambled objects | ? (?-?) |
| Chow, Kaup, Raabe & Greenlee | 2008 | fMRI | NO | EXPLICIT | VERBAL | VISUAL | HIGH | 15 | predictive&normal reading>pseudoword reading | 24 (?-?) |
| | | | | | | | | | normal reading>pseudoword reading | |
| | | | | | | | | | predictive reading>pseudoword reading | |
| Christensen, Antonucci, Lockwood, Kittleson & Plante | 2008 | fMRI | NO | EXPLICIT | VERBAL | AUDITORY | HIGH | 14 | diotic listening>reversed speech | ? (18-49) |
| | | | | | | | | | dichotic listening>reversed speech | |
| Clos, Langner, Meyer, Oechslin, Zilles & Eickhoff | 2014 | fMRI | NO | EXPLICIT | VERBAL | AUDITORY | HIGH | 29 | intelligibilty based on cue > unintelligible | 34.5 (?-?) |
| Damasio, Grabowski, Tranel, Ponto, Hichwa & Damasio | 2001 | PET | NO | EXPLICIT | NONVERBAL | VISUAL | LOW | 10 | actions without implement>control task | ? (23-55) |
| | | | | | | | | | actions with implement>control task | |
| Damasio, Tranel, Grabowski, Adolphs & Damasio | 2004 | PET | YES | EXPLICIT | NONVERBAL | VISUAL | HIGH | 55 | persons>face orientation judgement | ? (?-?) |
| | | | | | | | | | animals>scrambled pictures | |
| | | | | | | | | | tools>scrambled pictures | |
| Dapretto & Bookheimer | 1999 | fMRI | NO | EXPLICIT | VERBAL | AUDITORY | REST | 8 | semantic>rest | |
| D'Arcy, Bolster, Ryner, Mazerolle, Grant & Song | 2007 | fMRI | NO | EXPLICIT | BOTH | VISUAL | REST | 10 | basic-level living objects>baseline | 35.6 (?-?) |
| | | | | | | | | | basic-level non-living objects>baseline | |
| | | | | | | | | | superordinate-level living objects>baseline | |
| | | | | | | | | | superordinate-level non-living objects>baseline | |
| Davis, Meunier & Marslen-Wilson | 2004 | fMRI | NO | EXPLICIT | VERBAL | VISUAL | HIGH | 11 | words>letter strings | ? (18-40) |

| | | | | | | | | | | |
|---|---|---|---|---|---|---|---|---|---|---|
| Davis, Ford, Kherif & Johnsrude | 2011 | fMRI | NO | EXPLICIT | VERBAL | AUDITORY | LOW | 12 | clear & anomalous sentences>signal correlated noise | ? (18-45) |
| Demonet et al. | 1992 | PET | NO | EXPLICIT | VERBAL | AUDITORY | HIGH/LOW | 9 | words>phonemes | 35.7 (?-?) |
| | | | | | | | | | words>tones | |
| Devlin, Matthews & Rushworth | 2003 | fMRI | NO | EXPLICIT | VERBAL | VISUAL | HIGH | 12 | semantic>phonological judgement | ? (21-33) |
| Devlin et al. | 2002 | PET | NO | EXPLICIT | VERBAL | VISUAL | HIGH | 12 | all semantic>letter detection | 30 (21-51) |
| | | | | | | | | | all semantic>letter detection | |
| Devlin et al. | 2002 | PET | NO | EXPLICIT | VERBAL | VISUAL | HIGH | 8 | all semantic>letter categorisation | 28 (21-47) |
| Devlin et al. | 2002 | fMRI | NO | EXPLICIT | VERBAL | VISUAL | HIGH | 8 | all semantic> letter categorisation | 28 (21-47) |
| Devlin et al. | 2000 | PET | NO | EXPLICIT | VERBAL | VISUAL | HIGH | 8 | semantic categorisation>letter categorisation | 28 (21-47) |
| Devlin et al. | 2000 | fMRI | NO | EXPLICIT | VERBAL | VISUAL | HIGH | 8 | semantic categorisation>letter categorisation | |
| Diaz & McCarthy | 2009 | fMRI | NO | IMPLICIT | VERBAL | VISUAL | HIGH | 16 | all words>nonwords | 22.25 (19-34) |
| Dreyer & Pulvermueller | 2018 | fMRI | YES | IMPLICIT | VERBAL | VISUAL | LOW | 28 | all nouns>hashmarks | 23.7 (?-?) |
| | | | | | | | | | abstract emotional nouns>baseline (hashmarks) | |
| | | | | | | | | | abstract mental nouns>baseline (hashmarks) | |
| | | | | | | | | | food nouns>baseline (hashmarks) | |
| | | | | | | | | | tool nouns>baseline (hashmarks) | |
| Ebisch et al. | 2007 | fMRI | NO | EXPLICIT | BOTH | VISUAL | HIGH | 17 | functional&visuospatial pictures&words>meaningless drawings&pseudowords | 21.7 (?-?) |
| Elfgren, Westen, Passant, Larsson, Mannfolk & Fransson | 2006 | fMRI | YES | EXPLICIT | NONVERBAL | VISUAL | HIGH | 15 | familiar>unfamiliar faces (identification) | 23.3 (19-32) |
| Emmorey, Weisberg, McCullough & Petrich | 2013 | fMRI | NO | EXPLICIT /IMPLICI T | VERBAL | VISUAL | HIGH | 14 | words(semantic judgement)>false fonts | 25.4 (?-?) |
| | | | | | | | | | words(phonological)>false fonts | |

| | | | | | | | | | | |
|---|---|---|---|---|---|---|---|---|---|---|
| Emmorey, Xu, Gannon, Goldin-Meadow & Braun | 2010 | fMRI | NO | IMPLICIT | NONVERBAL | VISUAL | HIGH/REST | 14 | meaningful pantomimes>unknown sign language | 22.3 (19-43) |
| | | | | | | | | | meaningful pantomimes>rest | |
| Engelien et al. | 2006 | PET | NO | IMPLICIT | NONVERBAL | AUDITORY | HIGH/REST | 6 | meaningful>meaningless sounds | 34.5 (?-?) |
| | | | | | | | | | meaningful>rest | |
| Erb, Henry, Eisner & Obleser | 2013 | fMRI | NO | EXPLICIT | VERBAL | AUDITORY | HIGH | 30 | speech>vocoded speech | 25.9 (21-31) |
| Europa, Gitelman, Kiran & Thompson | 2019 | fMRI | NO | EXPLICIT | BOTH | BOTH | HIGH | 21 | sentences>baseline (reversed sentences) | 36.3 (24-67) |
| Foki, Gartus, Geissler & Beisteiner | 2008 | | | EXPLICIT | VERBAL | VISUAL | LOW | 23 | semantic judgement>tongue movements | 31 (?-?) |
| Friederici, Kotz, Scott & Obleser | 2010 | fMRI | NO | IMPLICIT | VERBAL | AUDITORY | HIGH | 17 | intelligible speech>rotated speech | ? (20-30) |
| Friese, Rutschmann, Raabe & Schmalhofer | 2008 | fMRI | NO | EXPLICIT | VERBAL | VISUAL | HIGH | 13 | words>pseudowords | 22.8 (?-?) |
| Garbin, Collina & Tabossi | 2012 | fMRI | NO | EXPLICIT | VERBAL | VISUAL | HIGH | 12 | object noun>pseudoword | 25 (21-29) |
| | | | | | | | | | event noun>pseudoword | |
| | | | | | | | | | verb>pseudoword | |
| Garn, Allen & Larsen | 2009 | fMRI | NO | EXPLICIT | NONVERBAL | VISUAL | HIGH | 26 | pictures>scrambled pictures | 23.6 (18-30) |
| | | | | | | | | | plants>scrambled pictures | |
| | | | | | | | | | tools>scrambled pictures | |
| Geranmayeh, Brownsett, Leech, Beckmann, Woodhead & Wise | 2012 | fMRI | NO | EXPLICIT | VERBAL | VISUAL | HIGH | 19 | speech>tongue movements | 30 (22-62) |
| Gerlach, Law, Gade & Paulson | 1999 | PET | NO | EXPLICIT | NONVERBAL | VISUAL | HIGH | 15 | object decision>pattern discrimination | 26 (22-30) |
| Gesierich et al. | 2012 | fMRI | YES | EXPLICIT | NONVERBAL | VISUAL | HIGH | 21 | familiar>scrambled faces | 28.4 (19-49) |
| | | | | | | | | | familiar > unfamiliar faces | |
| | | | | | | | | | familiar>unfamiliar faces | |
| Giraud & Price | 2001 | PET | NO | BOTH | BOTH | AUDITORY | BOTH | 12 | words+environmental sounds>syllables+noise | 36.6 (?-?) |
| Giraud et al. | 2004 | fMRI | NO | EXPLICIT | VERBAL | AUDITORY | HIGH | 8 | natural speech>speech envelope | 28.4 (24-38) |
| Gitelman, Nobre, Sonty, Parrish & Mesulam | 2005 | fMRI | NO | EXPLICIT | VERBAL | VISUAL | HIGH | 14 | semantic>control task | 29.9 (?-?) |

| | | | | | | | | | | |
|---|---|---|---|---|---|---|---|---|---|---|
| Gorno-Tempini et al. | 1998 | PET | YES | EXPLICIT | NONVERBAL /VERBAL | VISUAL | HIGH | 6 | famous faces>controls | ? (18-33) |
| | | | | | | | | | famous names>controls | |
| | | | | | | | | | double famous proper names>controls | |
| | | | | | | | | | non-famous faces>controls | |
| | | | | | | | | | Non-famous names>controls | |
| | | | | | | | | | Double common names>controls | |
| Grabowski, Damasio, Tranel, Boles Ponto, Hichwa & Damasio | 2001 | PET | YES | EXPLICIT | NONVERBAL | VISUAL | HIGH | 10 | naming persons>building orientation judgement | 28.8 (?-?) |
| | | | | | | | | | naming landmarks>face orientation judgement | |
| | | | | | | | | | naming persons>face orientation judgement | |
| | | | | | | | | | naming unique entities>baseline | |
| Graves, Binder, Desai, Conant & Seidenberg | 2010 | fMRI | NO | EXPLICIT | VERBAL | VISUAL | HIGH | 23 | forward>reverse phrases | 24.2 (?-?) |
| Graves, Binder, Desai, Conant & Seidenberg | 2010 | fMRI | NO | EXPLICIT | VERBAL | VISUAL | HIGH | 22 | forward>reverse phrases | 24.2 (?-?) |
| Grindrod, Garnett, Malyutina & den Ouden | 2014 | fMRI | NO | EXPLICIT | VERBAL | VISUAL | HIGH | 23 | words>nonwords | 23 (19-32) |
| Grossman et al. | 2002 a | fMRI | NO | IMPLICIT | VERBAL | VISUAL | HIGH | 16 | all nouns > pseudowords | 23.4 (?-?) |
| | | | | | | | | | implements>pseudowords | |
| | | | | | | | | | animals>pseudowords | |
| | | | | | | | | | abstract>pseudowords | |
| Grossman et al., 2002b | 2002 b | fMRI | NO | IMPLICIT | VERBAL | VISUAL | HIGH | 16 | verbs>pseudowords | 23.4 (19-28) |
| Groussard et al. | 2010 | PET | NO | EXPLICIT | VERBAL/NO NVERBAL | VISUAL/AU DITORY | HIGH/REST | 11 | verbal semantics>verbal reference | 23.6 (20-27) |
| | | | | | | | | 11 | musical semantic>musical reference | |
| | | | | | | | | 11 | verbal semantic>rest | |
| | | | | | | | | 11 | verbal semantic>rest | |

| | | | | | | | | | | |
|---|---|---|---|---|---|---|---|---|---|---|
| Guediche, Reilly, Santiago, Laurent & Blumstein | 2016 | fMRI | NO | IMPLICIT | VERBAL | AUDITORY | LOW | 16 | related > repeated sentence | 24.1 (18-34) |
| | | | | | | | | | unrelated > repeated sentence | |
| Gurd et al. | 2002 | fMRI | NO | EXPLICIT | VERBAL | AUDITORY | HIGH | 11 | category>rote fluency | 32 (?-?) |
| Haberling, Corballis &Corballis | 2016 | fMRI | NO | EXPLICIT | NONVERBAL /VERBAL | VISUAL | HIGH | 92 | meaningful pantomimes>unknown sign language | ? (17-35) |
| | | | | | | | | | meaningful pantomimes>dog videos | |
| | | | | | | | | | synonyms>letter strings | |
| Hagoort et al. | 1999 | PET | NO | IMPLICIT | VERBAL | VISUAL | HIGH/REST | 10 | words>pseudowords | 26.8 (25-30) |
| | | | | | | | | | words>fixation | |
| Harrington, Farias & Davis | 2009 | fMRI | NO | EXPLICIT | NONVERBAL | VISUAL | HIGH/REST | 8 | familiar>non objects | 32.8 (20-52) |
| | | | | | | | | | familiar objects>rest | |
| Hartung, Hagoort & Willems | 2017 | fMRI | NO | IMPLICIT | VERBAL | AUDITORY | HIGH | 52 | first-person speech>unintelligible reversed speech | 23.6 (18-35) |
| | | | | | | | | | third-person speech>unintelligible reversed speech | |
| Hauk & Pulvermueller | 2011 | fMRI | NO | IMPLICIT | VERBAL | VISUAL | LOW | 21 | action words>hashmarks | 25.8 (?-?) |
| | | | | | | | | | uni manual action words>hashmarks | |
| | | | | | | | | | uni manual action words>hashmarks | |
| Hayashi et al. | 2014 | fMRI | NO | EXPLICIT | VERBAL | VISUAL | LOW | 16 | concrete word>astersisks | 21.6 (20-36) |
| | | | | | | | | | abstract word>asterisks | |
| Heim, Eickhoff & Amunts | 2008 | fMRI | NO | EXPLICIT | VERBAL | VISUAL | HIGH/REST | 28 | semantic > phonological fluency | 29.4 (?-?) |
| | | | | | | | | | semantic fluency>rest | |
| Henke et al. | 1999 | PET | NO | EXPLICIT | VERBAL | VISUAL | HIGH | 12 | associative word learning>single word encoding | 19-30(23) |
| Herbster et al. | 1997 | PET | NO | EXPLICIT | VERBAL | VISUAL | HIGH | 10 | Irregular>zero order speak | 28.4 (12-18) |
| | | | | | | | | | regular>zero order speak | |

| | | | | | | | | | | |
|---|---|---|---|---|---|---|---|---|---|---|
| | | | | | | | | | irregular+regular>zero order speak | |
| Hervais-Adelman, Carlyon, Johnsrude & Davis | 2012 | fMRI | NO | EXPLICIT | VERBAL | AUDITORY | HIGH | 15 | clear>vocoded speech (uninteligible) | ? (18-35) |
| Higuchi, Moriguchi, Murakami, Katsunuma, Mishima &Uno | 2015 | fMRI | NO | IMPLICIT | VERBAL | VISUAL | LOW | 28 | all characters>checkerboard | 21.7 (18-28) |
| Hocking, McMahon & de Zubicaray | 2011 | fMRI | NO | EXPLICIT | NONVERBAL | AUDITORY | HIGH | 13 | all environmental sounds>perceptual baseline | 26.5 (20-34) |
| Holle, Gunter, Rueschemeyer, Hennenlotter & Iacoboni | 2008 | fMRI | NO | EXPLICIT | BOTH | BOTH | HIGH | 17 | iconic gesture of dominant meaning>grooming | 25.7 (21-30) |
| | | | | | | | | | iconic gesture of subordinate meaning>grooming | |
| Homae, Yahata & Sakai | 2003 | fMRI | NO | EXPLICIT | VERBAL | AUDITORY /VISUAL | HIGH | 10 | auditory sentences>auditory non words | ? (20-27) |
| | | | | | | | | | visual sentences>visual non words words | |
| Husain, Patkin, Kim, Braun & Horwitz | 2012 | fMRI | NO | EXPLICIT | NONVERBAL | VISUAL | HIGH | 16 | meaningful iconic>meaningless gestures | 30 (?-?) |
| Hwang, Palmer, Basho, Zadra & Muller | 2009 | fMRI | NO | EXPLICIT | VERBAL | AUDITORY | LOW | 13 | fluency generation>production baseline | 25.8 (21-37) |
| Ikuta et al. | 2006 | fMRI | NO | IMPLICIT | VERBAL | VISUAL | LOW | 34 | sentences>word lists | ? (18-33) |
| Jackson, Hoffman, Pobric, Lambon Ralph | 2015 | fMRI | NO | EXPLICIT | VERBAL | VISUAL | HIGH | 24 | words>letter strings | 25.48 (20-42) |
| Jensen, Hargreaves, Bass, Pexman, Goodyear & Federico | 2011 | fMRI | NO | EXPLICIT | VERBAL | VISUAL | HIGH/REST | 12 | words>pseudowords | ? (?-?) |
| | | | | | | | | | words>rest | |
| Jeon, Lee, Kim & Cho | 2009 | fMRI | NO | EXPLICIT | VERBAL | VISUAL | HIGH/REST | 16 | synonys>nonwords | 23 (21-30) |
| | | | | | | | | | antonyms>nonwords | |
| | | | | | | | | | English synonyma>nonwords | |
| | | | | | | | | | Korean synonyms>nonwords | |
| | | | | | | | | | honorific words>nonwords | |
| | | | | | | | | | synonyms>rest | |
| | | | | | | | | | antonyms>rest | |

| | | | | | | | | | | |
|---|---|---|---|---|---|---|---|---|---|---|
| | | | | | | | | | English synonyms>rest | |
| | | | | | | | | | Korean synonyms>nonwords | |
| | | | | | | | | | honorific words>nonwords | |
| Joubert et al. | 2004 | fMRI | NO | IMPLICIT | VERBAL | VISUAL | HIGH | 10 | low frequency words>nonwords | 26 (?-?) |
| | | | | | | | | | high frequency words>consonant strings | |
| | | | | | | | | | low frequency words>consonant strings | |
| Kang et al. | 2006 | PET | NO | EXPLICIT | VERBAL | BOTH/AUDITORY/VISUAL | HIGH/LOW | 17 | audio-visual speech>noises and facial movements | 24.9 (20-34) |
| | | | | | | | | | auditory speech>white noise | |
| | | | | | | | | | visual speech>facial movements (chewing gum) | |
| Khader, Jost, Mertens, Bien & Roesler | 2010 | fMRI | NO | EXPLICIT | VERBAL | VISUAL | HIGH | 16 | noun>rhyme generation | 22.7 (19-26) |
| | | | | | | | | | verb>rhyme generation | |
| | | | | | | | | | noun generation>letter detection | |
| | | | | | | | | | verb generation>letter detection | |
| Kim et al. | 2009 | fMRI | NO | EXPLICIT | VERBAL | VISUAL | HIGH/REST | 36 | sentences>word lists | 20.8 (?-?) |
| | | | | | | | | | sentences>rest | |
| Kinno, Kawamura, Shioda & Sakai | 2008 | fMRI | NO | EXPLICIT | BOTH | VISUAL | HIGH | 14 | canonical sentence>picture & letter strings | ? (20-31) |
| | | | | | | | | | active sentence>picture & letter strings | |
| | | | | | | | | | passive sentence>picture & letter strings | |
| Kotz, Cappa, Von Cramon & Friederici | 2002 | fMRI | NO | EXPLICIT | VERBAL | AUDITORY | HIGH | 13 | words>pseudowords | 23.5 (?-?) |
| Kuchinke et al. | 2005 | fMRI | YES | IMPLICIT | VERBAL | VISUAL | HIGH | 20 | emotion word>nonword | 26.3 (20-36) |
| Kumar | 2016 | fMRI | NO | EXPLICIT | VERBAL | VISUAL | HIGH | 20 | abstract+concrete words>pseudowords | 28.3 (?-?) |
| | | | | | | | | | abstract words>pseudowords | |
| Kuperberg et al. | 2000 | fMRI | NO | EXPLICIT | VERBAL | AUDITORY | HIGH | 4 | sentences>words strings | ? (?-?) |

| | | | | | | | | | | |
|---|---|---|---|---|---|---|---|---|---|---|
| Kyong, Scott, Rosen, Howe, Agnew & McGettigan | 2014 | fMRI | NO | IMPLICIT | VERBAL | AUDITORY | LOW | 19 | intelligible vocoded>inverted vocoded speech | ? (18-40) |
| Leff, Schofield, Stephan, Crinion, Friston & Price | 2008 | fMRI | NO | IMPLICIT | VERBAL | AUDITORY | HIGH | 26 | speech>reversed speech | 27.3 (21-38) |
| Leung & Alain | 2011 | fMRI | NO | EXPLICIT | NONVERBAL | AUDITORY | HIGH | 16 | semantic>location matching | 25.19 (18-30) |
| Leveroni et al. | 2000 | fMRI | YES | EXPLICIT | NONVERBAL | VISUAL | HIGH | 11 | familiar faces>foils(never seen faces) | 32.0 (25-36) |
| | | | | | | | | | newly learned faces>foils(never seen faces) | |
| Lin, Wang, Zhao, Liu, Li & Bi | 2015 | fMRI | NO | EXPLICIT | VERBAL | VISUAL | HIGH | 20 | words>pseudowords | 22.5 (?-?) |
| Liu et al. | 2009 | fMRI | NO | EXPLICIT | VERBAL | VISUAL/AUDITORY | LOW | 16 | words meaning>slashes | 22.8 (19.2-24.9) |
| | | | | | | | | | words rhyming>slashes | |
| | | | | | | | | | words meaning>tones | |
| | | | | | | | | | words rhyming>tones | |
| Liuzzi et al. | 2017 | fMRI | NO | EXPLICIT | VERBAL | BOTH | HIGH | 18 | semantic judgement > input modality detection | ? (18-28) |
| Ludersdorfer, Wimmer, Richlan, Schurz, Hutzler & Kronbichler | 2016 | fMRI | NO | EXPLICIT | VERBAL | AUDITORY | LOW/REST | 29 | words orthographic>tones | 26 (18-35) |
| | | | | | | | | | words semantic>tones | |
| | | | | | | | | | Auditory words orthographic>rest | |
| | | | | | | | | | Auditory words semantic>rest | |
| Ludersdorfer, Schurz, Richlan, Kronbichler & Wimmer | 2013 | fMRI | NO | EXPLICIT | VERBAL | VISUAL | HIGH | 29 | words>false fonts | 24.3 (19-48) |
| | | | | | | | | | words>pseudowords | |
| | | | | | | | | | speech>reversed speech | |
| | | | | | | | | | words>pseudowords | |
| Malins, Gumkowski, Buis, Molfese, Rueckl, Frost, Pugh, Morris & Mencl | 2016 | fMRI | NO | IMPLICIT | VERBAL | VISUAL | LOW/HIGH | 18 | unrelated words>false font | 24 (?-?) |

| | | | | | | | | | | |
|---|---|---|---|---|---|---|---|---|---|---|
| | | | | | | | | | unrelated>pseudowords | |
| Marques, Canessa & Cappa | 2009 | fMRI | NO | EXPLICIT | VERBAL | VISUAL | LOW | 21 | sentences>crosses | 26.09 (24-29) |
| Marques, Canessa, Siri, Catricala & Cappa | 2008 | fMRI | NO | EXPLICIT | VERBAL | VISUAL | LOW | 21 | semantic features>baseline task | 26.09 (24-29) |
| Mashal, Vishne, Laor & Titone | 2013 | fMRI | NO | EXPLICIT | VERBAL | VISUAL | HIGH | 14 | novel metaphor>unrelated words | ? (20-40) |
| | | | | | | | | | conventional metaphor>unrelated words | |
| Matchin, Liao, gaston & Lau | 2019 | fMRI | NO | EXPLICIT | VERBAL | VISUAL | HIGH/REST | 20 | verb phrase>list | 22 (18-27) |
| | | | | | | | | | noun phrase>list | |
| | | | | | | | | | sentence>rest | |
| Matchin, Hammerly & Lau | 2017 | fMRI | NO | EXPLICIT | VERBAL | VISUAL | HIGH | 16 | sentences>word lists | 24 (20-29) |
| | | | | | | | | | sentences>phrases | |
| | | | | | | | | | real>pseudoword lists | |
| | | | | | | | | | real>pseudoword phrases | |
| | | | | | | | | | real>pseudoword sentences | |
| Mellem, Jasmin, Peng & Martin | 2016 | fMRI | NO | IMPLICIT | VERBAL | VISUAL | LOW | 20 | longer>shorter phrase | 26 (?-?) |
| Menz, Blangero, Kunze & Binkofski | 2010 | fMRI | NO | EXPLICIT | NONVERBAL | VISUAL | HIGH | 20 | known>unknown objects | 26.6 (21-35) |
| Meyer, Alter, Friederici, Lohmann & Yves von Cramon | 2002 | fMRI | NO | IMPLICIT | VERBAL | AUDITORY | HIGH | 14 | word>pseudoword sentence | 25.2 (?-?) |
| Moberget, Gullesen, Andersson, Ivry & Endestad | 2014 | fMRI | NO | EXPLICIT | VERBAL | VISUAL | HIGH | 32 | incongruent>scrambled sentence | 26.2 (?-?) |
| | | | | | | | | | congruent>scrambled sentence | |
| Moseley, Carota, Hauk, Mohr & Pulvermueller | 2012 | fMRI | YES | IMPLICIT | VERBAL | VISUAL | LOW | 18 | all emotional words>hashmarks | 29 (?-?) |
| | | | | | | | | | abstract emotional words>hashmarks | |
| | | | | | | | | | arm+face+emotion words>hashmarks | |
| | | | | | | | | | face words>hashmarks | |

| | | | | | | | | | | |
|---|---|---|---|---|---|---|---|---|---|---|
| | | | | | | | | | arm words>hashmarks | |
| Mummery, Patterson, Hodges & Price | 1998 | PET | NO | EXPLICIT | VERBAL | VISUAL | HIGH | 10 | semantic>phonological decision | ? (25-31) |
| Nakamura et al. | 2000 | PET | YES | EXPLICIT | NONVERBAL | VISUAL | LOW | 7 | familiar faces>fixation cross | ? (23-29) |
| Nakamura et al. | 2001 | PET | YES | EXPLICIT | VERBAL | AUDITORY | HIGH | 9 | familiar voice>vowel discrimination | ? (20-34) |
| | | | | | | | | | self voice>vowel discrimination | |
| Nichelli, Grafman, Pietrini, Clark, Lee & Miletich | 1995 | PET | NO | EXPLICIT | VERBAL | VISUAL | HIGH | 9 | semantic>orthographic decision | ? (?-?) |
| Nielson et al. | 2010 | fMRI | YES | EXPLICIT | BOTH | VISUAL | HIGH | 17 | familiar>unfamiliar people | 28.8 (20-47) |
| Noppeney & Price | 2003 | PET | NO | EXPLICIT | VERBAL | AUDITORY | LOW | 9 | normal>reversed words | 23 (20-30) |
| Orfanidou, Marlsen-Wilson & Davis | 2006 | fMRI | NO | EXPLICIT | VERBAL | AUDITORY | HIGH | 13 | words>pseudowords | ? (18-40) |
| Pallier, Devauchelle & Dehaene | 2011 | fMRI | NO | EXPLICIT | VERBAL | VISUAL | HIGH | 40 | longer>shorter phrase | 24 (18-37) |
| | | | | | | | | | length of real>pseudoword sentences | |
| Peelle, Eason, Schmitter, Schwarzbauer & Davis | 2010 | fMRI | NO | EXPLICIT | VERBAL | AUDITORY | HIGH | 6 | sentences>signal correlated noise | ? (20-26) |
| Perani, Schnur, Tettamanti, Gorno-Tempini, Cappa & Fazio | 1999 | PET | NO | EXPLICIT | NONVERBAL | VISUAL | HIGH | 11 | living objects>shapes | ? (24-32) |
| | | | | | | | | | nonliving objects>shapes | |
| Perani, Schnur, Tettamanti, Gorno-Tempini, Cappa & Fazio | 1999 | PET | NO | EXPLICIT | VERBAL | VISUAL | HIGH | 8 | living words>pseudowords | ? (22-26) |
| | | | | | | | | | nonliving words>pseudowords | |
| Perrone-Bertolotti, Kauffmann, Pichat, Vidal & Baciu | 2017 | fMRI | NO | EXPLICIT | VERBAL | VISUAL | HIGH/REST | 24 | words>unreadable font | 26.91 (19-33) |
| | | | | | | | | | words>fixation | |
| Pilgrim, Fadili, Fletcher & Tyler | 2002 | fMRI | NO | EXPLICIT | VERBAL | VISUAL | HIGH | 14 | words>letter strings | 23 (18-29) |
| Price, Moore, Humphreys & Wise | 1997 | PET | NO | EXPLICIT | VERBAL | VISUAL | HIGH | 6 | semantic>phonological decision | ? (?-?) |

| | | | | | | | | | | |
|---|---|---|---|---|---|---|---|---|---|---|
| Pulvermueller, Cook & Hauk | 2012 | fMRI | NO | IMPLICIT | VERBAL | VISUAL | LOW | 23 | phrases>hashmarks | 22.8 (?-?) |
| | | | | | | | | | uninflected words>hashmarks | |
| | | | | | | | | | inflected words>hashmarks | |
| Raettig & Kotz | 2008 | fMRI | NO | EXPLICIT | VERBAL | AUDITORY | HIGH | 16 | real words>pseudowords | 26 (21-34) |
| Raposo, Frade & Alves | 2016 | fMRI | NO | IMPLICIT | VERBAL | VISUAL | HIGH | 18 | semantic>perceptual decision | ? (18-22) |
| Raposo, Moss, Stamatakis & Tyler | 2009 | fMRI | NO | IMPLICIT | VERBAL | AUDITORY | LOW | 22 | action sentences>SC noise | 23 (?-?) |
| Rapp & Lipka | 2011 | fMRI | NO | IMPLICIT | VERBAL | VISUAL | LOW/HIGH | 10 | words>checkerboards | ? (18-42) |
| | | | | | | | | | words>letter strings | |
| Redcay, Velnoskey & Rowe | 2016 | fMRI | NO | EXPLICIT | BOTH/NONVERBAL/VERBAL | VISUAL | HIGH | 24 | meaningful>meaningless stimuli | 22 (?-?) |
| | | | | | | | | | communicative>non-communicative gesture | |
| | | | | | | | | | real>pseudoword sentences | |
| Rissman, Eliassen & Blumstein | 2003 | fMRI | NO | IMPLICIT | VERBAL | AUDITORY | HIGH | 15 | words>pseudowords | 22.9 (18-44) |
| Robertson et al. | 2000 | fMRI | NO | IMPLICIT | VERBAL | VISUAL | HIGH | 8 | indefinite article sentence>letter strings | ? (?-?) |
| | | | | | | | | | definite article sentence>letter strings | |
| Rodd, Johnsrude & Davis | 2012 | fMRI | NO | EXPLICIT | VERBAL | AUDITORY | LOW | 15 | speech>SCN | ? (18-40) |
| Rodd, Longe, Randall & Tyler | 2010 | fMRI | NO | EXPLICIT | VERBAL | AUDITORY | LOW | 14 | speech>SCN | ? (19-37) |
| Rogalsky & Hickok | 2009 | fMRI | NO | IMPLICIT | VERBAL | AUDITORY | HIGH | 14 | sentences>word lists | 23 (19-31) |
| Rogalsky, Almeida, Sprouse & Hickok | 2015 | fMRI | NO | IMPLICIT | VERBAL | AUDITORY | HIGH | 15 | words>scrambled | 22 (18-29) |
| Rogers et al. | 2006 | PET | NO | EXPLICIT | BOTH/NONVERBAL | VISUAL | HIGH | 12 | pictures>scrambled pictures | 25 (19-39) |
| | | | | | | | | | specific-level judgement>baseline | |
| Roskies, Fiez, Balota, Raichle & Petersen | 2001 | PET | NO | EXPLICIT | VERBAL | VISUAL | HIGH | 20 | semantic>phonological decision | 25 (18-36) |
| Ross & Olson | 2012 | fMRI | YES | EXPLICIT | NONVERBAL | VISUAL | HIGH | 11 | famous>unknown faces & landmarks | 23 (?-?) |

| | | | | | | | | | | |
|---|---|---|---|---|---|---|---|---|---|---|
| Roxbury, McMahon & Copland | 2014 | fMRI | NO | EXPLICIT | VERBAL | AUDITORY | HIGH | 17 | concrete word>pseudoword | 27 (?-?) |
| | | | | | | | | | abstract word>pseudoword | |
| Ryan, Cox, Hayes & Nadel | 2008 | fMRI | NO | EXPLICIT | VERBAL | VISUAL | HIGH | 10 | semantic fluency (generate) > crosses | 24.5 (19-36) |
| | | | | | | | | | semantic fluency (recall) > crosses | |
| | | | | | | | | | semantic fluency (recall & geneerate)>crosses | |
| Ryan, Lin, Ketcham & Nadel | 2010 | fMRI | NO | EXPLICIT | VERBAL | VISUAL | HIGH | 15 | semantic spatial old>letter judgement | 21.7 (19-27) |
| | | | | | | | | | semantic spatial new>letter judgement | |
| | | | | | | | | | semantic non spatial old>letter judgement | |
| | | | | | | | | | semantic non spatial new>letter judgement | |
| | | | | | | | | | semantic new>episodic judgement | |
| Sabri, Binder, Desai, Medler, Leitl & Liebenthal | 2008 | fMRI | NO | EXPLICIT | VERBAL | AUDITORY | HIGH | 28 | speech>rotated speech | 26.5 (?-?) |
| | | | | | | | | | words>pseudowords | |
| Sachs, Weis, Krings, Huber & Kircher | 2008 | fMRI | NO | EXPLICIT | VERBAL | VISUAL | HIGH | 14 | biased thematic judgement>letters | 28 (?-?) |
| | | | | | | | | | biased taxonomic judgement>letters | |
| | | | | | | | | | balanced taxonomic judgement>letters | |
| | | | | | | | | | balanced taxonomic judgement>letters | |
| Saur et al. | 2008 | fMRI | NO | IMPLICIT | VERBAL | AUDITORY | HIGH | 33 | word>pseudoword sentences | 34 (18-71) |
| Schell, Zaccarella & Friederici | 2017 | fMRI | NO | EXPLICIT | VERBAL | AUDITORY | HIGH | 21 | phrases>non combinatorial words | 27.7 (21-36) |
| Schmitt, Auer & Ferstl | 2019 | fMRI | NO | EXPLICIT | VERBAL | AUDITORY | HIGH | 40 | known>unknown language | 23 (?-?) |
| Schuil, Smits & Zwaan | 2013 | fMRI | NO | EXPLICIT | VERBAL | VISUAL | HIGH | 20 | senteces>pseudowords | 22.1 (18-25) |
| | | | | | | | | | verbs>pseudowords | |

| | | | | | | | | | |
|---|---|---|---|---|---|---|---|---|---|
| | | | | | | | | literal sentences>pseudowords | |
| | | | | | | | | nonliteral sentences>pseudowords | |
| Scott, Blank, Rosen & Wise | 2000 | PET | NO | IMPLICIT | VERBAL | AUDITORY | HIGH | 8 | intelligible>unintelligible speech | ? (?-?) |
| Segal & Petrides | 2012 | fMRI | NO | EXPLICIT | NONVERBAL /VERBAL | VISUAL | HIGH | 90 | writing>copying | 26 (?-?) |
| | | | | | | | | | words>pseudowords | |
| Seghier, Josse, Leff & Price | 2011 | fMRI | NO | EXPLICIT | BOTH | VISUAL | HIGH | 60 | meaningful>meaningless stimuli | 32 (?-?) |
| Sergent, Otha & Macdonald | 1992 | PET | YES | EXPLICIT | VERBAL/NO NVERBAL | VISUAL | HIGH/LOW | 7 | face identity>gender discrimination | ? (22-31) |
| | | | | | | | | | object recognition>gratings | |
| Sheldon, McAndrews, Pruessner & Moscovitch | 2016 | fMRI | NO | EXPLICIT | VERBAL | VISUAL | HIGH | 15 | semantic fluency>perceptual task | 24.8 (?-?) |
| Simard, Monetta, Nagano-Saito & Monchi | 2013 | fMRI | NO | EXPLICIT | VERBAL | VISUAL | HIGH | 14 | semantic>control macthing | 26 (22-31) |
| | | | | | | | | | semantic matching>phonological decision (syllable rhyme) | |
| | | | | | | | | | semantic matching>phonological decision (syllable onset) matching | |
| Slioussar, Kireev, Chernigovskaya, Kataeva, Korotkov & Medvedev | 2014 | fMRI | NO | EXPLICIT | VERBAL | VISUAL | HIGH | 21 | real verbs>pseudowords | 19-32 |
| | | | | | | | | | real nouns>pseudowords | |
| Smith, Myers, Sethi, Pantazatos, Yanagihara & Hirsch | 2012 | fMRI | NO | EXPLICIT | VERBAL | VISUAL | HIGH | 14 | semantics>baseline | 28.7 (?-?) |
| Snijders, Vosse, Kempen, Van Berkum, Petersson & Hagoort | 2009 | fMRI | NO | IMPLICIT | VERBAL | VISUAL | HIGH | 28 | sentences>word lists | ? (18-35) |
| Stowe, Paans, Wijers, Zwarts, Mulder & Vaalburg | 1999 | PET | NO | IMPLICIT | VERBAL | VISUAL | HIGH/REST | 12 | sentences>scrambled word lists | 31 (19-47) |
| | | | | | | | | | sentences>rest | |

| | | | | | | | | | | |
|---|---|---|---|---|---|---|---|---|---|---|
| Straube, Green, Weis & Kircher | 2012 | fMRI | NO | IMPLICIT | VERBAL/NONVERBAL | AUDITORY/VISUAL | HIGH | 16 | known>unknown language | 28.8 (23-55) |
| | | | | | | | | | iconic>meaningless gesture | |
| Stringaris, Medford, Giampietro. Brammer & David | 2007 | fMRI | NO | EXPLICIT | VERBAL | VISUAL | HIGH | 11 | literal>meaningless sentences | 33.3 (?-?) |
| | | | | | | | | | methaphors>meaningless sentences | |
| Sugiura et al. | 2006 | fMRI | YES | EXPLICIT | VERBAL | VISUAL | HIGH | 24 | famous>unfamiliar names | ? (18-25) |
| | | | | | | | | | personal>unfamiliar names | |
| Sugiura et al. | 2001 | PET | YES | EXPLICIT | NONVERBAL | VISUAL | HIGH | 5 | identity discrimination>control | ? (23-28) |
| | | | | | | | | | identity discrimination>face direction | |
| Sugiura et al. | 2008 | fMRI | YES | EXPLICIT | VERBAL | VISUAL | HIGH | 25 | low familiar>unfamiliar names | ? (18-32) |
| | | | | | | | | | personal familiar>unfamiliar names | |
| | | | | | | | | | high familiar>unfamiliar names | |
| | | | | | | | | | high familiar>unfamiliar names | |
| Sun, Xue, Zhang, Zuo, Chen, Wang, Martin, Wang, Chen, He & Wang | 2017 | fMRI | NO | EXPLICIT | VERBAL | VISUAL | HIGH | 11 | semantic>orthography judgement | ? (?-?) |
| Szlachta, Bozic, Jelowicka & Marslen-Wilson | 2012 | fMRI | NO | IMPLICIT | VERBAL | VISUAL | LOW | 21 | words>musical rain | 25.2 (18-33) |
| | | | | | | | | | nouns>musical rain | |
| | | | | | | | | | inflected nouns>musical rain | |
| Takeichi, Koyama, Terao, Takeuchi, Toyosawa & Murohashi | 2010 | fMRI | NO | IMPLICIT | VERBAL | AUDITORY | HIGH | 23 | speech>reversed speech | 25 (20-38) |
| | | | | | | | | | speech>modulated speech | |
| Taminato, Miura, Sugiura & Kawashima | 2014 | fMRI | NO | EXPLICIT | NONVERBAL | VISUAL | LOW | 35 | object recognition>control task | ? (19-31) |
| | | | | | | | | | object recognition>control task | |

| | | | | | | | | | | |
|---|---|---|---|---|---|---|---|---|---|---|
| Taylor, Arsalidou, Bayless, Morris, Evans & Barbeau | 2009 | fMRI | YES | IMPLICIT | NONVERBAL | VISUAL | HIGH/REST | 10 | own>unfamiliar face | 35.4 (?-?) |
| | | | | | | | | | partner's>unfamiliar face | |
| | | | | | | | | | parent's>unfamiliar face | |
| | | | | | | | | | own face>baseline (rest) | |
| | | | | | | | | | parents face>baseline (rest) | |
| | | | | | | | | | famous face>baseline (rest) | |
| Thierry & Price | 2006 | PET | NO | EXPLICIT | VERBAL/NONVERBAL | AUDITORY | HIGH | 12 | auditory words>speech control(scrambled) | 26.3 (?-?) |
| | | | | | | | | | auditory sounds>sound control (scrambled) | |
| Thierry & Price | 2006 | PET | NO | EXPLICIT | VERBAL/NONVERBAL | VISUAL | HIGH | 12 | visual words>text control (scrambled letter strings) | 26.3 (?-?) |
| | | | | | | | | | visual videos>video control (distorted) | |
| Tieleman, Seurinck, Deblaere, Vandemaele, Vingerhoets & Achten | 2005 | fMRI | NO | EXPLICIT | VERBAL | VISUAL | HIGH | 22 | self-paced semantic>perceptual decision | 29 (22-47) |
| | | | | | | | | | fixed-paced semantic>perceptual decision | |
| Tyler, Stamatakis, Dick, Bright, Fletcher & Moss | 2003 | fMRI | NO | EXPLICIT | VERBAL | VISUAL | HIGH | 12 | animals>baseline | 24 (?-?) |
| | | | | | | | | | tool action words>baseline | |
| | | | | | | | | | biological action>baseline | |
| Vagharchakian, Dehaene-Lambertz, Pallier & Dehaene | 2012 | fMRI | NO | EXPLICIT | VERBAL | BOTH/AUDITORY/VISUAL | HIGH | 16 | intelligible>unintelligble compression rate | |
| | | | | | | | | | intelligible>unintelligble compression rate | |
| | | | | | | | | | intelligible>unintelligble compression rate | |
| Van Ettinger-Veenstra, McAllister, Lundberg, Karlsson & Engstrom | 2016 | fMRI | NO | EXPLICIT | VERBAL | VISUAL | HIGH | 27 | sentences>symbol strings | 25.5 (18-35) |
| van Leeuwen et al. | 2014 | fMRI | NO | EXPLICIT | BOTH | BOTH | HIGH | 16 | speech>reversed speech | ? (19-35) |

| | | | | | | | | | | |
|---|---|---|---|---|---|---|---|---|---|---|
| Vignali, Hawelka, Hutzler & Richlan | 2019 | fMRI | NO | EXPLICIT | VERBAL | VISUAL | HIGH | 21 | foveal & parafovealwords>foveal & parafoveal pseudowords | 25.8 (?-?) |
| Vingerhoets | 2008 | fMRI | NO | IMPLICIT | NONVERBAL | VISUAL | HIGH | 14 | familiar>unfamiliar tools | 22 (20-24) |
| Visser, Jefferies, Embleton & Lambon Ralph | 2012 | fMRI | NO | EXPLICIT | BOTH/NONVERBAL/VERBAL | VISUAL | HIGH | 15 | semantics>baseline | ? (?-?) |
| | | | | | | | | | pictures>baseline | |
| | | | | | | | | | words>baseline | |
| Vitello, Warren, Devlin & Rodd | 2014 | fMRI | NO | EXPLICIT | VERBAL | AUDITORY | LOW | 20 | sentences>SCN | 23.8 (18-35) |
| von Kriegstein, Eger, Kleinschmidt & Giraud | 2003 | fMRI | NO | EXPLICIT | VERBAL | AUDITORY | HIGH | 14 | sentence>speech envelope | ? (20-51) |
| Wang, Zhao, Zevin & Yang | 2016 | fMRI | NO | EXPLICIT | VERBAL | VISUAL | HIGH | 16 | words>nonsense strokes | ? (18-25) |
| Weiss, Katzir & Bitan | 2015 | fMRI | NO | IMPLICIT | VERBAL | VISUAL | LOW | 18 | pointed words>asterisks | 27.1 (22-32) |
| | | | | | | | | | unpointed words>asterisks | |
| Welcome & Joanisse | 2012 | fMRI | NO | EXPLICIT | VERBAL | VISUAL | HIGH | 20 | semantic>phonological/orthographic decision | 29.7 (19-59) |
| Wende, Straube, Stratmann, Sommer, Kircher & Nagels | 2012 | fMRI | NO | EXPLICIT | VERBAL | VISUAL | HIGH/REST | 18 | semantic>phonological fluency | 25.8 (20-45) |
| | | | | | | | | | causal fluency>rest | |
| | | | | | | | | | free association>rest | |
| Wirth, Jann, Dierks, Federspiel, Wiest & Horn | 2011 | fMRI | NO | EXPLICIT | VERBAL | VISUAL | HIGH | 19 | semantic>phonological & perceptual decision | 27 (?-?) |
| Wright et al. | 2008 | fMRI | NO | EXPLICIT | BOTH/VERBAL/NONVERBAL | VISUAL | HIGH | 10 | semantic>perceptual matching | 25 (19-35) |
| | | | | | | | | | semantic>perceptual matching | |
| | | | | | | | | | semantic>perceptual matching | |
| Wright et al. | 2008 | fMRI | NO | EXPLICIT | VERBAL/NONVERBAL | VISUAL | REST | 15 | words>rest | 28 (20-45) |
| | | | | | | | | | pictures>rest | |

| | | | | | | | | | | |
|---|---|---|---|---|---|---|---|---|---|---|
| Wright, Randall, Marslen-Wilson & Tyler | 2011 | fMRI | NO | BOTH | VERBAL | AUDITORY | LOW | 14 | speech>musical rain | 23.9 (19-34) |
| Wu, Mai, Tang, Ge, Luo & Liu | 2013 | fMRI | NO | IMPLICIT | VERBAL | VISUAL | LOW | 19 | arm words>checkerboard | 22.32 (19-25) |
| | | | | | | | | | leg words>checkerboard | |
| | | | | | | | | | mouth words>checkerboard | |
| Xiao et al. | 2005 | fMRI | NO | EXPLICIT | VERBAL | AUDITORY | HIGH/REST | 14 | words>pseudowords | 20.9 (18-23) |
| | | | | | | | | | words>rest | |
| Yang, Li, Fang, Shu, Liu & Chen | 2016 | fMRI | NO | IMPLICIT | VERBAL | VISUAL | LOW | 20 | opaque idioms>hashmarks | 21.7 (20-25) |
| | | | | | | | | | transparent idioms>hashmarks | |
| | | | | | | | | | literal phrases>hashmarks | |
| Zaccarella & Friederici | 2015 | fMRI | NO | EXPLICIT | VERBAL | VISUAL | HIGH | 22 | words>pseudowords | 28.5 (?-?) |
| Zhang, Liu & Zhang | 2014 | fMRI | NO | EXPLICIT | VERBAL | VISUAL | LOW | 18 | nonliving words>asterisks | 20.3 (17-23) |
| | | | | | | | | | living words>asterisks | |
| Zhang, Xiao & Weng | 2012 | fMRI | NO | EXPLICIT | VERBAL | VISUAL | HIGH/REST | 14 | words>pseudowords | 21.7 (18-25) |
| | | | | | | | | | words>rest | |
| Zhuang & Devereux | 2017 | fMRI | NO | EXPLICIT | VERBAL | AUDITORY | HIGH | 16 | phrases>words | ? (18-34) |
| Zou, Packard, Xia, Liu & Shu | 2016 | fMRI | NO | IMPLICIT | VERBAL | AUDITORY | LOW | 17 | speech>tone | 21.24 (?-?) |
| | | | | | | | | | speech>tone | |
| | | | | | | | | | speech>tone | |
| | | | | | | | | | identical speech>tone | |
| | | | | | | | | | words>tone | |
| Zvyagintsev, Clemens, Chechko, Mathiak, Sack & Mathiak | 2013 | fMRI | NO | IMPLICIT | NONE | NONE | HIGH | 15 | visual imagery>counting | 25.1 (?-?) |
| | | | | | | | | | auditory imagery>counting | |

**The references to access the listed studies are listed at the end of the document*

***Table M2 SC*** *List of studies included in the* ***semantic cognition MAIN high and low baselines (rest excluded)*** *(N = 211) meta-analysis.* ***Note:*** *This data set was used for the* ***MAIN*** *ToM & SC conjunction and contrast analyses. The list of excluded studies using rest as a baseline can be seen at the end of the document highlighted in grey. ToM= Theory of Mind, SC= Semantic Cognition, N= Sample Size*

| *Authors | Year | Imaging Method | Social Content | Instructional Cue | Stimulus Domain | Sensory Input Modality | Baseline Type | N | Contrast |
|---|---|---|---|---|---|---|---|---|---|
| AbdulSabur et al. | 2014 | fMRI | NO | EXPLICIT | VERBAL | VISUAL | LOW | 18 | narrative production>recitation |
| Abraham et al. | 2012 | fMRI | NO | EXPLICIT | VERBAL | VISUAL | HIGH | 19 | high&low divergent thinking (semantic)>1&2 back letter identity (working memory) |
| Alain, He & Grady | 2008 | fMRI | NO | EXPLICIT | VERBAL | AUDITORY | HIGH | 16 | sound category>sound location |
| Assadollahi, Meinzer, Flaisch, Obleser & Rockstroh | 2009 | fMRI | NO | IMPLICIT | VERBAL | VISUAL | HIGH | 20 | nouns followed by 1&3 argument verbs>letter strings followed by 1&3argument verbs |
| Axmacher, Bialleck, Weber, Helmstaedter, Elger & Fell | 2009 | fMRI | NO | EXPLICIT | VERBAL | VISUAL | HIGH | 32 | word decision>spatial decision |
| Bagga et al. | 2013 | fMRI | NO | EXPLICIT | VERBAL | VISUAL | HIGH | 18 | semantic>case matching judgement |
| Barros-Loscertales et al. | 2012 | fMRI | NO | EXPLICIT | VERBAL | VISUAL | LOW | 59 | control words>hashmarks baseline |
| Baumgaertener, Weiller & Buchel | 2002 | fMRI | NO | EXPLICIT | VERBAL | VISUAL | HIGH | 9 | word>pseudoword in sentence |
| Baumgaertner et al. | 2007 | fMRI | NO | IMPLICIT | VERBAL | AUDITORY | HIGH | 19 | sentences>reversed sentences |
| | | | | | | | | | videos>scrambled videos |
| Bautista & Wilson | 2016 | fMRI | NO | IMPLICIT | VERBAL | AUDITORY | HIGH | 12 | clear>scrambled rotated speech |
| Bick, Goelman & Frost | 2008 | fMRI | NO | EXPLICIT | VERBAL | VISUAL | LOW | 14 | semantic>visual control |
| | | | | | | | | | morphologiacl>visual control |
| | | | | | | | | | orthographic>visual control |
| | | | | | | | | | phonological>visual control |
| Binder, Frost, Hammeke, Bellgowan, Rao & Cox | 1999 | fMRI | NO | EXPLICIT | VERBAL | AUDITORY | HIGH | 30 | semantic>phonological decision |
| Binder et al. | 2003 | fMRI | NO | EXPLICIT | VERBAL | VISUAL | HIGH | 24 | word>nonword |

| | | | | | | | | | |
|---|---|---|---|---|---|---|---|---|---|
| Birn et al. | 2010 | fMRI | NO | EXPLICIT | VERBAL | VISUAL | LOW | 14 | category & letter fluency>months ( automatic speech) |
| | | | | | | | | | category>letter fluency |
| Bonhage, Fiebach, Bahlmann & Mueller | 2014 | fMRI | NO | EXPLICIT | VERBAL | VISUAL | HIGH | 18 | sentence fragments>ungrammatical word strings |
| Bonhage, Mueller, Friederici & Fiebach | 2015 | fMRI | NO | EXPLICIT | VERBAL | VISUAL | HIGH | 18 | sentences>jabberwocky sentences |
| Booth et al. | 2006 | fMRI | NO | EXPLICIT | VERBAL | VISUAL | HIGH | 13 | meaning (semantic) judgement>rhyming( phonological) judgement |
| | | | | | | | | | meaning(semantic)>control(symbols) |
| Boulenger, Hauk & Pulvermuller | 2009 | fMRI | NO | EXPLICIT | VERBAL | VISUAL | LOW | 18 | sentences> hashmarks |
| | | | | | | | | | sentences> hashmarks |
| Bozic & Marslen-Wilson | 2013 | fMRI | NO | IMPLICIT | VERBAL | AUDITORY | HIGH | 13 | speech>musical rain |
| Brambati, Benoit, Monetta, Belleville & Joubert | 2010 | fMRI | YES | EXPLICIT | BOTH | BOTH | HIGH | 12 | general & specific occupation judgement>baseline (scrambled face) |
| Bruffaerts, Dupont, Peeters, De Deyne, Storms & Vandenberghe | 2013 | fMRI | NO | EXPLICIT | BOTH | VISUAL | HIGH | 19 | real > scrambled pictures & words |
| Bulut, Hung, Tzeng & Wu | 2017 | fMRI | NO | IMPLICIT | VERBAL | VISUAL | HIGH | 20 | sentences>unstructured word lists |
| | | | | | | | | | sentences>unstructured character list |
| Cai, Kochiyama, Osaka & Wu | 2007 | fMRI | NO | IMPLICIT | VERBAL | AUDITORY | HIGH | 15 | words>nonsense words |
| Cao, Peng, Liu, Jin, Fan, Deng, & Booth | 2009 | fMRI | NO | EXPLICIT | VERBAL | VISUAL | LOW | 13 | meaning>perceptual decision |
| Cappa, Perani, Schnur, Tettamanti & Fazio | 1998 | PET | NO | EXPLICIT | VERBAL | VISUAL | HIGH | 13 | words>pseudowords |
| | | | | | | | | | animal visual knowledge>pseudowords |
| | | | | | | | | | tool visual knowledge>pseudowords |

| | | | | | | | | | |
|---|---|---|---|---|---|---|---|---|---|
| | | | | | | | | | animal associative knowledge>pseudowords |
| | | | | | | | | | tools functional knowledge>pseudowords |
| Carota, Kriegeskorte, Nili & Pulvermuller | 2017 | fMRI | NO | IMPLICIT | VERBAL | VISUAL | LOW | 23 | words>hashmarks |
| Carota, Moseley & Pulvermueller | 2012 | fMRI | NO | IMPLICIT | VERBAL | VISUAL | LOW | 18 | all words>hashmarks |
| | | | | | | | | | tool words>hashmarks |
| | | | | | | | | | animal words>hashmarks |
| | | | | | | | | | food words>hashmarks |
| Chan, Tang, Tang, Lee, Lo & Kwong | 2009 | fMRI | NO | IMPLICIT | VERBAL | VISUAL | HIGH | 22 | synonyms>pseudocharacters |
| | | | | | | | | | synonyms>Korean characters (unknown) |
| Chiao, Harada, Oby, Li, Parrish & Bridge | 2009 | fMRI | YES | EXPLICIT | NONVERBAL | VISUAL | LOW | 12 | uniform status judgement>colour change detection |
| | | | | | | | | | face status judgement>colour change detection |
| | | | | | | | | | car status judgement>colour change detection |
| Chou, Chen, Wu & Booth | 2009 | fMRI | NO | EXPLICIT | VERBAL | VISUAL | HIGH | 31 | related words>false font |
| | | | | | | | | | unrelated words>false font |
| Chou, Chen, Wu & Booth | 2009 | fMRI | NO | EXPLICIT | VERBAL | VISUAL | HIGH | 32 | related words>false font |
| | | | | | | | | | unrelated words>false font |
| Chouinard, Morrissey, Kohler & Goodale | 2008 | fMRI | NO | EXPLICIT | NONVERBAL | VISUAL | HIGH | 14 | objects>scrambled objects |
| Chow, Kaup, Raabe & Greenlee | 2008 | fMRI | NO | EXPLICIT | VERBAL | VISUAL | HIGH | 15 | predictive&normal reading>pseudoword reading |
| | | | | | | | | | normal reading>pseudoword reading |
| | | | | | | | | | predictive reading>pseudoword reading |
| Christensen, Antonucci, Lockwood, Kittleson & Plante | 2008 | fMRI | NO | EXPLICIT | VERBAL | AUDITORY | HIGH | 14 | diotic listening>reversed speech |

| | | | | | | | | | |
|---|---|---|---|---|---|---|---|---|---|
| | | | | | | | | | dichotic listening>reversed speech |
| Clos, Langner, Meyer, Oechslin, Zilles & Eickhoff | 2014 | fMRI | NO | EXPLICIT | VERBAL | AUDITORY | HIGH | 29 | intelligibilty based on cue > unintelligible |
| Damasio, Grabowski, Tranel, Ponto, Hichwa & Damasio | 2001 | PET | NO | EXPLICIT | NONVERBAL | VISUAL | LOW | 10 | actions without implement>control task |
| | | | | | | | | | actions with implement>control task |
| Damasio, Tranel, Grabowski, Adolphs & Damasio | 2004 | PET | YES | EXPLICIT | NONVERBAL | VISUAL | HIGH | 55 | persons>face orientation judgement |
| | | | | | | | | | animals>scrambled pictures |
| | | | | | | | | | tools>scrambled pictures |
| Davis, Meunier & Marslen-Wilson | 2004 | fMRI | NO | EXPLICIT | VERBAL | VISUAL | HIGH | 11 | words>letter strings |
| Davis, Ford, Kherif & Johnsrude | 2011 | fMRI | NO | EXPLICIT | VERBAL | AUDITORY | LOW | 12 | clear & anomalous sentences>signal correlated noise |
| Demonet et al. | 1992 | PET | NO | EXPLICIT | VERBAL | AUDITORY | HIGH | 9 | words>phonemes |
| | | | | | | | | | words>tones |
| Devlin, Matthews & Rushworth | 2003 | fMRI | NO | EXPLICIT | VERBAL | VISUAL | HIGH | 12 | semantic>phonological judgement |
| Devlin et al. | 2002 | PET | NO | EXPLICIT | VERBAL | VISUAL | HIGH | 12 | all semantic>letter detection |
| | | | | | | | | | all semantic>letter detection |
| Devlin et al. | 2002 | PET | NO | EXPLICIT | VERBAL | VISUAL | HIGH | 8 | all semantic>letter categorisation |
| Devlin et al. | 2002 | fMRI | NO | EXPLICIT | VERBAL | VISUAL | HIGH | 8 | all semantic> letter categorisation |
| Devlin et al. | 2000 | PET | NO | EXPLICIT | VERBAL | VISUAL | HIGH | 8 | semantic categorisation>letter categorisation |
| Devlin et al. | 2000 | fMRI | NO | EXPLICIT | VERBAL | VISUAL | HIGH | 8 | semantic categorisation>letter categorisation |
| Diaz & McCarthy | 2009 | fMRI | NO | IMPLICIT | VERBAL | VISUAL | HIGH | 16 | all words>nonwords |
| Dreyer & Pulvermueller | 2018 | fMRI | YES | IMPLICIT | VERBAL | VISUAL | LOW | 28 | all nouns>hashmarks |
| | | | | | | | | | abstract emotional nouns>baseline (hashmarks) |

| | | | | | | | | | |
|---|---|---|---|---|---|---|---|---|---|
| | | | | | | | | | abstract mental nouns>baseline (hashmarks) |
| | | | | | | | | | food nouns>baseline (hashmarks) |
| | | | | | | | | | tool nouns>baseline (hashmarks) |
| Ebisch et al. | 2007 | fMRI | NO | EXPLICIT | BOTH | VISUAL | HIGH | 17 | functional&visuospatial pictures&words>meaningless drawings&pseudowords |
| Elfgren, Westen, Passant, Larsson, Mannfolk & Fransson | 2006 | fMRI | YES | EXPLICIT | NONVERBAL | VISUAL | HIGH | 15 | familiar>unfamiliar faces (identification) |
| Emmorey, Weisberg, McCullough & Petrich | 2013 | fMRI | NO | EXPLICIT | VERBAL | VISUAL | HIGH | 14 | words(semantic judgement)>false fonts |
| | | | | | | | | | words(phonological)>false fonts |
| Emmorey, Xu, Gannon, Goldin-Meadow & Braun | 2010 | fMRI | NO | IMPLICIT | NONVERBAL | VISUAL | HIGH | 14 | meaningful pantomimes>unknown sign language |
| Engelien et al. | 2006 | PET | NO | IMPLICIT | NONVERBAL | AUDITORY | HIGH | 6 | meaningful>meaningless sounds |
| Erb, Henry, Eisner & Obleser | 2013 | fMRI | NO | EXPLICIT | VERBAL | AUDITORY | HIGH | 30 | speech>vocoded speech |
| Europa, Gitelman, Kiran & Thompson | 2019 | fMRI | NO | EXPLICIT | BOTH | BOTH | HIGH | 21 | sentences>baseline (reversed sentences) |
| Foki, Gartus, Geissler & Beisteiner | 2008 | fMRI | | EXPLICIT | VERBAL | VISUAL | LOW | 23 | semantic judgement>tongue movements |
| Friederici, Kotz, Scott & Obleser | 2010 | fMRI | NO | IMPLICIT | VERBAL | AUDITORY | HIGH | 17 | intelligible speech>rotated speech |
| Friese, Rutschmann, Raabe & Schmalhofer | 2008 | fMRI | NO | EXPLICIT | VERBAL | VISUAL | HIGH | 13 | words>pseudowords |
| Garbin, Collina & Tabossi | 2012 | fMRI | NO | EXPLICIT | VERBAL | VISUAL | HIGH | 12 | object noun>pseudoword |
| | | | | | | | | | event noun>pseudoword |
| | | | | | | | | | verb>pseudoword |
| Garn, Allen & Larsen | 2009 | fMRI | NO | EXPLICIT | NONVERBAL | VISUAL | HIGH | 26 | pictures>scrambled pictures |
| | | | | | | | | | plants>scrambled pictures |
| | | | | | | | | | tools>scrambled pictures |

| | | | | | | | | | |
|---|---|---|---|---|---|---|---|---|---|
| Geranmayeh, Brownsett, Leech, Beckmann, Woodhead & Wise | 2012 | fMRI | NO | EXPLICIT | VERBAL | VISUAL | HIGH | 19 | speech>tongue movements |
| Gerlach, Law, Gade & Paulson | 1999 | PET | NO | EXPLICIT | NONVERBAL | VISUAL | HIGH | 15 | object decision>pattern discrimination |
| Gesierich et al. | 2012 | fMRI | YES | EXPLICIT | NONVERBAL | VISUAL | HIGH | 21 | familiar>scrambled faces |
| | | | | | | | | | familiar > unfamiliar faces |
| | | | | | | | | | familiar>unfamiliar faces |
| Giraud & Price | 2001 | PET | NO | BOTH | BOTH | AUDITORY | BOTH | 12 | words+environmental sounds>syllables+noise |
| Giraud et al. | 2004 | fMRI | NO | EXPLICIT | VERBAL | AUDITORY | HIGH | 8 | natural speech>speech envelope |
| Gitelman, Nobre, Sonty, Parrish & Mesulam | 2005 | fMRI | NO | EXPLICIT | VERBAL | VISUAL | HIGH | 14 | semantic>control task |
| Gorno-Tempini et al. | 1998 | PET | YES | EXPLICIT | NONVERBAL | VISUAL | HIGH | 6 | famous faces>controls |
| | | | | | | | | | famous names>controls |
| | | | | | | | | | double famous proper names>controls |
| | | | | | | | | | non-famous faces>controls |
| | | | | | | | | | Non-famous names>controls |
| | | | | | | | | | Double common names>controls |
| Grabowski, Damasio, Tranel, Boles Ponto, Hichwa & Damasio | 2001 | PET | YES | EXPLICIT | NONVERBAL | VISUAL | HIGH | 10 | naming persons>building orientation judgement |
| | | | | | | | | | naming landmarks>face orientation judgement |
| | | | | | | | | | naming persons>face orientation judgement |
| | | | | | | | | | naming unique entities>baseline |
| Graves, Binder, Desai, Conant & Seidenberg | 2010 | fMRI | NO | EXPLICIT | VERBAL | VISUAL | HIGH | 23 | forward>reverse phrases |
| Graves, Binder, Desai, Conant & Seidenberg | 2010 | fMRI | NO | EXPLICIT | VERBAL | VISUAL | HIGH | 22 | forward>reverse phrases |
| Grindrod, Garnett, Malyutina & den Ouden | 2014 | fMRI | NO | EXPLICIT | VERBAL | VISUAL | HIGH | 23 | words>nonwords |
| Grossman et al. | 2002a | fMRI | NO | IMPLICIT | VERBAL | VISUAL | HIGH | 16 | all nouns > pseudowords |

| | | | | | | | | | |
|---|---|---|---|---|---|---|---|---|---|
| | | | | | | | | | implements>pseudowords |
| | | | | | | | | | animals>pseudowords |
| | | | | | | | | | abstract>pseudowords |
| Grossman et al., 2002b | 2002b | fMRI | NO | IMPLICIT | VERBAL | VISUAL | HIGH | 16 | verbs>pseudowords |
| Groussard et al. | 2010 | PET | NO | EXPLICIT | VERBAL | VISUAL | HIGH | 11 | verbal semantics>verbal reference |
| | | | | | | | | | musical semantic>musical reference |
| Guediche, Reilly, Santiago, Laurent & Blumstein | 2016 | fMRI | NO | IMPLICIT | VERBAL | AUDITORY | LOW | 16 | related > repeated sentence |
| | | | | | | | | | unrelated > repeated sentence |
| Gurd et al. | 2002 | fMRI | NO | EXPLICIT | VERBAL | AUDITORY | HIGH | 11 | category>rote fluency |
| Haberling, Corballis &Corballis | 2016 | fMRI | NO | EXPLICIT | NONVERBAL | VISUAL | HIGH | 92 | meaningful pantomimes>unknown sign language |
| | | | | | | | | | meaningful pantomimes>dog videos |
| | | | | | | | | | synonyms>letter strings |
| Hagoort et al. | 1999 | PET | NO | IMPLICIT | VERBAL | VISUAL | HIGH | 10 | words>pseudowords |
| Harrington, Farias & Davis | 2009 | fMRI | NO | EXPLICIT | NONVERBAL | VISUAL | HIGH | 8 | familiar>non objects |
| Hartung, Hagoort & Willems | 2017 | fMRI | NO | IMPLICIT | VERBAL | AUDITORY | HIGH | 52 | first-person speech>unintelligible reversed speech |
| | | | | | | | | | third-person speech>unintelligible reversed speech |
| Hauk & Pulvermueller | 2011 | fMRI | NO | IMPLICIT | VERBAL | VISUAL | LOW | 21 | action words>hashmarks |
| | | | | | | | | | uni manual action words>hashmarks |
| | | | | | | | | | uni manual action words>hashmarks |
| Hayashi et al. | 2014 | fMRI | NO | EXPLICIT | VERBAL | VISUAL | LOW | 16 | concrete word>astersisks |
| | | | | | | | | | abstract word>asterisks |
| Heim, Eickhoff & Amunts | 2008 | fMRI | NO | EXPLICIT | VERBAL | VISUAL | HIGH | 28 | semantic > phonological fluency |
| Henke et al. | 1999 | PET | NO | EXPLICIT | VERBAL | VISUAL | HIGH | 12 | associative word learning>single word encoding |

| | | | | | | | | | |
|---|---|---|---|---|---|---|---|---|---|
| Herbster et al. | 1997 | PET | NO | EXPLICIT | VERBAL | VISUAL | HIGH | 10 | Irregular>zero order speak |
| | | | | | | | | | regular>zero order speak |
| | | | | | | | | | irregular+regular>zero order speak |
| Hervais-Adelman, Carlyon, Johnsrude & Davis | 2012 | fMRI | NO | EXPLICIT | VERBAL | AUDITORY | HIGH | 15 | clear>vocoded speech (uninteligible) |
| Higuchi, Moriguchi, Murakami, Katsunuma, Mishima &Uno | 2015 | fMRI | NO | IMPLICIT | VERBAL | VISUAL | LOW | 28 | all characters>checkerboard |
| Hocking, McMahon & de Zubicaray | 2011 | fMRI | NO | EXPLICIT | NONVERBAL | AUDITORY | HIGH | 13 | all environmental sounds>perceptual baseline |
| Holle, Gunter, Rueschemeyer, Hennenlotter & Iacoboni | 2008 | fMRI | NO | EXPLICIT | BOTH | BOTH | HIGH | 17 | iconic gesture of dominant meaning>grooming |
| | | | | | | | | | iconic gesture of subordinate meaning>grooming |
| Homae, Yahata & Sakai | 2003 | fMRI | NO | EXPLICIT | VERBAL | AUDITORY | HIGH | 10 | auditory sentences>auditory non words |
| | | | | | | | | | visual sentences>visual non words words |
| Husain, Patkin, Kim, Braun & Horwitz | 2012 | fMRI | NO | EXPLICIT | NONVERBAL | VISUAL | HIGH | 16 | meaningful iconic>meaningless gestures |
| Hwang, Palmer, Basho, Zadra & Muller | 2009 | fMRI | NO | EXPLICIT | VERBAL | AUDITORY | LOW | 13 | fluency generation>production baseline |
| Ikuta et al. | 2006 | fMRI | NO | IMPLICIT | VERBAL | VISUAL | LOW | 34 | sentences>word lists |
| Jackson, Hoffman, Pobric, Lambon Ralph | 2015 | fMRI | NO | EXPLICIT | VERBAL | VISUAL | HIGH | 24 | words>letter strings |
| Jensen, Hargreaves, Bass, Pexman, Goodyear & Federico | 2011 | fMRI | NO | EXPLICIT | VERBAL | VISUAL | HIGH | 12 | words>pseudowords |
| Jeon, Lee, Kim & Cho | 2009 | fMRI | NO | EXPLICIT | VERBAL | VISUAL | HIGH | 16 | synonys>nonwords |
| | | | | | | | | | antonyms>nonwords |
| | | | | | | | | | English synonyma>nonwords |
| | | | | | | | | | Korean synonyms>nonwords |
| | | | | | | | | | honorific words>nonwords |
| Joubert et al. | 2004 | fMRI | NO | IMPLICIT | VERBAL | VISUAL | HIGH | 10 | low frequency words>nonwords |

| | | | | | | | | | |
|---|---|---|---|---|---|---|---|---|---|
| Kang et al. | 2006 | PET | NO | EXPLICIT | VERBAL | BOTH | HIGH | 17 | high frequency words>consonant strings |
| | | | | | | | | | low frequency words>consonant strings |
| | | | | | | | | | audio-visual speech>noises and facial movements |
| | | | | | | | | | auditory speech>white noise |
| | | | | | | | | | visual speech>facial movements (chewing gum) |
| Khader, Jost, Mertens, Bien & Roesler | 2010 | fMRI | NO | EXPLICIT | VERBAL | VISUAL | HIGH | 16 | noun>rhyme generation |
| | | | | | | | | | verb>rhyme generation |
| | | | | | | | | | noun generation>letter detection |
| | | | | | | | | | verb generation>letter detection |
| Kim et al. | 2009 | fMRI | NO | EXPLICIT | VERBAL | VISUAL | HIGH | 36 | sentences>word lists |
| Kinno, Kawamura, Shioda & Sakai | 2008 | fMRI | NO | EXPLICIT | BOTH | VISUAL | HIGH | 14 | canonical sentence>picture & letter strings |
| | | | | | | | | | active sentence>picture & letter strings |
| | | | | | | | | | passive sentence>picture & letter strings |
| Kotz, Cappa, Von Cramon & Friederici | 2002 | fMRI | NO | EXPLICIT | VERBAL | AUDITORY | HIGH | 13 | words>pseudowords |
| Kuchinke et al. | 2005 | fMRI | YES | IMPLICIT | VERBAL | VISUAL | HIGH | 20 | emotion word>nonword |
| Kumar | 2016 | fMRI | NO | EXPLICIT | VERBAL | VISUAL | HIGH | 20 | abstract+concrete words>pseudowords |
| | | | | | | | | | abstract words>pseudowords |
| Kuperberg et al. | 2000 | fMRI | NO | EXPLICIT | VERBAL | AUDITORY | HIGH | 4 | sentences>words strings |
| Kyong, Scott, Rosen, Howe, Agnew & McGettigan | 2014 | fMRI | NO | IMPLICIT | VERBAL | AUDITORY | LOW | 19 | intelligible vocoded>inverted vocoded speech |
| Leff, Schofield, Stephan, Crinion, Friston & Price | 2008 | fMRI | NO | IMPLICIT | VERBAL | AUDITORY | HIGH | 26 | speech>reversed speech |
| Leung & Alain | 2011 | fMRI | NO | EXPLICIT | NONVERBAL | AUDITORY | HIGH | 16 | semantic>location matching |
| Leveroni et al. | 2000 | fMRI | YES | EXPLICIT | NONVERBAL | VISUAL | HIGH | 11 | familiar faces>foils(never seen faces) |
| | | | | | | | | | newly learned faces>foils(never seen faces) |

| | | | | | | | | | |
|---|---|---|---|---|---|---|---|---|---|
| Lin, Wang, Zhao, Liu, Li & Bi | 2015 | fMRI | NO | EXPLICIT | VERBAL | VISUAL | HIGH | 20 | words>pseudowords |
| Liu et al. | 2009 | fMRI | NO | EXPLICIT | VERBAL | VISUAL | LOW | 16 | words meaning>slashes |
| | | | | | | | | | words rhyming>slashes |
| | | | | | | | | | words meaning>tones |
| | | | | | | | | | words rhyming>tones |
| Liuzzi et al. | 2017 | fMRI | NO | EXPLICIT | VERBAL | BOTH | HIGH | 18 | semantic judgement > input modality detection |
| Ludersdorfer, Wimmer, Richlan, Schurz, Hutzler & Kronbichler | 2016 | fMRI | NO | EXPLICIT | VERBAL | AUDITORY | LOW | 29 | words orthographic>tones |
| | | | | | | | | | words semantic>tones |
| Ludersdorfer, Schurz, Richlan, Kronbichler & Wimmer | 2013 | fMRI | NO | EXPLICIT | VERBAL | VISUAL | HIGH | 29 | words>false fonts |
| | | | | | | | | | words>pseudowords |
| | | | | | | | | | speech>reversed speech |
| | | | | | | | | | words>pseudowords |
| Malins, Gumkowski, Buis, Molfese, Rueckl, Frost, Pugh, Morris & Mencl | 2016 | fMRI | NO | IMPLICIT | VERBAL | VISUAL | LOW | 18 | unrelated words>false font |
| | | | | | | | | | unrelated>pseudowords |
| Marques, Canessa & Cappa | 2009 | fMRI | NO | EXPLICIT | VERBAL | VISUAL | LOW | 21 | sentences>crosses |
| Marques, Canessa, Siri, Catricala & Cappa | 2008 | fMRI | NO | EXPLICIT | VERBAL | VISUAL | LOW | 21 | semantic features>baseline task |
| Mashal, Vishne, Laor & Titone | 2013 | fMRI | NO | EXPLICIT | VERBAL | VISUAL | HIGH | 14 | novel metaphor>unrelated words |
| | | | | | | | | | conventional metaphor>unrelated words |
| Matchin, Liao, gaston & Lau | 2019 | fMRI | NO | EXPLICIT | VERBAL | VISUAL | HIGH | 20 | verb phrase>list |
| | | | | | | | | | noun phrase>list |
| Matchin, Hammerly & Lau | 2017 | fMRI | NO | EXPLICIT | VERBAL | VISUAL | HIGH | 16 | sentences>word lists |
| | | | | | | | | | sentences>phrases |
| | | | | | | | | | real>pseudoword lists |
| | | | | | | | | | real>pseudoword phrases |

| | | | | | | | | | |
|---|---|---|---|---|---|---|---|---|---|
| | | | | | | | | | real>pseudoword sentences |
| Mellem, Jasmin, Peng & Martin | 2016 | fMRI | NO | IMPLICIT | VERBAL | VISUAL | LOW | 20 | longer>shorter phrase |
| Menz, Blangero, Kunze & Binkofski | 2010 | fMRI | NO | EXPLICIT | NONVERBAL | VISUAL | HIGH | 20 | known>unknown objects |
| Meyer, Alter, Friederici, Lohmann & Yves von Cramon | 2002 | fMRI | NO | IMPLICIT | VERBAL | AUDITORY | HIGH | 14 | word>pseudoword sentence |
| Moberget, Gullesen, Andersson, Ivry & Endestad | 2014 | fMRI | NO | EXPLICIT | VERBAL | VISUAL | HIGH | 32 | incongruent>scrambled sentence |
| | | | | | | | | | congruent>scrambled sentence |
| Moseley, Carota, Hauk, Mohr & Pulvermueller | 2012 | fMRI | YES | IMPLICIT | VERBAL | VISUAL | LOW | 18 | all emotional words>hashmarks |
| | | | | | | | | | abstract emotional words>hashmarks |
| | | | | | | | | | arm+face+emotion words>hashmarks |
| | | | | | | | | | face words>hashmarks |
| | | | | | | | | | arm words>hashmarks |
| Mummery, Patterson, Hodges & Price | 1998 | PET | NO | EXPLICIT | VERBAL | VISUAL | HIGH | 10 | semantic>phonological decision |
| Nakamura et al. | 2000 | PET | YES | EXPLICIT | NONVERBAL | VISUAL | LOW | 7 | familiar faces>fixation cross |
| Nakamura et al. | 2001 | PET | YES | EXPLICIT | VERBAL | AUDITORY | HIGH | 9 | familiar voice>vowel discrimination |
| | | | | | | | | | self voice>vowel discrimination |
| Nichelli, Grafman, Pietrini, Clark, Lee & Miletich | 1995 | PET | NO | EXPLICIT | VERBAL | VISUAL | HIGH | 9 | semantic>orthographic decision |
| Nielson et al. | 2010 | fMRI | YES | EXPLICIT | BOTH | VISUAL | HIGH | 17 | familiar>unfamiliar people |
| Noppeney & Price | 2003 | PET | NO | EXPLICIT | VERBAL | AUDITORY | LOW | 9 | normal>reversed words |
| Orfanidou, Marlsen-Wilson & Davis | 2006 | fMRI | NO | EXPLICIT | VERBAL | AUDITORY | HIGH | 13 | words>pseudowords |
| Pallier, Devauchelle & Dehaene | 2011 | fMRI | NO | EXPLICIT | VERBAL | VISUAL | HIGH | 40 | longer>shorter phrase |
| | | | | | | | | | length of real>pseudoword sentences |
| Peelle, Eason, Schmitter, Schwarzbauer & Davis | 2010 | fMRI | NO | EXPLICIT | VERBAL | AUDITORY | HIGH | 6 | sentences>signal correlated noise |

| | | | | | | | | | |
|---|---|---|---|---|---|---|---|---|---|
| Perani, Schnur, Tettamanti, Gorno-Tempini, Cappa & Fazio | 1999 | PET | NO | EXPLICIT | NONVERBAL | VISUAL | HIGH | 11 | living objects>shapes |
| | | | | | | | | | nonliving objects>shapes |
| Perani, Schnur, Tettamanti, Gorno-Tempini, Cappa & Fazio | 1999 | PET | NO | EXPLICIT | VERBAL | VISUAL | HIGH | 8 | living words>pseudowords |
| | | | | | | | | | nonliving words>pseudowords |
| Perrone-Bertolotti, Kauffmann, Pichat, Vidal & Baciu | 2017 | fMRI | NO | EXPLICIT | VERBAL | VISUAL | HIGH | 24 | words>unreadable font |
| Pilgrim, Fadili, Fletcher & Tyler | 2002 | fMRI | NO | EXPLICIT | VERBAL | VISUAL | HIGH | 14 | words>letter strings |
| Price, Moore, Humphreys & Wise | 1997 | PET | NO | EXPLICIT | VERBAL | VISUAL | HIGH | 6 | semantic>phonological decision |
| Pulvermueller, Cook & Hauk | 2012 | fMRI | NO | IMPLICIT | VERBAL | VISUAL | LOW | 23 | phrases>hashmarks |
| | | | | | | | | | uninflected words>hashmarks |
| | | | | | | | | | inflected words>hashmarks |
| Raettig & Kotz | 2008 | fMRI | NO | EXPLICIT | VERBAL | AUDITORY | HIGH | 16 | real words>pseudowords |
| Raposo, Frade & Alves | 2016 | fMRI | NO | IMPLICIT | VERBAL | VISUAL | HIGH | 18 | semantic>perceptual decision |
| Raposo, Moss, Stamatakis & Tyler | 2009 | fMRI | NO | IMPLICIT | VERBAL | AUDITORY | LOW | 22 | action sentences>SC noise |
| Rapp & Lipka | 2011 | fMRI | NO | IMPLICIT | VERBAL | VISUAL | LOW | 10 | words>checkerboards |
| | | | | | | | | | words>letter strings |
| Redcay, Velnoskey & Rowe | 2016 | fMRI | NO | EXPLICIT | BOTH | VISUAL | HIGH | 24 | meaningful>meaningless stimuli |
| | | | | | | | | | communicative>non-communicative gesture |
| | | | | | | | | | real>pseudoword sentences |
| Rissman, Eliassen & Blumstein | 2003 | fMRI | NO | IMPLICIT | VERBAL | AUDITORY | HIGH | 15 | words>pseudowords |
| Robertson et al. | 2000 | fMRI | NO | IMPLICIT | VERBAL | VISUAL | HIGH | 8 | indefinite article sentence>letter strings |
| | | | | | | | | | definite article sentence>letter strings |
| Rodd, Johnsrude & Davis | 2012 | fMRI | NO | EXPLICIT | VERBAL | AUDITORY | LOW | 15 | speech>SCN |

| | | | | | | | | | |
|---|---|---|---|---|---|---|---|---|---|
| Rodd, Longe, Randall & Tyler | 2010 | fMRI | NO | EXPLICIT | VERBAL | AUDITORY | LOW | 14 | speech>SCN |
| Rogalsky & Hickok | 2009 | fMRI | NO | IMPLICIT | VERBAL | AUDITORY | HIGH | 14 | sentences>word lists |
| Rogalsky, Almeida, Sprouse & Hickok | 2015 | fMRI | NO | IMPLICIT | VERBAL | AUDITORY | HIGH | 15 | words>scrambled |
| Rogers et al. | 2006 | PET | NO | EXPLICIT | BOTH | VISUAL | HIGH | 12 | pictures>scrambled pictures |
| | | | | | | | | | specific-level judgement>baseline |
| Roskies, Fiez, Balota, Raichle & Petersen | 2001 | PET | NO | EXPLICIT | VERBAL | VISUAL | HIGH | 20 | semantic>phonological decision |
| Ross & Olson | 2012 | fMRI | YES | EXPLICIT | NONVERBAL | VISUAL | HIGH | 11 | famous>unknown faces & landmarks |
| Roxbury, McMahon & Copland | 2014 | fMRI | NO | EXPLICIT | VERBAL | AUDITORY | HIGH | 17 | concrete word>pseudoword |
| | | | | | | | | | abstract word>pseudoword |
| Ryan, Cox, Hayes & Nadel | 2008 | fMRI | NO | EXPLICIT | VERBAL | VISUAL | HIGH | 10 | semantic fluency (generate) > crosses |
| | | | | | | | | | semantic fluency (recall) > crosses |
| | | | | | | | | | semantic fluency (recall & geneerate)>crosses |
| Ryan, Lin, Ketcham & Nadel | 2010 | fMRI | NO | EXPLICIT | VERBAL | VISUAL | HIGH | 15 | semantic spatial old>letter judgement |
| | | | | | | | | | semantic spatial new>letter judgement |
| | | | | | | | | | semantic non spatial old>letter judgement |
| | | | | | | | | | semantic non spatial new>letter judgement |
| | | | | | | | | | semantic new>episodic judgement |
| Sabri, Binder, Desai, Medler, Leitl & Liebenthal | 2008 | fMRI | NO | EXPLICIT | VERBAL | AUDITORY | HIGH | 28 | speech>rotated speech |
| | | | | | | | | | words>pseudowords |
| Sachs, Weis, Krings, Huber & Kircher | 2008 | fMRI | NO | EXPLICIT | VERBAL | VISUAL | HIGH | 14 | biased thematic judgement>letters |
| | | | | | | | | | biased taxonomic judgement>letters |

| | | | | | | | | | |
|---|---|---|---|---|---|---|---|---|---|
| | | | | | | | | | balanced taxonomic judgement>letters |
| | | | | | | | | | balanced taxonomic judgement>letters |
| Saur et al. | 2008 | fMRI | NO | IMPLICIT | VERBAL | AUDITORY | HIGH | 33 | word>pseudoword sentences |
| Schell, Zaccarella & Friederici | 2017 | fMRI | NO | EXPLICIT | VERBAL | AUDITORY | HIGH | 21 | phrases>non combinatorial words |
| Schmitt, Auer & Ferstl | 2019 | fMRI | NO | EXPLICIT | VERBAL | AUDITORY | HIGH | 40 | known>unknown language |
| Schuil, Smits & Zwaan | 2013 | fMRI | NO | EXPLICIT | VERBAL | VISUAL | HIGH | 20 | senteces>pseudowords |
| | | | | | | | | | verbs>pseudowords |
| | | | | | | | | | literal sentences>pseudowords |
| | | | | | | | | | nonliteral sentences>pseudowords |
| Scott, Blank, Rosen & Wise | 2000 | PET | NO | IMPLICIT | VERBAL | AUDITORY | HIGH | 8 | intelligible>unintelligible speech |
| Segal & Petrides | 2012 | fMRI | NO | EXPLICIT | NONVERBAL | VISUAL | HIGH | 90 | writing>copying |
| | | | | | | | | | words>pseudowords |
| Seghier, Josse, Leff & Price | 2011 | fMRI | NO | EXPLICIT | BOTH | VISUAL | HIGH | 60 | meaningful>meaningless stimuli |
| Sergent, Otha & Macdonald | 1992 | PET | YES | EXPLICIT | VERBAL | VISUAL | HIGH | 7 | face identity>gender discrimination |
| | | | | | | | | | object recognition>gratings |
| Sheldon, McAndrews, Pruessner & Moscovitch | 2016 | fMRI | NO | EXPLICIT | VERBAL | VISUAL | HIGH | 15 | semantic fluency>perceptual task |
| Simard, Monetta, Nagano-Saito & Monchi | 2013 | fMRI | NO | EXPLICIT | VERBAL | VISUAL | HIGH | 14 | semantic>control macthing |
| | | | | | | | | | semantic matching>phonological decision (syllable rhyme) |
| | | | | | | | | | semantic matching>phonological decision (syllable onset) matching |
| Slioussar, Kireev, Chernigovskaya, Kataeva, Korotkov & Medvedev | 2014 | fMRI | NO | EXPLICIT | VERBAL | VISUAL | HIGH | 21 | real verbs>pseudowords |
| | | | | | | | | | real nouns>pseudowords |

| | | | | | | | | | |
|---|---|---|---|---|---|---|---|---|---|
| Smith, Myers, Sethi, Pantazatos, Yanagihara & Hirsch | 2012 | fMRI | NO | EXPLICIT | VERBAL | VISUAL | HIGH | 14 | semantics>baseline |
| Snijders, Vosse, Kempen, Van Berkum, Petersson & Hagoort | 2009 | fMRI | NO | IMPLICIT | VERBAL | VISUAL | HIGH | 28 | sentences>word lists |
| Stowe, Paans, Wijers, Zwarts, Mulder & Vaalburg | 1999 | PET | NO | IMPLICIT | VERBAL | VISUAL | HIGH | 12 | sentences>scrambled word lists |
| Straube, Green, Weis & Kircher | 2012 | fMRI | NO | IMPLICIT | VERBAL | AUDITORY | HIGH | 16 | known>unknown language |
| | | | | | | | | | iconic>meaningless gesture |
| Stringaris, Medford, Giampietro. Brammer & David | 2007 | fMRI | NO | EXPLICIT | VERBAL | VISUAL | HIGH | 11 | literal>meaningless sentences |
| | | | | | | | | | methaphors>meaningless sentences |
| Sugiura et al. | 2006 | fMRI | YES | EXPLICIT | VERBAL | VISUAL | HIGH | 24 | famous>unfamiliar names |
| | | | | | | | | | personal>unfamiliar names |
| Sugiura et al. | 2001 | PET | YES | EXPLICIT | NONVERBAL | VISUAL | HIGH | 5 | identity discrimination>control |
| | | | | | | | | | identity discrimination>face direction |
| Sugiura et al. | 2008 | fMRI | YES | EXPLICIT | VERBAL | VISUAL | HIGH | 25 | low familiar>unfamiliar names |
| | | | | | | | | | personal familiar>unfamiliar names |
| | | | | | | | | | high familiar>unfamiliar names |
| | | | | | | | | | high familiar>unfamiliar names |
| Sun, Xue, Zhang, Zuo, Chen, Wang, Martin, Wang, Chen, He & Wang | 2017 | fMRI | NO | EXPLICIT | VERBAL | VISUAL | HIGH | 11 | semantic>orthography judgement |
| Szlachta, Bozic, Jelowicka & Marslen-Wilson | 2012 | fMRI | NO | IMPLICIT | VERBAL | VISUAL | LOW | 21 | words>musical rain |
| | | | | | | | | | nouns>musical rain |
| | | | | | | | | | inflected nouns>musical rain |
| Takeichi, Koyama, Terao, Takeuchi, Toyosawa & Murohashi | 2010 | fMRI | NO | IMPLICIT | VERBAL | AUDITORY | HIGH | 23 | speech>reversed speech |

| | | | | | | | | | |
|---|---|---|---|---|---|---|---|---|---|
| | | | | | | | | | speech>modulated speech |
| Taminato, Miura, Sugiura & Kawashima | 2014 | fMRI | NO | EXPLICIT | NONVERBAL | VISUAL | LOW | 35 | object recognition>control task |
| | | | | | | | | | object recognition>control task |
| Taylor, Arsalidou, Bayless, Morris, Evans & Barbeau | 2009 | fMRI | YES | IMPLICIT | NONVERBAL | VISUAL | HIGH | 10 | own>unfamiliar face |
| | | | | | | | | | partner's>unfamiliar face |
| | | | | | | | | | parent's>unfamiliar face |
| Thierry & Price | 2006 | PET | NO | EXPLICIT | VERBAL | AUDITORY | HIGH | 12 | auditory words>speech control(scrambled) |
| | | | | | | | | | auditory sounds>sound control (scrambled) |
| Thierry & Price | 2006 | PET | NO | EXPLICIT | VERBAL | VISUAL | HIGH | 12 | visual words>text control (scrambled letter strings) |
| | | | | | | | | | visual videos>video control (distorted) |
| Tieleman, Seurinck, Deblaere, Vandemaele, Vingerhoets & Achten | 2005 | fMRI | NO | EXPLICIT | VERBAL | VISUAL | HIGH | 22 | self-paced semantic>perceptual decision |
| | | | | | | | | | fixed-paced semantic>perceptual decision |
| Tyler, Stamatakis, Dick, Bright, Fletcher & Moss | 2003 | fMRI | NO | EXPLICIT | VERBAL | VISUAL | HIGH | 12 | animals>baseline |
| | | | | | | | | | tool action words>baseline |
| | | | | | | | | | biological action>baseline |
| Vagharchakian, Dehaene-Lambertz, Pallier & Dehaene | 2012 | fMRI | NO | EXPLICIT | VERBAL | BOTH | HIGH | 16 | intelligible>unintelligble compression rate |
| | | | | | | | | | intelligible>unintelligble compression rate |
| | | | | | | | | | intelligible>unintelligble compression rate |
| Van Ettinger-Veenstra, McAllister, Lundberg, Karlsson & Engstrom | 2016 | fMRI | NO | EXPLICIT | VERBAL | VISUAL | HIGH | 27 | sentences>symbol strings |
| van Leeuwen et al. | 2014 | fMRI | NO | EXPLICIT | BOTH | BOTH | HIGH | 16 | speech>reversed speech |

| | | | | | | | | | |
|---|---|---|---|---|---|---|---|---|---|
| Vignali, Hawelka, Hutzler & Richlan | 2019 | fMRI | NO | EXPLICIT | VERBAL | VISUAL | HIGH | 21 | foveal & parafovealwords>foveal & parafoveal pseudowords |
| Vingerhoets | 2008 | fMRI | NO | IMPLICIT | NONVERBAL | VISUAL | HIGH | 14 | familiar>unfamiliar tools |
| Visser, Jefferies, Embleton & Lambon Ralph | 2012 | fMRI | NO | EXPLICIT | BOTH | VISUAL | HIGH | 15 | semantics>baseline |
| | | | | | | | | | pictures>baseline |
| | | | | | | | | | words>baseline |
| Vitello, Warren, Devlin & Rodd | 2014 | fMRI | NO | EXPLICIT | VERBAL | AUDITORY | LOW | 20 | sentences>SCN |
| von Kriegstein, Eger, Kleinschmidt & Giraud | 2003 | fMRI | NO | EXPLICIT | VERBAL | AUDITORY | HIGH | 14 | sentence>speech envelope |
| Wang, Zhao, Zevin & Yang | 2016 | fMRI | NO | EXPLICIT | VERBAL | VISUAL | HIGH | 16 | words>nonsense strokes |
| Weiss, Katzir & Bitan | 2015 | fMRI | NO | IMPLICIT | VERBAL | VISUAL | LOW | 18 | pointed words>asterisks |
| | | | | | | | | | unpointed words>asterisks |
| Welcome & Joanisse | 2012 | fMRI | NO | EXPLICIT | VERBAL | VISUAL | HIGH | 20 | semantic>phonological/orthographic decision |
| Wende, Straube, Stratmann, Sommer, Kircher & Nagels | 2012 | fMRI | NO | EXPLICIT | VERBAL | VISUAL | HIGH | 18 | semantic>phonological fluency |
| Wirth, Jann, Dierks, Federspiel, Wiest & Horn | 2011 | fMRI | NO | EXPLICIT | VERBAL | VISUAL | HIGH | 19 | semantic>phonological & perceptual decision |
| Wright et al. | 2008 | fMRI | NO | EXPLICIT | BOTH | VISUAL | HIGH | 10 | semantic>perceptual matching |
| | | | | | | | | | semantic>perceptual matching |
| | | | | | | | | | semantic>perceptual matching |
| Wright, Randall, Marslen-Wilson & Tyler | 2011 | fMRI | NO | BOTH | VERBAL | AUDITORY | LOW | 14 | speech>musical rain |
| Wu, Mai, Tang, Ge, Luo & Liu | 2013 | fMRI | NO | IMPLICIT | VERBAL | VISUAL | LOW | 19 | arm words>checkerboard |
| | | | | | | | | | leg words>checkerboard |
| | | | | | | | | | mouth words>checkerboard |
| Xiao et al. | 2005 | fMRI | NO | EXPLICIT | VERBAL | AUDITORY | HIGH | 14 | words>pseudowords |
| Yang, Li, Fang, Shu, Liu & Chen | 2016 | fMRI | NO | IMPLICIT | VERBAL | VISUAL | LOW | 20 | opaque idioms>hashmarks |
| | | | | | | | | | transparent idioms>hashmarks |
| | | | | | | | | | literal phrases>hashmarks |

| | | | | | | | | | |
|---|---|---|---|---|---|---|---|---|---|
| Zaccarella & Friederici | 2015 | fMRI | NO | EXPLICIT | VERBAL | VISUAL | HIGH | 22 | words>pseudowords |
| Zhang, Liu & Zhang | 2014 | fMRI | NO | EXPLICIT | VERBAL | VISUAL | LOW | 18 | nonliving words>asterisks |
| | | | | | | | | | living words>asterisks |
| Zhang, Xiao & Weng | 2012 | fMRI | NO | EXPLICIT | VERBAL | VISUAL | HIGH | 14 | words>pseudowords |
| Zhuang & Devereux | 2017 | fMRI | NO | EXPLICIT | VERBAL | AUDITORY | HIGH | 16 | phrases>words |
| Zou, Packard, Xia, Liu & Shu | 2016 | fMRI | NO | IMPLICIT | VERBAL | AUDITORY | LOW | 17 | speech>tone |
| | | | | | | | | | speech>tone |
| | | | | | | | | | speech>tone |
| | | | | | | | | | identical speech>tone |
| | | | | | | | | | words>tone |
| Zvyagintsev, Clemens, Chechko, Mathiak, Sack & Mathiak | 2013 | fMRI | NO | IMPLICIT | NONE | NONE | HIGH | 15 | visual imagery>counting |
| | | | | | | | | | auditory imagery>counting |
| EXCLUDED CONTRASTS WITH REST | | | | | | | | | |
| Alain, He & Grady | 2008 | fMRI | NO | EXPLICIT | VERBAL | AUDITORY | REST | 16 | sound category>rest |
| Assadollahi, Meinzer, Flaisch, Obleser & Rockstroh | 2009 | fMRI | NO | IMPLICIT | VERBAL | VISUAL | REST | 20 | nouns followed by 1 argument verbs>fixation cross (rest) |
| | | | | | | | | | nouns followed by 3 argument verbs>fixation cross (rest) |
| Cai, Kochiyama, Osaka & Wu | 2007 | fMRI | NO | IMPLICIT | VERBAL | AUDITORY | REST | 15 | words>rest |
| Chan, Tang, Tang, Lee, Lo & Kwong | 2009 | fMRI | NO | IMPLICIT | VERBAL | VISUAL | REST | 22 | synonyms>rest |
| Dapretto & Bookheimer | 1999 | fMRI | NO | EXPLICIT | VERBAL | AUDITORY | REST | 8 | semantic>rest |
| D'Arcy, Bolster, Ryner, Mazerolle, Grant & Song | 2007 | fMRI | NO | EXPLICIT | BOTH | VISUAL | REST | 10 | basic-level living objects>baseline |
| | | | | | | | | | basic-level non-living objects>baseline |
| | | | | | | | | | superordinate-level living objects>baseline |
| | | | | | | | | | superordinate-level non-living objects>baseline |

| | | | | | | | | | |
|---|---|---|---|---|---|---|---|---|---|
| Emmorey, Xu, Gannon, Goldin-Meadow & Braun | 2010 | fMRI | NO | IMPLICIT | NONVERBAL | VISUAL | REST | 14 | meaningful pantomimes>rest |
| Engelien et al. | 2006 | PET | NO | IMPLICIT | NONVERBAL | AUDITORY | REST | 6 | meaningful>rest |
| Groussard et al. | 2010 | PET | NO | EXPLICIT | VERBAL | VISUAL | REST | 11 | verbal semantic>rest |
| | | | | | | | | | verbal semantic>rest |
| Hagoort et al. | 1999 | PET | NO | IMPLICIT | VERBAL | VISUAL | REST | 10 | words>fixation |
| Harrington, Farias & Davis | 2009 | fMRI | NO | EXPLICIT | NONVERBAL | VISUAL | REST | 8 | familiar objects>rest |
| Heim, Eickhoff & Amunts | 2008 | fMRI | NO | EXPLICIT | VERBAL | VISUAL | REST | 28 | semantic fluency>rest |
| Jensen, Hargreaves, Bass, Pexman, Goodyear & Federico | 2011 | fMRI | NO | EXPLICIT | VERBAL | VISUAL | REST | 12 | words>rest |
| Jeon, Lee, Kim & Cho | 2009 | fMRI | NO | EXPLICIT | VERBAL | VISUAL | REST | 16 | synonyms>rest |
| | | | | | | | | | antonyms>rest |
| | | | | | | | | | English synonyms>rest |
| | | | | | | | | | Korean synonyms>nonwords |
| | | | | | | | | | honorific words>nonwords |
| Kim et al. | 2009 | fMRI | NO | EXPLICIT | VERBAL | VISUAL | REST | 36 | sentences>rest |
| Ludersdorfer, Wimmer, Richlan, Schurz, Hutzler & Kronbichler | 2016 | fMRI | NO | EXPLICIT | VERBAL | AUDITORY | REST | 29 | Auditory words orthographic>rest |
| | | | | | | | | | Auditory words semantic>rest |
| Matchin, Liao, gaston & Lau | 2019 | fMRI | NO | EXPLICIT | VERBAL | VISUAL | REST | 20 | sentence>rest |
| Perrone-Bertolotti, Kauffmann, Pichat, Vidal & Baciu | 2017 | fMRI | NO | EXPLICIT | VERBAL | VISUAL | REST | 24 | words>fixation |
| Stowe, Paans, Wijers, Zwarts, Mulder & Vaalburg | 1999 | PET | NO | IMPLICIT | VERBAL | VISUAL | REST | 12 | sentences>rest |
| Taylor, Arsalidou, Bayless, Morris, Evans & Barbeau | 2009 | fMRI | YES | IMPLICIT | NONVERBAL | VISUAL | REST | 10 | own face>baseline (rest) |
| | | | | | | | | | parents face>baseline (rest) |
| | | | | | | | | | famous face>baseline (rest) |
| Wende, Straube, Stratmann, Sommer, Kircher & Nagels | 2012 | fMRI | NO | EXPLICIT | VERBAL | VISUAL | REST | 18 | causal fluency>rest |
| | | | | | | | | | free association>rest |
| Wright et al. | 2008 | fMRI | NO | EXPLICIT | VERBAL | VISUAL | REST | 15 | words>rest |

| | | | | | | | | | |
|---|---|---|---|---|---|---|---|---|---|
| | | | | | | | | | pictures>rest |
| Xiao et al. | 2005 | fMRI | NO | EXPLICIT | VERBAL | AUDITORY | REST | 14 | words>rest |
| Zhang, Xiao & Weng | 2012 | fMRI | NO | EXPLICIT | VERBAL | VISUAL | REST | 14 | words>rest |

**The references to access the listed studies are listed at the end of the document*

***Table M3 SC*** *List of studies included in the* ***semantic cognition high baselines only (low baselines and rest excluded*** *(N = 170) meta-analysis.* ***Note:*** *The list of excluded studies using rest and low-level baselines can be seen at the end of the document highlighted in grey. SC= Semantic Cognition, N= Sample Size*

| *Authors | Year | Imaging Method | Social Content | Instructional Cue | Stimulus Domain | Sensory Input Modality | Baseline Type | N | Contrast |
|---|---|---|---|---|---|---|---|---|---|
| Abraham et al. | 2012 | fMRI | NO | EXPLICIT | VERBAL | VISUAL | HIGH | 19 | high&low divergent thinking (semantic)>1&2 back letter identity (working memory) |
| Alain, He & Grady | 2008 | fMRI | NO | EXPLICIT | VERBAL | AUDITORY | HIGH | 16 | sound category>sound location |
| Assadollahi, Meinzer, Flaisch, Obleser & Rockstroh | 2009 | fMRI | NO | IMPLICIT | VERBAL | VISUAL | HIGH | 20 | nouns followed by 1&3 argument verbs>letter strings followed by 1&3argument verbs |
| Axmacher, Bialleck, Weber, Helmstaedter, Elger & Fell | 2009 | fMRI | NO | EXPLICIT | VERBAL | VISUAL | HIGH | 32 | word decision>spatial decision |
| Bagga et al. | 2013 | fMRI | NO | EXPLICIT | VERBAL | VISUAL | HIGH | 18 | semantic>case matching judgement |
| Baumgaertener, Weiller & Buchel | 2002 | fMRI | NO | EXPLICIT | VERBAL | VISUAL | HIGH | 9 | word>pseudoword in sentence |
| Baumgaertner et al. | 2007 | fMRI | NO | IMPLICIT | VERBAL | AUDITORY | HIGH | 19 | sentences>reversed sentences |
| Bautista & Wilson | 2016 | fMRI | NO | IMPLICIT | VERBAL | AUDITORY | HIGH | 12 | videos>scrambled videos clear>scrambled rotated speech |
| Binder, Frost, Hammeke, Bellgowan, Rao & Cox | 1999 | fMRI | NO | EXPLICIT | VERBAL | AUDITORY | HIGH | 30 | semantic>phonological decision |
| Binder et al. | 2003 | fMRI | NO | EXPLICIT | VERBAL | VISUAL | HIGH | 24 | word>nonword |
| Birn et al. | 2010 | fMRI | NO | EXPLICIT | VERBAL | VISUAL | HIGH | 14 | category>letter fluency |
| Bonhage, Fiebach, Bahlmann & Mueller | 2014 | fMRI | NO | EXPLICIT | VERBAL | VISUAL | HIGH | 18 | sentence fragments>ungrammatical word strings |
| Bonhage, Mueller, Friederici & Fiebach | 2015 | fMRI | NO | EXPLICIT | VERBAL | VISUAL | HIGH | 18 | sentences>jabberwocky sentences |
| Booth et al. | 2006 | fMRI | NO | EXPLICIT | VERBAL | VISUAL | HIGH | 13 | meaning (semantic) judgement>rhyming( phonological) judgement |
| Bozic & Marslen-Wilson | 2013 | fMRI | NO | IMPLICIT | VERBAL | AUDITORY | HIGH | 13 | speech>musical rain |
| Brambati, Benoit, Monetta, Belleville & Joubert | 2010 | fMRI | YES | EXPLICIT | BOTH | BOTH | HIGH | 12 | general & specific occupation judgement>baseline (scrambled face) |
| Bruffaerts, Dupont, Peeters, De Deyne, Storms & Vandenberghe | 2013 | fMRI | NO | EXPLICIT | BOTH | VISUAL | HIGH | 19 | real > scrambled pictures & words |

| | | | | | | | | | |
|---|---|---|---|---|---|---|---|---|---|
| Bulut, Hung, Tzeng & Wu | 2017 | fMRI | NO | IMPLICIT | VERBAL | VISUAL | HIGH | 20 | sentences>unstructured word lists |
| | | | | | | | | | sentences>unstructured character list |
| Cai, Kochiyama, Osaka & Wu | 2007 | fMRI | NO | IMPLICIT | VERBAL | AUDITORY | HIGH | 15 | words>nonsense words |
| Cappa, Perani, Schnur, Tettamanti & Fazio | 1998 | PET | NO | EXPLICIT | VERBAL | VISUAL | HIGH | 13 | words>pseudowords |
| | | | | | | | | | animal visual knowledge>pseudowords |
| | | | | | | | | | tool visual knowledge>pseudowords |
| | | | | | | | | | animal associative knowledge>pseudowords |
| | | | | | | | | | tools functional knowledge>pseudowords |
| Chan, Tang, Tang, Lee, Lo & Kwong | 2009 | fMRI | NO | IMPLICIT | VERBAL | VISUAL | HIGH | 22 | synonyms>pseudocharacters |
| | | | | | | | | | synonyms>Korean characters (unknown) |
| Chou, Chen, Wu & Booth | 2009 | fMRI | NO | EXPLICIT | VERBAL | VISUAL | HIGH | 31 | related words>false font |
| | | | | | | | | | unrelated words>false font |
| Chou, Chen, Wu & Booth | 2009 | fMRI | NO | EXPLICIT | VERBAL | VISUAL | HIGH | 32 | related words>false font |
| | | | | | | | | | unrelated words>false font |
| Chouinard, Morrissey, Kohler & Goodale | 2008 | fMRI | NO | EXPLICIT | NONVERBAL | VISUAL | HIGH | 14 | objects>scrambled objects |
| Chow, Kaup, Raabe & Greenlee | 2008 | fMRI | NO | EXPLICIT | VERBAL | VISUAL | HIGH | 15 | predictive&normal reading>pseudoword reading |
| | | | | | | | | | normal reading>pseudoword reading |
| | | | | | | | | | predictive reading>pseudoword reading |
| Christensen, Antonucci, Lockwood, Kittleson & Plante | 2008 | fMRI | NO | EXPLICIT | VERBAL | AUDITORY | HIGH | 14 | diotic listening>reversed speech |
| | | | | | | | | | dichotic listening>reversed speech |
| Clos, Langner, Meyer, Oechslin, Zilles & Eickhoff | 2014 | fMRI | NO | EXPLICIT | VERBAL | AUDITORY | HIGH | 29 | intelligibilty based on cue > unintelligible |
| Damasio, Tranel, Grabowski, Adolphs & Damasio | 2004 | PET | YES | EXPLICIT | NONVERBAL | VISUAL | HIGH | 55 | persons>face orientation judgement |
| | | | | | | | | | animals>scrambled pictures |
| | | | | | | | | | tools>scrambled pictures |
| Davis, Meunier & Marslen-Wilson | 2004 | fMRI | NO | EXPLICIT | VERBAL | VISUAL | HIGH | 11 | words>letter strings |
| Demonet et al. | 1992 | PET | NO | EXPLICIT | VERBAL | AUDITORY | HIGH | 9 | words>phonemes |
| Devlin, Matthews & Rushworth | 2003 | fMRI | NO | EXPLICIT | VERBAL | VISUAL | HIGH | 12 | semantic>phonological judgement |

| | | | | | | | | | |
|---|---|---|---|---|---|---|---|---|---|
| Devlin et al. | 2002 | PET | NO | EXPLICIT | VERBAL | VISUAL | HIGH | 12 | all semantic>letter detection<br>all semantic>letter detection |
| Devlin et al. | 2002 | PET | NO | EXPLICIT | VERBAL | VISUAL | HIGH | 8 | all semantic>letter categorisation |
| Devlin et al. | 2002 | fMRI | NO | EXPLICIT | VERBAL | VISUAL | HIGH | 8 | all semantic> letter categorisation |
| Devlin et al. | 2000 | PET | NO | EXPLICIT | VERBAL | VISUAL | HIGH | 8 | semantic categorisation>letter categorisation |
| Devlin et al. | 2000 | fMRI | NO | EXPLICIT | VERBAL | VISUAL | HIGH | 8 | semantic categorisation>letter categorisation |
| Diaz & McCarthy | 2009 | fMRI | NO | IMPLICIT | VERBAL | VISUAL | HIGH | 16 | all words>nonwords |
| Ebisch et al. | 2007 | fMRI | NO | EXPLICIT | BOTH | VISUAL | HIGH | 17 | functional&visuospatial pictures&words>meaningless drawings&pseudowords |
| Elfgren, Westen, Passant, Larsson, Mannfolk & Fransson | 2006 | fMRI | YES | EXPLICIT | NONVERBAL | VISUAL | HIGH | 15 | familiar>unfamiliar faces (identification) |
| Emmorey, Weisberg, McCullough & Petrich | 2013 | fMRI | NO | EXPLICIT | VERBAL | VISUAL | HIGH | 14 | words(semantic judgement)>false fonts<br>words(phonological)>false fonts |
| Emmorey, Xu, Gannon, Goldin-Meadow & Braun | 2010 | fMRI | NO | IMPLICIT | NONVERBAL | VISUAL | HIGH | 14 | meaningful pantomimes>unknown sign language |
| Engelien et al. | 2006 | PET | NO | IMPLICIT | NONVERBAL | AUDITORY | HIGH | 6 | meaningful>meaningless sounds |
| Erb, Henry, Eisner & Obleser | 2013 | fMRI | NO | EXPLICIT | VERBAL | AUDITORY | HIGH | 30 | speech>vocoded speech |
| Europa, Gitelman, Kiran & Thompson | 2019 | fMRI | NO | EXPLICIT | BOTH | BOTH | HIGH | 21 | sentences>baseline (reversed sentences) |
| Friederici, Kotz, Scott & Obleser | 2010 | fMRI | NO | IMPLICIT | VERBAL | AUDITORY | HIGH | 17 | intelligible speech>rotated speech |
| Friese, Rutschmann, Raabe & Schmalhofer | 2008 | fMRI | NO | EXPLICIT | VERBAL | VISUAL | HIGH | 13 | words>pseudowords |
| Garbin, Collina & Tabossi | 2012 | fMRI | NO | EXPLICIT | VERBAL | VISUAL | HIGH | 12 | object noun>pseudoword<br>event noun>pseudoword<br>verb>pseudoword |
| Garn, Allen & Larsen | 2009 | fMRI | NO | EXPLICIT | NONVERBAL | VISUAL | HIGH | 26 | pictures>scrambled pictures<br>plants>scrambled pictures<br>tools>scrambled pictures |
| Geranmayeh, Brownsett, Leech, Beckmann, Woodhead & Wise | 2012 | fMRI | NO | EXPLICIT | VERBAL | VISUAL | HIGH | 19 | speech>tongue movements |
| Gerlach, Law, Gade & Paulson | 1999 | PET | NO | EXPLICIT | NONVERBAL | VISUAL | HIGH | 15 | object decision>pattern discrimination |
| Gesierich et al. | 2012 | fMRI | YES | EXPLICIT | NONVERBAL | VISUAL | HIGH | 21 | familiar>scrambled faces |

| | | | | | | | | | |
|---|---|---|---|---|---|---|---|---|---|
| | | | | | | | | | familiar > unfamiliar faces |
| | | | | | | | | | familiar>unfamiliar faces |
| Giraud et al. | 2004 | fMRI | NO | EXPLICIT | VERBAL | AUDITORY | HIGH | 8 | natural speech>speech envelope |
| Gitelman, Nobre, Sonty, Parrish & Mesulam | 2005 | fMRI | NO | EXPLICIT | VERBAL | VISUAL | HIGH | 14 | semantic>control task |
| Gorno-Tempini et al. | 1998 | PET | YES | EXPLICIT | NONVERBAL | VISUAL | HIGH | 6 | famous faces>controls |
| | | | | | | | | | famous names>controls |
| | | | | | | | | | double famous proper names>controls |
| | | | | | | | | | non-famous faces>controls |
| | | | | | | | | | Non-famous names>controls |
| | | | | | | | | | Double common names>controls |
| Grabowski, Damasio, Tranel, Boles Ponto, Hichwa & Damasio | 2001 | PET | YES | EXPLICIT | NONVERBAL | VISUAL | HIGH | 10 | naming persons>building orientation judgement |
| | | | | | | | | | naming landmarks>face orientation judgement |
| | | | | | | | | | naming persons>face orientation judgement |
| | | | | | | | | | naming unique entities>baseline |
| Graves, Binder, Desai, Conant & Seidenberg | 2010 | fMRI | NO | EXPLICIT | VERBAL | VISUAL | HIGH | 23 | forward>reverse phrases |
| Graves, Binder, Desai, Conant & Seidenberg | 2010 | fMRI | NO | EXPLICIT | VERBAL | VISUAL | HIGH | 22 | forward>reverse phrases |
| Grindrod, Garnett, Malyutina & den Ouden | 2014 | fMRI | NO | EXPLICIT | VERBAL | VISUAL | HIGH | 23 | words>nonwords |
| Grossman et al. | 2002a | fMRI | NO | IMPLICIT | VERBAL | VISUAL | HIGH | 16 | all nouns > pseudowords |
| | | | | | | | | | implements>pseudowords |
| | | | | | | | | | animals>pseudowords |
| | | | | | | | | | abstract>pseudowords |
| Grossman et al., 2002b | 2002b | fMRI | NO | IMPLICIT | VERBAL | VISUAL | HIGH | 16 | verbs>pseudowords |
| Groussard et al. | 2010 | PET | NO | EXPLICIT | VERBAL | VISUAL | HIGH | 11 | verbal semantics>verbal reference |
| | | | | | | | | | musical semantic>musical reference |
| Gurd et al. | 2002 | fMRI | NO | EXPLICIT | VERBAL | AUDITORY | HIGH | 11 | category>rote fluency |
| Haberling, Corballis &Corballis | 2016 | fMRI | NO | EXPLICIT | NONVERBAL | VISUAL | HIGH | 92 | meaningful pantomimes>unknown sign language |
| | | | | | | | | | meaningful pantomimes>dog videos |
| | | | | | | | | | synonyms>letter strings |
| Hagoort et al. | 1999 | PET | NO | IMPLICIT | VERBAL | VISUAL | HIGH | 10 | words>pseudowords |
| Harrington, Farias & Davis | 2009 | fMRI | NO | EXPLICIT | NONVERBAL | VISUAL | HIGH | 8 | familiar>non objects |
| Hartung, Hagoort & Willems | 2017 | fMRI | NO | IMPLICIT | VERBAL | AUDITORY | HIGH | 52 | first-person speech>unintelligible reversed speech |

| | | | | | | | | | |
|---|---|---|---|---|---|---|---|---|---|
| | | | | | | | | | third-person speech>unintelligible reversed speech |
| Heim, Eickhoff & Amunts | 2008 | fMRI | NO | EXPLICIT | VERBAL | VISUAL | HIGH | 28 | semantic > phonological fluency |
| Henke et al. | 1999 | PET | NO | EXPLICIT | VERBAL | VISUAL | HIGH | 12 | associative word learning>single word encoding |
| Herbster et al. | 1997 | PET | NO | EXPLICIT | VERBAL | VISUAL | HIGH | 10 | Irregular>zero order speak |
| | | | | | | | | | regular>zero order speak |
| | | | | | | | | | irregular+regular>zero order speak |
| Hervais-Adelman, Carlyon, Johnsrude & Davis | 2012 | fMRI | NO | EXPLICIT | VERBAL | AUDITORY | HIGH | 15 | clear>vocoded speech (uninteligible) |
| Hocking, McMahon & de Zubicaray | 2011 | fMRI | NO | EXPLICIT | NONVERBAL | AUDITORY | HIGH | 13 | all environmental sounds>perceptual baseline |
| Holle, Gunter, Rueschemeyer, Hennenlotter & Iacoboni | 2008 | fMRI | NO | EXPLICIT | BOTH | BOTH | HIGH | 17 | iconic gesture of dominant meaning>grooming |
| | | | | | | | | | iconic gesture of subordinate meaning>grooming |
| Homae, Yahata & Sakai | 2003 | fMRI | NO | EXPLICIT | VERBAL | AUDITORY | HIGH | 10 | auditory sentences>auditory non words |
| | | | | | | | | | visual sentences>visual non words words |
| Husain, Patkin, Kim, Braun & Horwitz | 2012 | fMRI | NO | EXPLICIT | NONVERBAL | VISUAL | HIGH | 16 | meaningful iconic>meaningless gestures |
| Jackson, Hoffman, Pobric, Lambon Ralph | 2015 | fMRI | NO | EXPLICIT | VERBAL | VISUAL | HIGH | 24 | words>letter strings |
| Jensen, Hargreaves, Bass, Pexman, Goodyear & Federico | 2011 | fMRI | NO | EXPLICIT | VERBAL | VISUAL | HIGH | 12 | words>pseudowords |
| Jeon, Lee, Kim & Cho | 2009 | fMRI | NO | EXPLICIT | VERBAL | VISUAL | HIGH | 16 | synonys>nonwords |
| | | | | | | | | | antonyms>nonwords |
| | | | | | | | | | English synonyma>nonwords |
| | | | | | | | | | Korean synonyms>nonwords |
| | | | | | | | | | honorific words>nonwords |
| Joubert et al. | 2004 | fMRI | NO | IMPLICIT | VERBAL | VISUAL | HIGH | 10 | low frequency words>nonwords |
| | | | | | | | | | high frequency words>consonant strings |
| | | | | | | | | | low frequency words>consonant strings |
| Kang et al. | 2006 | PET | NO | EXPLICIT | VERBAL | BOTH | HIGH | 17 | audio-visual speech>noises and facial movements |
| | | | | | | | | | visual speech>facial movements (chewing gum) |

| | | | | | | | | | |
|---|---|---|---|---|---|---|---|---|---|
| Khader, Jost, Mertens, Bien & Roesler | 2010 | fMRI | NO | EXPLICIT | VERBAL | VISUAL | HIGH | 16 | noun>rhyme generation |
| | | | | | | | | | verb>rhyme generation |
| | | | | | | | | | noun generation>letter detection |
| | | | | | | | | | verb generation>letter detection |
| Kim et al. | 2009 | fMRI | NO | EXPLICIT | VERBAL | VISUAL | HIGH | 36 | sentences>word lists |
| Kinno, Kawamura, Shioda & Sakai | 2008 | fMRI | NO | EXPLICIT | BOTH | VISUAL | HIGH | 14 | canonical sentence>picture & letter strings |
| | | | | | | | | | active sentence>picture & letter strings |
| | | | | | | | | | passive sentence>picture & letter strings |
| Kotz, Cappa, Von Cramon & Friederici | 2002 | fMRI | NO | EXPLICIT | VERBAL | AUDITORY | HIGH | 13 | words>pseudowords |
| Kuchinke et al. | 2005 | fMRI | YES | IMPLICIT | VERBAL | VISUAL | HIGH | 20 | emotion word>nonword |
| Kumar | 2016 | fMRI | NO | EXPLICIT | VERBAL | VISUAL | HIGH | 20 | abstract+concrete words>pseudowords |
| | | | | | | | | | abstract words>pseudowords |
| Kuperberg et al. | 2000 | fMRI | NO | EXPLICIT | VERBAL | AUDITORY | HIGH | 4 | sentences>words strings |
| Leff, Schofield, Stephan, Crinion, Friston & Price | 2008 | fMRI | NO | IMPLICIT | VERBAL | AUDITORY | HIGH | 26 | speech>reversed speech |
| Leung & Alain | 2011 | fMRI | NO | EXPLICIT | NONVERBAL | AUDITORY | HIGH | 16 | semantic>location matching |
| Leveroni et al. | 2000 | fMRI | YES | EXPLICIT | NONVERBAL | VISUAL | HIGH | 11 | familiar faces>foils(never seen faces) |
| | | | | | | | | | newly learned faces>foils(never seen faces) |
| Lin, Wang, Zhao, Liu, Li & Bi | 2015 | fMRI | NO | EXPLICIT | VERBAL | VISUAL | HIGH | 20 | words>pseudowords |
| Liuzzi et al. | 2017 | fMRI | NO | EXPLICIT | VERBAL | BOTH | HIGH | 18 | semantic judgement > input modality detection |
| Ludersdorfer, Schurz, Richlan, Kronbichler & Wimmer | 2013 | fMRI | NO | EXPLICIT | VERBAL | VISUAL | HIGH | 29 | words>false fonts |
| | | | | | | | | | words>pseudowords |
| | | | | | | | | | speech>reversed speech |
| | | | | | | | | | words>pseudowords |
| Malins, Gumkowski, Buis, Molfese, Rueckl, Frost, Pugh, Morris & Mencl | 2016 | fMRI | NO | IMPLICIT | VERBAL | VISUAL | HIGH | 18 | unrelated>pseudowords |
| Mashal, Vishne, Laor & Titone | 2013 | fMRI | NO | EXPLICIT | VERBAL | VISUAL | HIGH | 14 | novel metaphor>unrelated words |
| | | | | | | | | | conventional metaphor>unrelated words |
| Matchin, Liao, gaston & Lau | 2019 | fMRI | NO | EXPLICIT | VERBAL | VISUAL | HIGH | 20 | verb phrase>list |
| | | | | | | | | | noun phrase>list |

| | | | | | | | | | |
|---|---|---|---|---|---|---|---|---|---|
| Matchin, Hammerly & Lau | 2017 | fMRI | NO | EXPLICIT | VERBAL | VISUAL | HIGH | 16 | sentences>word lists<br>sentences>phrases<br>real>pseudoword lists<br>real>pseudoword phrases<br>real>pseudoword sentences |
| Menz, Blangero, Kunze & Binkofski | 2010 | fMRI | NO | EXPLICIT | NONVERBAL | VISUAL | HIGH | 20 | known>unknown objects |
| Meyer, Alter, Friederici, Lohmann & Yves von Cramon | 2002 | fMRI | NO | IMPLICIT | VERBAL | AUDITORY | HIGH | 14 | word>pseudoword sentence |
| Moberget, Gullesen, Andersson, Ivry & Endestad | 2014 | fMRI | NO | EXPLICIT | VERBAL | VISUAL | HIGH | 32 | incongruent>scrambled sentence |
| | | | | | | | | | congruent>scrambled sentence |
| Mummery, Patterson, Hodges & Price | 1998 | PET | NO | EXPLICIT | VERBAL | VISUAL | HIGH | 10 | semantic>phonological decision |
| Nakamura et al. | 2001 | PET | YES | EXPLICIT | VERBAL | AUDITORY | HIGH | 9 | familiar voice>vowel discrimination<br>self voice>vowel discrimination |
| Nichelli, Grafman, Pietrini, Clark, Lee & Miletich | 1995 | PET | NO | EXPLICIT | VERBAL | VISUAL | HIGH | 9 | semantic>orthographic decision |
| Nielson et al. | 2010 | fMRI | YES | EXPLICIT | BOTH | VISUAL | HIGH | 17 | familiar>unfamiliar people |
| Orfanidou, Marlsen-Wilson & Davis | 2006 | fMRI | NO | EXPLICIT | VERBAL | AUDITORY | HIGH | 13 | words>pseudowords |
| Pallier, Devauchelle & Dehaene | 2011 | fMRI | NO | EXPLICIT | VERBAL | VISUAL | HIGH | 40 | longer>shorter phrase |
| | | | | | | | | | length of real>pseudoword sentences |
| Peelle, Eason, Schmitter, Schwarzbauer & Davis | 2010 | fMRI | NO | EXPLICIT | VERBAL | AUDITORY | HIGH | 6 | sentences>signal correlated noise |
| Perani, Schnur, Tettamanti, Gorno-Tempini, Cappa & Fazio | 1999 | PET | NO | EXPLICIT | NONVERBAL | VISUAL | HIGH | 11 | living objects>shapes |
| | | | | | | | | | nonliving objects>shapes |
| Perani, Schnur, Tettamanti, Gorno-Tempini, Cappa & Fazio | 1999 | PET | NO | EXPLICIT | VERBAL | VISUAL | HIGH | 8 | living words>pseudowords |
| | | | | | | | | | nonliving words>pseudowords |
| Perrone-Bertolotti, Kauffmann, Pichat, Vidal & Baciu | 2017 | fMRI | NO | EXPLICIT | VERBAL | VISUAL | HIGH | 24 | words>unreadable font |

| | | | | | | | | | |
|---|---|---|---|---|---|---|---|---|---|
| Pilgrim, Fadili, Fletcher & Tyler | 2002 | fMRI | NO | EXPLICIT | VERBAL | VISUAL | HIGH | 14 | words>letter strings |
| Price, Moore, Humphreys & Wise | 1997 | PET | NO | EXPLICIT | VERBAL | VISUAL | HIGH | 6 | semantic>phonological decision |
| Raettig & Kotz | 2008 | fMRI | NO | EXPLICIT | VERBAL | AUDITORY | HIGH | 16 | real words>pseudowords |
| Raposo, Frade & Alves | 2016 | fMRI | NO | IMPLICIT | VERBAL | VISUAL | HIGH | 18 | semantic>perceptual decision |
| Rapp & Lipka | 2011 | fMRI | NO | IMPLICIT | VERBAL | VISUAL | HIGH | 10 | words>letter strings |
| Redcay, Velnoskey & Rowe | 2016 | fMRI | NO | EXPLICIT | BOTH | VISUAL | HIGH | 24 | meaningful>meaningless stimuli |
| | | | | | | | | | communicative>non-communicative gesture |
| | | | | | | | | | real>pseudoword sentences |
| Rissman, Eliassen & Blumstein | 2003 | fMRI | NO | IMPLICIT | VERBAL | AUDITORY | HIGH | 15 | words>pseudowords |
| Robertson et al. | 2000 | fMRI | NO | IMPLICIT | VERBAL | VISUAL | HIGH | 8 | indefinite article sentence>letter strings |
| | | | | | | | | | definite article sentence>letter strings |
| Rogalsky & Hickok | 2009 | fMRI | NO | IMPLICIT | VERBAL | AUDITORY | HIGH | 14 | sentences>word lists |
| Rogalsky, Almeida, Sprouse & Hickok | 2015 | fMRI | NO | IMPLICIT | VERBAL | AUDITORY | HIGH | 15 | words>scrambled |
| Rogers et al. | 2006 | PET | NO | EXPLICIT | BOTH | VISUAL | HIGH | 12 | pictures>scrambled pictures |
| | | | | | | | | | specific-level judgement>baseline |
| Roskies, Fiez, Balota, Raichle & Petersen | 2001 | PET | NO | EXPLICIT | VERBAL | VISUAL | HIGH | 20 | semantic>phonological decision |
| Ross & Olson | 2012 | fMRI | YES | EXPLICIT | NONVERBAL | VISUAL | HIGH | 11 | famous>unknown faces & landmarks |
| Roxbury, McMahon & Copland | 2014 | fMRI | NO | EXPLICIT | VERBAL | AUDITORY | HIGH | 17 | concrete word>pseudoword |
| | | | | | | | | | abstract word>pseudoword |
| Ryan, Cox, Hayes & Nadel | 2008 | fMRI | NO | EXPLICIT | VERBAL | VISUAL | HIGH | 10 | semantic fluency (generate) > crosses |
| | | | | | | | | | semantic fluency (recall) > crosses |
| | | | | | | | | | semantic fluency (recall & geneerate)>crosses |
| Ryan, Lin, Ketcham & Nadel | 2010 | fMRI | NO | EXPLICIT | VERBAL | VISUAL | HIGH | 15 | semantic spatial old>letter judgement |
| | | | | | | | | | semantic spatial new>letter judgement |
| | | | | | | | | | semantic non spatial old>letter judgement |
| | | | | | | | | | semantic non spatial new>letter judgement |
| | | | | | | | | | semantic new>episodic judgement |
| Sabri, Binder, Desai, Medler, Leitl & Liebenthal | 2008 | fMRI | NO | EXPLICIT | VERBAL | AUDITORY | HIGH | 28 | speech>rotated speech |
| | | | | | | | | | words>pseudowords |
| Sachs, Weis, Krings, Huber & Kircher | 2008 | fMRI | NO | EXPLICIT | VERBAL | VISUAL | HIGH | 14 | biased thematic judgement>letters |

| | | | | | | | | | |
|---|---|---|---|---|---|---|---|---|---|
| | | | | | | | | | biased taxonomic judgement>letters<br>balanced taxonomic judgement>letters<br>balanced taxonomic judgement>letters |
| Saur et al. | 2008 | fMRI | NO | IMPLICIT | VERBAL | AUDITORY | HIGH | 33 | word>pseudoword sentences |
| Schell, Zaccarella & Friederici | 2017 | fMRI | NO | EXPLICIT | VERBAL | AUDITORY | HIGH | 21 | phrases>non combinatorial words |
| Schmitt, Auer & Ferstl | 2019 | fMRI | NO | EXPLICIT | VERBAL | AUDITORY | HIGH | 40 | known>unknown language |
| Schuil, Smits & Zwaan | 2013 | fMRI | NO | EXPLICIT | VERBAL | VISUAL | HIGH | 20 | senteces>pseudowords<br>verbs>pseudowords<br>literal sentences>pseudowords<br>nonliteral sentences>pseudowords |
| Scott, Blank, Rosen & Wise | 2000 | PET | NO | IMPLICIT | VERBAL | AUDITORY | HIGH | 8 | intelligible>unintelligible speech |
| Segal & Petrides | 2012 | fMRI | NO | EXPLICIT | NONVERBAL | VISUAL | HIGH | 90 | writing>copying<br>words>pseudowords |
| Seghier, Josse, Leff & Price | 2011 | fMRI | NO | EXPLICIT | BOTH | VISUAL | HIGH | 60 | meaningful>meaningless stimuli |
| Sergent, Otha & Macdonald | 1992 | PET | YES | EXPLICIT | VERBAL | VISUAL | HIGH | 7 | face identity>gender discrimination |
| Sheldon, McAndrews, Pruessner & Moscovitch | 2016 | fMRI | NO | EXPLICIT | VERBAL | VISUAL | HIGH | 15 | semantic fluency>perceptual task |
| Simard, Monetta, Nagano-Saito & Monchi | 2013 | fMRI | NO | EXPLICIT | VERBAL | VISUAL | HIGH | 14 | semantic>control macthing<br>semantic matching>phonological decision (syllable rhyme)<br>semantic matching>phonological decision (syllable onset) matching |
| Slioussar, Kireev, Chernigovskaya, Kataeva, Korotkov & Medvedev | 2014 | fMRI | NO | EXPLICIT | VERBAL | VISUAL | HIGH | 21 | real verbs>pseudowords<br>real nouns>pseudowords |
| Smith, Myers, Sethi, Pantazatos, Yanagihara & Hirsch | 2012 | fMRI | NO | EXPLICIT | VERBAL | VISUAL | HIGH | 14 | semantics>baseline |
| Snijders, Vosse, Kempen, Van Berkum, Petersson & Hagoort | 2009 | fMRI | NO | IMPLICIT | VERBAL | VISUAL | HIGH | 28 | sentences>word lists |
| Stowe, Paans, Wijers, Zwarts, Mulder & Vaalburg | 1999 | PET | NO | IMPLICIT | VERBAL | VISUAL | HIGH | 12 | sentences>scrambled word lists |
| Straube, Green, Weis & Kircher | 2012 | fMRI | NO | IMPLICIT | VERBAL | AUDITORY | HIGH | 16 | known>unknown language<br>iconic>meaningless gesture |

| | | | | | | | | | |
|---|---|---|---|---|---|---|---|---|---|
| Stringaris, Medford, Giampietro. Brammer & David | 2007 | fMRI | NO | EXPLICIT | VERBAL | VISUAL | HIGH | 11 | literal>meaningless sentences |
| | | | | | | | | | methaphors>meaningless sentences |
| Sugiura et al. | 2006 | fMRI | YES | EXPLICIT | VERBAL | VISUAL | HIGH | 24 | famous>unfamiliar names |
| | | | | | | | | | personal>unfamiliar names |
| Sugiura et al. | 2001 | PET | YES | EXPLICIT | NONVERBAL | VISUAL | HIGH | 5 | identity discrimination>control |
| | | | | | | | | | identity discrimination>face direction |
| Sugiura et al. | 2008 | fMRI | YES | EXPLICIT | VERBAL | VISUAL | HIGH | 25 | low familiar>unfamiliar names |
| | | | | | | | | | personal familiar>unfamiliar names |
| | | | | | | | | | high familiar>unfamiliar names |
| | | | | | | | | | high familiar>unfamiliar names |
| Sun, Xue, Zhang, Zuo, Chen, Wang, Martin, Wang, Chen, He & Wang | 2017 | fMRI | NO | EXPLICIT | VERBAL | VISUAL | HIGH | 11 | semantic>orthography judgement |
| Takeichi, Koyama, Terao, Takeuchi, Toyosawa & Murohashi | 2010 | fMRI | NO | IMPLICIT | VERBAL | AUDITORY | HIGH | 23 | speech>reversed speech |
| | | | | | | | | | speech>modulated speech |
| Taylor, Arsalidou, Bayless, Morris, Evans & Barbeau | 2009 | fMRI | YES | IMPLICIT | NONVERBAL | VISUAL | HIGH | 10 | own>unfamiliar face |
| | | | | | | | | | partner's>unfamiliar face |
| | | | | | | | | | parent's>unfamiliar face |
| Thierry & Price | 2006 | PET | NO | EXPLICIT | VERBAL | AUDITORY | HIGH | 12 | auditory words>speech control(scrambled) |
| | | | | | | | | | auditory sounds>sound control (scrambled) |
| Thierry & Price | 2006 | PET | NO | EXPLICIT | VERBAL | VISUAL | HIGH | 12 | visual words>text control (scrambled letter strings) |
| | | | | | | | | | visual videos>video control (distorted) |
| Tieleman, Seurinck, Deblaere, Vandemaele, Vingerhoets & Achten | 2005 | fMRI | NO | EXPLICIT | VERBAL | VISUAL | HIGH | 22 | self-paced semantic>perceptual decision |
| | | | | | | | | | fixed-paced semantic>perceptual decision |
| Tyler, Stamatakis, Dick, Bright, Fletcher & Moss | 2003 | fMRI | NO | EXPLICIT | VERBAL | VISUAL | HIGH | 12 | animals>baseline |
| | | | | | | | | | tool action words>baseline |
| | | | | | | | | | biological action>baseline |
| Vagharchakian, Dehaene-Lambertz, Pallier & Dehaene | 2012 | fMRI | NO | EXPLICIT | VERBAL | BOTH | HIGH | 16 | intelligible>unintelligble compression rate |

| | | | | | | | | | |
|---|---|---|---|---|---|---|---|---|---|
| | | | | | | | | | intelligible>unintelligble compression rate |
| | | | | | | | | | intelligible>unintelligble compression rate |
| Van Ettinger-Veenstra, McAllister, Lundberg, Karlsson & Engstrom | 2016 | fMRI | NO | EXPLICIT | VERBAL | VISUAL | HIGH | 27 | sentences>symbol strings |
| van Leeuwen et al. | 2014 | fMRI | NO | EXPLICIT | BOTH | BOTH | HIGH | 16 | speech>reversed speech |
| Vignali, Hawelka, Hutzler & Richlan | 2019 | fMRI | NO | EXPLICIT | VERBAL | VISUAL | HIGH | 21 | foveal & parafovealwords>foveal & parafoveal pseudowords |
| Vingerhoets | 2008 | fMRI | NO | IMPLICIT | NONVERBAL | VISUAL | HIGH | 14 | familiar>unfamiliar tools |
| Visser, Jefferies, Embleton & Lambon Ralph | 2012 | fMRI | NO | EXPLICIT | BOTH | VISUAL | HIGH | 15 | semantics>baseline |
| | | | | | | | | | pictures>baseline |
| | | | | | | | | | words>baseline |
| von Kriegstein, Eger, Kleinschmidt & Giraud | 2003 | fMRI | NO | EXPLICIT | VERBAL | AUDITORY | HIGH | 14 | sentence>speech envelope |
| Wang, Zhao, Zevin & Yang | 2016 | fMRI | NO | EXPLICIT | VERBAL | VISUAL | HIGH | 16 | words>nonsense strokes |
| Welcome & Joanisse | 2012 | fMRI | NO | EXPLICIT | VERBAL | VISUAL | HIGH | 20 | semantic>phonological/orthographic decision |
| Wende, Straube, Stratmann, Sommer, Kircher & Nagels | 2012 | fMRI | NO | EXPLICIT | VERBAL | VISUAL | HIGH | 18 | semantic>phonological fluency |
| Wirth, Jann, Dierks, Federspiel, Wiest & Horn | 2011 | fMRI | NO | EXPLICIT | VERBAL | VISUAL | HIGH | 19 | semantic>phonological & perceptual decision |
| Wright et al. | 2008 | fMRI | NO | EXPLICIT | BOTH | VISUAL | HIGH | 10 | semantic>perceptual matching |
| | | | | | | | | | semantic>perceptual matching |
| | | | | | | | | | semantic>perceptual matching |
| Xiao et al. | 2005 | fMRI | NO | EXPLICIT | VERBAL | AUDITORY | HIGH | 14 | words>pseudowords |
| Zaccarella & Friederici | 2015 | fMRI | NO | EXPLICIT | VERBAL | VISUAL | HIGH | 22 | words>pseudowords |
| Zhang, Xiao & Weng | 2012 | fMRI | NO | EXPLICIT | VERBAL | VISUAL | HIGH | 14 | words>pseudowords |
| Zhuang & Devereux | 2017 | fMRI | NO | EXPLICIT | VERBAL | AUDITORY | HIGH | 16 | phrases>words |
| Zvyagintsev, Clemens, Chechko, Mathiak, Sack & Mathiak | 2013 | fMRI | NO | IMPLICIT | NONE | NONE | HIGH | 15 | visual imagery>counting |
| | | | | | | | | | auditory imagery>counting |
| EXCLUDED CONTRASTS WITH REST AND LOW LEVEL BASELINES | | | | | | | | | |
| AbdulSabur et al. | 2014 | fMRI | NO | EXPLICIT | VERBAL | VISUAL | LOW | 18 | narrative production>recitation |
| Alain, He & Grady | 2008 | fMRI | NO | EXPLICIT | VERBAL | AUDITORY | REST | 16 | sound category>rest |
| Assadollahi, Meinzer, Flaisch, Obleser & Rockstroh | 2009 | fMRI | NO | IMPLICIT | VERBAL | VISUAL | REST | 20 | nouns followed by 1 argument verbs>fixation cross (rest) |

| | | | | | | | | | |
|---|---|---|---|---|---|---|---|---|---|
| | | | | | | | | | nouns followed by 3 argument verbs>fixation cross (rest) |
| Barros-Loscertales et al. | 2012 | fMRI | NO | EXPLICIT | VERBAL | VISUAL | LOW | 59 | control words>hashmarks baseline |
| Bick, Goelman & Frost | 2008 | fMRI | NO | EXPLICIT | VERBAL | VISUAL | LOW | 14 | semantic>visual control |
| | | | | | | | | | morphologiacl>visual control |
| | | | | | | | | | orthographic>visual control |
| | | | | | | | | | phonological>visual control |
| Birn et al. | 2010 | fMRI | NO | EXPLICIT | VERBAL | VISUAL | LOW | 14 | category & letter fluency>months ( automatic speech) |
| Booth et al. | 2006 | fMRI | NO | EXPLICIT | VERBAL | VISUAL | LOW | 13 | meaning(semantic)>control(symbols) |
| Boulenger, Hauk & Pulvermuller | 2009 | fMRI | NO | EXPLICIT | VERBAL | VISUAL | LOW | 18 | sentences> hashmarks |
| | | | | | | | | | sentences> hashmarks |
| Cai, Kochiyama, Osaka & Wu | 2007 | fMRI | NO | IMPLICIT | VERBAL | AUDITORY | REST | 15 | words>rest |
| Cao, Peng, Liu, Jin, Fan, Deng, & Booth | 2009 | fMRI | NO | EXPLICIT | VERBAL | VISUAL | LOW | 13 | meaning>perceptual decision |
| Carota, Kriegeskorte, Nili & Pulvermuller | 2017 | fMRI | NO | IMPLICIT | VERBAL | VISUAL | LOW | 23 | words>hashmarks |
| Carota, Moseley & Pulvermueller | 2012 | fMRI | NO | IMPLICIT | VERBAL | VISUAL | LOW | 18 | all words>hashmarks |
| | | | | | | | | | tool words>hashmarks |
| | | | | | | | | | animal words>hashmarks |
| | | | | | | | | | food words>hashmarks |
| Chan, Tang, Tang, Lee, Lo & Kwong | 2009 | fMRI | NO | IMPLICIT | VERBAL | VISUAL | REST | 22 | synonyms>rest |
| Chiao, Harada, Oby, Li, Parrish & Bridge | 2009 | fMRI | YES | EXPLICIT | NONVERBAL | VISUAL | LOW | 12 | uniform status judgement>colour change detection |
| | | | | | | | | | face status judgement>colour change detection |
| | | | | | | | | | car status judgement>colour change detection |
| Damasio, Grabowski, Tranel, Ponto, Hichwa & Damasio | 2001 | PET | NO | EXPLICIT | NONVERBAL | VISUAL | LOW | 10 | actions without implement>control task |
| | | | | | | | | | actions with implement>control task |
| Dapretto & Bookheimer | 1999 | fMRI | NO | EXPLICIT | VERBAL | AUDITORY | REST | 8 | semantic>rest |
| D'Arcy, Bolster, Ryner, Mazerolle, Grant & Song | 2007 | fMRI | NO | EXPLICIT | BOTH | VISUAL | REST | 10 | basic-level living objects>baseline |
| | | | | | | | | | basic-level non-living objects>baseline |

| | | | | | | | | | |
|---|---|---|---|---|---|---|---|---|---|
| | | | | | | | | | superordinate-level living objects>baseline<br>superordinate-level non-living objects>baseline |
| Davis, Ford, Kherif & Johnsrude | 2011 | fMRI | NO | EXPLICIT | VERBAL | AUDITORY | LOW | 12 | clear & anomalous sentences>signal correlated noise |
| Demonet et al. | 1992 | PET | NO | EXPLICIT | VERBAL | AUDITORY | LOW | 9 | words>tones |
| Dreyer & Pulvermueller | 2018 | fMRI | YES | IMPLICIT | VERBAL | VISUAL | LOW | 28 | all nouns>hashmarks<br>abstract emotional nouns>baseline (hashmarks)<br>abstract mental nouns>baseline (hashmarks)<br>food nouns>baseline (hashmarks)<br>tool nouns>baseline (hashmarks) |
| Emmorey, Xu, Gannon, Goldin-Meadow & Braun | 2010 | fMRI | NO | IMPLICIT | NONVERBAL | VISUAL | REST | 14 | meaningful pantomimes>rest |
| Engelien et al. | 2006 | PET | NO | IMPLICIT | NONVERBAL | AUDITORY | REST | 6 | meaningful>rest |
| Foki, Gartus, Geissler & Beisteiner | 2008 | fMRI | | EXPLICIT | VERBAL | VISUAL | LOW | 23 | semantic judgement>tongue movements |
| Groussard et al. | 2010 | PET | NO | EXPLICIT | VERBAL | VISUAL | REST | 11 | verbal semantic>rest<br>verbal semantic>rest |
| Guediche, Reilly, Santiago, Laurent & Blumstein | 2016 | fMRI | NO | IMPLICIT | VERBAL | AUDITORY | LOW | 16 | related > repeated sentence<br>unrelated > repeated sentence |
| Hagoort et al. | 1999 | PET | NO | IMPLICIT | VERBAL | VISUAL | REST | 10 | words>fixation |
| Harrington, Farias & Davis | 2009 | fMRI | NO | EXPLICIT | NONVERBAL | VISUAL | REST | 8 | familiar objects>rest |
| Hauk & Pulvermueller | 2011 | fMRI | NO | IMPLICIT | VERBAL | VISUAL | LOW | 21 | action words>hashmarks<br>uni manual action words>hashmarks<br>uni manual action words>hashmarks |
| Hayashi et al. | 2014 | fMRI | NO | EXPLICIT | VERBAL | VISUAL | LOW | 16 | concrete word>astersisks<br>abstract word>asterisks |
| Heim, Eickhoff & Amunts | 2008 | fMRI | NO | EXPLICIT | VERBAL | VISUAL | REST | 28 | semantic fluency>rest |
| Higuchi, Moriguchi, Murakami, Katsunuma, Mishima &Uno | 2015 | fMRI | NO | IMPLICIT | VERBAL | VISUAL | LOW | 28 | all characters>checkerboard |
| Hwang, Palmer, Basho, Zadra & Muller | 2009 | fMRI | NO | EXPLICIT | VERBAL | AUDITORY | LOW | 13 | fluency generation>production baseline |
| Ikuta et al. | 2006 | fMRI | NO | IMPLICIT | VERBAL | VISUAL | LOW | 34 | sentences>word lists |
| Jensen, Hargreaves, Bass, Pexman, Goodyear & Federico | 2011 | fMRI | NO | EXPLICIT | VERBAL | VISUAL | REST | 12 | words>rest |

| | | | | | | | | | |
|---|---|---|---|---|---|---|---|---|---|
| Jeon, Lee, Kim & Cho | 2009 | fMRI | NO | EXPLICIT | VERBAL | VISUAL | REST | 16 | synonyms>rest |
| | | | | | | | | | antonyms>rest |
| | | | | | | | | | English synonyms>rest |
| | | | | | | | | | Korean synonyms>nonwords |
| | | | | | | | | | honorific words>nonwords |
| Kang et al. | 2006 | PET | NO | EXPLICIT | VERBAL | AUDITORY | LOW | 17 | auditory speech>white noise |
| Kim et al. | 2009 | fMRI | NO | EXPLICIT | VERBAL | VISUAL | REST | 36 | sentences>rest |
| Kyong, Scott, Rosen, Howe, Agnew & McGettigan | 2014 | fMRI | NO | IMPLICIT | VERBAL | AUDITORY | LOW | 19 | intelligible vocoded>inverted vocoded speech |
| Liu et al. | 2009 | fMRI | NO | EXPLICIT | VERBAL | VISUAL | LOW | 16 | words meaning>slashes |
| | | | | | | | | | words rhyming>slashes |
| | | | | | | | | | words meaning>tones |
| | | | | | | | | | words rhyming>tones |
| Ludersdorfer, Wimmer, Richlan, Schurz, Hutzler & Kronbichler | 2016 | fMRI | NO | EXPLICIT | VERBAL | AUDITORY | LOW | 29 | words orthographic>tones |
| | | | | | | | | | words semantic>tones |
| | | | | | | | | | Auditory words orthographic>rest |
| | | | | | | | | | Auditory words semantic>rest |
| Malins, Gumkowski, Buis, Molfese, Rueckl, Frost, Pugh, Morris & Mencl | 2016 | fMRI | NO | IMPLICIT | VERBAL | VISUAL | LOW | 18 | unrelated words>false font |
| Marques, Canessa & Cappa | 2009 | fMRI | NO | EXPLICIT | VERBAL | VISUAL | LOW | 21 | sentences>crosses |
| Marques, Canessa, Siri, Catricala & Cappa | 2008 | fMRI | NO | EXPLICIT | VERBAL | VISUAL | LOW | 21 | semantic features>baseline task |
| Matchin, Liao, gaston & Lau | 2019 | fMRI | NO | EXPLICIT | VERBAL | VISUAL | REST | 20 | sentence>rest |
| Mellem, Jasmin, Peng & Martin | 2016 | fMRI | NO | IMPLICIT | VERBAL | VISUAL | LOW | 20 | longer>shorter phrase |
| Moseley, Carota, Hauk, Mohr & Pulvermueller | 2012 | fMRI | YES | IMPLICIT | VERBAL | VISUAL | LOW | 18 | all emotional words>hashmarks |
| | | | | | | | | | abstract emotional words>hashmarks |
| | | | | | | | | | arm+face+emotion words>hashmarks |
| | | | | | | | | | face words>hashmarks |
| | | | | | | | | | arm words>hashmarks |
| Nakamura et al. | 2000 | PET | YES | EXPLICIT | NONVERBAL | VISUAL | LOW | 7 | familiar faces>fixation cross |
| Noppeney & Price | 2003 | PET | NO | EXPLICIT | VERBAL | AUDITORY | LOW | 9 | normal>reversed words |
| Perrone-Bertolotti, Kauffmann, Pichat, Vidal & Baciu | 2017 | fMRI | NO | EXPLICIT | VERBAL | VISUAL | REST | 24 | words>fixation |

| | | | | | | | | | |
|---|---|---|---|---|---|---|---|---|---|
| Pulvermueller, Cook & Hauk | 2012 | fMRI | NO | IMPLICIT | VERBAL | VISUAL | LOW | 23 | phrases>hashmarks |
| | | | | | | | | | uninflected words>hashmarks<br>inflected words>hashmarks |
| Raposo, Moss, Stamatakis & Tyler | 2009 | fMRI | NO | IMPLICIT | VERBAL | AUDITORY | LOW | 22 | action sentences>SC noise |
| Rapp & Lipka | 2011 | fMRI | NO | IMPLICIT | VERBAL | VISUAL | LOW | 10 | words>checkerboards |
| Rodd, Johnsrude & Davis | 2012 | fMRI | NO | EXPLICIT | VERBAL | AUDITORY | LOW | 15 | speech>SCN |
| Rodd, Longe, Randall & Tyler | 2010 | fMRI | NO | EXPLICIT | VERBAL | AUDITORY | LOW | 14 | speech>SCN |
| Sergent, Otha & Macdonald | 1992 | PET | YES | EXPLICIT | NONVERBAL | VISUAL | LOW | 7 | object recognition>gratings |
| Stowe, Paans, Wijers, Zwarts, Mulder & Vaalburg | 1999 | PET | NO | IMPLICIT | VERBAL | VISUAL | REST | 12 | sentences>rest |
| Szlachta, Bozic, Jelowicka & Marslen-Wilson | 2012 | fMRI | NO | IMPLICIT | VERBAL | VISUAL | LOW | 21 | words>musical rain |
| | | | | | | | | | nouns>musical rain<br>inflected nouns>musical rain |
| Taminato, Miura, Sugiura & Kawashima | 2014 | fMRI | NO | EXPLICIT | NONVERBAL | VISUAL | LOW | 35 | object recognition>control task |
| | | | | | | | | | object recognition>control task |
| Taylor, Arsalidou, Bayless, Morris, Evans & Barbeau | 2009 | fMRI | YES | IMPLICIT | NONVERBAL | VISUAL | REST | 10 | own face>baseline (rest) |
| | | | | | | | | | parents face>baseline (rest)<br>famous face>baseline (rest) |
| Vitello, Warren, Devlin & Rodd | 2014 | fMRI | NO | EXPLICIT | VERBAL | AUDITORY | LOW | 20 | sentences>SCN |
| Weiss, Katzir & Bitan | 2015 | fMRI | NO | IMPLICIT | VERBAL | VISUAL | LOW | 18 | pointed words>asterisks<br>unpointed words>asterisks |
| Wende, Straube, Stratmann, Sommer, Kircher & Nagels | 2012 | fMRI | NO | EXPLICIT | VERBAL | VISUAL | REST | 18 | causal fluency>rest |
| Wright et al. | 2008 | fMRI | NO | EXPLICIT | VERBAL | VISUAL | REST | 15 | free association>rest<br>words>rest<br>pictures>rest |
| Wright, Randall, Marslen-Wilson & Tyler | 2011 | fMRI | NO | BOTH | VERBAL | AUDITORY | LOW | 14 | speech>musical rain |
| Wu, Mai, Tang, Ge, Luo & Liu | 2013 | fMRI | NO | IMPLICIT | VERBAL | VISUAL | LOW | 19 | arm words>checkerboard |
| | | | | | | | | | leg words>checkerboard<br>mouth words>checkerboard |

| | | | | | | | | | |
|---|---|---|---|---|---|---|---|---|---|
| Xiao et al. | 2005 | fMRI | NO | EXPLICIT | VERBAL | AUDITORY | REST | 14 | words>rest |
| Yang, Li, Fang, Shu, Liu & Chen | 2016 | fMRI | NO | IMPLICIT | VERBAL | VISUAL | LOW | 20 | opaque idioms>hashmarks |
| | | | | | | | | | transparent idioms>hashmarks |
| | | | | | | | | | literal phrases>hashmarks |
| Zhang, Liu & Zhang | 2014 | fMRI | NO | EXPLICIT | VERBAL | VISUAL | LOW | 18 | nonliving words>asterisks |
| | | | | | | | | | living words>asterisks |
| Zhang, Xiao & Weng | 2012 | fMRI | NO | EXPLICIT | VERBAL | VISUAL | REST | 14 | words>rest |
| Zou, Packard, Xia, Liu & Shu | 2016 | fMRI | NO | IMPLICIT | VERBAL | AUDITORY | LOW | 17 | speech>tone |
| | | | | | | | | | speech>tone |
| | | | | | | | | | speech>tone |
| | | | | | | | | | identical speech>tone |
| | | | | | | | | | words>tone |
| Giraud & Price | 2001 | PET | NO | BOTH | BOTH | AUDITORY | BOTH | 12 | words+environmental sounds>syllables+noise |

***The references to access the listed studies are listed at the end of the document**

***Table M2.0 SC NO SOC*** *List of studies included in the* ***semantic cognition MAIN (high & low baselines) analysis after further excluding contrasts with a degree of social content*** *(N = 193) meta-analysis.* ***Note:*** *The list of excluded studies with social content in the stimuli can be seen at the end of the document highlighted in grey. SC= Semantic Cognition, N= Sample Size, SOC= stimuli with a degree of social content*

| *Authors | Year | Imaging Method | Social Content | Instructional Cue | Stimulus Domain | Sensory Input Modality | Baseline Type | N | Contrast |
|---|---|---|---|---|---|---|---|---|---|
| AbdulSabur et al. | 2014 | fMRI | NO | EXPLICIT | VERBAL | VISUAL | LOW | 18 | narrative production>recitation |
| Abraham et al. | 2012 | fMRI | NO | EXPLICIT | VERBAL | VISUAL | HIGH | 19 | high&low divergent thinking (semantic)>1&2 back letter identity (working memory) |
| Alain, He & Grady | 2008 | fMRI | NO | EXPLICIT | VERBAL | AUDITORY | HIGH | 16 | sound category>sound location |
| Assadollahi, Meinzer, Flaisch, Obleser & Rockstroh | 2009 | fMRI | NO | IMPLICIT | VERBAL | VISUAL | HIGH | 20 | nouns followed by 1&3 argument verbs>letter strings followed by 1&3argument verbs |
| Axmacher, Bialleck, Weber, Helmstaedter, Elger & Fell | 2009 | fMRI | NO | EXPLICIT | VERBAL | VISUAL | HIGH | 32 | word decision>spatial decision |
| Bagga et al. | 2013 | fMRI | NO | EXPLICIT | VERBAL | VISUAL | HIGH | 18 | semantic>case matching judgement |
| Barros-Loscertales et al. | 2012 | fMRI | NO | EXPLICIT | VERBAL | VISUAL | LOW | 59 | control words>hashmarks baseline |
| Baumgaertener, Weiller & Buchel | 2002 | fMRI | NO | EXPLICIT | VERBAL | VISUAL | HIGH | 9 | word>pseudoword in sentence |
| Baumgaertner et al. | 2007 | fMRI | NO | IMPLICIT | VERBAL | AUDITORY | HIGH | 19 | sentences>reversed sentences |
| | | | | | | | | | videos>scrambled videos |
| Bautista & Wilson | 2016 | fMRI | NO | IMPLICIT | VERBAL | AUDITORY | HIGH | 12 | clear>scrambled rotated speech |
| Bick, Goelman & Frost | 2008 | fMRI | NO | EXPLICIT | VERBAL | VISUAL | LOW | 14 | semantic>visual control |
| | | | | | | | | | morphologiacl>visual control |
| | | | | | | | | | orthographic>visual control |
| | | | | | | | | | phonological>visual control |
| Binder, Frost, Hammeke, Bellgowan, Rao & Cox | 1999 | fMRI | NO | EXPLICIT | VERBAL | AUDITORY | HIGH | 30 | semantic>phonological decision |
| Binder et al. | 2003 | fMRI | NO | EXPLICIT | VERBAL | VISUAL | HIGH | 24 | word>nonword |
| Birn et al. | 2010 | fMRI | NO | EXPLICIT | VERBAL | VISUAL | LOW | 14 | category & letter fluency>months ( automatic speech) |
| | | | | | | | | | category>letter fluency |
| Bonhage, Fiebach, Bahlmann & Mueller | 2014 | fMRI | NO | EXPLICIT | VERBAL | VISUAL | HIGH | 18 | sentence fragments>ungrammatical word strings |

| | | | | | | | | | |
|---|---|---|---|---|---|---|---|---|---|
| Bonhage, Mueller, Friederici & Fiebach | 2015 | fMRI | NO | EXPLICIT | VERBAL | VISUAL | HIGH | 18 | sentences>jabberwocky sentences |
| Booth et al. | 2006 | fMRI | NO | EXPLICIT | VERBAL | VISUAL | HIGH | 13 | meaning (semantic) judgement>rhyming( phonological) judgement |
| | | | | | | | | | meaning(semantic)>control(symbols) |
| Boulenger, Hauk & Pulvermuller | 2009 | fMRI | NO | EXPLICIT | VERBAL | VISUAL | LOW | 18 | sentences> hashmarks |
| | | | | | | | | | sentences> hashmarks |
| Bozic & Marslen-Wilson | 2013 | fMRI | NO | IMPLICIT | VERBAL | AUDITORY | HIGH | 13 | speech>musical rain |
| Bruffaerts, Dupont, Peeters, De Deyne, Storms & Vandenberghe | 2013 | fMRI | NO | EXPLICIT | BOTH | VISUAL | HIGH | 19 | real > scrambled pictures & words |
| Bulut, Hung, Tzeng & Wu | 2017 | fMRI | NO | IMPLICIT | VERBAL | VISUAL | HIGH | 20 | sentences>unstructured word lists |
| | | | | | | | | | sentences>unstructured character list |
| Cai, Kochiyama, Osaka & Wu | 2007 | fMRI | NO | IMPLICIT | VERBAL | AUDITORY | HIGH | 15 | words>nonsense words |
| Cao, Peng, Liu, Jin, Fan, Deng, & Booth | 2009 | fMRI | NO | EXPLICIT | VERBAL | VISUAL | LOW | 13 | meaning>perceptual decision |
| Cappa, Perani, Schnur, Tettamanti & Fazio | 1998 | PET | NO | EXPLICIT | VERBAL | VISUAL | HIGH | 13 | words>pseudowords |
| | | | | | | | | | animal visual knowledge>pseudowords |
| | | | | | | | | | tool visual knowledge>pseudowords |
| | | | | | | | | | animal associative knowledge>pseudowords |
| | | | | | | | | | tools functional knowledge>pseudowords |
| Carota, Kriegeskorte, Nili & Pulvermuller | 2017 | fMRI | NO | IMPLICIT | VERBAL | VISUAL | LOW | 23 | words>hashmarks |
| Carota, Moseley & Pulvermueller | 2012 | fMRI | NO | IMPLICIT | VERBAL | VISUAL | LOW | 18 | all words>hashmarks |
| | | | | | | | | | tool words>hashmarks |
| | | | | | | | | | animal words>hashmarks |
| | | | | | | | | | food words>hashmarks |
| Chan, Tang, Tang, Lee, Lo & Kwong | 2009 | fMRI | NO | IMPLICIT | VERBAL | VISUAL | HIGH | 22 | synonyms>pseudocharacters |
| | | | | | | | | | synonyms>Korean characters (unknown) |
| Chou, Chen, Wu & Booth | 2009 | fMRI | NO | EXPLICIT | VERBAL | VISUAL | HIGH | 31 | related words>false font |

| | | | | | | | | | |
|---|---|---|---|---|---|---|---|---|---|
| | | | | | | | | | unrelated words>false font |
| Chou, Chen, Wu & Booth | 2009 | fMRI | NO | EXPLICIT | VERBAL | VISUAL | HIGH | 32 | related words>false font |
| | | | | | | | | | unrelated words>false font |
| Chouinard, Morrissey, Kohler & Goodale | 2008 | fMRI | NO | EXPLICIT | NONVERBAL | VISUAL | HIGH | 14 | objects>scrambled objects |
| Chow, Kaup, Raabe & Greenlee | 2008 | fMRI | NO | EXPLICIT | VERBAL | VISUAL | HIGH | 15 | predictive&normal reading>pseudoword reading |
| | | | | | | | | | normal reading>pseudoword reading |
| | | | | | | | | | predictive reading>pseudoword reading |
| Christensen, Antonucci, Lockwood, Kittleson & Plante | 2008 | fMRI | NO | EXPLICIT | VERBAL | AUDITORY | HIGH | 14 | diotic listening>reversed speech |
| | | | | | | | | | dichotic listening>reversed speech |
| Clos, Langner, Meyer, Oechslin, Zilles & Eickhoff | 2014 | fMRI | NO | EXPLICIT | VERBAL | AUDITORY | HIGH | 29 | intelligibilty based on cue > unintelligible |
| Damasio, Grabowski, Tranel, Ponto, Hichwa & Damasio | 2001 | PET | NO | EXPLICIT | NONVERBAL | VISUAL | LOW | 10 | actions without implement>control task |
| | | | | | | | | | actions with implement>control task |
| Damasio, Tranel, Grabowski, Adolphs & Damasio | | PET | NO | EXPLICIT | NONVERBAL | VISUAL | HIGH | 55 | animals>scrambled pictures |
| | | | | | | | | | tools>scrambled pictures |
| Davis, Meunier & Marslen-Wilson | 2004 | fMRI | NO | EXPLICIT | VERBAL | VISUAL | HIGH | 11 | words>letter strings |
| Davis, Ford, Kherif & Johnsrude | 2011 | fMRI | NO | EXPLICIT | VERBAL | AUDITORY | LOW | 12 | clear & anomalous sentences>signal correlated noise |
| Demonet et al. | 1992 | PET | NO | EXPLICIT | VERBAL | AUDITORY | HIGH | 9 | words>phonemes |
| | | | | | | | | | words>tones |
| Devlin, Matthews & Rushworth | 2003 | fMRI | NO | EXPLICIT | VERBAL | VISUAL | HIGH | 12 | semantic>phonological judgement |
| Devlin et al. | 2002 | PET | NO | EXPLICIT | VERBAL | VISUAL | HIGH | 12 | all semantic>letter detection |
| | | | | | | | | | all semantic>letter detection |
| Devlin et al. | 2002 | PET | NO | EXPLICIT | VERBAL | VISUAL | HIGH | 8 | all semantic>letter categorisation |
| Devlin et al. | 2002 | fMRI | NO | EXPLICIT | VERBAL | VISUAL | HIGH | 8 | all semantic> letter categorisation |

| | | | | | | | | | |
|---|---|---|---|---|---|---|---|---|---|
| Devlin et al. | 2000 | PET | NO | EXPLICIT | VERBAL | VISUAL | HIGH | 8 | semantic categorisation>letter categorisation |
| Devlin et al. | 2000 | fMRI | NO | EXPLICIT | VERBAL | VISUAL | HIGH | 8 | semantic categorisation>letter categorisation |
| Diaz & McCarthy | 2009 | fMRI | NO | IMPLICIT | VERBAL | VISUAL | HIGH | 16 | all words>nonwords |
| Ebisch et al. | 2007 | fMRI | NO | EXPLICIT | BOTH | VISUAL | HIGH | 17 | functional&visuospatial pictures&words>meaningless drawings&pseudowords |
| Emmorey, Weisberg, McCullough & Petrich | 2013 | fMRI | NO | EXPLICIT | VERBAL | VISUAL | HIGH | 14 | words(semantic judgement)>false fonts |
| | | | | | | | | | words(phonological)>false fonts |
| Emmorey, Xu, Gannon, Goldin-Meadow & Braun | 2010 | fMRI | NO | IMPLICIT | NONVERBAL | VISUAL | HIGH | 14 | meaningful pantomimes>unknown sign language |
| Engelien et al. | 2006 | PET | NO | IMPLICIT | NONVERBAL | AUDITORY | HIGH | 6 | meaningful>meaningless sounds |
| Erb, Henry, Eisner & Obleser | 2013 | fMRI | NO | EXPLICIT | VERBAL | AUDITORY | HIGH | 30 | speech>vocoded speech |
| Europa, Gitelman, Kiran & Thompson | 2019 | fMRI | NO | EXPLICIT | BOTH | BOTH | HIGH | 21 | sentences>baseline (reversed sentences) |
| Foki, Gartus, Geissler & Beisteiner | 2008 | fMRI | | EXPLICIT | VERBAL | VISUAL | LOW | 23 | semantic judgement>tongue movements |
| Friederici, Kotz, Scott & Obleser | 2010 | fMRI | NO | IMPLICIT | VERBAL | AUDITORY | HIGH | 17 | intelligible speech>rotated speech |
| Friese, Rutschmann, Raabe & Schmalhofer | 2008 | fMRI | NO | EXPLICIT | VERBAL | VISUAL | HIGH | 13 | words>pseudowords |
| Garbin, Collina & Tabossi | 2012 | fMRI | NO | EXPLICIT | VERBAL | VISUAL | HIGH | 12 | object noun>pseudoword |
| | | | | | | | | | event noun>pseudoword |
| | | | | | | | | | verb>pseudoword |
| Garn, Allen & Larsen | 2009 | fMRI | NO | EXPLICIT | NONVERBAL | VISUAL | HIGH | 26 | pictures>scrambled pictures |
| | | | | | | | | | plants>scrambled pictures |
| | | | | | | | | | tools>scrambled pictures |
| Geranmayeh, Brownsett, Leech, Beckmann, Woodhead & Wise | 2012 | fMRI | NO | EXPLICIT | VERBAL | VISUAL | HIGH | 19 | speech>tongue movements |
| Gerlach, Law, Gade & Paulson | 1999 | PET | NO | EXPLICIT | NONVERBAL | VISUAL | HIGH | 15 | object decision>pattern discrimination |
| Giraud & Price | 2001 | PET | NO | BOTH | BOTH | AUDITORY | BOTH | 12 | words+environmental sounds>syllables+noise |
| Giraud et al. | 2004 | fMRI | NO | EXPLICIT | VERBAL | AUDITORY | HIGH | 8 | natural speech>speech envelope |

| | | | | | | | | | |
|---|---|---|---|---|---|---|---|---|---|
| Gitelman, Nobre, Sonty, Parrish & Mesulam | 2005 | fMRI | NO | EXPLICIT | VERBAL | VISUAL | HIGH | 14 | semantic>control task |
| Grabowski, Damasio, Tranel, Boles Ponto, Hichwa & Damasio | | PET | NO | EXPLICIT | NONVERBAL | VISUAL | HIGH | 10 | naming landmarks>face orientation judgement |
| Graves, Binder, Desai, Conant & Seidenberg | 2010 | fMRI | NO | EXPLICIT | VERBAL | VISUAL | HIGH | 23 | forward>reverse phrases |
| Graves, Binder, Desai, Conant & Seidenberg | 2010 | fMRI | NO | EXPLICIT | VERBAL | VISUAL | HIGH | 22 | forward>reverse phrases |
| Grindrod, Garnett, Malyutina & den Ouden | 2014 | fMRI | NO | EXPLICIT | VERBAL | VISUAL | HIGH | 23 | words>nonwords |
| Grossman et al. | 2002a | fMRI | NO | IMPLICIT | VERBAL | VISUAL | HIGH | 16 | all nouns > pseudowords |
| | | | | | | | | | implements>pseudowords |
| | | | | | | | | | animals>pseudowords |
| | | | | | | | | | abstract>pseudowords |
| Grossman et al., 2002b | 2002 b | fMRI | NO | IMPLICIT | VERBAL | VISUAL | HIGH | 16 | verbs>pseudowords |
| Groussard et al. | 2010 | PET | NO | EXPLICIT | VERBAL | VISUAL | HIGH | 11 | verbal semantics>verbal reference |
| | | | | | | | | | musical semantic>musical reference |
| Guediche, Reilly, Santiago, Laurent & Blumstein | 2016 | fMRI | NO | IMPLICIT | VERBAL | AUDITORY | LOW | 16 | related > repeated sentence |
| | | | | | | | | | unrelated > repeated sentence |
| Gurd et al. | 2002 | fMRI | NO | EXPLICIT | VERBAL | AUDITORY | HIGH | 11 | category>rote fluency |
| Haberling, Corballis &Corballis | 2016 | fMRI | NO | EXPLICIT | NONVERBAL | VISUAL | HIGH | 92 | meaningful pantomimes>unknown sign language |
| | | | | | | | | | meaningful pantomimes>dog videos |
| | | | | | | | | | synonyms>letter strings |
| Hagoort et al. | 1999 | PET | NO | IMPLICIT | VERBAL | VISUAL | HIGH | 10 | words>pseudowords |
| Harrington, Farias & Davis | 2009 | fMRI | NO | EXPLICIT | NONVERBAL | VISUAL | HIGH | 8 | familiar>non objects |
| Hartung, Hagoort & Willems | 2017 | fMRI | NO | IMPLICIT | VERBAL | AUDITORY | HIGH | 52 | first-person speech>unintelligible reversed speech |
| | | | | | | | | | third-person speech>unintelligible reversed speech |
| Hauk & Pulvermueller | 2011 | fMRI | NO | IMPLICIT | VERBAL | VISUAL | LOW | 21 | action words>hashmarks |
| | | | | | | | | | uni manual action words>hashmarks |

| | | | | | | | | | |
|---|---|---|---|---|---|---|---|---|---|
| | | | | | | | | | uni manual action words>hashmarks |
| Hayashi et al. | 2014 | fMRI | NO | EXPLICIT | VERBAL | VISUAL | LOW | 16 | concrete word>astersisks |
| | | | | | | | | | abstract word>asterisks |
| Heim, Eickhoff & Amunts | 2008 | fMRI | NO | EXPLICIT | VERBAL | VISUAL | HIGH | 28 | semantic > phonological fluency |
| Henke et al. | 1999 | PET | NO | EXPLICIT | VERBAL | VISUAL | HIGH | 12 | associative word learning>single word encoding |
| Herbster et al. | 1997 | PET | NO | EXPLICIT | VERBAL | VISUAL | HIGH | 10 | Irregular>zero order speak |
| | | | | | | | | | regular>zero order speak |
| | | | | | | | | | irregular+regular>zero order speak |
| Hervais-Adelman, Carlyon, Johnsrude & Davis | 2012 | fMRI | NO | EXPLICIT | VERBAL | AUDITORY | HIGH | 15 | clear>vocoded speech (uninteligible) |
| Higuchi, Moriguchi, Murakami, Katsunuma, Mishima &Uno | 2015 | fMRI | NO | IMPLICIT | VERBAL | VISUAL | LOW | 28 | all characters>checkerboard |
| Hocking, McMahon & de Zubicaray | 2011 | fMRI | NO | EXPLICIT | NONVERBAL | AUDITORY | HIGH | 13 | all environmental sounds>perceptual baseline |
| Holle, Gunter, Rueschemeyer, Hennenlotter & Iacoboni | 2008 | fMRI | NO | EXPLICIT | BOTH | BOTH | HIGH | 17 | iconic gesture of dominant meaning>grooming |
| | | | | | | | | | iconic gesture of subordinate meaning>grooming |
| Homae, Yahata & Sakai | 2003 | fMRI | NO | EXPLICIT | VERBAL | AUDITORY | HIGH | 10 | auditory sentences>auditory non words |
| | | | | | | | | | visual sentences>visual non words words |
| Husain, Patkin, Kim, Braun & Horwitz | 2012 | fMRI | NO | EXPLICIT | NONVERBAL | VISUAL | HIGH | 16 | meaningful iconic>meaningless gestures |
| Hwang, Palmer, Basho, Zadra & Muller | 2009 | fMRI | NO | EXPLICIT | VERBAL | AUDITORY | LOW | 13 | fluency generation>production baseline |
| Ikuta et al. | 2006 | fMRI | NO | IMPLICIT | VERBAL | VISUAL | LOW | 34 | sentences>word lists |
| Jackson, Hoffman, Pobric, Lambon Ralph | 2015 | fMRI | NO | EXPLICIT | VERBAL | VISUAL | HIGH | 24 | words>letter strings |
| Jensen, Hargreaves, Bass, Pexman, Goodyear & Federico | 2011 | fMRI | NO | EXPLICIT | VERBAL | VISUAL | HIGH | 12 | words>pseudowords |
| Jeon, Lee, Kim & Cho | 2009 | fMRI | NO | EXPLICIT | VERBAL | VISUAL | HIGH | 16 | synonys>nonwords |
| | | | | | | | | | antonyms>nonwords |
| | | | | | | | | | English synonyma>nonwords |
| | | | | | | | | | Korean synonyms>nonwords |

| | | | | | | | | | |
|---|---|---|---|---|---|---|---|---|---|
| | | | | | | | | | honorific words>nonwords |
| Joubert et al. | 2004 | fMRI | NO | IMPLICIT | VERBAL | VISUAL | HIGH | 10 | low frequency words>nonwords |
| | | | | | | | | | high frequency words>consonant strings |
| | | | | | | | | | low frequency words>consonant strings |
| Kang et al. | 2006 | PET | NO | EXPLICIT | VERBAL | BOTH | HIGH | 17 | audio-visual speech>noises and facial movements |
| | | | | | | | | | auditory speech>white noise |
| | | | | | | | | | visual speech>facial movements (chewing gum) |
| Khader, Jost, Mertens, Bien & Roesler | 2010 | fMRI | NO | EXPLICIT | VERBAL | VISUAL | HIGH | 16 | noun>rhyme generation |
| | | | | | | | | | verb>rhyme generation |
| | | | | | | | | | noun generation>letter detection |
| | | | | | | | | | verb generation>letter detection |
| Kim et al. | 2009 | fMRI | NO | EXPLICIT | VERBAL | VISUAL | HIGH | 36 | sentences>word lists |
| Kinno, Kawamura, Shioda & Sakai | 2008 | fMRI | NO | EXPLICIT | BOTH | VISUAL | HIGH | 14 | canonical sentence>picture & letter strings |
| | | | | | | | | | active sentence>picture & letter strings |
| | | | | | | | | | passive sentence>picture & letter strings |
| Kotz, Cappa, Von Cramon & Friederici | 2002 | fMRI | NO | EXPLICIT | VERBAL | AUDITORY | HIGH | 13 | words>pseudowords |
| Kumar | 2016 | fMRI | NO | EXPLICIT | VERBAL | VISUAL | HIGH | 20 | abstract+concrete words>pseudowords |
| | | | | | | | | | abstract words>pseudowords |
| Kuperberg et al. | 2000 | fMRI | NO | EXPLICIT | VERBAL | AUDITORY | HIGH | 4 | sentences>words strings |
| Kyong, Scott, Rosen, Howe, Agnew & McGettigan | 2014 | fMRI | NO | IMPLICIT | VERBAL | AUDITORY | LOW | 19 | intelligible vocoded>inverted vocoded speech |
| Leff, Schofield, Stephan, Crinion, Friston & Price | 2008 | fMRI | NO | IMPLICIT | VERBAL | AUDITORY | HIGH | 26 | speech>reversed speech |
| Leung & Alain | 2011 | fMRI | NO | EXPLICIT | NONVERBAL | AUDITORY | HIGH | 16 | semantic>location matching |
| Lin, Wang, Zhao, Liu, Li & Bi | 2015 | fMRI | NO | EXPLICIT | VERBAL | VISUAL | HIGH | 20 | words>pseudowords |
| Liu et al. | 2009 | fMRI | NO | EXPLICIT | VERBAL | VISUAL | LOW | 16 | words meaning>slashes |
| | | | | | | | | | words rhyming>slashes |
| | | | | | | | | | words meaning>tones |
| | | | | | | | | | words rhyming>tones |
| Liuzzi et al. | 2017 | fMRI | NO | EXPLICIT | VERBAL | BOTH | HIGH | 18 | semantic judgement > input modality detection |

| | | | | | | | | | |
|---|---|---|---|---|---|---|---|---|---|
| Ludersdorfer, Wimmer, Richlan, Schurz, Hutzler & Kronbichler | 2016 | fMRI | NO | EXPLICIT | VERBAL | AUDITORY | LOW | 29 | words orthographic>tones |
| | | | | | | | | | words semantic>tones |
| Ludersdorfer, Schurz, Richlan, Kronbichler & Wimmer | 2013 | fMRI | NO | EXPLICIT | VERBAL | VISUAL | HIGH | 29 | words>false fonts |
| | | | | | | | | | words>pseudowords |
| | | | | | | | | | speech>reversed speech |
| | | | | | | | | | words>pseudowords |
| Malins, Gumkowski, Buis, Molfese, Rueckl, Frost, Pugh, Morris & Mencl | 2016 | fMRI | NO | IMPLICIT | VERBAL | VISUAL | LOW | 18 | unrelated words>false font |
| | | | | | | | | | unrelated>pseudowords |
| Marques, Canessa & Cappa | 2009 | fMRI | NO | EXPLICIT | VERBAL | VISUAL | LOW | 21 | sentences>crosses |
| Marques, Canessa, Siri, Catricala & Cappa | 2008 | fMRI | NO | EXPLICIT | VERBAL | VISUAL | LOW | 21 | semantic features>baseline task |
| Mashal, Vishne, Laor & Titone | 2013 | fMRI | NO | EXPLICIT | VERBAL | VISUAL | HIGH | 14 | novel metaphor>unrelated words |
| | | | | | | | | | conventional metaphor>unrelated words |
| Matchin, Liao, gaston & Lau | 2019 | fMRI | NO | EXPLICIT | VERBAL | VISUAL | HIGH | 20 | verb phrase>list |
| | | | | | | | | | noun phrase>list |
| Matchin, Hammerly & Lau | 2017 | fMRI | NO | EXPLICIT | VERBAL | VISUAL | HIGH | 16 | sentences>word lists |
| | | | | | | | | | sentences>phrases |
| | | | | | | | | | real>pseudoword lists |
| | | | | | | | | | real>pseudoword phrases |
| | | | | | | | | | real>pseudoword sentences |
| Mellem, Jasmin, Peng & Martin | 2016 | fMRI | NO | IMPLICIT | VERBAL | VISUAL | LOW | 20 | longer>shorter phrase |
| Menz, Blangero, Kunze & Binkofski | 2010 | fMRI | NO | EXPLICIT | NONVERBAL | VISUAL | HIGH | 20 | known>unknown objects |
| Meyer, Alter, Friederici, Lohmann & Yves von Cramon | 2002 | fMRI | NO | IMPLICIT | VERBAL | AUDITORY | HIGH | 14 | word>pseudoword sentence |

| | | | | | | | | | |
|---|---|---|---|---|---|---|---|---|---|
| Moberget, Gullesen, Andersson, Ivry & Endestad | 2014 | fMRI | NO | EXPLICIT | VERBAL | VISUAL | HIGH | 32 | incongruent>scrambled sentence |
| | | | | | | | | | congruent>scrambled sentence |
| Mummery, Patterson, Hodges & Price | 1998 | PET | NO | EXPLICIT | VERBAL | VISUAL | HIGH | 10 | semantic>phonological decision |
| Nichelli, Grafman, Pietrini, Clark, Lee & Miletich | 1995 | PET | NO | EXPLICIT | VERBAL | VISUAL | HIGH | 9 | semantic>orthographic decision |
| Noppeney & Price | 2003 | PET | NO | EXPLICIT | VERBAL | AUDITORY | LOW | 9 | normal>reversed words |
| Orfanidou, Marlsen-Wilson & Davis | 2006 | fMRI | NO | EXPLICIT | VERBAL | AUDITORY | HIGH | 13 | words>pseudowords |
| Pallier, Devauchelle & Dehaene | 2011 | fMRI | NO | EXPLICIT | VERBAL | VISUAL | HIGH | 40 | longer>shorter phrase |
| | | | | | | | | | length of real>pseudoword sentences |
| Peelle, Eason, Schmitter, Schwarzbauer & Davis | 2010 | fMRI | NO | EXPLICIT | VERBAL | AUDITORY | HIGH | 6 | sentences>signal correlated noise |
| Perani, Schnur, Tettamanti, Gorno-Tempini, Cappa & Fazio | 1999 | PET | NO | EXPLICIT | NONVERBAL | VISUAL | HIGH | 11 | living objects>shapes |
| | | | | | | | | | nonliving objects>shapes |
| Perani, Schnur, Tettamanti, Gorno-Tempini, Cappa & Fazio | 1999 | PET | NO | EXPLICIT | VERBAL | VISUAL | HIGH | 8 | living words>pseudowords |
| | | | | | | | | | nonliving words>pseudowords |
| Perrone-Bertolotti, Kauffmann, Pichat, Vidal & Baciu | 2017 | fMRI | NO | EXPLICIT | VERBAL | VISUAL | HIGH | 24 | words>unreadable font |
| Pilgrim, Fadili, Fletcher & Tyler | 2002 | fMRI | NO | EXPLICIT | VERBAL | VISUAL | HIGH | 14 | words>letter strings |
| Price, Moore, Humphreys & Wise | 1997 | PET | NO | EXPLICIT | VERBAL | VISUAL | HIGH | 6 | semantic>phonological decision |
| Pulvermueller, Cook & Hauk | 2012 | fMRI | NO | IMPLICIT | VERBAL | VISUAL | LOW | 23 | phrases>hashmarks |
| | | | | | | | | | uninflected words>hashmarks |
| | | | | | | | | | inflected words>hashmarks |
| Raettig & Kotz | 2008 | fMRI | NO | EXPLICIT | VERBAL | AUDITORY | HIGH | 16 | real words>pseudowords |
| Raposo, Frade & Alves | 2016 | fMRI | NO | IMPLICIT | VERBAL | VISUAL | HIGH | 18 | semantic>perceptual decision |

| | | | | | | | | | |
|---|---|---|---|---|---|---|---|---|---|
| Raposo, Moss, Stamatakis & Tyler | 2009 | fMRI | NO | IMPLICIT | VERBAL | AUDITORY | LOW | 22 | action sentences>SC noise |
| Rapp & Lipka | 2011 | fMRI | NO | IMPLICIT | VERBAL | VISUAL | LOW | 10 | words>checkerboards |
| | | | | | | | | | words>letter strings |
| Redcay, Velnoskey & Rowe | 2016 | fMRI | NO | EXPLICIT | BOTH | VISUAL | HIGH | 24 | meaningful>meaningless stimuli |
| | | | | | | | | | communicative>non-communicative gesture |
| | | | | | | | | | real>pseudoword sentences |
| Rissman, Eliassen & Blumstein | 2003 | fMRI | NO | IMPLICIT | VERBAL | AUDITORY | HIGH | 15 | words>pseudowords |
| Robertson et al. | 2000 | fMRI | NO | IMPLICIT | VERBAL | VISUAL | HIGH | 8 | indefinite article sentence>letter strings |
| | | | | | | | | | definite article sentence>letter strings |
| Rodd, Johnsrude & Davis | 2012 | fMRI | NO | EXPLICIT | VERBAL | AUDITORY | LOW | 15 | speech>SCN |
| Rodd, Longe, Randall & Tyler | 2010 | fMRI | NO | EXPLICIT | VERBAL | AUDITORY | LOW | 14 | speech>SCN |
| Rogalsky & Hickok | 2009 | fMRI | NO | IMPLICIT | VERBAL | AUDITORY | HIGH | 14 | sentences>word lists |
| Rogalsky, Almeida, Sprouse & Hickok | 2015 | fMRI | NO | IMPLICIT | VERBAL | AUDITORY | HIGH | 15 | words>scrambled |
| Rogers et al. | 2006 | PET | NO | EXPLICIT | BOTH | VISUAL | HIGH | 12 | pictures>scrambled pictures |
| | | | | | | | | | specific-level judgement>baseline |
| Roskies, Fiez, Balota, Raichle & Petersen | 2001 | PET | NO | EXPLICIT | VERBAL | VISUAL | HIGH | 20 | semantic>phonological decision |
| Roxbury, McMahon & Copland | 2014 | fMRI | NO | EXPLICIT | VERBAL | AUDITORY | HIGH | 17 | concrete word>pseudoword |
| | | | | | | | | | abstract word>pseudoword |
| Ryan, Cox, Hayes & Nadel | 2008 | fMRI | NO | EXPLICIT | VERBAL | VISUAL | HIGH | 10 | semantic fluency (generate) > crosses |
| | | | | | | | | | semantic fluency (recall) > crosses |
| | | | | | | | | | semantic fluency (recall & geneerate)>crosses |
| Ryan, Lin, Ketcham & Nadel | 2010 | fMRI | NO | EXPLICIT | VERBAL | VISUAL | HIGH | 15 | semantic spatial old>letter judgement |
| | | | | | | | | | semantic spatial new>letter judgement |
| | | | | | | | | | semantic non spatial old>letter judgement |
| | | | | | | | | | semantic non spatial new>letter judgement |
| | | | | | | | | | semantic new>episodic judgement |

| | | | | | | | | | |
|---|---|---|---|---|---|---|---|---|---|
| Sabri, Binder, Desai, Medler, Leitl & Liebenthal | 2008 | fMRI | NO | EXPLICIT | VERBAL | AUDITORY | HIGH | 28 | speech>rotated speech |
| | | | | | | | | | words>pseudowords |
| Sachs, Weis, Krings, Huber & Kircher | 2008 | fMRI | NO | EXPLICIT | VERBAL | VISUAL | HIGH | 14 | biased thematic judgement>letters |
| | | | | | | | | | biased taxonomic judgement>letters |
| | | | | | | | | | balanced taxonomic judgement>letters |
| | | | | | | | | | balanced taxonomic judgement>letters |
| Saur et al. | 2008 | fMRI | NO | IMPLICIT | VERBAL | AUDITORY | HIGH | 33 | word>pseudoword sentences |
| Schell, Zaccarella & Friederici | 2017 | fMRI | NO | EXPLICIT | VERBAL | AUDITORY | HIGH | 21 | phrases>non combinatorial words |
| Schmitt, Auer & Ferstl | 2019 | fMRI | NO | EXPLICIT | VERBAL | AUDITORY | HIGH | 40 | known>unknown language |
| Schuil, Smits & Zwaan | 2013 | fMRI | NO | EXPLICIT | VERBAL | VISUAL | HIGH | 20 | senteces>pseudowords |
| | | | | | | | | | verbs>pseudowords |
| | | | | | | | | | literal sentences>pseudowords |
| | | | | | | | | | nonliteral sentences>pseudowords |
| Scott, Blank, Rosen & Wise | 2000 | PET | NO | IMPLICIT | VERBAL | AUDITORY | HIGH | 8 | intelligible>unintelligible speech |
| Segal & Petrides | 2012 | fMRI | NO | EXPLICIT | NONVERBAL | VISUAL | HIGH | 90 | writing>copying |
| | | | | | | | | | words>pseudowords |
| Seghier, Josse, Leff & Price | 2011 | fMRI | NO | EXPLICIT | BOTH | VISUAL | HIGH | 60 | meaningful>meaningless stimuli |
| Sheldon, McAndrews, Pruessner & Moscovitch | 2016 | fMRI | NO | EXPLICIT | VERBAL | VISUAL | HIGH | 15 | semantic fluency>perceptual task |
| Simard, Monetta, Nagano-Saito & Monchi | 2013 | fMRI | NO | EXPLICIT | VERBAL | VISUAL | HIGH | 14 | semantic>control macthing |
| | | | | | | | | | semantic matching>phonological decision (syllable rhyme) |
| | | | | | | | | | semantic matching>phonological decision (syllable onset) matching |
| Slioussar, Kireev, Chernigovskaya, Kataeva, Korotkov & Medvedev | 2014 | fMRI | NO | EXPLICIT | VERBAL | VISUAL | HIGH | 21 | real verbs>pseudowords |
| | | | | | | | | | real nouns>pseudowords |
| Smith, Myers, Sethi, Pantazatos, Yanagihara & Hirsch | 2012 | fMRI | NO | EXPLICIT | VERBAL | VISUAL | HIGH | 14 | semantics>baseline |

| | | | | | | | | | |
|---|---|---|---|---|---|---|---|---|---|
| Snijders, Vosse, Kempen, Van Berkum, Petersson & Hagoort | 2009 | fMRI | NO | IMPLICIT | VERBAL | VISUAL | HIGH | 28 | sentences>word lists |
| Stowe, Paans, Wijers, Zwarts, Mulder & Vaalburg | 1999 | PET | NO | IMPLICIT | VERBAL | VISUAL | HIGH | 12 | sentences>scrambled word lists |
| Straube, Green, Weis & Kircher | 2012 | fMRI | NO | IMPLICIT | VERBAL | AUDITORY | HIGH | 16 | known>unknown language |
| | | | | | | | | | iconic>meaningless gesture |
| Stringaris, Medford, Giampietro. Brammer & David | 2007 | fMRI | NO | EXPLICIT | VERBAL | VISUAL | HIGH | 11 | literal>meaningless sentences |
| | | | | | | | | | methaphors>meaningless sentences |
| Sun, Xue, Zhang, Zuo, Chen, Wang, Martin, Wang, Chen, He & Wang | 2017 | fMRI | NO | EXPLICIT | VERBAL | VISUAL | HIGH | 11 | semantic>orthography judgement |
| Szlachta, Bozic, Jelowicka & Marslen-Wilson | 2012 | fMRI | NO | IMPLICIT | VERBAL | VISUAL | LOW | 21 | words>musical rain |
| | | | | | | | | | nouns>musical rain |
| | | | | | | | | | inflected nouns>musical rain |
| Takeichi, Koyama, Terao, Takeuchi, Toyosawa & Murohashi | 2010 | fMRI | NO | IMPLICIT | VERBAL | AUDITORY | HIGH | 23 | speech>reversed speech |
| | | | | | | | | | speech>modulated speech |
| Taminato, Miura, Sugiura & Kawashima | 2014 | fMRI | NO | EXPLICIT | NONVERBAL | VISUAL | LOW | 35 | object recognition>control task |
| | | | | | | | | | object recognition>control task |
| Thierry & Price | 2006 | PET | NO | EXPLICIT | VERBAL | AUDITORY | HIGH | 12 | auditory words>speech control(scrambled) |
| | | | | | | | | | auditory sounds>sound control (scrambled) |
| Thierry & Price | 2006 | PET | NO | EXPLICIT | VERBAL | VISUAL | HIGH | 12 | visual words>text control (scrambled letter strings) |
| | | | | | | | | | visual videos>video control (distorted) |
| Tieleman, Seurinck, Deblaere, Vandemaele, Vingerhoets & Achten | 2005 | fMRI | NO | EXPLICIT | VERBAL | VISUAL | HIGH | 22 | self-paced semantic>perceptual decision |
| | | | | | | | | | fixed-paced semantic>perceptual decision |

| | | | | | | | | | |
|---|---|---|---|---|---|---|---|---|---|
| Tyler, Stamatakis, Dick, Bright, Fletcher & Moss | 2003 | fMRI | NO | EXPLICIT | VERBAL | VISUAL | HIGH | 12 | animals>baseline |
| | | | | | | | | | tool action words>baseline |
| | | | | | | | | | biological action>baseline |
| Vagharchakian, Dehaene-Lambertz, Pallier & Dehaene | 2012 | fMRI | NO | EXPLICIT | VERBAL | BOTH | HIGH | 16 | intelligible>unintelligble compression rate |
| | | | | | | | | | intelligible>unintelligble compression rate |
| | | | | | | | | | intelligible>unintelligble compression rate |
| Van Ettinger-Veenstra, McAllister, Lundberg, Karlsson & Engstrom | 2016 | fMRI | NO | EXPLICIT | VERBAL | VISUAL | HIGH | 27 | sentences>symbol strings |
| van Leeuwen et al. | 2014 | fMRI | NO | EXPLICIT | BOTH | BOTH | HIGH | 16 | speech>reversed speech |
| Vignali, Hawelka, Hutzler & Richlan | 2019 | fMRI | NO | EXPLICIT | VERBAL | VISUAL | HIGH | 21 | foveal & parafovealwords>foveal & parafoveal pseudowords |
| Vingerhoets | 2008 | fMRI | NO | IMPLICIT | NONVERBAL | VISUAL | HIGH | 14 | familiar>unfamiliar tools |
| Visser, Jefferies, Embleton & Lambon Ralph | 2012 | fMRI | NO | EXPLICIT | BOTH | VISUAL | HIGH | 15 | semantics>baseline |
| | | | | | | | | | pictures>baseline |
| | | | | | | | | | words>baseline |
| Vitello, Warren, Devlin & Rodd | 2014 | fMRI | NO | EXPLICIT | VERBAL | AUDITORY | LOW | 20 | sentences>SCN |
| von Kriegstein, Eger, Kleinschmidt & Giraud | 2003 | fMRI | NO | EXPLICIT | VERBAL | AUDITORY | HIGH | 14 | sentence>speech envelope |
| Wang, Zhao, Zevin & Yang | 2016 | fMRI | NO | EXPLICIT | VERBAL | VISUAL | HIGH | 16 | words>nonsense strokes |
| Weiss, Katzir & Bitan | 2015 | fMRI | NO | IMPLICIT | VERBAL | VISUAL | LOW | 18 | pointed words>asterisks |
| | | | | | | | | | unpointed words>asterisks |
| Welcome & Joanisse | 2012 | fMRI | NO | EXPLICIT | VERBAL | VISUAL | HIGH | 20 | semantic>phonological/orthographic decision |
| Wende, Straube, Stratmann, Sommer, Kircher & Nagels | 2012 | fMRI | NO | EXPLICIT | VERBAL | VISUAL | HIGH | 18 | semantic>phonological fluency |
| Wirth, Jann, Dierks, Federspiel, Wiest & Horn | 2011 | fMRI | NO | EXPLICIT | VERBAL | VISUAL | HIGH | 19 | semantic>phonological & perceptual decision |
| Wright et al. | 2008 | fMRI | NO | EXPLICIT | BOTH | VISUAL | HIGH | 10 | semantic>perceptual matching |
| | | | | | | | | | semantic>perceptual matching |
| | | | | | | | | | semantic>perceptual matching |

| | | | | | | | | | |
|---|---|---|---|---|---|---|---|---|---|
| Wright, Randall, Marslen-Wilson & Tyler | 2011 | fMRI | NO | BOTH | VERBAL | AUDITORY | LOW | 14 | speech>musical rain |
| Wu, Mai, Tang, Ge, Luo & Liu | 2013 | fMRI | NO | IMPLICIT | VERBAL | VISUAL | LOW | 19 | arm words>checkerboard |
| | | | | | | | | | leg words>checkerboard |
| | | | | | | | | | mouth words>checkerboard |
| Xiao et al. | 2005 | fMRI | NO | EXPLICIT | VERBAL | AUDITORY | HIGH | 14 | words>pseudowords |
| Yang, Li, Fang, Shu, Liu & Chen | 2016 | fMRI | NO | IMPLICIT | VERBAL | VISUAL | LOW | 20 | opaque idioms>hashmarks |
| | | | | | | | | | transparent idioms>hashmarks |
| | | | | | | | | | literal phrases>hashmarks |
| Zaccarella & Friederici | 2015 | fMRI | NO | EXPLICIT | VERBAL | VISUAL | HIGH | 22 | words>pseudowords |
| Zhang, Liu & Zhang | 2014 | fMRI | NO | EXPLICIT | VERBAL | VISUAL | LOW | 18 | nonliving words>asterisks |
| | | | | | | | | | living words>asterisks |
| Zhang, Xiao & Weng | 2012 | fMRI | NO | EXPLICIT | VERBAL | VISUAL | HIGH | 14 | words>pseudowords |
| Zhuang & Devereux | 2017 | fMRI | NO | EXPLICIT | VERBAL | AUDITORY | HIGH | 16 | phrases>words |
| Zou, Packard, Xia, Liu & Shu | 2016 | fMRI | NO | IMPLICIT | VERBAL | AUDITORY | LOW | 17 | speech>tone |
| | | | | | | | | | speech>tone |
| | | | | | | | | | speech>tone |
| | | | | | | | | | identical speech>tone |
| | | | | | | | | | words>tone |
| Zvyagintsev, Clemens, Chechko, Mathiak, Sack & Mathiak | 2013 | fMRI | NO | IMPLICIT | NONE | NONE | HIGH | 15 | visual imagery>counting |
| | | | | | | | | | auditory imagery>counting |
| EXCLUDED CONTRASTS WITH STIMULI CONTAINING SOCIAL CONTENT | | | | | | | | | |
| Brambati, Benoit, Monetta, Belleville & Joubert | 2010 | fMRI | YES | EXPLICIT | BOTH | BOTH | HIGH | 12 | general & specific occupation judgement>baseline (scrambled face) |
| Chiao, Harada, Oby, Li, Parrish & Bridge | 2009 | fMRI | YES | EXPLICIT | NONVERBAL | VISUAL | LOW | 12 | uniform status judgement>colour change detection |
| | | | | | | | | | face status judgement>colour change detection |
| | | | | | | | | | car status judgement>colour change detection |

| | | | | | | | | | |
|---|---|---|---|---|---|---|---|---|---|
| Damasio, Tranel, Grabowski, Adolphs & Damasio | 2004 | PET | YES | EXPLICIT | NONVERBAL | VISUAL | HIGH | 55 | persons>face orientation judgement |
| Dreyer & Pulvermueller | 2018 | fMRI | YES | IMPLICIT | VERBAL | VISUAL | LOW | 28 | all nouns>hashmarks |
| | | | | | | | | | abstract emotional nouns>baseline (hashmarks) |
| | | | | | | | | | abstract mental nouns>baseline (hashmarks) |
| | | | | | | | | | food nouns>baseline (hashmarks) |
| | | | | | | | | | tool nouns>baseline (hashmarks) |
| Elfgren, Westen, Passant, Larsson, Mannfolk & Fransson | 2006 | fMRI | YES | EXPLICIT | NONVERBAL | VISUAL | HIGH | 15 | familiar>unfamiliar faces (identification) |
| Gesierich et al. | 2012 | fMRI | YES | EXPLICIT | NONVERBAL | VISUAL | HIGH | 21 | familiar>scrambled faces |
| | | | | | | | | | familiar > unfamiliar faces |
| | | | | | | | | | familiar>unfamiliar faces |
| Gorno-Tempini et al. | 1998 | PET | YES | EXPLICIT | NONVERBAL | VISUAL | HIGH | 6 | famous faces>controls |
| | | | | | | | | | famous names>controls |
| | | | | | | | | | double famous proper names>controls |
| | | | | | | | | | non-famous faces>controls |
| | | | | | | | | | Non-famous names>controls |
| | | | | | | | | | Double common names>controls |
| Grabowski, Damasio, Tranel, Boles Ponto, Hichwa & Damasio | 2001 | PET | YES | EXPLICIT | NONVERBAL | VISUAL | HIGH | 10 | naming persons>building orientation judgement |
| | | | | | | | | | naming persons>face orientation judgement |
| Kuchinke et al. | 2005 | fMRI | YES | IMPLICIT | VERBAL | VISUAL | HIGH | 20 | emotion word>nonword |
| Leveroni et al. | 2000 | fMRI | YES | EXPLICIT | NONVERBAL | VISUAL | HIGH | 11 | familiar faces>foils(never seen faces) |
| | | | | | | | | | newly learned faces>foils(never seen faces) |
| Moseley, Carota, Hauk, Mohr & Pulvermueller | 2012 | fMRI | YES | IMPLICIT | VERBAL | VISUAL | LOW | 18 | all emotional words>hashmarks |
| | | | | | | | | | abstract emotional words>hashmarks |
| | | | | | | | | | arm+face+emotion words>hashmarks |
| | | | | | | | | | face words>hashmarks |
| | | | | | | | | | arm words>hashmarks |

| | | | | | | | | | |
|---|---|---|---|---|---|---|---|---|---|
| Nakamura et al. | 2000 | PET | YES | EXPLICIT | NONVERBAL | VISUAL | LOW | 7 | familiar faces>fixation cross |
| Nakamura et al. | 2001 | PET | YES | EXPLICIT | VERBAL | AUDITORY | HIGH | 9 | familiar voice>vowel discrimination |
| | | | | | | | | | self voice>vowel discrimination |
| Nielson et al. | 2010 | fMRI | YES | EXPLICIT | BOTH | VISUAL | HIGH | 17 | familiar>unfamiliar people |
| Ross & Olson | 2012 | fMRI | YES | EXPLICIT | NONVERBAL | VISUAL | HIGH | 11 | famous>unknown faces & landmarks |
| Sergent, Otha & Macdonald | 1992 | PET | YES | EXPLICIT | VERBAL | VISUAL | HIGH | 7 | face identity>gender discrimination |
| | | | | | | | | | object recognition>gratings |
| Sugiura et al. | 2006 | fMRI | YES | EXPLICIT | VERBAL | VISUAL | HIGH | 24 | famous>unfamiliar names |
| | | | | | | | | | personal>unfamiliar names |
| Sugiura et al. | 2001 | PET | YES | EXPLICIT | NONVERBAL | VISUAL | HIGH | 5 | identity discrimination>control |
| | | | | | | | | | identity discrimination>face direction |
| Sugiura et al. | 2008 | fMRI | YES | EXPLICIT | VERBAL | VISUAL | HIGH | 25 | low familiar>unfamiliar names |
| | | | | | | | | | personal familiar>unfamiliar names |
| | | | | | | | | | high familiar>unfamiliar names |
| | | | | | | | | | high familiar>unfamiliar names |
| Taylor, Arsalidou, Bayless, Morris, Evans & Barbeau | 2009 | fMRI | YES | IMPLICIT | NONVERBAL | VISUAL | HIGH | 10 | own>unfamiliar face |
| | | | | | | | | | partner's>unfamiliar face |
| | | | | | | | | | parent's>unfamiliar face |

**The references to access the listed studies are listed at the end of the document*

***Table M2.1 SC VERBAL*** *List of studies included in the* ***semantic cognition VERBAL STIMULUS DOMAIN*** *(N = 175) meta-analysis after excluding NON-VERBAL contrasts.* ***Note:*** *The list of excluded studies with NON-VERBAL STIMULUS DOMAIN can be seen in Table M.2.2 SC NON-VERBAL. Contrasts with BOTH VEBAL and NON-VERBAL stimuli were also excluded and are listed at the end of the document highlighted in grey. SC= Semantic Cognition, N= Sample Size*

| *Authors | Year | Imaging Method | Social Content | Instructional Cue | Stimulus Domain | Sensory Input Modality | Baseline Type | N | Contrast |
|---|---|---|---|---|---|---|---|---|---|
| AbdulSabur et al. | 2014 | fMRI | NO | EXPLICIT | VERBAL | VISUAL | LOW | 18 | narrative production>recitation |
| Abraham et al. | 2012 | fMRI | NO | EXPLICIT | VERBAL | VISUAL | HIGH | 19 | high&low divergent thinking (semantic)>1&2 back letter identity (working memory) |
| Alain, He & Grady | 2008 | fMRI | NO | EXPLICIT | VERBAL | AUDITORY | HIGH | 16 | sound category>sound location |
| Assadollahi, Meinzer, Flaisch, Obleser & Rockstroh | 2009 | fMRI | NO | IMPLICIT | VERBAL | VISUAL | HIGH | 20 | nouns followed by 1&3 argument verbs>letter strings followed by 1&3argument verbs |
| Axmacher, Bialleck, Weber, Helmstaedter, Elger & Fell | 2009 | fMRI | NO | EXPLICIT | VERBAL | VISUAL | HIGH | 32 | word decision>spatial decision |
| Bagga et al. | 2013 | fMRI | NO | EXPLICIT | VERBAL | VISUAL | HIGH | 18 | semantic>case matching judgement |
| Barros-Loscertales et al. | 2012 | fMRI | NO | EXPLICIT | VERBAL | VISUAL | LOW | 59 | control words>hashmarks baseline |
| Baumgaertener, Weiller & Buchel | 2002 | fMRI | NO | EXPLICIT | VERBAL | VISUAL | HIGH | 9 | word>pseudoword in sentence |
| Baumgaertner et al. | 2007 | fMRI | NO | IMPLICIT | VERBAL | AUDITORY | HIGH | 19 | sentences>reversed sentences |
| Bautista & Wilson | 2016 | fMRI | NO | IMPLICIT | VERBAL | AUDITORY | HIGH | 12 | clear>scrambled rotated speech |
| Bick, Goelman & Frost | 2008 | fMRI | NO | EXPLICIT | VERBAL | VISUAL | LOW | 14 | semantic>visual control |
| | | | | | | | | | morphologiacl>visual control |
| | | | | | | | | | orthographic>visual control |
| | | | | | | | | | phonological>visual control |
| Binder, Frost, Hammeke, Bellgowan, Rao & Cox | 1999 | fMRI | NO | EXPLICIT | VERBAL | AUDITORY | HIGH | 30 | semantic>phonological decision |
| Binder et al. | 2003 | fMRI | NO | EXPLICIT | VERBAL | VISUAL | HIGH | 24 | word>nonword |
| Birn et al. | 2010 | fMRI | NO | EXPLICIT | VERBAL | VISUAL | LOW/HIGH | 14 | category & letter fluency>months ( automatic speech) |
| | | | | | | | | | category>letter fluency |

| | | | | | | | | | |
|---|---|---|---|---|---|---|---|---|---|
| Bonhage, Fiebach, Bahlmann & Mueller | 2014 | fMRI | NO | EXPLICIT | VERBAL | VISUAL | HIGH | 18 | sentence fragments>ungrammatical word strings |
| Bonhage, Mueller, Friederici & Fiebach | 2015 | fMRI | NO | EXPLICIT | VERBAL | VISUAL | HIGH | 18 | sentences>jabberwocky sentences |
| Booth et al. | 2006 | fMRI | NO | EXPLICIT | VERBAL | VISUAL | HIGH/LOW | 13 | meaning (semantic) judgement>rhyming( phonological) judgement |
| | | | | | | | | | meaning(semantic)>control(symbols) |
| Boulenger, Hauk & Pulvermuller | 2009 | fMRI | NO | EXPLICIT | VERBAL | VISUAL | LOW | 18 | sentences> hashmarks |
| | | | | | | | | | sentences> hashmarks |
| Bozic & Marslen-Wilson | 2013 | fMRI | NO | IMPLICIT | VERBAL | AUDITORY | HIGH | 13 | speech>musical rain |
| Bulut, Hung, Tzeng & Wu | 2017 | fMRI | NO | IMPLICIT | VERBAL | VISUAL | HIGH | 20 | sentences>unstructured word lists |
| | | | | | | | | | sentences>unstructured character list |
| Cai, Kochiyama, Osaka & Wu | 2007 | fMRI | NO | IMPLICIT | VERBAL | AUDITORY | HIGH | 15 | words>nonsense words |
| Cao, Peng, Liu, Jin, Fan, Deng, & Booth | 2009 | fMRI | NO | EXPLICIT | VERBAL | VISUAL | LOW | 13 | meaning>perceptual decision |
| Cappa, Perani, Schnur, Tettamanti & Fazio | 1998 | PET | NO | EXPLICIT | VERBAL | VISUAL | HIGH | 13 | words>pseudowords |
| | | | | | | | | | animal visual knowledge>pseudowords |
| | | | | | | | | | tool visual knowledge>pseudowords |
| | | | | | | | | | animal associative knowledge>pseudowords |
| | | | | | | | | | tools functional knowledge>pseudowords |
| Carota, Kriegeskorte, Nili & Pulvermuller | 2017 | fMRI | NO | IMPLICIT | VERBAL | VISUAL | LOW | 23 | words>hashmarks |
| Carota, Moseley & Pulvermueller | 2012 | fMRI | NO | IMPLICIT | VERBAL | VISUAL | LOW | 18 | all words>hashmarks |
| | | | | | | | | | tool words>hashmarks |
| | | | | | | | | | animal words>hashmarks |
| | | | | | | | | | food words>hashmarks |
| Chan, Tang, Tang, Lee, Lo & Kwong | 2009 | fMRI | NO | IMPLICIT | VERBAL | VISUAL | HIGH | 22 | synonyms>pseudocharacters |
| | | | | | | | | | synonyms>Korean characters (unknown) |
| Chou, Chen, Wu & Booth | 2009 | fMRI | NO | EXPLICIT | VERBAL | VISUAL | HIGH | 31 | related words>false font |
| | | | | | | | | | unrelated words>false font |

| | | | | | | | | | |
|---|---|---|---|---|---|---|---|---|---|
| Chou, Chen, Wu & Booth | 2009 | fMRI | NO | EXPLICIT | VERBAL | VISUAL | HIGH | 32 | related words>false font |
| | | | | | | | | | unrelated words>false font |
| Chow, Kaup, Raabe & Greenlee | 2008 | fMRI | NO | EXPLICIT | VERBAL | VISUAL | HIGH | 15 | predictive&normal reading>pseudoword reading |
| | | | | | | | | | normal reading>pseudoword reading |
| | | | | | | | | | predictive reading>pseudoword reading |
| Christensen, Antonucci, Lockwood, Kittleson & Plante | 2008 | fMRI | NO | EXPLICIT | VERBAL | AUDITORY | HIGH | 14 | diotic listening>reversed speech |
| | | | | | | | | | dichotic listening>reversed speech |
| Clos, Langner, Meyer, Oechslin, Zilles & Eickhoff | 2014 | fMRI | NO | EXPLICIT | VERBAL | AUDITORY | HIGH | 29 | intelligibilty based on cue > unintelligible |
| Davis, Meunier & Marslen-Wilson | 2004 | fMRI | NO | EXPLICIT | VERBAL | VISUAL | HIGH | 11 | words>letter strings |
| Davis, Ford, Kherif & Johnsrude | 2011 | fMRI | NO | EXPLICIT | VERBAL | AUDITORY | LOW | 12 | clear & anomalous sentences>signal correlated noise |
| Demonet et al. | 1992 | PET | NO | EXPLICIT | VERBAL | AUDITORY | HIGH/LOW | 9 | words>phonemes |
| | | | | | | | | | words>tones |
| Devlin, Matthews & Rushworth | 2003 | fMRI | NO | EXPLICIT | VERBAL | VISUAL | HIGH | 12 | semantic>phonological judgement |
| Devlin et al. | 2002 | PET | NO | EXPLICIT | VERBAL | VISUAL | HIGH | 12 | all semantic>letter detection |
| | | | | | | | | | all semantic>letter detection |
| Devlin et al. | 2002 | PET | NO | EXPLICIT | VERBAL | VISUAL | HIGH | 8 | all semantic>letter categorisation |
| Devlin et al. | 2002 | fMRI | NO | EXPLICIT | VERBAL | VISUAL | HIGH | 8 | all semantic> letter categorisation |
| Devlin et al. | 2000 | PET | NO | EXPLICIT | VERBAL | VISUAL | HIGH | 8 | semantic categorisation>letter categorisation |
| Devlin et al. | 2000 | fMRI | NO | EXPLICIT | VERBAL | VISUAL | HIGH | 8 | semantic categorisation>letter categorisation |
| Diaz & McCarthy | 2009 | fMRI | NO | IMPLICIT | VERBAL | VISUAL | HIGH | 16 | all words>nonwords |
| Dreyer & Pulvermueller | 2018 | fMRI | YES | IMPLICIT | VERBAL | VISUAL | LOW | 28 | all nouns>hashmarks |
| | | | | | | | | | abstract emotional nouns>baseline (hashmarks) |
| | | | | | | | | | abstract mental nouns>baseline (hashmarks) |
| | | | | | | | | | food nouns>baseline (hashmarks) |
| | | | | | | | | | tool nouns>baseline (hashmarks) |

| | | | | | | | | | |
|---|---|---|---|---|---|---|---|---|---|
| Emmorey, Weisberg, McCullough & Petrich | 2013 | fMRI | NO | EXPLICIT | VERBAL | VISUAL | HIGH | 14 | words(semantic judgement)>false fonts |
| | | | | | | | | | words(phonological)>false fonts |
| Erb, Henry, Eisner & Obleser | 2013 | fMRI | NO | EXPLICIT | VERBAL | AUDITORY | HIGH | 30 | speech>vocoded speech |
| Foki, Gartus, Geissler & Beisteiner | 2008 | fMRI | | EXPLICIT | VERBAL | VISUAL | LOW | 23 | semantic judgement>tongue movements |
| Friederici, Kotz, Scott & Obleser | 2010 | fMRI | NO | IMPLICIT | VERBAL | AUDITORY | HIGH | 17 | intelligible speech>rotated speech |
| Friese, Rutschmann, Raabe & Schmalhofer | 2008 | fMRI | NO | EXPLICIT | VERBAL | VISUAL | HIGH | 13 | words>pseudowords |
| Garbin, Collina & Tabossi | 2012 | fMRI | NO | EXPLICIT | VERBAL | VISUAL | HIGH | 12 | object noun>pseudoword |
| | | | | | | | | | event noun>pseudoword |
| | | | | | | | | | verb>pseudoword |
| Geranmayeh, Brownsett, Leech, Beckmann, Woodhead & Wise | 2012 | fMRI | NO | EXPLICIT | VERBAL | VISUAL | HIGH | 19 | speech>tongue movements |
| Giraud et al. | 2004 | fMRI | NO | EXPLICIT | VERBAL | AUDITORY | HIGH | 8 | natural speech>speech envelope |
| Gitelman, Nobre, Sonty, Parrish & Mesulam | 2005 | fMRI | NO | EXPLICIT | VERBAL | VISUAL | HIGH | 14 | semantic>control task |
| Gorno-Tempini et al. | 1998 | PET | YES | EXPLICIT | VERBAL | VISUAL | HIGH | 6 | famous names>controls |
| | | | | | | | | | double famous proper names>controls |
| | | | | | | | | | Non-famous names>controls |
| | | | | | | | | | Double common names>controls |
| Graves, Binder, Desai, Conant & Seidenberg | 2010 | fMRI | NO | EXPLICIT | VERBAL | VISUAL | HIGH | 23 | forward>reverse phrases |
| Graves, Binder, Desai, Conant & Seidenberg | 2010 | fMRI | NO | EXPLICIT | VERBAL | VISUAL | HIGH | 22 | forward>reverse phrases |
| Grindrod, Garnett, Malyutina & den Ouden | 2014 | fMRI | NO | EXPLICIT | VERBAL | VISUAL | HIGH | 23 | words>nonwords |
| Grossman et al. | 2002a | fMRI | NO | IMPLICIT | VERBAL | VISUAL | HIGH | 16 | all nouns > pseudowords |
| | | | | | | | | | implements>pseudowords |
| | | | | | | | | | animals>pseudowords |
| | | | | | | | | | abstract>pseudowords |
| Grossman et al., 2002b | 2002 b | fMRI | NO | IMPLICIT | VERBAL | VISUAL | HIGH | 16 | verbs>pseudowords |

| | | | | | | | | | |
|---|---|---|---|---|---|---|---|---|---|
| Groussard et al. | 2010 | PET | NO | EXPLICIT | VERBAL | VISUAL | HIGH | 11 | verbal semantics>verbal reference |
| Guediche, Reilly, Santiago, Laurent & Blumstein | 2016 | fMRI | NO | IMPLICIT | VERBAL | AUDITORY | LOW | 16 | related > repeated sentence |
| | | | | | | | | | unrelated > repeated sentence |
| Gurd et al. | 2002 | fMRI | NO | EXPLICIT | VERBAL | AUDITORY | HIGH | 11 | category>rote fluency |
| Haberling, Corballis &Corballis | 2016 | fMRI | NO | EXPLICIT | VERBAL | VISUAL | HIGH | 92 | synonyms>letter strings |
| Hagoort et al. | 1999 | PET | NO | IMPLICIT | VERBAL | VISUAL | HIGH | 10 | words>pseudowords |
| Hartung, Hagoort & Willems | 2017 | fMRI | NO | IMPLICIT | VERBAL | AUDITORY | HIGH | 52 | first-person speech>unintelligible reversed speech |
| | | | | | | | | | third-person speech>unintelligible reversed speech |
| Hauk & Pulvermueller | 2011 | fMRI | NO | IMPLICIT | VERBAL | VISUAL | LOW | 21 | action words>hashmarks |
| | | | | | | | | | uni manual action words>hashmarks |
| | | | | | | | | | uni manual action words>hashmarks |
| Hayashi et al. | 2014 | fMRI | NO | EXPLICIT | VERBAL | VISUAL | LOW | 16 | concrete word>astersisks |
| | | | | | | | | | abstract word>asterisks |
| Heim, Eickhoff & Amunts | 2008 | fMRI | NO | EXPLICIT | VERBAL | VISUAL | HIGH | 28 | semantic > phonological fluency |
| Henke et al. | 1999 | PET | NO | EXPLICIT | VERBAL | VISUAL | HIGH | 12 | associative word learning>single word encoding |
| Herbster et al. | 1997 | PET | NO | EXPLICIT | VERBAL | VISUAL | HIGH | 10 | Irregular>zero order speak |
| | | | | | | | | | regular>zero order speak |
| | | | | | | | | | irregular+regular>zero order speak |
| Hervais-Adelman, Carlyon, Johnsrude & Davis | 2012 | fMRI | NO | EXPLICIT | VERBAL | AUDITORY | HIGH | 15 | clear>vocoded speech (uninteligible) |
| Higuchi, Moriguchi, Murakami, Katsunuma, Mishima &Uno | 2015 | fMRI | NO | IMPLICIT | VERBAL | VISUAL | LOW | 28 | all characters>checkerboard |
| Homae, Yahata & Sakai | 2003 | fMRI | NO | EXPLICIT | VERBAL | AUDITORY/ VISUAL | HIGH | 10 | auditory sentences>auditory non words |
| | | | | | | | | | visual sentences>visual non words words |
| Hwang, Palmer, Basho, Zadra & Muller | 2009 | fMRI | NO | EXPLICIT | VERBAL | AUDITORY | LOW | 13 | fluency generation>production baseline |
| Ikuta et al. | 2006 | fMRI | NO | IMPLICIT | VERBAL | VISUAL | LOW | 34 | sentences>word lists |

| | | | | | | | | | |
|---|---|---|---|---|---|---|---|---|---|
| Jackson, Hoffman, Pobric, Lambon Ralph | 2015 | fMRI | NO | EXPLICIT | VERBAL | VISUAL | HIGH | 24 | words>letter strings |
| Jensen, Hargreaves, Bass, Pexman, Goodyear & Federico | 2011 | fMRI | NO | EXPLICIT | VERBAL | VISUAL | HIGH | 12 | words>pseudowords |
| Jeon, Lee, Kim & Cho | 2009 | fMRI | NO | EXPLICIT | VERBAL | VISUAL | HIGH | 16 | synonys>nonwords |
| | | | | | | | | | antonyms>nonwords |
| | | | | | | | | | English synonyma>nonwords |
| | | | | | | | | | Korean synonyms>nonwords |
| | | | | | | | | | honorific words>nonwords |
| Joubert et al. | 2004 | fMRI | NO | IMPLICIT | VERBAL | VISUAL | HIGH | 10 | low frequency words>nonwords |
| | | | | | | | | | high frequency words>consonant strings |
| | | | | | | | | | low frequency words>consonant strings |
| Kang et al. | 2006 | PET | NO | EXPLICIT | VERBAL | BOTH/AUDITORY/VISUAL | HIGH/LOW | 17 | audio-visual speech>noises and facial movements |
| | | | | | | | | | auditory speech>white noise |
| | | | | | | | | | visual speech>facial movements (chewing gum) |
| Khader, Jost, Mertens, Bien & Roesler | 2010 | fMRI | NO | EXPLICIT | VERBAL | VISUAL | HIGH | 16 | noun>rhyme generation |
| | | | | | | | | | verb>rhyme generation |
| | | | | | | | | | noun generation>letter detection |
| | | | | | | | | | verb generation>letter detection |
| Kim et al. | 2009 | fMRI | NO | EXPLICIT | VERBAL | VISUAL | HIGH | 36 | sentences>word lists |
| Kotz, Cappa, Von Cramon & Friederici | 2002 | fMRI | NO | EXPLICIT | VERBAL | AUDITORY | HIGH | 13 | words>pseudowords |
| Kuchinke et al. | 2005 | fMRI | YES | IMPLICIT | VERBAL | VISUAL | HIGH | 20 | emotion word>nonword |
| Kumar | 2016 | fMRI | NO | EXPLICIT | VERBAL | VISUAL | HIGH | 20 | abstract+concrete words>pseudowords |
| | | | | | | | | | abstract words>pseudowords |
| Kuperberg et al. | 2000 | fMRI | NO | EXPLICIT | VERBAL | AUDITORY | HIGH | 4 | sentences>words strings |
| Kyong, Scott, Rosen, Howe, Agnew & McGettigan | 2014 | fMRI | NO | IMPLICIT | VERBAL | AUDITORY | LOW | 19 | intelligible vocoded>inverted vocoded speech |
| Leff, Schofield, Stephan, Crinion, Friston & Price | 2008 | fMRI | NO | IMPLICIT | VERBAL | AUDITORY | HIGH | 26 | speech>reversed speech |

| | | | | | | | | | |
|---|---|---|---|---|---|---|---|---|---|
| Lin, Wang, Zhao, Liu, Li & Bi | 2015 | fMRI | NO | EXPLICIT | VERBAL | VISUAL | HIGH | 20 | words>pseudowords |
| Liu et al. | 2009 | fMRI | NO | EXPLICIT | VERBAL | VISUAL/AUDITORY | LOW | 16 | words meaning>slashes |
| | | | | | | | | | words rhyming>slashes |
| | | | | | | | | | words meaning>tones |
| | | | | | | | | | words rhyming>tones |
| Liuzzi et al. | 2017 | fMRI | NO | EXPLICIT | VERBAL | BOTH | HIGH | 18 | semantic judgement > input modality detection |
| Ludersdorfer, Wimmer, Richlan, Schurz, Hutzler & Kronbichler | 2016 | fMRI | NO | EXPLICIT | VERBAL | AUDITORY | LOW | 29 | words orthographic>tones |
| | | | | | | | | | words semantic>tones |
| Ludersdorfer, Schurz, Richlan, Kronbichler & Wimmer | 2013 | fMRI | NO | EXPLICIT | VERBAL | VISUAL/AUDITORY | HIGH | 29 | words>false fonts |
| | | | | | | | | | words>pseudowords |
| | | | | | | | | | speech>reversed speech |
| | | | | | | | | | words>pseudowords |
| Malins, Gumkowski, Buis, Molfese, Rueckl, Frost, Pugh, Morris & Mencl | 2016 | fMRI | NO | IMPLICIT | VERBAL | VISUAL | LOW | 18 | unrelated words>false font |
| | | | | | | | | | unrelated>pseudowords |
| Marques, Canessa & Cappa | 2009 | fMRI | NO | EXPLICIT | VERBAL | VISUAL | LOW | 21 | sentences>crosses |
| Marques, Canessa, Siri, Catricala & Cappa | 2008 | fMRI | NO | EXPLICIT | VERBAL | VISUAL | LOW | 21 | semantic features>baseline task |
| Mashal, Vishne, Laor & Titone | 2013 | fMRI | NO | EXPLICIT | VERBAL | VISUAL | HIGH | 14 | novel metaphor>unrelated words |
| | | | | | | | | | conventional metaphor>unrelated words |
| Matchin, Liao, gaston & Lau | 2019 | fMRI | NO | EXPLICIT | VERBAL | VISUAL | HIGH | 20 | verb phrase>list |
| | | | | | | | | | noun phrase>list |
| Matchin, Hammerly & Lau | 2017 | fMRI | NO | EXPLICIT | VERBAL | VISUAL | HIGH | 16 | sentences>word lists |
| | | | | | | | | | sentences>phrases |
| | | | | | | | | | real>pseudoword lists |
| | | | | | | | | | real>pseudoword phrases |

| | | | | | | | | | |
|---|---|---|---|---|---|---|---|---|---|
| | | | | | | | | | real>pseudoword sentences |
| Mellem, Jasmin, Peng & Martin | 2016 | fMRI | NO | IMPLICIT | VERBAL | VISUAL | LOW | 20 | longer>shorter phrase |
| Meyer, Alter, Friederici, Lohmann & Yves von Cramon | 2002 | fMRI | NO | IMPLICIT | VERBAL | AUDITORY | HIGH | 14 | word>pseudoword sentence |
| Moberget, Gullesen, Andersson, Ivry & Endestad | 2014 | fMRI | NO | EXPLICIT | VERBAL | VISUAL | HIGH | 32 | incongruent>scrambled sentence |
| | | | | | | | | | congruent>scrambled sentence |
| Moseley, Carota, Hauk, Mohr & Pulvermueller | 2012 | fMRI | YES | IMPLICIT | VERBAL | VISUAL | LOW | 18 | all emotional words>hashmarks |
| | | | | | | | | | abstract emotional words>hashmarks |
| | | | | | | | | | arm+face+emotion words>hashmarks |
| | | | | | | | | | face words>hashmarks |
| | | | | | | | | | arm words>hashmarks |
| Mummery, Patterson, Hodges & Price | 1998 | PET | NO | EXPLICIT | VERBAL | VISUAL | HIGH | 10 | semantic>phonological decision |
| Nakamura et al. | 2001 | PET | YES | EXPLICIT | VERBAL | AUDITORY | HIGH | 9 | familiar voice>vowel discrimination |
| | | | | | | | | | self voice>vowel discrimination |
| Nichelli, Grafman, Pietrini, Clark, Lee & Miletich | 1995 | PET | NO | EXPLICIT | VERBAL | VISUAL | HIGH | 9 | semantic>orthographic decision |
| Noppeney & Price | 2003 | PET | NO | EXPLICIT | VERBAL | AUDITORY | LOW | 9 | normal>reversed words |
| Orfanidou, Marlsen-Wilson & Davis | 2006 | fMRI | NO | EXPLICIT | VERBAL | AUDITORY | HIGH | 13 | words>pseudowords |
| Pallier, Devauchelle & Dehaene | 2011 | fMRI | NO | EXPLICIT | VERBAL | VISUAL | HIGH | 40 | longer>shorter phrase |
| | | | | | | | | | length of real>pseudoword sentences |
| Peelle, Eason, Schmitter, Schwarzbauer & Davis | 2010 | fMRI | NO | EXPLICIT | VERBAL | AUDITORY | HIGH | 6 | sentences>signal correlated noise |
| Perani, Schnur, Tettamanti, Gorno-Tempini, Cappa & Fazio | 1999 | PET | NO | EXPLICIT | VERBAL | VISUAL | HIGH | 8 | living words>pseudowords |
| | | | | | | | | | nonliving words>pseudowords |
| Perrone-Bertolotti, Kauffmann, Pichat, Vidal & Baciu | 2017 | fMRI | NO | EXPLICIT | VERBAL | VISUAL | HIGH | 24 | words>unreadable font |

| | | | | | | | | | |
|---|---|---|---|---|---|---|---|---|---|
| Pilgrim, Fadili, Fletcher & Tyler | 2002 | fMRI | NO | EXPLICIT | VERBAL | VISUAL | HIGH | 14 | words>letter strings |
| Price, Moore, Humphreys & Wise | 1997 | PET | NO | EXPLICIT | VERBAL | VISUAL | HIGH | 6 | semantic>phonological decision |
| Pulvermueller, Cook & Hauk | 2012 | fMRI | NO | IMPLICIT | VERBAL | VISUAL | LOW | 23 | phrases>hashmarks |
| | | | | | | | | | uninflected words>hashmarks |
| | | | | | | | | | inflected words>hashmarks |
| Raettig & Kotz | 2008 | fMRI | NO | EXPLICIT | VERBAL | AUDITORY | HIGH | 16 | real words>pseudowords |
| Raposo, Frade & Alves | 2016 | fMRI | NO | IMPLICIT | VERBAL | VISUAL | HIGH | 18 | semantic>perceptual decision |
| Raposo, Moss, Stamatakis & Tyler | 2009 | fMRI | NO | IMPLICIT | VERBAL | AUDITORY | LOW | 22 | action sentences>SC noise |
| Rapp & Lipka | 2011 | fMRI | NO | IMPLICIT | VERBAL | VISUAL | LOW | 10 | words>checkerboards |
| | | | | | | | | | words>letter strings |
| Redcay, Velnoskey & Rowe | 2016 | fMRI | NO | EXPLICIT | VERBAL | VISUAL | HIGH | 24 | real>pseudoword sentences |
| Rissman, Eliassen & Blumstein | 2003 | fMRI | NO | IMPLICIT | VERBAL | AUDITORY | HIGH | 15 | words>pseudowords |
| Robertson et al. | 2000 | fMRI | NO | IMPLICIT | VERBAL | VISUAL | HIGH | 8 | indefinite article sentence>letter strings |
| | | | | | | | | | definite article sentence>letter strings |
| Rodd, Johnsrude & Davis | 2012 | fMRI | NO | EXPLICIT | VERBAL | AUDITORY | LOW | 15 | speech>SCN |
| Rodd, Longe, Randall & Tyler | 2010 | fMRI | NO | EXPLICIT | VERBAL | AUDITORY | LOW | 14 | speech>SCN |
| Rogalsky & Hickok | 2009 | fMRI | NO | IMPLICIT | VERBAL | AUDITORY | HIGH | 14 | sentences>word lists |
| Rogalsky, Almeida, Sprouse & Hickok | 2015 | fMRI | NO | IMPLICIT | VERBAL | AUDITORY | HIGH | 15 | words>scrambled |
| Roskies, Fiez, Balota, Raichle & Petersen | 2001 | PET | NO | EXPLICIT | VERBAL | VISUAL | HIGH | 20 | semantic>phonological decision |
| Roxbury, McMahon & Copland | 2014 | fMRI | NO | EXPLICIT | VERBAL | AUDITORY | HIGH | 17 | concrete word>pseudoword |
| | | | | | | | | | abstract word>pseudoword |
| Ryan, Cox, Hayes & Nadel | 2008 | fMRI | NO | EXPLICIT | VERBAL | VISUAL | HIGH | 10 | semantic fluency (generate) > crosses |
| | | | | | | | | | semantic fluency (recall) > crosses |
| | | | | | | | | | semantic fluency (recall & geneerate)>crosses |

| | | | | | | | | | |
|---|---|---|---|---|---|---|---|---|---|
| Ryan, Lin, Ketcham & Nadel | 2010 | fMRI | NO | EXPLICIT | VERBAL | VISUAL | HIGH | 15 | semantic spatial old>letter judgement |
| | | | | | | | | | semantic spatial new>letter judgement |
| | | | | | | | | | semantic non spatial old>letter judgement |
| | | | | | | | | | semantic non spatial new>letter judgement |
| | | | | | | | | | semantic new>episodic judgement |
| Sabri, Binder, Desai, Medler, Leitl & Liebenthal | 2008 | fMRI | NO | EXPLICIT | VERBAL | AUDITORY | HIGH | 28 | speech>rotated speech |
| | | | | | | | | | words>pseudowords |
| Sachs, Weis, Krings, Huber & Kircher | 2008 | fMRI | NO | EXPLICIT | VERBAL | VISUAL | HIGH | 14 | biased thematic judgement>letters |
| | | | | | | | | | biased taxonomic judgement>letters |
| | | | | | | | | | balanced taxonomic judgement>letters |
| | | | | | | | | | balanced taxonomic judgement>letters |
| Saur et al. | 2008 | fMRI | NO | IMPLICIT | VERBAL | AUDITORY | HIGH | 33 | word>pseudoword sentences |
| Schell, Zaccarella & Friederici | 2017 | fMRI | NO | EXPLICIT | VERBAL | AUDITORY | HIGH | 21 | phrases>non combinatorial words |
| Schmitt, Auer & Ferstl | 2019 | fMRI | NO | EXPLICIT | VERBAL | AUDITORY | HIGH | 40 | known>unknown language |
| Schuil, Smits & Zwaan | 2013 | fMRI | NO | EXPLICIT | VERBAL | VISUAL | HIGH | 20 | senteces>pseudowords |
| | | | | | | | | | verbs>pseudowords |
| | | | | | | | | | literal sentences>pseudowords |
| | | | | | | | | | nonliteral sentences>pseudowords |
| Scott, Blank, Rosen & Wise | 2000 | PET | NO | IMPLICIT | VERBAL | AUDITORY | HIGH | 8 | intelligible>unintelligible speech |
| Segal & Petrides | 2012 | fMRI | NO | EXPLICIT | VERBAL | VISUAL | HIGH | 90 | words>pseudowords |
| Sergent, Otha & Macdonald | 1992 | PET | YES | EXPLICIT | VERBAL | VISUAL | HIGH | 7 | face identity>gender discrimination |
| Sheldon, McAndrews, Pruessner & Moscovitch | 2016 | fMRI | NO | EXPLICIT | VERBAL | VISUAL | HIGH | 15 | semantic fluency>perceptual task |
| Simard, Monetta, Nagano-Saito & Monchi | 2013 | fMRI | NO | EXPLICIT | VERBAL | VISUAL | HIGH | 14 | semantic>control macthing |
| | | | | | | | | | semantic matching>phonological decision (syllable rhyme) |
| | | | | | | | | | semantic matching>phonological decision (syllable onset) matching |

| | | | | | | | | | |
|---|---|---|---|---|---|---|---|---|---|
| Slioussar, Kireev, Chernigovskaya, Kataeva, Korotkov & Medvedev | 2014 | fMRI | NO | EXPLICIT | VERBAL | VISUAL | HIGH | 21 | real verbs>pseudowords |
| | | | | | | | | | real nouns>pseudowords |
| Smith, Myers, Sethi, Pantazatos, Yanagihara & Hirsch | 2012 | fMRI | NO | EXPLICIT | VERBAL | VISUAL | HIGH | 14 | semantics>baseline |
| Snijders, Vosse, Kempen, Van Berkum, Petersson & Hagoort | 2009 | fMRI | NO | IMPLICIT | VERBAL | VISUAL | HIGH | 28 | sentences>word lists |
| Stowe, Paans, Wijers, Zwarts, Mulder & Vaalburg | 1999 | PET | NO | IMPLICIT | VERBAL | VISUAL | HIGH | 12 | sentences>scrambled word lists |
| Straube, Green, Weis & Kircher | 2012 | fMRI | NO | IMPLICIT | VERBAL | AUDITORY | HIGH | 16 | known>unknown language |
| Stringaris, Medford, Giampietro. Brammer & David | 2007 | fMRI | NO | EXPLICIT | VERBAL | VISUAL | HIGH | 11 | literal>meaningless sentences |
| | | | | | | | | | methaphors>meaningless sentences |
| Sugiura et al. | 2006 | fMRI | YES | EXPLICIT | VERBAL | VISUAL | HIGH | 24 | famous>unfamiliar names |
| | | | | | | | | | personal>unfamiliar names |
| Sugiura et al. | 2008 | fMRI | YES | EXPLICIT | VERBAL | VISUAL | HIGH | 25 | low familiar>unfamiliar names |
| | | | | | | | | | personal familiar>unfamiliar names |
| | | | | | | | | | high familiar>unfamiliar names |
| | | | | | | | | | high familiar>unfamiliar names |
| Sun, Xue, Zhang, Zuo, Chen, Wang, Martin, Wang, Chen, He & Wang | 2017 | fMRI | NO | EXPLICIT | VERBAL | VISUAL | HIGH | 11 | semantic>orthography judgement |
| Szlachta, Bozic, Jelowicka & Marslen-Wilson | 2012 | fMRI | NO | IMPLICIT | VERBAL | VISUAL | LOW | 21 | words>musical rain |
| | | | | | | | | | nouns>musical rain |
| | | | | | | | | | inflected nouns>musical rain |
| Takeichi, Koyama, Terao, Takeuchi, Toyosawa & Murohashi | 2010 | fMRI | NO | IMPLICIT | VERBAL | AUDITORY | HIGH | 23 | speech>reversed speech |
| | | | | | | | | | speech>modulated speech |
| Thierry & Price | 2006 | PET | NO | EXPLICIT | VERBAL | AUDITORY | HIGH | 12 | auditory words>speech control(scrambled) |

| | | | | | | | | | |
|---|---|---|---|---|---|---|---|---|---|
| Thierry & Price | 2006 | PET | NO | EXPLICIT | VERBAL | VISUAL | HIGH | 12 | visual words>text control (scrambled letter strings) |
| Tieleman, Seurinck, Deblaere, Vandemaele, Vingerhoets & Achten | 2005 | fMRI | NO | EXPLICIT | VERBAL | VISUAL | HIGH | 22 | self-paced semantic>perceptual decision |
| | | | | | | | | | fixed-paced semantic>perceptual decision |
| Tyler, Stamatakis, Dick, Bright, Fletcher & Moss | 2003 | fMRI | NO | EXPLICIT | VERBAL | VISUAL | HIGH | 12 | animals>baseline |
| | | | | | | | | | tool action words>baseline |
| | | | | | | | | | biological action>baseline |
| Vagharchakian, Dehaene-Lambertz, Pallier & Dehaene | 2012 | fMRI | NO | EXPLICIT | VERBAL | BOTH/AUDITORY | HIGH | 16 | intelligible>unintelligble compression rate |
| | | | | | | | | | intelligible>unintelligble compression rate |
| | | | | | | | | | intelligible>unintelligble compression rate |
| Van Ettinger-Veenstra, McAllister, Lundberg, Karlsson & Engstrom | 2016 | fMRI | NO | EXPLICIT | VERBAL | VISUAL | HIGH | 27 | sentences>symbol strings |
| Vignali, Hawelka, Hutzler & Richlan | 2019 | fMRI | NO | EXPLICIT | VERBAL | VISUAL | HIGH | 21 | foveal & parafovealwords>foveal & parafoveal pseudowords |
| Visser, Jefferies, Embleton & Lambon Ralph | 2012 | fMRI | NO | EXPLICIT | VERBAL | VISUAL | HIGH | 15 | words>baseline |
| Vitello, Warren, Devlin & Rodd | 2014 | fMRI | NO | EXPLICIT | VERBAL | AUDITORY | LOW | 20 | sentences>SCN |
| von Kriegstein, Eger, Kleinschmidt & Giraud | 2003 | fMRI | NO | EXPLICIT | VERBAL | AUDITORY | HIGH | 14 | sentence>speech envelope |
| Wang, Zhao, Zevin & Yang | 2016 | fMRI | NO | EXPLICIT | VERBAL | VISUAL | HIGH | 16 | words>nonsense strokes |
| Weiss, Katzir & Bitan | 2015 | fMRI | NO | IMPLICIT | VERBAL | VISUAL | LOW | 18 | pointed words>asterisks |
| | | | | | | | | | unpointed words>asterisks |
| Welcome & Joanisse | 2012 | fMRI | NO | EXPLICIT | VERBAL | VISUAL | HIGH | 20 | semantic>phonological/orthographic decision |
| Wende, Straube, Stratmann, Sommer, Kircher & Nagels | 2012 | fMRI | NO | EXPLICIT | VERBAL | VISUAL | HIGH | 18 | semantic>phonological fluency |
| Wirth, Jann, Dierks, Federspiel, Wiest & Horn | 2011 | fMRI | NO | EXPLICIT | VERBAL | VISUAL | HIGH | 19 | semantic>phonological & perceptual decision |
| Wright et al. | 2008 | fMRI | NO | EXPLICIT | VERBAL | VISUAL | HIGH | 10 | semantic>perceptual matching |

| | | | | | | | | | |
|---|---|---|---|---|---|---|---|---|---|
| Wright, Randall, Marslen-Wilson & Tyler | 2011 | fMRI | NO | BOTH | VERBAL | AUDITORY | LOW | 14 | speech>musical rain |
| Wu, Mai, Tang, Ge, Luo & Liu | 2013 | fMRI | NO | IMPLICIT | VERBAL | VISUAL | LOW | 19 | arm words>checkerboard |
| | | | | | | | | | leg words>checkerboard |
| | | | | | | | | | mouth words>checkerboard |
| Xiao et al. | 2005 | fMRI | NO | EXPLICIT | VERBAL | AUDITORY | HIGH | 14 | words>pseudowords |
| Yang, Li, Fang, Shu, Liu & Chen | 2016 | fMRI | NO | IMPLICIT | VERBAL | VISUAL | LOW | 20 | opaque idioms>hashmarks |
| | | | | | | | | | transparent idioms>hashmarks |
| | | | | | | | | | literal phrases>hashmarks |
| Zaccarella & Friederici | 2015 | fMRI | NO | EXPLICIT | VERBAL | VISUAL | HIGH | 22 | words>pseudowords |
| Zhang, Liu & Zhang | 2014 | fMRI | NO | EXPLICIT | VERBAL | VISUAL | LOW | 18 | nonliving words>asterisks |
| | | | | | | | | | living words>asterisks |
| Zhang, Xiao & Weng | 2012 | fMRI | NO | EXPLICIT | VERBAL | VISUAL | HIGH | 14 | words>pseudowords |
| Zhuang & Devereux | 2017 | fMRI | NO | EXPLICIT | VERBAL | AUDITORY | HIGH | 16 | phrases>words |
| Zou, Packard, Xia, Liu & Shu | 2016 | fMRI | NO | IMPLICIT | VERBAL | AUDITORY | LOW | 17 | speech>tone |
| | | | | | | | | | speech>tone |
| | | | | | | | | | speech>tone |
| | | | | | | | | | identical speech>tone |
| | | | | | | | | | words>tone |
| EXCLUDED CONTRASTS CONTAINING BOTH VERBAL AND NON VERBAL STIMULI. THE EXLUDED NON VERBAL EXPERIMENTS CAN BE SEEN IN Table M.2.2 SC NON-VERBAL | | | | | | | | | |
| Brambati, Benoit, Monetta, Belleville & Joubert | 2010 | fMRI | YES | EXPLICIT | BOTH | BOTH | HIGH | 12 | general & specific occupation judgement>baseline (scrambled face) |
| Bruffaerts, Dupont, Peeters, De Deyne, Storms & Vandenberghe | 2013 | fMRI | NO | EXPLICIT | BOTH | VISUAL | HIGH | 19 | real > scrambled pictures & words |
| Ebisch et al. | 2007 | fMRI | NO | EXPLICIT | BOTH | VISUAL | HIGH | 17 | functional&visuospatial pictures&words>meaningless drawings&pseudowords |
| Europa, Gitelman, Kiran & Thompson | 2019 | fMRI | NO | EXPLICIT | BOTH | BOTH | HIGH | 21 | sentences>baseline (reversed sentences) |

| | | | | | | | | | |
|---|---|---|---|---|---|---|---|---|---|
| Giraud & Price | 2001 | PET | NO | BOTH | BOTH | AUDITORY | BOTH | 12 | words+environmental sounds>syllables+noise |
| Holle, Gunter, Rueschemeyer, Hennenlotter & Iacoboni | 2008 | fMRI | NO | EXPLICIT | BOTH | BOTH | HIGH | 17 | iconic gesture of dominant meaning>grooming |
| | | fMRI | NO | EXPLICIT | BOTH | BOTH | HIGH | 17 | iconic gesture of subordinate meaning>grooming |
| Kinno, Kawamura, Shioda & Sakai | 2008 | fMRI | NO | EXPLICIT | BOTH | VISUAL | HIGH | 14 | canonical sentence>picture & letter strings |
| | | fMRI | NO | EXPLICIT | BOTH | VISUAL | HIGH | 14 | active sentence>picture & letter strings |
| | | fMRI | NO | EXPLICIT | BOTH | VISUAL | HIGH | 14 | passive sentence>picture & letter strings |
| Nielson et al. | 2010 | fMRI | YES | EXPLICIT | BOTH | VISUAL | HIGH | 17 | familiar>unfamiliar people |
| Redcay, Velnoskey & Rowe | 2016 | fMRI | NO | EXPLICIT | BOTH | VISUAL | HIGH | 24 | meaningful>meaningless stimuli |
| Rogers et al. | 2006 | PET | NO | EXPLICIT | BOTH | VISUAL | HIGH | 12 | pictures>scrambled pictures |
| Seghier, Josse, Leff & Price | 2011 | fMRI | NO | EXPLICIT | BOTH | VISUAL | HIGH | 60 | meaningful>meaningless stimuli |
| van Leeuwen et al. | 2014 | fMRI | NO | EXPLICIT | BOTH | BOTH | HIGH | 16 | speech>reversed speech |
| Visser, Jefferies, Embleton & Lambon Ralph | 2012 | fMRI | NO | EXPLICIT | BOTH | VISUAL | HIGH | 15 | semantics>baseline |
| Wright et al. | 2008 | fMRI | NO | EXPLICIT | BOTH | VISUAL | HIGH | 10 | semantic>perceptual matching |
| Zvyagintsev, Clemens, Chechko, Mathiak, Sack & Mathiak | 2013 | fMRI | NO | IMPLICIT | NONE | NONE | HIGH | 15 | visual imagery>counting |
| | | fMRI | NO | IMPLICIT | NONE | NONE | HIGH | 15 | auditory imagery>counting |

**The references to access the listed studies are listed at the end of the document*

***Table M2.2 SC NON-VERBAL*** *List of studies included in the* **semantic cognition NON-VERBAL STIMULUS DOMAIN** *(N = 37) meta-analysis after excluding VERBAL contrasts.* ***Note:*** *The list of excluded studies with VERBAL STIMULUS DOMAIN and contrasts with BOTH VEBAL and NON-VERBAL stimuli that were also excluded can be seen in Table M.2.1 SC VERBAL. SC= Semantic Cognition, N= Sample Size*

| *Authors | Year | Imaging Method | Social Content | Instructional Cue | Stimulus Domain | Sensory Input Modality | Baseline Type | N | Contrast |
|---|---|---|---|---|---|---|---|---|---|
| Baumgaertner et al. | 2007 | fMRI | NO | IMPLICIT | NONVERBAL | VISUAL | HIGH | 19 | videos>scrambled videos |
| Chiao, Harada, Oby, Li, Parrish & Bridge | 2009 | fMRI | YES | EXPLICIT | NONVERBAL | VISUAL | LOW | 12 | uniform status judgement>colour change detection |
| | | | | | | | | | face status judgement>colour change detection |
| | | | | | | | | | car status judgement>colour change detection |
| Chouinard, Morrissey, Kohler & Goodale | 2008 | fMRI | NO | EXPLICIT | NONVERBAL | VISUAL | HIGH | 14 | objects>scrambled objects |
| Damasio, Grabowski, Tranel, Ponto, Hichwa & Damasio | 2001 | PET | NO | EXPLICIT | NONVERBAL | VISUAL | LOW | 10 | actions without implement>control task |
| | | | | | | | | | actions with implement>control task |
| Damasio, Tranel, Grabowski, Adolphs & Damasio | 2004 | PET | YES | EXPLICIT | NONVERBAL | VISUAL | HIGH | 55 | persons>face orientation judgement |
| | | | | | | | | | animals>scrambled pictures |
| | | | | | | | | | tools>scrambled pictures |
| Elfgren, Westen, Passant, Larsson, Mannfolk & Fransson | 2006 | fMRI | YES | EXPLICIT | NONVERBAL | VISUAL | HIGH | 15 | familiar>unfamiliar faces (identification) |
| Emmorey, Xu, Gannon, Goldin-Meadow & Braun | 2010 | fMRI | NO | IMPLICIT | NONVERBAL | VISUAL | HIGH | 14 | meaningful pantomimes>unknown sign language |
| Engelien et al. | 2006 | PET | NO | IMPLICIT | NONVERBAL | AUDITORY | HIGH | 6 | meaningful>meaningless sounds |
| Garn, Allen & Larsen | 2009 | fMRI | NO | EXPLICIT | NONVERBAL | VISUAL | HIGH | 26 | pictures>scrambled pictures |
| | | | | | | | | | plants>scrambled pictures |
| | | | | | | | | | tools>scrambled pictures |
| Gerlach, Law, Gade & Paulson | 1999 | PET | NO | EXPLICIT | NONVERBAL | VISUAL | HIGH | 15 | object decision>pattern discrimination |
| Gesierich et al. | 2012 | fMRI | YES | EXPLICIT | NONVERBAL | VISUAL | HIGH | 21 | familiar>scrambled faces |

| | | | | | | | | | |
|---|---|---|---|---|---|---|---|---|---|
| | | | | | | | | | familiar > unfamiliar faces |
| | | | | | | | | | familiar>unfamiliar faces |
| Gorno-Tempini et al. | 1998 | PET | YES | EXPLICIT | NONVERBAL | VISUAL | HIGH | 6 | famous faces>controls |
| | | | | | | | | | non-famous faces>controls |
| Grabowski, Damasio, Tranel, Boles Ponto, Hichwa & Damasio | 2001 | PET | YES | EXPLICIT | NONVERBAL | VISUAL | HIGH | 10 | naming persons>building orientation judgement |
| | | | | | | | | | naming landmarks>face orientation judgement |
| | | | | | | | | | naming persons>face orientation judgement |
| | | | | | | | | | naming unique entities>baseline |
| Groussard et al. | 2010 | PET | NO | EXPLICIT | NONVERBAL | AUDITORY | HIGH | 11 | musical semantic>musical reference |
| Haberling, Corballis &Corballis | 2016 | fMRI | NO | EXPLICIT | NONVERBAL | VISUAL | HIGH | 92 | meaningful pantomimes>unknown sign language |
| | | | | | | | | | meaningful pantomimes>dog videos |
| Harrington, Farias & Davis | 2009 | fMRI | NO | EXPLICIT | NONVERBAL | VISUAL | HIGH | 8 | familiar>non objects |
| Hocking, McMahon & de Zubicaray | 2011 | fMRI | NO | EXPLICIT | NONVERBAL | AUDITORY | HIGH | 13 | all environmental sounds>perceptual baseline |
| Husain, Patkin, Kim, Braun & Horwitz | 2012 | fMRI | NO | EXPLICIT | NONVERBAL | VISUAL | HIGH | 16 | meaningful iconic>meaningless gestures |
| Leung & Alain | 2011 | fMRI | NO | EXPLICIT | NONVERBAL | AUDITORY | HIGH | 16 | semantic>location matching |
| Leveroni et al. | 2000 | fMRI | YES | EXPLICIT | NONVERBAL | VISUAL | HIGH | 11 | familiar faces>foils(never seen faces) |
| | | | | | | | | | newly learned faces>foils(never seen faces) |
| Menz, Blangero, Kunze & Binkofski | 2010 | fMRI | NO | EXPLICIT | NONVERBAL | VISUAL | HIGH | 20 | known>unknown objects |
| Nakamura et al. | 2000 | PET | YES | EXPLICIT | NONVERBAL | VISUAL | LOW | 7 | familiar faces>fixation cross |
| Perani, Schnur, Tettamanti, Gorno-Tempini, Cappa & Fazio | 1999 | PET | NO | EXPLICIT | NONVERBAL | VISUAL | HIGH | 11 | living objects>shapes |
| | | | | | | | | | nonliving objects>shapes |
| Redcay, Velnoskey & Rowe | 2016 | fMRI | NO | EXPLICIT | NONVERBAL | VISUAL | HIGH | 24 | communicative>non-communicative gesture |
| Rogers et al. | 2006 | PET | NO | EXPLICIT | NONVERBAL | VISUAL | HIGH | 12 | specific-level judgement>baseline |
| Ross & Olson | 2012 | fMRI | YES | EXPLICIT | NONVERBAL | VISUAL | HIGH | 11 | famous>unknown faces & landmarks |
| Segal & Petrides | 2012 | fMRI | NO | EXPLICIT | NONVERBAL | VISUAL | HIGH | 90 | writing>copying |
| Sergent, Otha & Macdonald | 1992 | PET | YES | EXPLICIT | NONVERBAL | VISUAL | LOW | 7 | object recognition>gratings |

| | | | | | | | | | |
|---|---|---|---|---|---|---|---|---|---|
| Straube, Green, Weis & Kircher | 2012 | fMRI | NO | IMPLICIT | NONVERBAL | VISUAL | HIGH | 16 | iconic>meaningless gesture |
| Sugiura et al. | 2001 | PET | YES | EXPLICIT | NONVERBAL | VISUAL | HIGH | 5 | identity discrimination>control |
| | | | | | | | | | identity discrimination>face direction |
| Taminato, Miura, Sugiura & Kawashima | 2014 | fMRI | NO | EXPLICIT | NONVERBAL | VISUAL | LOW | 35 | object recognition>control task |
| | | | | | | | | | object recognition>control task |
| Taylor, Arsalidou, Bayless, Morris, Evans & Barbeau | 2009 | fMRI | YES | IMPLICIT | NONVERBAL | VISUAL | HIGH | 10 | own>unfamiliar face |
| | | | | | | | | | partner's>unfamiliar face |
| | | | | | | | | | parent's>unfamiliar face |
| Thierry & Price | 2006 | PET | NO | EXPLICIT | NONVERBAL | AUDITORY | HIGH | 12 | auditory sounds>sound control (scrambled) |
| Thierry & Price | 2006 | PET | NO | EXPLICIT | NONVERBAL | VISUAL | HIGH | 12 | visual videos>video control (distorted) |
| Vingerhoets | 2008 | fMRI | NO | IMPLICIT | NONVERBAL | VISUAL | HIGH | 14 | familiar>unfamiliar tools |
| Visser, Jefferies, Embleton & Lambon Ralph | 2012 | fMRI | NO | EXPLICIT | NONVERBAL | VISUAL | HIGH | 15 | pictures>baseline |
| Wright et al. | 2008 | fMRI | NO | EXPLICIT | NONVERBAL | VISUAL | HIGH | 10 | semantic>perceptual matching |
| EXCLUDED CONTRASTS CONTAINING BOTH VERBAL AND NON VERBAL STIMULI. THE EXLUDED VERBAL EXPERIMENTS CAN BE SEEN IN Table M.2.1 SC VERBAL | | | | | | | | | |
| Brambati, Benoit, Monetta, Belleville & Joubert | 2010 | fMRI | YES | EXPLICIT | BOTH | BOTH | HIGH | 12 | general & specific occupation judgement>baseline (scrambled face) |
| Bruffaerts, Dupont, Peeters, De Deyne, Storms & Vandenberghe | 2013 | fMRI | NO | EXPLICIT | BOTH | VISUAL | HIGH | 19 | real > scrambled pictures & words |
| Ebisch et al. | 2007 | fMRI | NO | EXPLICIT | BOTH | VISUAL | HIGH | 17 | functional&visuospatial pictures&words>meaningless drawings&pseudowords |
| Europa, Gitelman, Kiran & Thompson | 2019 | fMRI | NO | EXPLICIT | BOTH | BOTH | HIGH | 21 | sentences>baseline (reversed sentences) |
| Giraud & Price | 2001 | PET | NO | BOTH | BOTH | AUDITORY | BOTH | 12 | words+environmental sounds>syllables+noise |
| Holle, Gunter, Rueschemeyer, Hennenlotter & Iacoboni | 2008 | fMRI | NO | EXPLICIT | BOTH | BOTH | HIGH | 17 | iconic gesture of dominant meaning>grooming |
| | | fMRI | NO | EXPLICIT | BOTH | BOTH | HIGH | 17 | iconic gesture of subordinate meaning>grooming |
| Kinno, Kawamura, Shioda & Sakai | 2008 | fMRI | NO | EXPLICIT | BOTH | VISUAL | HIGH | 14 | canonical sentence>picture & letter strings |

| | | | | | | | | | |
|---|---|---|---|---|---|---|---|---|---|
| | | fMRI | NO | EXPLICIT | BOTH | VISUAL | HIGH | 14 | active sentence>picture & letter strings |
| | | fMRI | NO | EXPLICIT | BOTH | VISUAL | HIGH | 14 | passive sentence>picture & letter strings |
| Nielson et al. | 2010 | fMRI | YES | EXPLICIT | BOTH | VISUAL | HIGH | 17 | familiar>unfamiliar people |
| Redcay, Velnoskey & Rowe | 2016 | fMRI | NO | EXPLICIT | BOTH | VISUAL | HIGH | 24 | meaningful>meaningless stimuli |
| Rogers et al. | 2006 | PET | NO | EXPLICIT | BOTH | VISUAL | HIGH | 12 | pictures>scrambled pictures |
| Seghier, Josse, Leff & Price | 2011 | fMRI | NO | EXPLICIT | BOTH | VISUAL | HIGH | 60 | meaningful>meaningless stimuli |
| van Leeuwen et al. | 2014 | fMRI | NO | EXPLICIT | BOTH | BOTH | HIGH | 16 | speech>reversed speech |
| Visser, Jefferies, Embleton & Lambon Ralph | 2012 | fMRI | NO | EXPLICIT | BOTH | VISUAL | HIGH | 15 | semantics>baseline |
| Wright et al. | 2008 | fMRI | NO | EXPLICIT | BOTH | VISUAL | HIGH | 10 | semantic>perceptual matching |
| Zvyagintsev, Clemens, Chechko, Mathiak, Sack & Mathiak | 2013 | fMRI | NO | IMPLICIT | NONE | NONE | HIGH | 15 | visual imagery>counting |
| | | fMRI | NO | IMPLICIT | NONE | NONE | HIGH | 15 | auditory imagery>counting |

*****The references to access the listed studies are listed at the end of the document*

***Table M2.3 SC VISUAL*** *List of studies included in the* ***semantic cognition VISUAL INPUT MODALITY*** *(N = 152) meta-analysis after excluding AUDITORY contrasts.* ***Note:*** *The list of excluded studies with AUDITORY INPUT MODALITY can be seen in Table M.2.4 SC AUDITORY. Contrasts with BOTH VISUAL and AUDITORY stimuli were also excluded and are listed at the end of the document highlighted in grey. SC= Semantic Cognition, N= Sample Size*

| *Authors | Year | Imaging Method | Social Content | Instructional Cue | Stimulus Domain | Sensory Input Modality | Baseline Type | N | Contrast |
|---|---|---|---|---|---|---|---|---|---|
| AbdulSabur et al. | 2014 | fMRI | NO | EXPLICIT | VERBAL | VISUAL | LOW | 18 | narrative production>recitation |
| Abraham et al. | 2012 | fMRI | NO | EXPLICIT | VERBAL | VISUAL | HIGH | 19 | high&low divergent thinking (semantic)>1&2 back letter identity (working memory) |
| Assadollahi, Meinzer, Flaisch, Obleser & Rockstroh | 2009 | fMRI | NO | IMPLICIT | VERBAL | VISUAL | HIGH | 20 | nouns followed by 1&3 argument verbs>letter strings followed by 1&3argument verbs |
| Axmacher, Bialleck, Weber, Helmstaedter, Elger & Fell | 2009 | fMRI | NO | EXPLICIT | VERBAL | VISUAL | HIGH | 32 | word decision>spatial decision |
| Bagga et al. | 2013 | fMRI | NO | EXPLICIT | VERBAL | VISUAL | HIGH | 18 | semantic>case matching judgement |
| Barros-Loscertales et al. | 2012 | fMRI | NO | EXPLICIT | VERBAL | VISUAL | LOW | 59 | control words>hashmarks baseline |
| Baumgaertener, Weiller & Buchel | 2002 | fMRI | NO | EXPLICIT | VERBAL | VISUAL | HIGH | 9 | word>pseudoword in sentence |
| Baumgaertner et al. | 2007 | fMRI | NO | IMPLICIT | NONVERBAL | VISUAL | HIGH | 19 | videos>scrambled videos |
| Bick, Goelman & Frost | 2008 | fMRI | NO | EXPLICIT | VERBAL | VISUAL | LOW | 14 | semantic>visual control |
| | | | | | | | | | morphologiacl>visual control |
| | | | | | | | | | orthographic>visual control |
| | | | | | | | | | phonological>visual control |
| Binder et al. | 2003 | fMRI | NO | EXPLICIT | VERBAL | VISUAL | HIGH | 24 | word>nonword |
| Birn et al. | 2010 | fMRI | NO | EXPLICIT | VERBAL | VISUAL | LOW/HIGH | 14 | category & letter fluency>months ( automatic speech) |
| | | | | | | | | | category>letter fluency |
| Bonhage, Fiebach, Bahlmann & Mueller | 2014 | fMRI | NO | EXPLICIT | VERBAL | VISUAL | HIGH | 18 | sentence fragments>ungrammatical word strings |
| Bonhage, Mueller, Friederici & Fiebach | 2015 | fMRI | NO | EXPLICIT | VERBAL | VISUAL | HIGH | 18 | sentences>jabberwocky sentences |
| Booth et al. | 2006 | fMRI | NO | EXPLICIT | VERBAL | VISUAL | HIGH | 13 | meaning (semantic) judgement>rhyming( phonological) judgement |

| | | | | | | | | | |
|---|---|---|---|---|---|---|---|---|---|
| | | | | | | | | | meaning(semantic)>control(symbols) |
| Boulenger, Hauk & Pulvermuller | 2009 | fMRI | NO | EXPLICIT | VERBAL | VISUAL | LOW | 18 | sentences> hashmarks |
| | | | | | | | | | sentences> hashmarks |
| Bruffaerts, Dupont, Peeters, De Deyne, Storms & Vandenberghe | 2013 | fMRI | NO | EXPLICIT | BOTH | VISUAL | HIGH | 19 | real > scrambled pictures & words |
| Bulut, Hung, Tzeng & Wu | 2017 | fMRI | NO | IMPLICIT | VERBAL | VISUAL | HIGH | 20 | sentences>unstructured word lists |
| | | | | | | | | | sentences>unstructured character list |
| Cao, Peng, Liu, Jin, Fan, Deng, & Booth | 2009 | fMRI | NO | EXPLICIT | VERBAL | VISUAL | LOW | 13 | meaning>perceptual decision |
| Cappa, Perani, Schnur, Tettamanti & Fazio | 1998 | PET | NO | EXPLICIT | VERBAL | VISUAL | HIGH | 13 | words>pseudowords |
| | | | | | | | | | animal visual knowledge>pseudowords |
| | | | | | | | | | tool visual knowledge>pseudowords |
| | | | | | | | | | animal associative knowledge>pseudowords |
| | | | | | | | | | tools functional knowledge>pseudowords |
| Carota, Kriegeskorte, Nili & Pulvermuller | 2017 | fMRI | NO | IMPLICIT | VERBAL | VISUAL | LOW | 23 | words>hashmarks |
| Carota, Moseley & Pulvermueller | 2012 | fMRI | NO | IMPLICIT | VERBAL | VISUAL | LOW | 18 | all words>hashmarks |
| | | | | | | | | | tool words>hashmarks |
| | | | | | | | | | animal words>hashmarks |
| | | | | | | | | | food words>hashmarks |
| Chan, Tang, Tang, Lee, Lo & Kwong | 2009 | fMRI | NO | IMPLICIT | VERBAL | VISUAL | HIGH | 22 | synonyms>pseudocharacters |
| | | | | | | | | | synonyms>Korean characters (unknown) |
| Chiao, Harada, Oby, Li, Parrish & Bridge | 2009 | fMRI | YES | EXPLICIT | NONVERBAL | VISUAL | LOW | 12 | uniform status judgement>colour change detection |
| | | | | | | | | | face status judgement>colour change detection |
| | | | | | | | | | car status judgement>colour change detection |
| Chou, Chen, Wu & Booth | 2009 | fMRI | NO | EXPLICIT | VERBAL | VISUAL | HIGH | 31 | related words>false font |
| | | | | | | | | | unrelated words>false font |
| Chou, Chen, Wu & Booth | 2009 | fMRI | NO | EXPLICIT | VERBAL | VISUAL | HIGH | 32 | related words>false font |
| | | | | | | | | | unrelated words>false font |

| | | | | | | | | | |
|---|---|---|---|---|---|---|---|---|---|
| Chouinard, Morrissey, Kohler & Goodale | 2008 | fMRI | NO | EXPLICIT | NONVERBAL | VISUAL | HIGH | 14 | objects>scrambled objects |
| Chow, Kaup, Raabe & Greenlee | 2008 | fMRI | NO | EXPLICIT | VERBAL | VISUAL | HIGH | 15 | predictive&normal reading>pseudoword reading |
| | | | | | | | | | normal reading>pseudoword reading |
| | | | | | | | | | predictive reading>pseudoword reading |
| Damasio, Grabowski, Tranel, Ponto, Hichwa & Damasio | 2001 | PET | NO | EXPLICIT | NONVERBAL | VISUAL | LOW | 10 | actions without implement>control task |
| | | | | | | | | | actions with implement>control task |
| Damasio, Tranel, Grabowski, Adolphs & Damasio | 2004 | PET | YES | EXPLICIT | NONVERBAL | VISUAL | HIGH | 55 | persons>face orientation judgement |
| | | | | | | | | | animals>scrambled pictures |
| | | | | | | | | | tools>scrambled pictures |
| Davis, Meunier & Marslen-Wilson | 2004 | fMRI | NO | EXPLICIT | VERBAL | VISUAL | HIGH | 11 | words>letter strings |
| Devlin, Matthews & Rushworth | 2003 | fMRI | NO | EXPLICIT | VERBAL | VISUAL | HIGH | 12 | semantic>phonological judgement |
| Devlin et al. | 2002 | PET | NO | EXPLICIT | VERBAL | VISUAL | HIGH | 12 | all semantic>letter detection |
| | | | | | | | | | all semantic>letter detection |
| Devlin et al. | 2002 | PET | NO | EXPLICIT | VERBAL | VISUAL | HIGH | 8 | all semantic>letter categorisation |
| Devlin et al. | 2002 | fMRI | NO | EXPLICIT | VERBAL | VISUAL | HIGH | 8 | all semantic> letter categorisation |
| Devlin et al. | 2000 | PET | NO | EXPLICIT | VERBAL | VISUAL | HIGH | 8 | semantic categorisation>letter categorisation |
| Devlin et al. | 2000 | fMRI | NO | EXPLICIT | VERBAL | VISUAL | HIGH | 8 | semantic categorisation>letter categorisation |
| Diaz & McCarthy | 2009 | fMRI | NO | IMPLICIT | VERBAL | VISUAL | HIGH | 16 | all words>nonwords |
| Dreyer & Pulvermueller | 2018 | fMRI | YES | IMPLICIT | VERBAL | VISUAL | LOW | 28 | all nouns>hashmarks |
| | | | | | | | | | abstract emotional nouns>baseline (hashmarks) |
| | | | | | | | | | abstract mental nouns>baseline (hashmarks) |
| | | | | | | | | | food nouns>baseline (hashmarks) |
| | | | | | | | | | tool nouns>baseline (hashmarks) |
| Ebisch et al. | 2007 | fMRI | NO | EXPLICIT | BOTH | VISUAL | HIGH | 17 | functional&visuospatial pictures&words>meaningless drawings&pseudowords |

| | | | | | | | | | |
|---|---|---|---|---|---|---|---|---|---|
| Elfgren, Westen, Passant, Larsson, Mannfolk & Fransson | 2006 | fMRI | YES | EXPLICIT | NONVERBAL | VISUAL | HIGH | 15 | familiar>unfamiliar faces (identification) |
| Emmorey, Weisberg, McCullough & Petrich | 2013 | fMRI | NO | EXPLICIT | VERBAL | VISUAL | HIGH | 14 | words(semantic judgement)>false fonts |
| | | | | | | | | | words(phonological)>false fonts |
| Emmorey, Xu, Gannon, Goldin-Meadow & Braun | 2010 | fMRI | NO | IMPLICIT | NONVERBAL | VISUAL | HIGH | 14 | meaningful pantomimes>unknown sign language |
| Foki, Gartus, Geissler & Beisteiner | 2008 | fMRI | | EXPLICIT | VERBAL | VISUAL | LOW | 23 | semantic judgement>tongue movements |
| Friese, Rutschmann, Raabe & Schmalhofer | 2008 | fMRI | NO | EXPLICIT | VERBAL | VISUAL | HIGH | 13 | words>pseudowords |
| Garbin, Collina & Tabossi | 2012 | fMRI | NO | EXPLICIT | VERBAL | VISUAL | HIGH | 12 | object noun>pseudoword |
| | | | | | | | | | event noun>pseudoword |
| | | | | | | | | | verb>pseudoword |
| Garn, Allen & Larsen | 2009 | fMRI | NO | EXPLICIT | NONVERBAL | VISUAL | HIGH | 26 | pictures>scrambled pictures |
| | | | | | | | | | plants>scrambled pictures |
| | | | | | | | | | tools>scrambled pictures |
| Geranmayeh, Brownsett, Leech, Beckmann, Woodhead & Wise | 2012 | fMRI | NO | EXPLICIT | VERBAL | VISUAL | HIGH | 19 | speech>tongue movements |
| Gerlach, Law, Gade & Paulson | 1999 | PET | NO | EXPLICIT | NONVERBAL | VISUAL | HIGH | 15 | object decision>pattern discrimination |
| Gesierich et al. | 2012 | fMRI | YES | EXPLICIT | NONVERBAL | VISUAL | HIGH | 21 | familiar>scrambled faces |
| | | | | | | | | | familiar > unfamiliar faces |
| | | | | | | | | | familiar>unfamiliar faces |
| Gitelman, Nobre, Sonty, Parrish & Mesulam | 2005 | fMRI | NO | EXPLICIT | VERBAL | VISUAL | HIGH | 14 | semantic>control task |
| Gorno-Tempini et al. | 1998 | PET | YES | EXPLICIT | NONVERBAL/ VERBAL | VISUAL | HIGH | 6 | famous faces>controls |
| | | | | | | | | | famous names>controls |
| | | | | | | | | | double famous proper names>controls |
| | | | | | | | | | non-famous faces>controls |
| | | | | | | | | | Non-famous names>controls |
| | | | | | | | | | Double common names>controls |

| | | | | | | | | | |
|---|---|---|---|---|---|---|---|---|---|
| Grabowski, Damasio, Tranel, Boles Ponto, Hichwa & Damasio | 2001 | PET | YES | EXPLICIT | NONVERBAL | VISUAL | HIGH | 10 | naming persons>building orientation judgement |
| | | | | | | | | | naming landmarks>face orientation judgement |
| | | | | | | | | | naming persons>face orientation judgement |
| | | | | | | | | | naming unique entities>baseline |
| Graves, Binder, Desai, Conant & Seidenberg | 2010 | fMRI | NO | EXPLICIT | VERBAL | VISUAL | HIGH | 23 | forward>reverse phrases |
| Graves, Binder, Desai, Conant & Seidenberg | 2010 | fMRI | NO | EXPLICIT | VERBAL | VISUAL | HIGH | 22 | forward>reverse phrases |
| Grindrod, Garnett, Malyutina & den Ouden | 2014 | fMRI | NO | EXPLICIT | VERBAL | VISUAL | HIGH | 23 | words>nonwords |
| Grossman et al. | 2002a | fMRI | NO | IMPLICIT | VERBAL | VISUAL | HIGH | 16 | all nouns > pseudowords |
| | | | | | | | | | implements>pseudowords |
| | | | | | | | | | animals>pseudowords |
| | | | | | | | | | abstract>pseudowords |
| Grossman et al., 2002b | 2002 b | fMRI | NO | IMPLICIT | VERBAL | VISUAL | HIGH | 16 | verbs>pseudowords |
| Groussard et al. | 2010 | PET | NO | EXPLICIT | VERBAL | VISUAL | HIGH | 11 | verbal semantics>verbal reference |
| Haberling, Corballis &Corballis | 2016 | fMRI | NO | EXPLICIT | NONVERBAL/ VERBAL | VISUAL | HIGH | 92 | meaningful pantomimes>unknown sign language |
| | | | | | | | | | meaningful pantomimes>dog videos |
| | | | | | | | | | synonyms>letter strings |
| Hagoort et al. | 1999 | PET | NO | IMPLICIT | VERBAL | VISUAL | HIGH | 10 | words>pseudowords |
| Harrington, Farias & Davis | 2009 | fMRI | NO | EXPLICIT | NONVERBAL | VISUAL | HIGH | 8 | familiar>non objects |
| Hauk & Pulvermueller | 2011 | fMRI | NO | IMPLICIT | VERBAL | VISUAL | LOW | 21 | action words>hashmarks |
| | | | | | | | | | uni manual action words>hashmarks |
| | | | | | | | | | uni manual action words>hashmarks |
| Hayashi et al. | 2014 | fMRI | NO | EXPLICIT | VERBAL | VISUAL | LOW | 16 | concrete word>astersisks |
| | | | | | | | | | abstract word>asterisks |
| Heim, Eickhoff & Amunts | 2008 | fMRI | NO | EXPLICIT | VERBAL | VISUAL | HIGH | 28 | semantic > phonological fluency |
| Henke et al. | 1999 | PET | NO | EXPLICIT | VERBAL | VISUAL | HIGH | 12 | associative word learning>single word encoding |
| Herbster et al. | 1997 | PET | NO | EXPLICIT | VERBAL | VISUAL | HIGH | 10 | Irregular>zero order speak |
| | | | | | | | | | regular>zero order speak |

| | | | | | | | | | |
|---|---|---|---|---|---|---|---|---|---|
| | | | | | | | | | irregular+regular>zero order speak |
| Higuchi, Moriguchi, Murakami, Katsunuma, Mishima &Uno | 2015 | fMRI | NO | IMPLICIT | VERBAL | VISUAL | LOW | 28 | all characters>checkerboard |
| Homae, Yahata & Sakai | 2003 | fMRI | NO | EXPLICIT | VERBAL | VISUAL | HIGH | 10 | visual sentences>visual non words words |
| Husain, Patkin, Kim, Braun & Horwitz | 2012 | fMRI | NO | EXPLICIT | NONVERBAL | VISUAL | HIGH | 16 | meaningful iconic>meaningless gestures |
| Ikuta et al. | 2006 | fMRI | NO | IMPLICIT | VERBAL | VISUAL | LOW | 34 | sentences>word lists |
| Jackson, Hoffman, Pobric, Lambon Ralph | 2015 | fMRI | NO | EXPLICIT | VERBAL | VISUAL | HIGH | 24 | words>letter strings |
| Jensen, Hargreaves, Bass, Pexman, Goodyear & Federico | 2011 | fMRI | NO | EXPLICIT | VERBAL | VISUAL | HIGH | 12 | words>pseudowords |
| Jeon, Lee, Kim & Cho | 2009 | fMRI | NO | EXPLICIT | VERBAL | VISUAL | HIGH | 16 | synonys>nonwords |
| | | | | | | | | | antonyms>nonwords |
| | | | | | | | | | English synonyma>nonwords |
| | | | | | | | | | Korean synonyms>nonwords |
| | | | | | | | | | honorific words>nonwords |
| Joubert et al. | 2004 | fMRI | NO | IMPLICIT | VERBAL | VISUAL | HIGH | 10 | low frequency words>nonwords |
| | | | | | | | | | high frequency words>consonant strings |
| | | | | | | | | | low frequency words>consonant strings |
| Kang et al. | 2006 | PET | NO | EXPLICIT | VERBAL | VISUAL | HIGH | 17 | visual speech>facial movements (chewing gum) |
| Khader, Jost, Mertens, Bien & Roesler | 2010 | fMRI | NO | EXPLICIT | VERBAL | VISUAL | HIGH | 16 | noun>rhyme generation |
| | | | | | | | | | verb>rhyme generation |
| | | | | | | | | | noun generation>letter detection |
| | | | | | | | | | verb generation>letter detection |
| Kim et al. | 2009 | fMRI | NO | EXPLICIT | VERBAL | VISUAL | HIGH | 36 | sentences>word lists |
| Kinno, Kawamura, Shioda & Sakai | 2008 | fMRI | NO | EXPLICIT | BOTH | VISUAL | HIGH | 14 | canonical sentence>picture & letter strings |
| | | | | | | | | | active sentence>picture & letter strings |
| | | | | | | | | | passive sentence>picture & letter strings |
| Kuchinke et al. | 2005 | fMRI | YES | IMPLICIT | VERBAL | VISUAL | HIGH | 20 | emotion word>nonword |
| Kumar | 2016 | fMRI | NO | EXPLICIT | VERBAL | VISUAL | HIGH | 20 | abstract+concrete words>pseudowords |

| | | | | | | | | | |
|---|---|---|---|---|---|---|---|---|---|
| | | | | | | | | | abstract words>pseudowords |
| Leveroni et al. | 2000 | fMRI | YES | EXPLICIT | NONVERBAL | VISUAL | HIGH | 11 | familiar faces>foils(never seen faces) |
| | | | | | | | | | newly learned faces>foils(never seen faces) |
| Lin, Wang, Zhao, Liu, Li & Bi | 2015 | fMRI | NO | EXPLICIT | VERBAL | VISUAL | HIGH | 20 | words>pseudowords |
| Liu et al. | 2009 | fMRI | NO | EXPLICIT | VERBAL | VISUAL | LOW | 16 | words meaning>slashes |
| | | | | | | | | | words rhyming>slashes |
| Ludersdorfer, Schurz, Richlan, Kronbichler & Wimmer | 2013 | fMRI | NO | EXPLICIT | VERBAL | VISUAL | HIGH | 29 | words>false fonts |
| | | | | | | | | | words>pseudowords |
| Malins, Gumkowski, Buis, Molfese, Rueckl, Frost, Pugh, Morris & Mencl | 2016 | fMRI | NO | IMPLICIT | VERBAL | VISUAL | LOW | 18 | unrelated words>false font |
| | | | | | | | | | unrelated>pseudowords |
| Marques, Canessa & Cappa | 2009 | fMRI | NO | EXPLICIT | VERBAL | VISUAL | LOW | 21 | sentences>crosses |
| Marques, Canessa, Siri, Catricala & Cappa | 2008 | fMRI | NO | EXPLICIT | VERBAL | VISUAL | LOW | 21 | semantic features>baseline task |
| Mashal, Vishne, Laor & Titone | 2013 | fMRI | NO | EXPLICIT | VERBAL | VISUAL | HIGH | 14 | novel metaphor>unrelated words |
| | | | | | | | | | conventional metaphor>unrelated words |
| Matchin, Liao, gaston & Lau | 2019 | fMRI | NO | EXPLICIT | VERBAL | VISUAL | HIGH | 20 | verb phrase>list |
| | | | | | | | | | noun phrase>list |
| Matchin, Hammerly & Lau | 2017 | fMRI | NO | EXPLICIT | VERBAL | VISUAL | HIGH | 16 | sentences>word lists |
| | | | | | | | | | sentences>phrases |
| | | | | | | | | | real>pseudoword lists |
| | | | | | | | | | real>pseudoword phrases |
| | | | | | | | | | real>pseudoword sentences |
| Mellem, Jasmin, Peng & Martin | 2016 | fMRI | NO | IMPLICIT | VERBAL | VISUAL | LOW | 20 | longer>shorter phrase |
| Menz, Blangero, Kunze & Binkofski | 2010 | fMRI | NO | EXPLICIT | NONVERBAL | VISUAL | HIGH | 20 | known>unknown objects |
| Moberget, Gullesen, Andersson, Ivry & Endestad | 2014 | fMRI | NO | EXPLICIT | VERBAL | VISUAL | HIGH | 32 | incongruent>scrambled sentence |
| | | | | | | | | | congruent>scrambled sentence |

| | | | | | | | | | |
|---|---|---|---|---|---|---|---|---|---|
| Moseley, Carota, Hauk, Mohr & Pulvermueller | 2012 | fMRI | YES | IMPLICIT | VERBAL | VISUAL | LOW | 18 | all emotional words>hashmarks |
| | | | | | | | | | abstract emotional words>hashmarks |
| | | | | | | | | | arm+face+emotion words>hashmarks |
| | | | | | | | | | face words>hashmarks |
| | | | | | | | | | arm words>hashmarks |
| Mummery, Patterson, Hodges & Price | 1998 | PET | NO | EXPLICIT | VERBAL | VISUAL | HIGH | 10 | semantic>phonological decision |
| Nakamura et al. | 2000 | PET | YES | EXPLICIT | NONVERBAL | VISUAL | LOW | 7 | familiar faces>fixation cross |
| Nichelli, Grafman, Pietrini, Clark, Lee & Miletich | 1995 | PET | NO | EXPLICIT | VERBAL | VISUAL | HIGH | 9 | semantic>orthographic decision |
| Nielson et al. | 2010 | fMRI | YES | EXPLICIT | BOTH | VISUAL | HIGH | 17 | familiar>unfamiliar people |
| Pallier, Devauchelle & Dehaene | 2011 | fMRI | NO | EXPLICIT | VERBAL | VISUAL | HIGH | 40 | longer>shorter phrase |
| | | | | | | | | | length of real>pseudoword sentences |
| Perani, Schnur, Tettamanti, Gorno-Tempini, Cappa & Fazio | 1999 | PET | NO | EXPLICIT | NONVERBAL | VISUAL | HIGH | 11 | living objects>shapes |
| | | | | | | | | | nonliving objects>shapes |
| Perani, Schnur, Tettamanti, Gorno-Tempini, Cappa & Fazio | 1999 | PET | NO | EXPLICIT | VERBAL | VISUAL | HIGH | 8 | living words>pseudowords |
| | | | | | | | | | nonliving words>pseudowords |
| Perrone-Bertolotti, Kauffmann, Pichat, Vidal & Baciu | 2017 | fMRI | NO | EXPLICIT | VERBAL | VISUAL | HIGH | 24 | words>unreadable font |
| Pilgrim, Fadili, Fletcher & Tyler | 2002 | fMRI | NO | EXPLICIT | VERBAL | VISUAL | HIGH | 14 | words>letter strings |
| Price, Moore, Humphreys & Wise | 1997 | PET | NO | EXPLICIT | VERBAL | VISUAL | HIGH | 6 | semantic>phonological decision |
| Pulvermueller, Cook & Hauk | 2012 | fMRI | NO | IMPLICIT | VERBAL | VISUAL | LOW | 23 | phrases>hashmarks |
| | | | | | | | | | uninflected words>hashmarks |
| | | | | | | | | | inflected words>hashmarks |
| Raposo, Frade & Alves | 2016 | fMRI | NO | IMPLICIT | VERBAL | VISUAL | HIGH | 18 | semantic>perceptual decision |
| Rapp & Lipka | 2011 | fMRI | NO | IMPLICIT | VERBAL | VISUAL | LOW | 10 | words>checkerboards |

| | | | | | | | | | |
|---|---|---|---|---|---|---|---|---|---|
| | | | | | | | | | words>letter strings |
| Redcay, Velnoskey & Rowe | 2016 | fMRI | NO | EXPLICIT | BOTH/NONVERBAL/VERBAL | VISUAL | HIGH | 24 | meaningful>meaningless stimuli |
| | | | | | | | | | communicative>non-communicative gesture |
| | | | | | | | | | real>pseudoword sentences |
| Robertson et al. | 2000 | fMRI | NO | IMPLICIT | VERBAL | VISUAL | HIGH | 8 | indefinite article sentence>letter strings |
| | | | | | | | | | definite article sentence>letter strings |
| Rogers et al. | 2006 | PET | NO | EXPLICIT | BOTH/NONVERBAL | VISUAL | HIGH | 12 | pictures>scrambled pictures |
| | | | | | | | | | specific-level judgement>baseline |
| Roskies, Fiez, Balota, Raichle & Petersen | 2001 | PET | NO | EXPLICIT | VERBAL | VISUAL | HIGH | 20 | semantic>phonological decision |
| Ross & Olson | 2012 | fMRI | YES | EXPLICIT | NONVERBAL | VISUAL | HIGH | 11 | famous>unknown faces & landmarks |
| Ryan, Cox, Hayes & Nadel | 2008 | fMRI | NO | EXPLICIT | VERBAL | VISUAL | HIGH | 10 | semantic fluency (generate) > crosses |
| | | | | | | | | | semantic fluency (recall) > crosses |
| | | | | | | | | | semantic fluency (recall & geneerate)>crosses |
| Ryan, Lin, Ketcham & Nadel | 2010 | fMRI | NO | EXPLICIT | VERBAL | VISUAL | HIGH | 15 | semantic spatial old>letter judgement |
| | | | | | | | | | semantic spatial new>letter judgement |
| | | | | | | | | | semantic non spatial old>letter judgement |
| | | | | | | | | | semantic non spatial new>letter judgement |
| | | | | | | | | | semantic new>episodic judgement |
| Sachs, Weis, Krings, Huber & Kircher | 2008 | fMRI | NO | EXPLICIT | VERBAL | VISUAL | HIGH | 14 | biased thematic judgement>letters |
| | | | | | | | | | biased taxonomic judgement>letters |
| | | | | | | | | | balanced taxonomic judgement>letters |
| | | | | | | | | | balanced taxonomic judgement>letters |
| Schuil, Smits & Zwaan | 2013 | fMRI | NO | EXPLICIT | VERBAL | VISUAL | HIGH | 20 | senteces>pseudowords |
| | | | | | | | | | verbs>pseudowords |
| | | | | | | | | | literal sentences>pseudowords |
| | | | | | | | | | nonliteral sentences>pseudowords |
| Segal & Petrides | 2012 | fMRI | NO | EXPLICIT | NONVERBAL/VERBAL | VISUAL | HIGH | 90 | writing>copying |

| | | | | | | | | | |
|---|---|---|---|---|---|---|---|---|---|
| | | | | | | | | | words>pseudowords |
| Seghier, Josse, Leff & Price | 2011 | fMRI | NO | EXPLICIT | BOTH | VISUAL | HIGH | 60 | meaningful>meaningless stimuli |
| Sergent, Otha & Macdonald | 1992 | PET | YES | EXPLICIT | VERBAL/NON VERBAL | VISUAL | HIGH/LOW | 7 | face identity>gender discrimination |
| | | | | | | | | | object recognition>gratings |
| Sheldon, McAndrews, Pruessner & Moscovitch | 2016 | fMRI | NO | EXPLICIT | VERBAL | VISUAL | HIGH | 15 | semantic fluency>perceptual task |
| Simard, Monetta, Nagano-Saito & Monchi | 2013 | fMRI | NO | EXPLICIT | VERBAL | VISUAL | HIGH | 14 | semantic>control macthing |
| | | | | | | | | | semantic matching>phonological decision (syllable rhyme) |
| | | | | | | | | | semantic matching>phonological decision (syllable onset) matching |
| Slioussar, Kireev, Chernigovskaya, Kataeva, Korotkov & Medvedev | 2014 | fMRI | NO | EXPLICIT | VERBAL | VISUAL | HIGH | 21 | real verbs>pseudowords |
| | | | | | | | | | real nouns>pseudowords |
| Smith, Myers, Sethi, Pantazatos, Yanagihara & Hirsch | 2012 | fMRI | NO | EXPLICIT | VERBAL | VISUAL | HIGH | 14 | semantics>baseline |
| Snijders, Vosse, Kempen, Van Berkum, Petersson & Hagoort | 2009 | fMRI | NO | IMPLICIT | VERBAL | VISUAL | HIGH | 28 | sentences>word lists |
| Stowe, Paans, Wijers, Zwarts, Mulder & Vaalburg | 1999 | PET | NO | IMPLICIT | VERBAL | VISUAL | HIGH | 12 | sentences>scrambled word lists |
| Straube, Green, Weis & Kircher | 2012 | fMRI | NO | IMPLICIT | NONVERBAL | VISUAL | HIGH | 16 | iconic>meaningless gesture |
| Stringaris, Medford, Giampietro. Brammer & David | 2007 | fMRI | NO | EXPLICIT | VERBAL | VISUAL | HIGH | 11 | literal>meaningless sentences |
| | | | | | | | | | methaphors>meaningless sentences |
| Sugiura et al. | 2006 | fMRI | YES | EXPLICIT | VERBAL | VISUAL | HIGH | 24 | famous>unfamiliar names |
| | | | | | | | | | personal>unfamiliar names |
| Sugiura et al. | 2001 | PET | YES | EXPLICIT | NONVERBAL | VISUAL | HIGH | 5 | identity discrimination>control |
| | | | | | | | | | identity discrimination>face direction |
| Sugiura et al. | 2008 | fMRI | YES | EXPLICIT | VERBAL | VISUAL | HIGH | 25 | low familiar>unfamiliar names |

| | | | | | | | | | |
|---|---|---|---|---|---|---|---|---|---|
| | | | | | | | | | personal familiar>unfamiliar names |
| | | | | | | | | | high familiar>unfamiliar names |
| | | | | | | | | | high familiar>unfamiliar names |
| Sun, Xue, Zhang, Zuo, Chen, Wang, Martin, Wang, Chen, He & Wang | 2017 | fMRI | NO | EXPLICIT | VERBAL | VISUAL | HIGH | 11 | semantic>orthography judgement |
| Szlachta, Bozic, Jelowicka & Marslen-Wilson | 2012 | fMRI | NO | IMPLICIT | VERBAL | VISUAL | LOW | 21 | words>musical rain |
| | | | | | | | | | nouns>musical rain |
| | | | | | | | | | inflected nouns>musical rain |
| Taminato, Miura, Sugiura & Kawashima | 2014 | fMRI | NO | EXPLICIT | NONVERBAL | VISUAL | LOW | 35 | object recognition>control task |
| | | | | | | | | | object recognition>control task |
| Taylor, Arsalidou, Bayless, Morris, Evans & Barbeau | 2009 | fMRI | YES | IMPLICIT | NONVERBAL | VISUAL | HIGH | 10 | own>unfamiliar face |
| | | | | | | | | | partner's>unfamiliar face |
| | | | | | | | | | parent's>unfamiliar face |
| Thierry & Price | 2006 | PET | NO | EXPLICIT | VERBAL/NON VERBAL | VISUAL | HIGH | 12 | visual words>text control (scrambled letter strings) |
| | | | | | | | | | visual videos>video control (distorted) |
| Tieleman, Seurinck, Deblaere, Vandemaele, Vingerhoets & Achten | 2005 | fMRI | NO | EXPLICIT | VERBAL | VISUAL | HIGH | 22 | self-paced semantic>perceptual decision |
| | | | | | | | | | fixed-paced semantic>perceptual decision |
| Tyler, Stamatakis, Dick, Bright, Fletcher & Moss | 2003 | fMRI | NO | EXPLICIT | VERBAL | VISUAL | HIGH | 12 | animals>baseline |
| | | | | | | | | | tool action words>baseline |
| | | | | | | | | | biological action>baseline |
| Van Ettinger-Veenstra, McAllister, Lundberg, Karlsson & Engstrom | 2016 | fMRI | NO | EXPLICIT | VERBAL | VISUAL | HIGH | 27 | sentences>symbol strings |
| Vignali, Hawelka, Hutzler & Richlan | 2019 | fMRI | NO | EXPLICIT | VERBAL | VISUAL | HIGH | 21 | foveal & parafovealwords>foveal & parafoveal pseudowords |
| Vingerhoets | 2008 | fMRI | NO | IMPLICIT | NONVERBAL | VISUAL | HIGH | 14 | familiar>unfamiliar tools |

| | | | | | | | | | |
|---|---|---|---|---|---|---|---|---|---|
| Visser, Jefferies, Embleton & Lambon Ralph | 2012 | fMRI | NO | EXPLICIT | BOTH/NONVERBAL/VERBAL | VISUAL | HIGH | 15 | semantics>baseline |
| | | | | | | | | | pictures>baseline |
| | | | | | | | | | words>baseline |
| Wang, Zhao, Zevin & Yang | 2016 | fMRI | NO | EXPLICIT | VERBAL | VISUAL | HIGH | 16 | words>nonsense strokes |
| Weiss, Katzir & Bitan | 2015 | fMRI | NO | IMPLICIT | VERBAL | VISUAL | LOW | 18 | pointed words>asterisks |
| | | | | | | | | | unpointed words>asterisks |
| Welcome & Joanisse | 2012 | fMRI | NO | EXPLICIT | VERBAL | VISUAL | HIGH | 20 | semantic>phonological/orthographic decision |
| Wende, Straube, Stratmann, Sommer, Kircher & Nagels | 2012 | fMRI | NO | EXPLICIT | VERBAL | VISUAL | HIGH | 18 | semantic>phonological fluency |
| Wirth, Jann, Dierks, Federspiel, Wiest & Horn | 2011 | fMRI | NO | EXPLICIT | VERBAL | VISUAL | HIGH | 19 | semantic>phonological & perceptual decision |
| Wright et al. | 2008 | fMRI | NO | EXPLICIT | BOTH/VERBAL/NONVERBAL | VISUAL | HIGH | 10 | semantic>perceptual matching |
| | | | | | | | | | semantic>perceptual matching |
| | | | | | | | | | semantic>perceptual matching |
| Wu, Mai, Tang, Ge, Luo & Liu | 2013 | fMRI | NO | IMPLICIT | VERBAL | VISUAL | LOW | 19 | arm words>checkerboard |
| | | | | | | | | | leg words>checkerboard |
| | | | | | | | | | mouth words>checkerboard |
| Yang, Li, Fang, Shu, Liu & Chen | 2016 | fMRI | NO | IMPLICIT | VERBAL | VISUAL | LOW | 20 | opaque idioms>hashmarks |
| | | | | | | | | | transparent idioms>hashmarks |
| | | | | | | | | | literal phrases>hashmarks |
| Zaccarella & Friederici | 2015 | fMRI | NO | EXPLICIT | VERBAL | VISUAL | HIGH | 22 | words>pseudowords |
| Zhang, Liu & Zhang | 2014 | fMRI | NO | EXPLICIT | VERBAL | VISUAL | LOW | 18 | nonliving words>asterisks |
| | | | | | | | | | living words>asterisks |
| Zhang, Xiao & Weng | 2012 | fMRI | NO | EXPLICIT | VERBAL | VISUAL | HIGH | 14 | words>pseudowords |
| EXCLUDED CONTRASTS CONTAINING BOTH VISUAL AND AUDITORI STIMULI. THE EXLUDED AUDITORY EXPERIMENTS CAN BE SEEN IN Table M.2.4 SC NON-AUDITORY | | | | | | | | | |
| Brambati, Benoit, Monetta, Belleville & Joubert | 2010 | fMRI | YES | EXPLICIT | BOTH | BOTH | HIGH | 12 | general & specific occupation judgement>baseline (scrambled face) |

| | | | | | | | | | |
|---|---|---|---|---|---|---|---|---|---|
| Europa, Gitelman, Kiran & Thompson | 2019 | fMRI | NO | EXPLICIT | BOTH | BOTH | HIGH | 21 | sentences>baseline (reversed sentences) |
| Holle, Gunter, Rueschemeyer, Hennenlotter & Iacoboni | 2008 | fMRI | NO | EXPLICIT | BOTH | BOTH | HIGH | 17 | iconic gesture of dominant meaning>grooming |
| | | fMRI | NO | EXPLICIT | BOTH | BOTH | HIGH | 17 | iconic gesture of subordinate meaning>grooming |
| Kang et al. | 2006 | PET | NO | EXPLICIT | VERBAL | BOTH | HIGH | 17 | audio-visual speech>noises and facial movements |
| Liuzzi et al. | 2017 | fMRI | NO | EXPLICIT | VERBAL | BOTH | HIGH | 18 | semantic judgement > input modality detection |
| Vagharchakian, Dehaene-Lambertz, Pallier & Dehaene | 2012 | fMRI | NO | EXPLICIT | VERBAL | BOTH | HIGH | 16 | intelligible>unintelligble compression rate |
| | | fMRI | NO | EXPLICIT | VERBAL | BOTH | HIGH | 16 | intelligible>unintelligble compression rate |
| van Leeuwen et al. | 2014 | fMRI | NO | EXPLICIT | BOTH | BOTH | HIGH | 16 | speech>reversed speech |
| Zvyagintsev, Clemens, Chechko, Mathiak, Sack & Mathiak | 2013 | fMRI | NO | IMPLICIT | NONE | NONE | HIGH | 15 | visual imagery>counting |
| | | fMRI | NO | IMPLICIT | NONE | NONE | HIGH | 15 | auditory imagery>counting |

**The references to access the listed studies are listed at the end of the document*

***Table M2.4 SC AUDITORY** List of studies included in the **semantic cognition AUDITORY INPUT MODALITY** (N = 60) meta-analysis after excluding VISUAL contrasts. **Note:** The list of excluded studies with VISUAL INPUT MODALITY and contrasts with BOTH VISUAL and AUDITORY stimuli that were also excluded can be seen in Table M.2.3 SC VISUAL.*

| *Authors | Year | Imaging Method | Social Content | Instructional Cue | Stimulus Domain | Sensory Input Modality | Baseline Type | N | Contrast |
|---|---|---|---|---|---|---|---|---|---|
| Alain, He & Grady | 2008 | fMRI | NO | EXPLICIT | VERBAL | AUDITORY | HIGH | 16 | sound category>sound location |
| Baumgaertner et al. | 2007 | fMRI | NO | IMPLICIT | VERBAL | AUDITORY | HIGH | 19 | sentences>reversed sentences |
| Bautista & Wilson | 2016 | fMRI | NO | IMPLICIT | VERBAL | AUDITORY | HIGH | 12 | clear>scrambled rotated speech |
| Binder, Frost, Hammeke, Bellgowan, Rao & Cox | 1999 | fMRI | NO | EXPLICIT | VERBAL | AUDITORY | HIGH | 30 | semantic>phonological decision |
| Bozic & Marslen-Wilson | 2013 | fMRI | NO | IMPLICIT | VERBAL | AUDITORY | HIGH | 13 | speech>musical rain |
| Cai, Kochiyama, Osaka & Wu | 2007 | fMRI | NO | IMPLICIT | VERBAL | AUDITORY | HIGH | 15 | words>nonsense words |
| Christensen, Antonucci, Lockwood, Kittleson & Plante | 2008 | fMRI | NO | EXPLICIT | VERBAL | AUDITORY | HIGH | 14 | diotic listening>reversed speech |
| | | | | | | | | | dichotic listening>reversed speech |
| Clos, Langner, Meyer, Oechslin, Zilles & Eickhoff | 2014 | fMRI | NO | EXPLICIT | VERBAL | AUDITORY | HIGH | 29 | intelligibilty based on cue > unintelligible |
| Davis, Ford, Kherif & Johnsrude | 2011 | fMRI | NO | EXPLICIT | VERBAL | AUDITORY | LOW | 12 | clear & anomalous sentences>signal correlated noise |
| Demonet et al. | 1992 | PET | NO | EXPLICIT | VERBAL | AUDITORY | HIGH | 9 | words>phonemes |
| | | | | | | | | | words>tones |
| Engelien et al. | 2006 | PET | NO | IMPLICIT | NONVERBAL | AUDITORY | HIGH | 6 | meaningful>meaningless sounds |
| Erb, Henry, Eisner & Obleser | 2013 | fMRI | NO | EXPLICIT | VERBAL | AUDITORY | HIGH | 30 | speech>vocoded speech |
| Friederici, Kotz, Scott & Obleser | 2010 | fMRI | NO | IMPLICIT | VERBAL | AUDITORY | HIGH | 17 | intelligible speech>rotated speech |
| Giraud & Price | 2001 | PET | NO | BOTH | BOTH | AUDITORY | BOTH | 12 | words+environmental sounds>syllables+noise |
| Giraud et al. | 2004 | fMRI | NO | EXPLICIT | VERBAL | AUDITORY | HIGH | 8 | natural speech>speech envelope |
| Groussard et al. | 2010 | PET | NO | EXPLICIT | NONVERBAL | AUDITORY | HIGH | 11 | musical semantic>musical reference |
| Guediche, Reilly, Santiago, Laurent & Blumstein | 2016 | fMRI | NO | IMPLICIT | VERBAL | AUDITORY | LOW | 16 | related > repeated sentence |

| | | | | | | | | | |
|---|---|---|---|---|---|---|---|---|---|
| | | | | | | | | | unrelated > repeated sentence |
| Gurd et al. | 2002 | fMRI | NO | EXPLICIT | VERBAL | AUDITORY | HIGH | 11 | category>rote fluency |
| Hartung, Hagoort & Willems | 2017 | fMRI | NO | IMPLICIT | VERBAL | AUDITORY | HIGH | 52 | first-person speech>unintelligible reversed speech |
| | | | | | | | | | third-person speech>unintelligible reversed speech |
| Hervais-Adelman, Carlyon, Johnsrude & Davis | 2012 | fMRI | NO | EXPLICIT | VERBAL | AUDITORY | HIGH | 15 | clear>vocoded speech (uninteligible) |
| Hocking, McMahon & de Zubicaray | 2011 | fMRI | NO | EXPLICIT | NONVERBAL | AUDITORY | HIGH | 13 | all environmental sounds>perceptual baseline |
| Homae, Yahata & Sakai | 2003 | fMRI | NO | EXPLICIT | VERBAL | AUDITORY | HIGH | 10 | auditory sentences>auditory non words |
| Hwang, Palmer, Basho, Zadra & Muller | 2009 | fMRI | NO | EXPLICIT | VERBAL | AUDITORY | LOW | 13 | fluency generation>production baseline |
| Kang et al. | 2006 | PET | NO | EXPLICIT | VERBAL | AUDITORY | LOW | 17 | auditory speech>white noise |
| Kotz, Cappa, Von Cramon & Friederici | 2002 | fMRI | NO | EXPLICIT | VERBAL | AUDITORY | HIGH | 13 | words>pseudowords |
| Kuperberg et al. | 2000 | fMRI | NO | EXPLICIT | VERBAL | AUDITORY | HIGH | 4 | sentences>words strings |
| Kyong, Scott, Rosen, Howe, Agnew & McGettigan | 2014 | fMRI | NO | IMPLICIT | VERBAL | AUDITORY | LOW | 19 | intelligible vocoded>inverted vocoded speech |
| Leff, Schofield, Stephan, Crinion, Friston & Price | 2008 | fMRI | NO | IMPLICIT | VERBAL | AUDITORY | HIGH | 26 | speech>reversed speech |
| Leung & Alain | 2011 | fMRI | NO | EXPLICIT | NONVERBAL | AUDITORY | HIGH | 16 | semantic>location matching |
| Liu et al. | 2009 | fMRI | NO | EXPLICIT | VERBAL | AUDITORY | LOW | 16 | words meaning>tones |
| | | | | | | | | | words rhyming>tones |
| Ludersdorfer, Wimmer, Richlan, Schurz, Hutzler & Kronbichler | 2016 | fMRI | NO | EXPLICIT | VERBAL | AUDITORY | LOW | 29 | words orthographic>tones |
| | | | | | | | | | words semantic>tones |
| Ludersdorfer, Schurz, Richlan, Kronbichler & Wimmer | 2013 | fMRI | NO | EXPLICIT | VERBAL | AUDITORY | HIGH | 29 | speech>reversed speech |
| | | | | | | | | | words>pseudowords |
| Meyer, Alter, Friederici, Lohmann & Yves von Cramon | 2002 | fMRI | NO | IMPLICIT | VERBAL | AUDITORY | HIGH | 14 | word>pseudoword sentence |
| Nakamura et al. | 2001 | PET | YES | EXPLICIT | VERBAL | AUDITORY | HIGH | 9 | familiar voice>vowel discrimination |

| | | | | | | | | | |
|---|---|---|---|---|---|---|---|---|---|
| | | | | | | | | | self voice>vowel discrimination |
| Noppeney & Price | 2003 | PET | NO | EXPLICIT | VERBAL | AUDITORY | LOW | 9 | normal>reversed words |
| Orfanidou, Marlsen-Wilson & Davis | 2006 | fMRI | NO | EXPLICIT | VERBAL | AUDITORY | HIGH | 13 | words>pseudowords |
| Peelle, Eason, Schmitter, Schwarzbauer & Davis | 2010 | fMRI | NO | EXPLICIT | VERBAL | AUDITORY | HIGH | 6 | sentences>signal correlated noise |
| Raettig & Kotz | 2008 | fMRI | NO | EXPLICIT | VERBAL | AUDITORY | HIGH | 16 | real words>pseudowords |
| Raposo, Moss, Stamatakis & Tyler | 2009 | fMRI | NO | IMPLICIT | VERBAL | AUDITORY | LOW | 22 | action sentences>SC noise |
| Rissman, Eliassen & Blumstein | 2003 | fMRI | NO | IMPLICIT | VERBAL | AUDITORY | HIGH | 15 | words>pseudowords |
| Rodd, Johnsrude & Davis | 2012 | fMRI | NO | EXPLICIT | VERBAL | AUDITORY | LOW | 15 | speech>SCN |
| Rodd, Longe, Randall & Tyler | 2010 | fMRI | NO | EXPLICIT | VERBAL | AUDITORY | LOW | 14 | speech>SCN |
| Rogalsky & Hickok | 2009 | fMRI | NO | IMPLICIT | VERBAL | AUDITORY | HIGH | 14 | sentences>word lists |
| Rogalsky, Almeida, Sprouse & Hickok | 2015 | fMRI | NO | IMPLICIT | VERBAL | AUDITORY | HIGH | 15 | words>scrambled |
| Roxbury, McMahon & Copland | 2014 | fMRI | NO | EXPLICIT | VERBAL | AUDITORY | HIGH | 17 | concrete word>pseudoword |
| | | | | | | | | | abstract word>pseudoword |
| Sabri, Binder, Desai, Medler, Leitl & Liebenthal | 2008 | fMRI | NO | EXPLICIT | VERBAL | AUDITORY | HIGH | 28 | speech>rotated speech |
| | | | | | | | | | words>pseudowords |
| Saur et al. | 2008 | fMRI | NO | IMPLICIT | VERBAL | AUDITORY | HIGH | 33 | word>pseudoword sentences |
| Schell, Zaccarella & Friederici | 2017 | fMRI | NO | EXPLICIT | VERBAL | AUDITORY | HIGH | 21 | phrases>non combinatorial words |
| Schmitt, Auer & Ferstl | 2019 | fMRI | NO | EXPLICIT | VERBAL | AUDITORY | HIGH | 40 | known>unknown language |
| Scott, Blank, Rosen & Wise | 2000 | PET | NO | IMPLICIT | VERBAL | AUDITORY | HIGH | 8 | intelligible>unintelligible speech |
| Straube, Green, Weis & Kircher | 2012 | fMRI | NO | IMPLICIT | VERBAL | AUDITORY | HIGH | 16 | known>unknown language |
| Takeichi, Koyama, Terao, Takeuchi, Toyosawa & Murohashi | 2010 | fMRI | NO | IMPLICIT | VERBAL | AUDITORY | HIGH | 23 | speech>reversed speech |
| | | | | | | | | | speech>modulated speech |
| Thierry & Price | 2006 | PET | NO | EXPLICIT | VERBAL/NON VERBAL | AUDITORY | HIGH | 12 | auditory words>speech control(scrambled) |

| | | | | | | | | | |
|---|---|---|---|---|---|---|---|---|---|
| | | | | | | | | | auditory sounds>sound control (scrambled) |
| Vagharchakian, Dehaene-Lambertz, Pallier & Dehaene | 2012 | fMRI | NO | EXPLICIT | VERBAL | AUDITORY | HIGH | 16 | intelligible>unintelligble compression rate |
| Vitello, Warren, Devlin & Rodd | 2014 | fMRI | NO | EXPLICIT | VERBAL | AUDITORY | LOW | 20 | sentences>SCN |
| von Kriegstein, Eger, Kleinschmidt & Giraud | 2003 | fMRI | NO | EXPLICIT | VERBAL | AUDITORY | HIGH | 14 | sentence>speech envelope |
| Wright, Randall, Marslen-Wilson & Tyler | 2011 | fMRI | NO | BOTH | VERBAL | AUDITORY | LOW | 14 | speech>musical rain |
| Xiao et al. | 2005 | fMRI | NO | EXPLICIT | VERBAL | AUDITORY | HIGH | 14 | words>pseudowords |
| Zhuang & Devereux | 2017 | fMRI | NO | EXPLICIT | VERBAL | AUDITORY | HIGH | 16 | phrases>words |
| Zou, Packard, Xia, Liu & Shu | 2016 | fMRI | NO | IMPLICIT | VERBAL | AUDITORY | LOW | 17 | speech>tone |
| | | | | | | | | | speech>tone |
| | | | | | | | | | speech>tone |
| | | | | | | | | | identical speech>tone |
| | | | | | | | | | words>tone |
| EXCLUDED CONTRASTS CONTAINING BOTH VISUAL AND AUDITORI STIMULI. THE EXLUDED VISAUAL EXPERIMENTS CAN BE SEEN IN Table M.2.3 SC VISUAL | | | | | | | | | |
| Brambati, Benoit, Monetta, Belleville & Joubert | 2010 | fMRI | YES | EXPLICIT | BOTH | BOTH | HIGH | 12 | general & specific occupation judgement>baseline (scrambled face) |
| Europa, Gitelman, Kiran & Thompson | 2019 | fMRI | NO | EXPLICIT | BOTH | BOTH | HIGH | 21 | sentences>baseline (reversed sentences) |
| Holle, Gunter, Rueschemeyer, Hennenlotter & Iacoboni | 2008 | fMRI | NO | EXPLICIT | BOTH | BOTH | HIGH | 17 | iconic gesture of dominant meaning>grooming |
| | | fMRI | NO | EXPLICIT | BOTH | BOTH | HIGH | 17 | iconic gesture of subordinate meaning>grooming |
| Kang et al. | 2006 | PET | NO | EXPLICIT | VERBAL | BOTH | HIGH | 17 | audio-visual speech>noises and facial movements |
| Liuzzi et al. | 2017 | fMRI | NO | EXPLICIT | VERBAL | BOTH | HIGH | 18 | semantic judgement > input modality detection |
| Vagharchakian, Dehaene-Lambertz, Pallier & Dehaene | 2012 | fMRI | NO | EXPLICIT | VERBAL | BOTH | HIGH | 16 | intelligible>unintelligble compression rate |
| | | fMRI | NO | EXPLICIT | VERBAL | BOTH | HIGH | 16 | intelligible>unintelligble compression rate |
| van Leeuwen et al. | 2014 | fMRI | NO | EXPLICIT | BOTH | BOTH | HIGH | 16 | speech>reversed speech |

| | | | | | | | | | |
|---|---|---|---|---|---|---|---|---|---|
| Zvyagintsev, Clemens, Chechko, Mathiak, Sack & Mathiak | 2013 | fMRI | NO | IMPLICIT | NONE | NONE | HIGH | 15 | visual imagery>counting |
| | | fMRI | NO | IMPLICIT | NONE | NONE | HIGH | 15 | auditory imagery>counting |

**The references to access the listed studies are listed at the end of the document*

#### *Theory of Mind Data Sets and Subsets*

***Table M1 ToM*** *List of studies included in the* ***theory of mind all baselines*** *(N =114) meta-analysis.* ***Note:*** *ToM= Theory of Mind, N= Sample Size*

| *Authors | Year | Imaging Method | Instructional Cue | Stimulus Domain | Sensory Input Modality | Baseline Type | N | Contrast | Age Range |
|---|---|---|---|---|---|---|---|---|---|
| Abraham et al. | 2008 | fMRI | Explicit | Verbal | visual | high | 17 | stories about mental states > control stories | 25.65 (22-30) |
| Abraham et al. | 2010 | fMRI | Explicit | Verbal | visual | high | 22 | Mental state inference > syllogistic reasoning (belief and desire conjunction) | 26.14 (21-35) |
| Adams et al. | 2010 | fMRI | Explicit | Non-verbal | visual | high | 28 | mental state > gender judgement | ? (18-27) |
| Aichhorn et al. | 2009 | fMRI | Explicit | Verbal | visual | high | 21 | false beliefs > false photographs (timepoint 1 story) | 24 (21-41) |
| | | | | | | | | false beliefs > false photographs (timepoint 2 question) | |
| Alderson-Day et al. | 2016 | fMRI | Explicit | Non-verbal | visual | high | 21 | ToM > physical causality | 24.38 (?-?) |
| Bahnemann et al. | 2010 | fMRI | Explicit | Non-verbal | visual | high | 25 | mental state inference > gender judgements | 26 (?-?) |
| Baron-Cohen et al. | 1999 | fMRI | Explicit | Non-verbal | visual | high | 12 | mental state > gender judgement | 25.5 (?-?) |
| Bartholomeusz et al. | 2018 | fMRI | Explicit | Non-verbal | visual | high | 22 | ToM > physical causality | ? (15-25) |
| Bliksted et al. | 2019 | fMRI | Implicit | Non-verbal | visual | high | 17 | intentional > random movement | 23.59 (?-?) |
| Bodden et al. | 2013 | fMRI | Explicit | Non-verbal | visual | high | 30 | affective ToM > physical | 25.3 (?-?) |
| | | | | | | | | cognitive ToM > physical | |
| Briend et al. | 2019 | fMRI | Explicit | Verbal | auditory | low | 28 | known (ToM) > unknown (non-ToM) language | 37.62 (?-?) |
| Brune et al. | 2008 | fMRI | Explicit | Non-verbal | visual | high | 13 | correct (ToM) > jumbled (non-ToM) cartoon sequence | 26.46 (22-38) |
| Brunet et al. | 2000 | PET | Explicit | Non-verbal | visual | high | 8 | ToM > physical causality | 23.3 (?-?) |
| Canessa et al. | 2012 | fMRI | Implicit | Non-verbal | visual | high | 27 | cooperative social interactions > landscapes | 24.9 & 26.3 |
| | | | | | | | | affective social interactions > landscapes | |
| Cassidy et al. | 2020 | fMRI | Explicit | Verbal | visual | high | 40 | false belief > false photograph | 21.58 (18-33) |
| Castelli et al. | 2000 | PET | Explicit/ Implicit | Non-verbal | visual | high | 6 | intentional > random movement | 24.5 (20-31) |

| | | | | | | | | | |
|---|---|---|---|---|---|---|---|---|---|
| Castelli et al. | 2010 | fMRI | Explicit | Non-verbal | visual | high | 12 | mental state > gender judgement | 25.2 (21-30) |
| Chakroff et al. | 2016 | fMRI | Explicit | Verbal | visual | high | 23 | false belief > false photograph | 27 (?-?) |
| Cheung et al. | 2012 | fMRI | Explicit | Non-verbal | visual | high | 20 | false belief > physical judgements | 23.5 (22-26) |
| | | | | | | | | true belief > physical judgements | |
| | | | | | | | | false belief > physical judgements | |
| | | | | | | | | true belief > physical judgements | |
| Cole et al. | 2019 | fMRI | Explicit | Non-verbal | visual | high | 20 | ToM > non-ToM judgements | 29.6 (?-?) |
| Contreras et al. | 2013 | fMRI | Explicit | Non-verbal | visual | high | 25 | ToM (group) > physical judgements | 22 (19-27) |
| | | | | | | | | ToM(group member) > physical judgements | |
| Contreras et al. | 2013 | fMRI | Explicit | Non-verbal | visual | high | 13 | ToM > physical inference | 19.9 (18-23) |
| | | | | | | | | false belief > false photograph | |
| Corradi-Dell'Acqua et al., | 2014 | fMRI | Explicit | Verbal | visual | high | 46 | mental state > physical judgements | ? (18-31) |
| | | | | | | | | mental state > physical judgements | |
| Das et al. | 2012 | fMRI | Implicit | Non-verbal | visual | high | 19 | ToM > random movement | 33.5 (?-?) |
| de Achaval et al. | 2012 | fMRI | Explicit | Non-verbal | visual | high | 14 | mental state > gender judgement | 28.4 (?-?) |
| | | | | | | | | mental state > gender judgement | |
| Deuse et al. | 2016 | fMRI | Explicit | Non-verbal | visual | high | 38 | social > non-social scenes | 23.88 (18-30) |
| Dodell-Feder et al. | 2011 | fMRI | Explicit | Verbal | visual | high | 62 | false beliefs > false photographs | 22 (18-35) |
| Dodell-Feder et al. | 2014 | fMRI | Explicit | Verbal | visual | high | 18 | stories about thoughts > appearance | ? (15-32) |
| | | | | | | | | stories about thoughts > appearance | |
| Dohnel et al. | 2012 | fMRI | Explicit | Non-verbal | visual | high | 22 | mental state inference > physical judgements | 25.27 (21-40) |
| Dufour et al. | 2013 | fMRI | Explicit | Verbal | BOTH | high | 27 | false beliefs > false photographs | 31 (18-52) |
| Ferstl & von Cramon | 2002 | fMRI | Explicit | Verbal | auditory | high | 9 | ToM stories > pseudo-sentences | 24 (22-27) |
| Fletcher et al. | 1995 | PET | Explicit | Verbal | visual | high | 6 | ToM > physical stories | 38 (24-65) |
| | | | | | | | | ToM stories > unlinked sentences | |
| Focquaert et al | 2010 | fMRI | Explicit | Non-verbal | visual | high | 12 | mental state > gender judgement | 27 (18-38) |
| Focquaert et al. | 2010 | fMRI | Explicit | Non-verbal | visual | high | 12 | mental state > gender judgement | 27 (18-38) |
| Gallagher et al. | 2000 | fMRI | Implicit | Non-verbal | visual | high | 6 | ToM (attribution of false belief/ignorance) > non- ToM (no mental state attribution) cartoons | 30 (23-36) |

| | | | | | | | | | |
|---|---|---|---|---|---|---|---|---|---|
| | | | | | | | | ToM > non-ToM stories | |
| Geiger et al. | 2019 | fMRI | Explicit | Non-verbal | visual | high | 32 | mental state inference > movement identification | 25.75 (?-?) |
| Gobbini et al. | 2007 | fMRI | Explicit | Non-verbal | visual | high | 12 | Intentional social > random movement | 22.2 (?-?) |
| | | | | | | | | false beliefs > physical stories | |
| Gweon et al. | 2012 | fMRI | Explicit | Verbal | auditory | high | 8 | ToM > physical judgements | 21.5 (18-25) |
| Hartwright et al. | 2015 | fMRI | Explicit | Non-verbal | visual | high | 21 | false belief > false photograph | 22 (19-28) |
| Herve et al. | 2013 | fMRI | Explicit | Verbal | visual | | 42 | ToM > semantic judgements | 30.9 (18-53) |
| Hooker et al. | 2008 | fMRI | Explicit | Non-verbal | visual | high | 20 | infer emotion > recognize emotion (false belief trials) | 21 (19-26) |
| Hooker et al. | 2010 | fMRI | Explicit | Non-verbal | visual | high | 15 | change in the protagonist's mental state > no change | 21 (18-25) |
| Jack & Pelphre | 2015 | fMRI | Implicit | Non-verbal | visual | high | 34 | ToM > random movement | 25.73 (20-34) |
| Jacoby et al. | 2016 | fMRI | Explicit | Verbal | visual | high | 17 | false belief > false photograph | 25.3 (18-39) |
| | | | | | | | | ToM > pain events | |
| Jenkins & Mitchell | 2010 | fMRI | Explicit | Verbal | visual | high | 15 | ToM > physical inference | 19.8 (18-22) |
| Jenkins et al. | 2014 | | Implicit | Verbal | visual | high | 19 | ToM > nonToM control stories | 21 (19-25) |
| | | | | | | | | false belief > false photograph | |
| Jimura et al. | 2010 | fMRI | Explicit | Verbal | visual | high | 34 | ToM > factual judgements | ? (20-28) |
| Kana et al. | 2009 | fMRI | Explicit | Non-verbal | visual | high | 12 | Intentional social > random movement | 24.4 (?-?) |
| Kandylaki et al. | 2015 | fMRI | Explicit | Verbal | auditory | high | 20 | false belief > physical causality stories | 24.3 (?-?) |
| | | | | | | | | ToM > physical judgements | |
| Kanske et al. | 2015 | fMRI | Explicit | Non-verbal | visual | high | 25 | ToM > factual reasoning | 32.6 (?-?) |
| | | | | | | | | false belief > false photograph | |
| Kirkovski et al. | 2016 | fMRI | Explicit | Non-verbal | visual | high | 23 | ToM > random movement | ? (19-56) |
| | | | | | | | | ToM > goal-directed movement | |
| | | | | | | rest | | ToM > rest | |
| Kliemann et al. | 2008 | fMRI | Explicit | Verbal | visual | high | 26 | false beliefs > false photographs | ? (19-33) |
| Kobayashi et al. | 2006 | fMRI | Explicit | Verbal | visual | high | 16 | false belief > physical causality | 28.42 (18-49) |
| Kobayashi et al. | 2006 | fMRI | Explicit | Verbal | visual | high | 16 | false belief > physical causality | 28.42 (18-49) age reported for merged groups |
| Koelkebeck et al. | 2011 | fMRI | Explicit | Non-verbal | visual | high | 15 | Intentional social > random movement | 30.9 (?-?) |

| | | | | | | | | | |
|---|---|---|---|---|---|---|---|---|---|
| Lavoie et al. | 2016 | fMRI | Explicit | Verbal | visual | high | 19 | ToM > physical inference (action description phase) | 28.84 (20-53) |
| | | | | | | | | ToM > physical inference (contextual information phase) | |
| | | | | | | | | ToM > physical inference (response phase) | |
| Lee & McCarthy | 2016 | fMRI | Explicit | Verbal | visual | high | 19 | false belief > false photograph | 24.2 (?-?) |
| Lewis et al. | 2017 | fMRI | Explicit | Verbal | visual | high | 17 | ToM > factual judgements | 22 (?-?) |
| Libero et al. | 2014 | fMRI | Explicit | Non-verbal | visual | high | 22 | intention > means judgement | 24.9 (19-36) |
| Lin et al. | 2018 | fMRI | Explicit | Verbal | visual | high | 39 | false belief > false photograph | 22.2 (?-?) |
| Malhi et al. | 2008 | fMRI | Implicit | Non-verbal | visual | high | 20 | Intentional social > random movement | 35.8 (22.56) |
| Marjoram et al. | 2006 | fMRI | Explicit | Non-verbal | visual | high | 13 | ToM> non-ToM cartoons | 29.6 (?-?) |
| Martin & Weisberg | 2003 | fMRI | Explicit | Non-verbal | visual | high | 12 | social interaction > mechanical movement | 27.5 (23.34) |
| Mason et al. | 2008 | fMRI | Explicit | Verbal | visual | rest | 18 | ToM inference > rest | 27.4 (?-?) |
| | | | | | | rest | | ToM inference > rest | |
| McAdams & Krawczyk | 2013 | fMRI | Explicit | Non-verbal | visual | high | 17 | Intentional social > random movement | 24.5 (18-39) |
| Mier et al. | 2010 | fMRI | Explicit | Non-verbal | visual | high | 16 | Mental state inference > emotion categorization | 37.0 (?-?) |
| | | | | | | rest | | Mental state inference > rest | |
| Mitchell | 2008 | fMRI | Explicit | Verbal | visual | high | 20 | false beliefs > false photographs | 23 (19-29) |
| Modinos et al. | 2010 | fMRI | Explicit | Non-verbal | visual | high | 36 | ToM > physical | |
| Moessnang et al. | 2016 | fMRI | Explicit | Non-verbal | visual | high | 46 | ToM > goal-directed movement | 24.7 (?-?) |
| | | | | | | | | ToM > random movement | |
| Mohnke et al. | 2016 | fMRI | Explicit | Non-verbal | visual | high | 29 7 | ToM > physical judgements | 33.58 (?-?) |
| Moran et al. | 2012 | fMRI | Implicit | Non-verbal | visual | high | 31 | Intentional social > random movement | 23 (?-?) |
| | | | | | | | | false belief > false photograph | |
| Naughtin et al. | 2017 | fMRI | Explicit | Verbal | visual | high | 22 | false belief > false photograph | 24 (?-?) |
| Nieminen-von Wendt et al. | 2003 | PET | Explicit | Verbal | auditory | high | 8 | ToM > physical stories | 28.1 (19.2-38.2) |
| Oliver et al. | 2018 | fMRI | Explicit | Verbal | visual | high | 35 | false belief > false photograph | 21.5 (18-26) |
| Otsuka et al. | 2009 | fMRI | Explicit | Verbal | visual | high | 22 | mental state > tense judgement | 24 (19-31) |
| | | | | | | rest | | mental state judgement > rest | |

| | | | | | | | | | |
|---|---|---|---|---|---|---|---|---|---|
| Perner et al. | 2007 | fMRI | Explicit | Verbal | visual | high | 19 | false belief > false photograph | 25 (19-34) |
| | | | | | | | | false belief > temporal change | |
| Platek et al. | 2004 | fMRI | Explicit | Non-verbal | visual | low | 5 | face stimuli > checkerboard | ? (?-?) |
| Powell et al. | 2017 | fMRI | Explicit | Non-verbal | visual | high | 12 | ToM > physical causality | 36.42 (20-58) |
| Roser et al. | 2012 | fMRI | Explicit | Non-verbal | visual | high | 14 | correct (ToM) > jumbled (non-ToM) cartoon sequence | 27.3 (?-?) |
| Ross & Olson | 2010 | fMRI | Explicit | Non-verbal | visual | high | 15 | Intentional social > random movement | 27.86 & 26.25 (?-?) |
| Russel et al | 2000 | fMRI | Explicit | Non-verbal | visual | high | 7 | mental state > gender judgement | 40 (26-58) |
| Saft et al. | 2013 | fMRI | Explicit | Non-verbal | visual | high | 26 | correct (ToM) > jumbled (non-ToM) cartoon sequence | 28.8 (?-?) |
| Samson et al. | 2008 | fMRI | Implicit | Non-verbal | visual | high | 17 | ToM > non-ToM visual puns cartoons | 26.06 (?-?) |
| | | | | | | | | ToM > non-ToM semantic cartoons | |
| Saxe et al. | 2006 | fMRI | Explicit | Verbal | visual | high | 12 | false belief > false photograph | ? (18-26) |
| Saxe & Kaniwisher | 2003 | fMRI | Implicit | Verbal | visual | high | 25 | false belief > mechanical inference stories | ? (?-?) |
| Saxe & Kaniwisher | 2003 | fMRI | Explicit | Verbal | visual | high | 28 | false beliefs > false photographs | ? (?-?) |
| Saxe & Powell | 2006 | fMRI | Explicit | Verbal | visual | high | 12 | false beliefs > false photographs | ? (19-26) |
| Schiffer et al. | 2013 | fMRI | Explicit | Non-verbal | visual | high | 22 | mental state > gender judgement | 35.6 (?-?) |
| Schlaffke et al. | 2014 | | Explicit | Non-verbal | visual | high | 39 | correct (ToM) > jumbled (non-ToM) cartoon sequence | 25.9 (?-?) |
| | | | | | | | | correct (ToM) > jumbled (non-ToM) cartoon sequence | |
| Schmitgen et al. | 2016 | fMRI | Explicit | Non-verbal | visual | high | 21 | ToM > physical judgements | 23.73 (?-?) |
| Schneider et al. | 2014 | fMRI | Explicit | Non-verbal | visual | high | 16 | false belief > false photograph | 22.70 (?-?) |
| Shimada et al. | 2018 | fMRI | Explicit | Non-verbal | visual | high | 30 | ToM > gender judgement | 35.3 (27-43) |
| Sommer et al. | 2010 | fMRI | Explicit | BOTH | visual | high | 14 | ToM inference > physical judgement | 26 (23-37) |
| Specht & Wigglesworth | 2018 | fMRI | Explicit | Non-verbal | visual | high | 18 | ToM > physical causality | 25.7 (21-29) |
| Spunt & Adolphs | 2014 | fMRI | Explicit | Non-verbal | visual | high | 29 | why > how inferences | 27.10 (19-38) |
| Spunt & Adolphs | 2014 | fMRI | Explicit | Non-verbal | visual | high | 21 | why > how inferences | 27.62 (19-38) |
| Spunt & Lieberman | 2012 a | fMRI | Explicit | BOTH | visual | high | 21 | why > how judgements (conjunction video & text) | 21.7 (19-32) |
| Spunt & Lieberman | 2012 b | fMRI | Explicit | Non-verbal | visual | high | 22 | why > how judgements | 21.59 (19-32) |

| | | | | | | | | | |
|---|---|---|---|---|---|---|---|---|---|
| Spunt et al. | 2011 | fMRI | Explicit | Non-verbal | visual | high | 15 | why > how judgements | 19.47 (?-?) |
| | | | | | | | | why > what judgements | |
| Tholen et al. | 2020 | fMRI | Explicit | Non-verbal | visual | high | 130 | ToM > factual reasoning | 40.4 (?-?) |
| Thye et al. | 2018 | fMRI | Explicit | Non-verbal | visual | high | 18 | mental state > gender judgement | 20.15 (18-24) |
| | | | | | | | | mental state > gender judgement | |
| | | | | | | | | ToM > physical causality | |
| Van der Meer et al. | 2011 | fMRI | Explicit | Non-verbal | visual | high | 19 | mental state > physical inference | 21.6 (?-?) |
| | | | | | | rest | | mental state inference > rest | |
| Van Hoeck et al. | 2014 | fMRI | Explicit | Verbal | visual | high | 19 | false belief > physical inference | 22 (19-29) |
| Vanderwal et al | 2008 | fMRI | Explicit | Non-verbal | visual | high | 17 | Intentional social > random movement | 26.4 (?-?) |
| Vogeley et al. | 2001 | fMRI | Explicit | Verbal | visual | high | 8 | ToM > non-ToM stories | ? (25-36) |
| | | | | | | | | ToM (self not in the story) > unlinked senentences | |
| | | | | | | | | ToM (self as one of the agents in the story) > unlinked sentences | |
| Vollm et al. | 2006 | fMRI | Explicit | Non-verbal | visual | high | 13 | ToM > physical causality | 24.9 (19-36) |
| Walter et al. | 2004 | fMRI | Explicit | Non-verbal | visual | high | 13 | communicative intention > physical causality | 25.15 (20-28) |
| | | | | | | | | private intention (one agent) > physical causality | |
| | | | | | | | | private intention (two agents) > physical causality | |
| Walter et al. | 2004 | fMRI | Explicit | Non-verbal | visual | high | 12 | private intention > physical causality | 24.75 (19-27) |
| | | | | | | | | prospective intention > physical causality | |
| | | | | | | | | communicative intention > physical causality | |
| Walter et al. | 2009 | fMRI | Explicit | Non-verbal | visual | high | 12 | private intention > physical causality | 24.75 (?-?) |
| | | | | | | | | prospective social intention > physical causality | |
| | | | | | | | | communicative intention > physical causality | |
| Wang et al. | 2015 | fMRI | Explicit | Non-verbal | visual | high | 56 | ToM > physical causality | 19.3 (17-21) |
| Willert et al. | 2015 | fMRI | Explicit | Non-verbal | visual | high | 81 | ToM > physical judgements | 35.57 (?-?) |

| | | | | | | | | | |
|---|---|---|---|---|---|---|---|---|---|
| Wolf et al. | 2010 | fMRI | Explicit | BOTH | BOTH | high | 18 | ToM > non-ToM scenes (movie watching phase)<br>ToM > physical inferences (response phase 1)<br>ToM > physical inferences (response phase 2) | 31.94 (20-45) |
| Young et al. | 2011 | fMRI | Explicit | Verbal | visual | high | 17 | false beliefs > false photographs | ? (18-22) |
| Zaitchik et al. | 2010 | fMRI | Explicit | Verbal | visual | high | 15 | stories about beliefs > non-ToM stories<br>stories about emotions > non-ToM stories | 22.4 (20-24) |

**The references to access the listed studies are listed at the end of the document*

***Table M2 ToM*** *List of studies included in the* ***theory of mind MAIN high and low baselines (rest excluded)*** *(N = ) meta-analysis.* ***Note:*** *This data set was used for the* ***MAIN*** *ToM & SC conjunction and contrast analyses. The list of excluded studies using rest as a baseline can be seen at the end of the document highlighted in grey. ToM= Theory of Mind, SC= Semantic Cognition, N= Sample Size*

| *Authors | Year | Imaging Method | Instructional Cue | Stimulus Domain | Sensory Input Modality | Baseline Type | N | Contrast |
|---|---|---|---|---|---|---|---|---|
| Abraham et al. | 2008 | fMRI | Explicit | Verbal | visual | high | 17 | stories about mental states > control stories |
| Abraham et al. | 2010 | fMRI | Explicit | Verbal | visual | high | 22 | Mental state inference > syllogistic reasoning (belief and desire conjunction) |
| Adams et al. | 2010 | fMRI | Explicit | Non-verbal | visual | high | 28 | mental state > gender judgement |
| Aichhorn et al. | 2009 | fMRI | Explicit | Verbal | visual | high | 21 | false beliefs > false photographs (timepoint 1 story) |
| | | | | | | | | false beliefs > false photographs (timepoint 2 question) |
| Alderson-Day et al. | 2016 | fMRI | Explicit | Non-verbal | visual | high | 21 | ToM > physical causality |
| Bahnemann et al. | 2010 | fMRI | Explicit | Non-verbal | visual | high | 25 | mental state inference > gender judgements |
| Baron-Cohen et al. | 1999 | fMRI | Explicit | Non-verbal | visual | high | 12 | mental state > gender judgement |
| Bartholomeusz et al. | 2018 | fMRI | Explicit | Non-verbal | visual | high | 22 | ToM > physical causality |
| Bliksted et al. | 2019 | fMRI | Implicit | Non-verbal | visual | high | 17 | intentional > random movement |
| Bodden et al. | 2013 | fMRI | Explicit | Non-verbal | visual | high | 30 | affective ToM > physical |
| | | | | | | | | cognitive ToM > physical |
| Briend et al. | 2019 | fMRI | Explicit | Verbal | auditory | low | 28 | known (ToM) > unknown (non-ToM) language |
| Brune et al. | 2008 | fMRI | Explicit | Non-verbal | visual | high | 13 | correct (ToM) > jumbled (non-ToM) cartoon sequence |
| Brunet et al. | 2000 | PET | Explicit | Non-verbal | visual | high | 8 | ToM > physical causality |
| Canessa et al. | 2012 | fMRI | Implicit | Non-verbal | visual | high | 27 | cooperative social interactions > landscapes |
| | | | | | | | | affective social interactions > landscapes |
| Cassidy et al. | 2020 | fMRI | Explicit | Verbal | visual | high | 40 | false belief > false photograph |
| Castelli et al. | 2000 | PET | Explicit/Implicit | Non-verbal | visual | high | 6 | intentional > random movement |
| Castelli et al. | 2010 | fMRI | Explicit | Non-verbal | visual | high | 12 | mental state > gender judgement |
| Chakroff et al. | 2016 | fMRI | Explicit | Verbal | visual | high | 23 | false belief > false photograph |
| Cheung et al. | 2012 | fMRI | Explicit | Non-verbal | visual | high | 20 | false belief > physical judgements |
| | | | | | | | | true belief > physical judgements |
| | | | | | | | | false belief > physical judgements |

| | | | | | | | | |
|---|---|---|---|---|---|---|---|---|
| | | | | | | | | true belief > physical judgements |
| Cole et al. | 2019 | fMRI | Explicit | Non-verbal | visual | high | 20 | ToM > non-ToM judgements |
| Contreras et al. | 2013 | fMRI | Explicit | Non-verbal | visual | high | 25 | ToM (group) > physical judgements |
| | | | | | | | | ToM(group member) > physical judgements |
| Contreras et al. | 2013 | fMRI | Explicit | Non-verbal | visual | high | 13 | ToM > physical inference |
| | | | | | | | | false belief > false photograph |
| Corradi-Dell'Acqua et al., | 2014 | fMRI | Explicit | Verbal | visual | high | 46 | mental state > physical judgements |
| | | | | | | | | mental state > physical judgements |
| Das et al. | 2012 | fMRI | Implicit | Non-verbal | visual | high | 19 | ToM > random movement |
| de Achaval et al. | 2012 | fMRI | Explicit | Non-verbal | visual | high | 14 | mental state > gender judgement |
| | | | | | | | | mental state > gender judgement |
| Deuse et al. | 2016 | fMRI | Explicit | Non-verbal | visual | high | 38 | social > non-social scenes |
| Dodell-Feder et al. | 2011 | fMRI | Explicit | Verbal | visual | high | 62 | false beliefs > false photographs |
| Dodell-Feder et al. | 2014 | fMRI | Explicit | Verbal | visual | high | 18 | stories about thoughts > appearance |
| | | | | | | | | stories about thoughts > appearance |
| Dohnel et al. | 2012 | fMRI | Explicit | Non-verbal | visual | high | 22 | mental state inference > physical judgements |
| Dufour et al. | 2013 | fMRI | Explicit | Verbal | BOTH | high | 27 | false beliefs > false photographs |
| Ferstl & von Cramon | 2002 | fMRI | Explicit | Verbal | auditory | high | 9 | ToM stories > pseudo-sentences |
| Fletcher et al. | 1995 | PET | Explicit | Verbal | visual | high | 6 | ToM > physical stories |
| | | | | | | | | ToM stories > unlinked sentences |
| Focquaert et al | 2010 | fMRI | Explicit | Non-verbal | visual | high | 12 | mental state > gender judgement |
| Focquaert et al. | 2010 | fMRI | Explicit | Non-verbal | visual | high | 12 | mental state > gender judgement |
| Gallagher et al. | 2000 | fMRI | Implicit | Non-verbal | visual | high | 6 | ToM (attribution of false belief/ignorance) > non- ToM (no mental state attribution) cartoons |
| | | | | | | | | ToM > non-ToM stories |
| Geiger et al. | 2019 | fMRI | Explicit | Non-verbal | visual | high | 32 | mental state inference > movement identification |
| Gobbini et al. | 2007 | fMRI | Explicit | Non-verbal | visual | high | 12 | Intentional social > random movement |
| | | | | | | | | false beliefs > physical stories |
| Gweon et al. | 2012 | fMRI | Explicit | Verbal | auditory | high | 8 | ToM > physical judgements |
| Hartwright et al. | 2015 | fMRI | Explicit | Non-verbal | visual | high | 21 | false belief > false photograph |
| Herve et al. | 2013 | fMRI | Explicit | Verbal | visual | high | 42 | ToM > semantic judgements |
| Hooker et al. | 2008 | fMRI | Explicit | Non-verbal | visual | high | 20 | infer emotion > recognize emotion (false belief trials) |

| | | | | | | | | |
|---|---|---|---|---|---|---|---|---|
| Hooker et al. | 2010 | fMRI | Explicit | Non-verbal | visual | high | 15 | change in the protagonist's mental state > no change |
| Jack & Pelphre | 2015 | fMRI | Implicit | Non-verbal | visual | high | 34 | ToM > random movement |
| Jacoby et al. | 2016 | fMRI | Explicit | Verbal | visual | high | 17 | false belief > false photograph |
| | | | | | | | | ToM > pain events |
| Jenkins & Mitchell | 2010 | fMRI | Explicit | Verbal | visual | high | 15 | ToM > physical inference |
| Jenkins et al. | 2014 | | Implicit | Verbal | visual | high | 19 | ToM > nonToM control stories |
| | | | | | | | | false belief > false photograph |
| Jimura et al. | 2010 | fMRI | Explicit | Verbal | visual | high | 34 | ToM > factual judgements |
| Kana et al. | 2009 | fMRI | Explicit | Non-verbal | visual | high | 12 | Intentional social > random movement |
| Kandylaki et al. | 2015 | fMRI | Explicit | Verbal | auditory | high | 20 | false belief > physical causality stories |
| | | | | | | | | ToM > physical judgements |
| Kanske et al. | 2015 | fMRI | Explicit | Non-verbal | visual | high | 25 | ToM > factual reasoning |
| | | | | | | | | false belief > false photograph |
| Kirkovski et al. | 2016 | fMRI | Explicit | Non-verbal | visual | high | 23 | ToM > random movement |
| | | | | | | | | ToM > goal-directed movement |
| Kliemann et al. | 2008 | fMRI | Explicit | Verbal | visual | high | 26 | false beliefs > false photographs |
| Kobayashi et al. | 2006 | fMRI | Explicit | Verbal | visual | high | 16 | false belief > physical causality |
| Kobayashi et al. | 2006 | fMRI | Explicit | Verbal | visual | high | 16 | false belief > physical causality |
| Koelkebeck et al. | 2011 | fMRI | Explicit | Non-verbal | visual | high | 15 | Intentional social > random movement |
| Lavoie et al. | 2016 | fMRI | Explicit | Verbal | visual | high | 19 | ToM > physical inference (action description phase) |
| | | | | | | | | ToM > physical inference (contextual information phase) |
| | | | | | | | | ToM > physical inference (response phase) |
| Lee & McCarthy | 2016 | fMRI | Explicit | Verbal | visual | high | 19 | false belief > false photograph |
| Lewis et al. | 2017 | fMRI | Explicit | Verbal | visual | high | 17 | ToM > factual judgements |
| Libero et al. | 2014 | fMRI | Explicit | Non-verbal | visual | high | 22 | intention > means judgement |
| Lin et al. | 2018 | fMRI | Explicit | Verbal | visual | high | 39 | false belief > false photograph |
| Malhi et al. | 2008 | fMRI | Implicit | Non-verbal | visual | high | 20 | Intentional social > random movement |
| Marjoram et al. | 2006 | fMRI | Explicit | Non-verbal | visual | high | 13 | ToM> non-ToM cartoons |
| Martin & Weisberg | 2003 | fMRI | Explicit | Non-verbal | visual | high | 12 | social interaction > mechanical movement |
| McAdams & Krawczyk | 2013 | fMRI | Explicit | Non-verbal | visual | high | 17 | Intentional social > random movement |
| Mier et al. | 2010 | fMRI | Explicit | Non-verbal | visual | high | 16 | Mental state inference > emotion categorization |

| | | | | | | | | |
|---|---|---|---|---|---|---|---|---|
| Mitchell | 2008 | fMRI | Explicit | Verbal | visual | high | 20 | false beliefs > false photographs |
| Modinos et al. | 2010 | fMRI | Explicit | Non-verbal | visual | high | 36 | ToM > physical |
| Moessnang et al. | 2016 | fMRI | Explicit | Non-verbal | visual | high | 46 | ToM > goal-directed movement |
| | | | | | | | | ToM > random movement |
| Mohnke et al. | 2016 | fMRI | Explicit | Non-verbal | visual | high | 297 | ToM > physical judgements |
| Moran et al. | 2012 | fMRI | Implicit | Non-verbal | visual | high | 31 | Intentional social > random movement |
| | | | | | | | | false belief > false photograph |
| Naughtin et al. | 2017 | fMRI | Explicit | Verbal | visual | high | 22 | false belief > false photograph |
| Nieminen-von Wendt et al. | 2003 | PET | Explicit | Verbal | auditory | high | 8 | ToM > physical stories |
| Oliver et al. | 2018 | fMRI | Explicit | Verbal | visual | high | 35 | false belief > false photograph |
| Otsuka et al. | 2009 | fMRI | Explicit | Verbal | visual | high | 22 | mental state > tense judgement |
| Perner et al. | 2007 | fMRI | Explicit | Verbal | visual | high | 19 | false belief > false photograph |
| | | | | | | | | false belief > temporal change |
| Platek et al. | 2004 | fMRI | Explicit | Non-verbal | visual | low | 5 | face stimuli > checkerboard |
| Powell et al. | 2017 | fMRI | Explicit | Non-verbal | visual | high | 12 | ToM > physical causality |
| Roser et al. | 2012 | fMRI | Explicit | Non-verbal | visual | high | 14 | correct (ToM) > jumbled (non-ToM) cartoon sequence |
| Ross & Olson | 2010 | fMRI | Explicit | Non-verbal | visual | high | 15 | Intentional social > random movement |
| Russel et al | 2000 | fMRI | Explicit | Non-verbal | visual | high | 7 | mental state > gender judgement |
| Saft et al. | 2013 | fMRI | Explicit | Non-verbal | visual | high | 26 | correct (ToM) > jumbled (non-ToM) cartoon sequence |
| Samson et al. | 2008 | fMRI | Implicit | Non-verbal | visual | high | 17 | ToM > non-ToM visual puns cartoons |
| | | | | | | | | ToM > non-ToM semantic cartoons |
| Saxe et al. | 2006 | fMRI | Explicit | Verbal | visual | high | 12 | false belief > false photograph |
| Saxe & Kaniwisher | 2003 | fMRI | Implicit | Verbal | visual | high | 25 | false belief > mechanical inference stories |
| Saxe & Kaniwisher | 2003 | fMRI | Explicit | Verbal | visual | high | 28 | false beliefs > false photographs |
| Saxe & Powell | 2006 | fMRI | Explicit | Verbal | visual | high | 12 | false beliefs > false photographs |
| Schiffer et al. | 2013 | fMRI | Explicit | Non-verbal | visual | high | 22 | mental state > gender judgement |
| Schlaffke et al. | 2014 | | Explicit | Non-verbal | visual | high | 39 | correct (ToM) > jumbled (non-ToM) cartoon sequence |
| | | | | | | | | correct (ToM) > jumbled (non-ToM) cartoon sequence |
| Schmitgen et al. | 2016 | fMRI | Explicit | Non-verbal | visual | high | 21 | ToM > physical judgements |
| Schneider et al. | 2014 | fMRI | Explicit | Non-verbal | visual | high | 16 | false belief > false photograph |
| Shimada et al. | 2018 | fMRI | Explicit | Non-verbal | visual | high | 30 | ToM > gender judgement |

| | | | | | | | | |
|---|---|---|---|---|---|---|---|---|
| Sommer et al. | 2010 | fMRI | Explicit | BOTH | visual | high | 14 | ToM inference > physical judgement |
| Specht & Wigglesworth | 2018 | fMRI | Explicit | Non-verbal | visual | high | 18 | ToM > physical causality |
| Spunt & Adolphs | 2014 | fMRI | Explicit | Non-verbal | visual | high | 29 | why > how inferences |
| Spunt & Adolphs | 2014 | fMRI | Explicit | Non-verbal | visual | high | 21 | why > how inferences |
| Spunt & Lieberman | 2012a | fMRI | Explicit | BOTH | visual | high | 21 | why > how judgements (conjunction video & text) |
| Spunt & Lieberman | 2012b | fMRI | Explicit | Non-verbal | visual | high | 22 | why > how judgements |
| Spunt et al. | 2011 | fMRI | Explicit | Non-verbal | visual | high | 15 | why > how judgements |
| | | | | | | | | why > what judgements |
| Tholen et al. | 2020 | fMRI | Explicit | Non-verbal | visual | high | 130 | ToM > factual reasoning |
| Thye et al. | 2018 | fMRI | Explicit | Non-verbal | visual | high | 18 | mental state > gender judgement |
| | | | | | | | | mental state > gender judgement |
| | | | | | | | | ToM > physical causality |
| Van der Meer et al. | 2011 | fMRI | Explicit | Non-verbal | visual | high | 19 | mental state > physical inference |
| Van Hoeck et al. | 2014 | fMRI | Explicit | Verbal | visual | high | 19 | false belief > physical inference |
| Vanderwal et al | 2008 | fMRI | Explicit | Non-verbal | visual | high | 17 | Intentional social > random movement |
| Vogeley et al. | 2001 | fMRI | Explicit | Verbal | visual | high | 8 | ToM > non-ToM stories |
| | | | | | | | | ToM (self not in the story) > unlinked senentences |
| | | | | | | | | ToM (self as one of the agents in the story) > unlinked sentences |
| Vollm et al. | 2006 | fMRI | Explicit | Non-verbal | visual | high | 13 | ToM > physical causality |
| Walter et al. | 2004 | fMRI | Explicit | Non-verbal | visual | high | 13 | communicative intention > physical causality |
| | | | | | | | | private intention (one agent) > physical causality |
| | | | | | | | | private intention (two agents) > physical causality |
| Walter et al. | 2004 | fMRI | Explicit | Non-verbal | visual | high | 12 | private intention > physical causality |
| | | | | | | | | prospective intention > physical causality |
| | | | | | | | | communicative intention > physical causality |
| Walter et al. | 2009 | fMRI | Explicit | Non-verbal | visual | high | 12 | private intention > physical causality |
| | | | | | | | | prospective social intention > physical causality |
| | | | | | | | | communicative intention > physical causality |
| Wang et al. | 2015 | fMRI | Explicit | Non-verbal | visual | high | 56 | ToM > physical causality |
| Willert et al. | 2015 | fMRI | Explicit | Non-verbal | visual | high | 81 | ToM > physical judgements |
| Wolf et al. | 2010 | fMRI | Explicit | BOTH | BOTH | high | 18 | ToM > non-ToM scenes (movie watching phase) |

| | | | | | | | | |
|---|---|---|---|---|---|---|---|---|
| | | | | | | | | ToM > physical inferences (response phase 1) |
| | | | | | | | | ToM > physical inferences (response phase 2) |
| Young et al. | 2011 | fMRI | Explicit | Verbal | visual | high | 17 | false beliefs > false photographs |
| Zaitchik et al. | 2010 | fMRI | Explicit | Verbal | visual | high | 15 | stories about beliefs > non-ToM stories |
| | | | | | | | | stories about emotions > non-ToM stories |
| EXCLUDED CONTRASTS WITH REST | | | | | | | | |
| Kirkovski et al. | 2016 | | Explicit | Non-verbal | visual | rest | 23 | ToM > rest |
| Mason et al. | 2008 | fMRI | Explicit | Verbal | visual | rest | 18 | ToM inference > rest |
| | | | | | | | | ToM inference > rest |
| Mier et al. | 2010 | | Explicit | Non-verbal | visual | rest | 16 | Mental state inference > rest |
| Otsuka et al. | 2009 | | Explicit | Verbal | visual | rest | 22 | mental state judgement > rest |
| Van der Meer et al. | 2011 | | Explicit | Non-verbal | visual | rest | 19 | mental state inference > rest |

**The references to access the listed studies are listed at the end of the document*

***Table M3 ToM*** *List of studies included in the* ***theory of mind high baselines only (low baselines and rest excluded*** *(N = 111) meta-analysis.* ***Note:*** *The list of excluded studies using rest and low-level baselines can be seen at the end of the document highlighted in grey. ToM= Theory of Mind, N= Sample Size*

| *Authors | Year | Imaging Method | Instructional Cue | Stimulus Domain | Sensory Input Modality | Baseline Type | N | Contrast |
|---|---|---|---|---|---|---|---|---|
| Abraham et al. | 2008 | fMRI | Explicit | Verbal | visual | high | 17 | stories about mental states > control stories |
| Abraham et al. | 2010 | fMRI | Explicit | Verbal | visual | high | 22 | Mental state inference > syllogistic reasoning (belief and desire conjunction) |
| Adams et al. | 2010 | fMRI | Explicit | Non-verbal | visual | high | 28 | mental state > gender judgement |
| Aichhorn et al. | 2009 | fMRI | Explicit | Verbal | visual | high | 21 | false beliefs > false photographs (timepoint 1 story) |
| | | | | | | | | false beliefs > false photographs (timepoint 2 question) |
| Alderson-Day et al. | 2016 | fMRI | Explicit | Non-verbal | visual | high | 21 | ToM > physical causality |
| Bahnemann et al. | 2010 | fMRI | Explicit | Non-verbal | visual | high | 25 | mental state inference > gender judgements |
| Baron-Cohen et al. | 1999 | fMRI | Explicit | Non-verbal | visual | high | 12 | mental state > gender judgement |
| Bartholomeusz et al. | 2018 | fMRI | Explicit | Non-verbal | visual | high | 22 | ToM > physical causality |
| Bliksted et al. | 2019 | fMRI | Implicit | Non-verbal | visual | high | 17 | intentional > random movement |
| Bodden et al. | 2013 | fMRI | Explicit | Non-verbal | visual | high | 30 | affective ToM > physical |
| | | | | | | | | cognitive ToM > physical |
| Brune et al. | 2008 | fMRI | Explicit | Non-verbal | visual | high | 13 | correct (ToM) > jumbled (non-ToM) cartoon sequence |
| Brunet et al. | 2000 | PET | Explicit | Non-verbal | visual | high | 8 | ToM > physical causality |
| Canessa et al. | 2012 | fMRI | Implicit | Non-verbal | visual | high | 27 | cooperative social interactions > landscapes |
| | | | | | | | | affective social interactions > landscapes |
| Cassidy et al. | 2020 | fMRI | Explicit | Verbal | visual | high | 40 | false belief > false photograph |
| Castelli et al. | 2000 | PET | Explicit/Implicit | Non-verbal | visual | high | 6 | intentional > random movement |
| Castelli et al. | 2010 | fMRI | Explicit | Non-verbal | visual | high | 12 | mental state > gender judgement |
| Chakroff et al. | 2016 | fMRI | Explicit | Verbal | visual | high | 23 | false belief > false photograph |
| Cheung et al. | 2012 | fMRI | Explicit | Non-verbal | visual | high | 20 | false belief > physical judgements |
| | | | | | | | | true belief > physical judgements |
| | | | | | | | | false belief > physical judgements |
| | | | | | | | | true belief > physical judgements |

| | | | | | | | | |
|---|---|---|---|---|---|---|---|---|
| Cole et al. | 2019 | fMRI | Explicit | Non-verbal | visual | high | 20 | ToM > non-ToM judgements |
| Contreras et al. | 2013 | fMRI | Explicit | Non-verbal | visual | high | 25 | ToM (group) > physical judgements |
| | | | | | | | | ToM(group member) > physical judgements |
| Contreras et al. | 2013 | fMRI | Explicit | Non-verbal | visual | high | 13 | ToM > physical inference |
| | | | | | | | | false belief > false photograph |
| Corradi-Dell'Acqua et al., | 2014 | fMRI | Explicit | Verbal | visual | high | 46 | mental state > physical judgements |
| | | | | | | | | mental state > physical judgements |
| Das et al. | 2012 | fMRI | Implicit | Non-verbal | visual | high | 19 | ToM > random movement |
| de Achaval et al. | 2012 | fMRI | Explicit | Non-verbal | visual | high | 14 | mental state > gender judgement |
| | | | | | | | | mental state > gender judgement |
| Deuse et al. | 2016 | fMRI | Explicit | Non-verbal | visual | high | 38 | social > non-social scenes |
| Dodell-Feder et al. | 2011 | fMRI | Explicit | Verbal | visual | high | 62 | false beliefs > false photographs |
| Dodell-Feder et al. | 2014 | fMRI | Explicit | Verbal | visual | high | 18 | stories about thoughts > appearance |
| | | | | | | | | stories about thoughts > appearance |
| Dohnel et al. | 2012 | fMRI | Explicit | Non-verbal | visual | high | 22 | mental state inference > physical judgements |
| Dufour et al. | 2013 | fMRI | Explicit | Verbal | BOTH | high | 27 | false beliefs > false photographs |
| Ferstl & von Cramon | 2002 | fMRI | Explicit | Verbal | auditory | high | 9 | ToM stories > pseudo-sentences |
| Fletcher et al. | 1995 | PET | Explicit | Verbal | visual | high | 6 | ToM > physical stories |
| | | | | | | | | ToM stories > unlinked sentences |
| Focquaert et al | 2010 | fMRI | Explicit | Non-verbal | visual | high | 12 | mental state > gender judgement |
| Focquaert et al. | 2010 | fMRI | Explicit | Non-verbal | visual | high | 12 | mental state > gender judgement |
| Gallagher et al. | 2000 | fMRI | Implicit | Non-verbal | visual | high | 6 | ToM (attribution of false belief/ignorance) > non- ToM (no mental state attribution) cartoons |
| | | | | | | | | ToM > non-ToM stories |
| Geiger et al. | 2019 | fMRI | Explicit | Non-verbal | visual | high | 32 | mental state inference > movement identification |
| Gobbini et al. | 2007 | fMRI | Explicit | Non-verbal | visual | high | 12 | Intentional social > random movement |
| | | | | | | | | false beliefs > physical stories |
| Gweon et al. | 2012 | fMRI | Explicit | Verbal | auditory | high | 8 | ToM > physical judgements |
| Hartwright et al. | 2015 | fMRI | Explicit | Non-verbal | visual | high | 21 | false belief > false photograph |
| Herve et al. | 2013 | fMRI | Explicit | Verbal | visual | high | 42 | ToM > semantic judgements |
| Hooker et al. | 2008 | fMRI | Explicit | Non-verbal | visual | high | 20 | infer emotion > recognize emotion (false belief trials) |
| Hooker et al. | 2010 | fMRI | Explicit | Non-verbal | visual | high | 15 | change in the protagonist's mental state > no change |

| | | | | | | | | |
|---|---|---|---|---|---|---|---|---|
| Jack & Pelphre | 2015 | fMRI | Implicit | Non-verbal | visual | high | 34 | ToM > random movement |
| Jacoby et al. | 2016 | fMRI | Explicit | Verbal | visual | high | 17 | false belief > false photograph |
| | | | | | | | | ToM > pain events |
| Jenkins & Mitchell | 2010 | fMRI | Explicit | Verbal | visual | high | 15 | ToM > physical inference |
| Jenkins et al. | 2014 | | Implicit | Verbal | visual | high | 19 | ToM > nonToM control stories |
| | | | | | | | | false belief > false photograph |
| Jimura et al. | 2010 | fMRI | Explicit | Verbal | visual | high | 34 | ToM > factual judgements |
| Kana et al. | 2009 | fMRI | Explicit | Non-verbal | visual | high | 12 | Intentional social > random movement |
| Kandylaki et al. | 2015 | fMRI | Explicit | Verbal | auditory | high | 20 | false belief > physical causality stories |
| | | | | | | | | ToM > physical judgements |
| Kanske et al. | 2015 | fMRI | Explicit | Non-verbal | visual | high | 25 | ToM > factual reasoning |
| | | | | | | | | false belief > false photograph |
| Kirkovski et al. | 2016 | fMRI | Explicit | Non-verbal | visual | high | 23 | ToM > random movement |
| | | | | | | | | ToM > goal-directed movement |
| Kliemann et al. | 2008 | fMRI | Explicit | Verbal | visual | high | 26 | false beliefs > false photographs |
| Kobayashi et al. | 2006 | fMRI | Explicit | Verbal | visual | high | 16 | false belief > physical causality |
| Kobayashi et al. | 2006 | fMRI | Explicit | Verbal | visual | high | 16 | false belief > physical causality |
| Koelkebeck et al. | 2011 | fMRI | Explicit | Non-verbal | visual | high | 15 | Intentional social > random movement |
| Lavoie et al. | 2016 | fMRI | Explicit | Verbal | visual | high | 19 | ToM > physical inference (action description phase) |
| | | | | | | | | ToM > physical inference (contextual information phase) |
| | | | | | | | | ToM > physical inference (response phase) |
| Lee & McCarthy | 2016 | fMRI | Explicit | Verbal | visual | high | 19 | false belief > false photograph |
| Lewis et al. | 2017 | fMRI | Explicit | Verbal | visual | high | 17 | ToM > factual judgements |
| Libero et al. | 2014 | fMRI | Explicit | Non-verbal | visual | high | 22 | intention > means judgement |
| Lin et al. | 2018 | fMRI | Explicit | Verbal | visual | high | 39 | false belief > false photograph |
| Malhi et al. | 2008 | fMRI | Implicit | Non-verbal | visual | high | 20 | Intentional social > random movement |
| Marjoram et al. | 2006 | fMRI | Explicit | Non-verbal | visual | high | 13 | ToM> non-ToM cartoons |
| Martin & Weisberg | 2003 | fMRI | Explicit | Non-verbal | visual | high | 12 | social interaction > mechanical movement |
| McAdams & Krawczyk | 2013 | fMRI | Explicit | Non-verbal | visual | high | 17 | Intentional social > random movement |
| Mier et al. | 2010 | fMRI | Explicit | Non-verbal | visual | high | 16 | Mental state inference > emotion categorization |
| Mitchell | 2008 | fMRI | Explicit | Verbal | visual | high | 20 | false beliefs > false photographs |

| | | | | | | | | |
|---|---|---|---|---|---|---|---|---|
| Modinos et al. | 2010 | fMRI | Explicit | Non-verbal | visual | high | 36 | ToM > physical |
| Moessnang et al. | 2016 | fMRI | Explicit | Non-verbal | visual | high | 46 | ToM > goal-directed movement |
| | | | | | | | | ToM > random movement |
| Mohnke et al. | 2016 | fMRI | Explicit | Non-verbal | visual | high | 297 | ToM > physical judgements |
| Moran et al. | 2012 | fMRI | Implicit | Non-verbal | visual | high | 31 | Intentional social > random movement |
| | | | | | | | | false belief > false photograph |
| Naughtin et al. | 2017 | fMRI | Explicit | Verbal | visual | high | 22 | false belief > false photograph |
| Nieminen-von Wendt et al. | 2003 | PET | Explicit | Verbal | auditory | high | 8 | ToM > physical stories |
| Oliver et al. | 2018 | fMRI | Explicit | Verbal | visual | high | 35 | false belief > false photograph |
| Otsuka et al. | 2009 | fMRI | Explicit | Verbal | visual | high | 22 | mental state > tense judgement |
| Perner et al. | 2007 | fMRI | Explicit | Verbal | visual | high | 19 | false belief > false photograph |
| | | | | | | | | false belief > temporal change |
| Powell et al. | 2017 | fMRI | Explicit | Non-verbal | visual | high | 12 | ToM > physical causality |
| Roser et al. | 2012 | fMRI | Explicit | Non-verbal | visual | high | 14 | correct (ToM) > jumbled (non-ToM) cartoon sequence |
| Ross & Olson | 2010 | fMRI | Explicit | Non-verbal | visual | high | 15 | Intentional social > random movement |
| Russel et al | 2000 | fMRI | Explicit | Non-verbal | visual | high | 7 | mental state > gender judgement |
| Saft et al. | 2013 | fMRI | Explicit | Non-verbal | visual | high | 26 | correct (ToM) > jumbled (non-ToM) cartoon sequence |
| Samson et al. | 2008 | fMRI | Implicit | Non-verbal | visual | high | 17 | ToM > non-ToM visual puns cartoons |
| | | | | | | | | ToM > non-ToM semantic cartoons |
| Saxe et al. | 2006 | fMRI | Explicit | Verbal | visual | high | 12 | false belief > false photograph |
| Saxe & Kaniwisher | 2003 | fMRI | Implicit | Verbal | visual | high | 25 | false belief > mechanical inference stories |
| Saxe & Kaniwisher | 2003 | fMRI | Explicit | Verbal | visual | high | 28 | false beliefs > false photographs |
| Saxe & Powell | 2006 | fMRI | Explicit | Verbal | visual | high | 12 | false beliefs > false photographs |
| Schiffer et al. | 2013 | fMRI | Explicit | Non-verbal | visual | high | 22 | mental state > gender judgement |
| Schlaffke et al. | 2014 | | Explicit | Non-verbal | visual | high | 39 | correct (ToM) > jumbled (non-ToM) cartoon sequence |
| | | | | | | | | correct (ToM) > jumbled (non-ToM) cartoon sequence |
| Schmitgen et al. | 2016 | fMRI | Explicit | Non-verbal | visual | high | 21 | ToM > physical judgements |
| Schneider et al. | 2014 | fMRI | Explicit | Non-verbal | visual | high | 16 | false belief > false photograph |
| Shimada et al. | 2018 | fMRI | Explicit | Non-verbal | visual | high | 30 | ToM > gender judgement |
| Sommer et al. | 2010 | fMRI | Explicit | BOTH | visual | high | 14 | ToM inference > physical judgement |
| Specht & Wigglesworth | 2018 | fMRI | Explicit | Non-verbal | visual | high | 18 | ToM > physical causality |

| | | | | | | | | |
|---|---|---|---|---|---|---|---|---|
| Spunt & Adolphs | 2014 | fMRI | Explicit | Non-verbal | visual | high | 29 | why > how inferences |
| Spunt & Adolphs | 2014 | fMRI | Explicit | Non-verbal | visual | high | 21 | why > how inferences |
| Spunt & Lieberman | 2012a | fMRI | Explicit | BOTH | visual | high | 21 | why > how judgements (conjunction video & text) |
| Spunt & Lieberman | 2012b | fMRI | Explicit | Non-verbal | visual | high | 22 | why > how judgements |
| Spunt et al. | 2011 | fMRI | Explicit | Non-verbal | visual | high | 15 | why > how judgements |
| | | | | | | | | why > what judgements |
| Tholen et al. | 2020 | fMRI | Explicit | Non-verbal | visual | high | 130 | ToM > factual reasoning |
| Thye et al. | 2018 | fMRI | Explicit | Non-verbal | visual | high | 18 | mental state > gender judgement |
| | | | | | | | | mental state > gender judgement |
| | | | | | | | | ToM > physical causality |
| Van der Meer et al. | 2011 | fMRI | Explicit | Non-verbal | visual | high | 19 | mental state > physical inference |
| Van Hoeck et al. | 2014 | fMRI | Explicit | Verbal | visual | high | 19 | false belief > physical inference |
| Vanderwal et al | 2008 | fMRI | Explicit | Non-verbal | visual | high | 17 | Intentional social > random movement |
| Vogeley et al. | 2001 | fMRI | Explicit | Verbal | visual | high | 8 | ToM > non-ToM stories |
| | | | | | | | | ToM (self not in the story) > unlinked senentences |
| | | | | | | | | ToM (self as one of the agents in the story) > unlinked sentences |
| Vollm et al. | 2006 | fMRI | Explicit | Non-verbal | visual | high | 13 | ToM > physical causality |
| Walter et al. | 2004 | fMRI | Explicit | Non-verbal | visual | high | 13 | communicative intention > physical causality |
| | | | | | | | | private intention (one agent) > physical causality |
| | | | | | | | | private intention (two agents) > physical causality |
| Walter et al. | 2004 | fMRI | Explicit | Non-verbal | visual | high | 12 | private intention > physical causality |
| | | | | | | | | prospective intention > physical causality |
| | | | | | | | | communicative intention > physical causality |
| Walter et al. | 2009 | fMRI | Explicit | Non-verbal | visual | high | 12 | private intention > physical causality |
| | | | | | | | | prospective social intention > physical causality |
| | | | | | | | | communicative intention > physical causality |
| Wang et al. | 2015 | fMRI | Explicit | Non-verbal | visual | high | 56 | ToM > physical causality |
| Willert et al. | 2015 | fMRI | Explicit | Non-verbal | visual | high | 81 | ToM > physical judgements |
| Wolf et al. | 2010 | fMRI | Explicit | BOTH | BOTH | high | 18 | ToM > non-ToM scenes (movie watching phase) |
| | | | | | | | | ToM > physical inferences (response phase 1) |
| | | | | | | | | ToM > physical inferences (response phase 2) |

| | | | | | | | | |
|---|---|---|---|---|---|---|---|---|
| Young et al. | 2011 | fMRI | Explicit | Verbal | visual | high | 17 | false beliefs > false photographs |
| Zaitchik et al. | 2010 | fMRI | Explicit | Verbal | visual | high | 15 | stories about beliefs > non-ToM stories |
| | | | | | | | | stories about emotions > non-ToM stories |
| EXCLUDED CONTRASTS WITH REST AND LOW LEVEL BASELINES | | | | | | | | |
| Briend et al. | 2019 | fMRI | Explicit | Verbal | auditory | low | 28 | known (ToM) > unknown (non-ToM) language |
| Kirkovski et al. | 2016 | | Explicit | Non-verbal | visual | rest | 23 | ToM > rest |
| Mason et al. | 2008 | fMRI | Explicit | Verbal | visual | rest | 18 | ToM inference > rest |
| | | | | | | | | ToM inference > rest |
| Mier et al. | 2010 | | Explicit | Non-verbal | visual | rest | 16 | Mental state inference > rest |
| Otsuka et al. | 2009 | | Explicit | Verbal | visual | rest | 22 | mental state judgement > rest |
| Platek et al. | 2004 | fMRI | Explicit | Non-verbal | visual | low | 5 | face stimuli > checkerboard |
| Van der Meer et al. | 2011 | | Explicit | Non-verbal | visual | rest | 19 | mental state inference > rest |

**The references to access the listed studies are listed at the end of the document*

***Table M2.1 ToM VERBAL*** *List of studies included in the* ***theory of mind VERBAL STIMULUS DOMAIN*** *(N = 46) meta-analysis after excluding NON-VERBAL contrasts.* ***Note:*** *The list of excluded studies with NON-VERBAL STIMULUS DOMAIN can be seen in Table M.2.2 ToM NON-VERBAL. Contrasts with BOTH VEBAL and NON-VERBAL stimuli were also excluded and are listed at the end of the document highlighted in grey. ToM= Theory of Mind, N= Sample Size*

| *Authors | Year | Imaging Method | Instructional Cue | Stimulus Domain | Sensory Input Modality | Baseline Type | N | Contrast |
|---|---|---|---|---|---|---|---|---|
| Abraham et al. | 2008 | fMRI | Explicit | Verbal | visual | high | 17 | stories about mental states > control stories |
| Abraham et al. | 2010 | fMRI | Explicit | Verbal | visual | high | 22 | Mental state inference > syllogistic reasoning (belief and desire conjunction) |
| Aichhorn et al. | 2009 | fMRI | Explicit | Verbal | visual | high | 21 | false beliefs > false photographs (timepoint 1 story) |
| | | | | | | | | false beliefs > false photographs (timepoint 2 question) |
| Briend et al. | 2019 | fMRI | Explicit | Verbal | auditory | low | 28 | known (ToM) > unknown (non-ToM) language |
| Cassidy et al. | 2020 | fMRI | Explicit | Verbal | visual | high | 40 | false belief > false photograph |
| Chakroff et al. | 2016 | fMRI | Explicit | Verbal | visual | high | 23 | false belief > false photograph |
| Cheung et al. | 2012 | fMRI | Explicit | verbal | visual | high | 20 | false belief > physical judgements |
| | | | | verbal | | | | true belief > physical judgements |
| Contreras et al. | 2013 | fMRI | Explicit | verbal | visual | high | 13 | false belief > false photograph |
| Corradi-Dell’Acqua et al., | 2014 | fMRI | Explicit | Verbal | visual | high | 46 | mental state > physical judgements |
| | | | | | | | | mental state > physical judgements |
| Dodell-Feder et al. | 2011 | fMRI | Explicit | Verbal | visual | high | 62 | false beliefs > false photographs |
| Dodell-Feder et al. | 2014 | fMRI | Explicit | Verbal | visual | high | 18 | stories about thoughts > appearance |
| | | | | | | | | stories about thoughts > appearance |
| Dufour et al. | 2013 | fMRI | Explicit | Verbal | BOTH | high | 27 | false beliefs > false photographs |
| Ferstl & von Cramon | 2002 | fMRI | Explicit | Verbal | auditory | high | 9 | ToM stories > pseudo-sentences |
| Fletcher et al. | 1995 | PET | Explicit | Verbal | visual | high | 6 | ToM > physical stories |
| | | | | | | | | ToM stories > unlinked sentences |
| Gallagher et al. | 2000 | fMRI | Implicit | verbal | visual | high | 6 | ToM > non-ToM stories |
| Gobbini et al. | 2007 | fMRI | Explicit | verbal | visual | high | 12 | false beliefs > physical stories |
| Gweon et al. | 2012 | fMRI | Explicit | Verbal | auditory | high | 8 | ToM > physical judgements |
| Herve et al. | 2013 | fMRI | Explicit | Verbal | visual | high | 42 | ToM > semantic judgements |

| | | | | | | | | |
|---|---|---|---|---|---|---|---|---|
| Jacoby et al. | 2016 | fMRI | Explicit | Verbal | visual | high | 17 | false belief > false photograph |
| Jenkins & Mitchell | 2010 | fMRI | Explicit | Verbal | visual | high | 15 | ToM > physical inference |
| Jenkins et al. | 2014 | | Implicit | Verbal | visual | high | 19 | ToM > nonToM control stories |
| | | | | | | | | false belief > false photograph |
| Jimura et al. | 2010 | fMRI | Explicit | Verbal | visual | high | 34 | ToM > factual judgements |
| Kandylaki et al. | 2015 | fMRI | Explicit | Verbal | auditory | high | 20 | false belief > physical causality stories |
| | | | | | | | | ToM > physical judgements |
| Kliemann et al. | 2008 | fMRI | Explicit | Verbal | visual | high | 26 | false beliefs > false photographs |
| Kobayashi et al. | 2006 | fMRI | Explicit | Verbal | visual | high | 16 | false belief > physical causality |
| Kobayashi et al. | 2006 | fMRI | Explicit | Verbal | visual | high | 16 | false belief > physical causality |
| Lavoie et al. | 2016 | fMRI | Explicit | Verbal | visual | high | 19 | ToM > physical inference (action description phase) |
| | | | | | | | | ToM > physical inference (contextual information phase) |
| | | | | | | | | ToM > physical inference (response phase) |
| Lee & McCarthy | 2016 | fMRI | Explicit | Verbal | visual | high | 19 | false belief > false photograph |
| Lewis et al. | 2017 | fMRI | Explicit | Verbal | visual | high | 17 | ToM > factual judgements |
| Lin et al. | 2018 | fMRI | Explicit | Verbal | visual | high | 39 | false belief > false photograph |
| Mitchell | 2008 | fMRI | Explicit | Verbal | visual | high | 20 | false beliefs > false photographs |
| Moran et al. | 2012 | fMRI | Implicit | verbal | visual | high | 31 | false belief > false photograph |
| Naughtin et al. | 2017 | fMRI | Explicit | Verbal | visual | high | 22 | false belief > false photograph |
| Nieminen-von Wendt et al. | 2003 | PET | Explicit | Verbal | auditory | high | 8 | ToM > physical stories |
| Oliver et al. | 2018 | fMRI | Explicit | Verbal | visual | high | 35 | false belief > false photograph |
| Otsuka et al. | 2009 | fMRI | Explicit | Verbal | visual | high | 22 | mental state > tense judgement |
| Perner et al. | 2007 | fMRI | Explicit | Verbal | visual | high | 19 | false belief > false photograph |
| | | | | | | | | false belief > temporal change |
| Saxe et al. | 2006 | fMRI | Explicit | Verbal | visual | high | 12 | false belief > false photograph |
| Saxe & Kaniwisher | 2003 | fMRI | Implicit | Verbal | visual | high | 25 | false belief > mechanical inference stories |
| Saxe & Kaniwisher | 2003 | fMRI | Explicit | Verbal | visual | high | 28 | false beliefs > false photographs |
| Saxe & Powell | 2006 | fMRI | Explicit | Verbal | visual | high | 12 | false beliefs > false photographs |
| Thye et al. | 2018 | fMRI | Explicit | verbal | visual | high | 18 | mental state > gender judgement |
| Van Hoeck et al. | 2014 | fMRI | Explicit | Verbal | visual | high | 19 | false belief > physical inference |
| Vogeley et al. | 2001 | fMRI | Explicit | Verbal | visual | high | 8 | ToM > non-ToM stories |

| | | | | | | | | |
|---|---|---|---|---|---|---|---|---|
| | | | | | | | | ToM (self not in the story) > unlinked senentences |
| | | | | | | | | ToM (self as one of the agents in the story) > unlinked sentences |
| Young et al. | 2011 | fMRI | Explicit | Verbal | visual | high | 17 | false beliefs > false photographs |
| Zaitchik et al. | 2010 | fMRI | Explicit | Verbal | visual | high | 15 | stories about beliefs > non-ToM stories |
| | | | | | | | | stories about emotions > non-ToM stories |
| EXCLUDED CONTRASTS CONTAINING BOTH VERBAL AND NON VERBAL STIMULI. THE EXLUDED NON VERBAL EXPERIMENTS CAN BE SEEN IN Table M.2.2 ToM NON-VERBAL | | | | | | | | |
| Sommer et al. | 2010 | fMRI | Explicit | BOTH | visual | high | 14 | ToM inference > physical judgement |
| Spunt & Lieberman | 2012a | fMRI | Explicit | BOTH | visual | high | 21 | why > how judgements (conjunction video & text) |
| Wolf et al. | 2010 | fMRI | Explicit | BOTH | BOTH | high | 18 | ToM > non-ToM scenes (movie watching phase) |
| | | | | | | | | ToM > physical inferences (response phase 1) |
| | | | | | | | | ToM > physical inferences (response phase 2) |

**The references to access the listed studies are listed at the end of the document*

***Table M2.2 ToM NON-VERBAL*** *List of studies included in the* ***theory of mind NON-VERBAL STIMULUS DOMAIN*** *(N = 71) meta-analysis after excluding VERBAL contrasts.* ***Note:*** *The list of excluded studies with VERBAL STIMULUS DOMAIN and contrasts with BOTH VEBAL and NON-VERBAL stimuli that were also excluded can be seen in Table M.2.1 ToM VERBAL. ToM= Theory of Mind, N= Sample Size*

| *Authors | Year | Imaging Method | Instructional Cue | Stimulus Domain | Sensory Input Modality | Baseline Type | N | Contrast |
|---|---|---|---|---|---|---|---|---|
| Adams et al. | 2010 | fMRI | Explicit | Non-verbal | visual | high | 28 | mental state > gender judgement |
| Alderson-Day et al. | 2016 | fMRI | Explicit | Non-verbal | visual | high | 21 | ToM > physical causality |
| Bahnemann et al. | 2010 | fMRI | Explicit | Non-verbal | visual | high | 25 | mental state inference > gender judgements |
| Baron-Cohen et al. | 1999 | fMRI | Explicit | Non-verbal | visual | high | 12 | mental state > gender judgement |
| Bartholomeusz et al. | 2018 | fMRI | Explicit | Non-verbal | visual | high | 22 | ToM > physical causality |
| Bliksted et al. | 2019 | fMRI | Implicit | Non-verbal | visual | high | 17 | intentional > random movement |
| Bodden et al. | 2013 | fMRI | Explicit | Non-verbal | visual | high | 30 | affective ToM > physical |
| | | | | | | | | cognitive ToM > physical |
| Brune et al. | 2008 | fMRI | Explicit | Non-verbal | visual | high | 13 | correct (ToM) > jumbled (non-ToM) cartoon sequence |
| Brunet et al. | 2000 | PET | Explicit | Non-verbal | visual | high | 8 | ToM > physical causality |
| Canessa et al. | 2012 | fMRI | Implicit | Non-verbal | visual | high | 27 | cooperative social interactions > landscapes |
| | | | | | | | | affective social interactions > landscapes |
| Castelli et al. | 2000 | PET | Explicit/Implicit | Non-verbal | visual | high | 6 | intentional > random movement |
| Castelli et al. | 2010 | fMRI | Explicit | Non-verbal | visual | high | 12 | mental state > gender judgement |
| Cheung et al. | 2012 | fMRI | Explicit | Non-verbal | visual | high | 20 | false belief > physical judgements |
| | | | | | | | | true belief > physical judgements |
| Cole et al. | 2019 | fMRI | Explicit | Non-verbal | visual | high | 20 | ToM > non-ToM judgements |
| Contreras et al. | 2013 | fMRI | Explicit | Non-verbal | visual | high | 25 | ToM (group) > physical judgements |
| | | | | | | | | ToM(group member) > physical judgements |
| Contreras et al. | 2013 | fMRI | Explicit | Non-verbal | visual | high | 13 | ToM > physical inference |
| Das et al. | 2012 | fMRI | Implicit | Non-verbal | visual | high | 19 | ToM > random movement |
| de Achaval et al. | 2012 | fMRI | Explicit | Non-verbal | visual | high | 14 | mental state > gender judgement |
| | | | | | | | | mental state > gender judgement |
| Deuse et al. | 2016 | fMRI | Explicit | Non-verbal | visual | high | 38 | social > non-social scenes |

| | | | | | | | | |
|---|---|---|---|---|---|---|---|---|
| Dohnel et al. | 2012 | fMRI | Explicit | Non-verbal | visual | high | 22 | mental state inference > physical judgements |
| Focquaert et al | 2010 | fMRI | Explicit | Non-verbal | visual | high | 12 | mental state > gender judgement |
| Focquaert et al. | 2010 | fMRI | Explicit | Non-verbal | visual | high | 12 | mental state > gender judgement |
| Gallagher et al. | 2000 | fMRI | Implicit | Non-verbal | visual | high | 6 | ToM (attribution of false belief/ignorance) > non- ToM (no mental state attribution) cartoons |
| Geiger et al. | 2019 | fMRI | Explicit | Non-verbal | visual | high | 32 | mental state inference > movement identification |
| Gobbini et al. | 2007 | fMRI | Explicit | Non-verbal | visual | high | 12 | Intentional social > random movement |
| Hartwright et al. | 2015 | fMRI | Explicit | Non-verbal | visual | high | 21 | false belief > false photograph |
| Hooker et al. | 2008 | fMRI | Explicit | Non-verbal | visual | high | 20 | infer emotion > recognize emotion (false belief trials) |
| Hooker et al. | 2010 | fMRI | Explicit | Non-verbal | visual | high | 15 | change in the protagonist's mental state > no change |
| Jack & Pelphre | 2015 | fMRI | Implicit | Non-verbal | visual | high | 34 | ToM > random movement |
| Jacoby et al. | 2016 | fMRI | Explicit | Non-verbal | visual | high | 17 | ToM > pain events |
| Kana et al. | 2009 | fMRI | Explicit | Non-verbal | visual | high | 12 | Intentional social > random movement |
| Kanske et al. | 2015 | fMRI | Explicit | Non-verbal | visual | high | 25 | ToM > factual reasoning |
| | | | | | | | | false belief > false photograph |
| Kirkovski et al. | 2016 | fMRI | Explicit | Non-verbal | visual | high | 23 | ToM > random movement |
| | | | | | | | | ToM > goal-directed movement |
| Koelkebeck et al. | 2011 | fMRI | Explicit | Non-verbal | visual | high | 15 | Intentional social > random movement |
| Libero et al. | 2014 | fMRI | Explicit | Non-verbal | visual | high | 22 | intention > means judgement |
| Malhi et al. | 2008 | fMRI | Implicit | Non-verbal | visual | high | 20 | Intentional social > random movement |
| Marjoram et al. | 2006 | fMRI | Explicit | Non-verbal | visual | high | 13 | ToM> non-ToM cartoons |
| Martin & Weisberg | 2003 | fMRI | Explicit | Non-verbal | visual | high | 12 | social interaction > mechanical movement |
| McAdams & Krawczyk | 2013 | fMRI | Explicit | Non-verbal | visual | high | 17 | Intentional social > random movement |
| Mier et al. | 2010 | fMRI | Explicit | Non-verbal | visual | high | 16 | Mental state inference > emotion categorization |
| Modinos et al. | 2010 | fMRI | Explicit | Non-verbal | visual | high | 36 | ToM > physical |
| Moessnang et al. | 2016 | fMRI | Explicit | Non-verbal | visual | high | 46 | ToM > goal-directed movement |
| | | | | | | | | ToM > random movement |
| Mohnke et al. | 2016 | fMRI | Explicit | Non-verbal | visual | high | 29 7 | ToM > physical judgements |
| Moran et al. | 2012 | fMRI | Implicit | Non-verbal | visual | high | 31 | Intentional social > random movement |
| Platek et al. | 2004 | fMRI | Explicit | Non-verbal | visual | low | 5 | face stimuli > checkerboard |
| Powell et al. | 2017 | fMRI | Explicit | Non-verbal | visual | high | 12 | ToM > physical causality |

| | | | | | | | | |
|---|---|---|---|---|---|---|---|---|
| Roser et al. | 2012 | fMRI | Explicit | Non-verbal | visual | high | 14 | correct (ToM) > jumbled (non-ToM) cartoon sequence |
| Ross & Olson | 2010 | fMRI | Explicit | Non-verbal | visual | high | 15 | Intentional social > random movement |
| Russel et al | 2000 | fMRI | Explicit | Non-verbal | visual | high | 7 | mental state > gender judgement |
| Saft et al. | 2013 | fMRI | Explicit | Non-verbal | visual | high | 26 | correct (ToM) > jumbled (non-ToM) cartoon sequence |
| Samson et al. | 2008 | fMRI | Implicit | Non-verbal | visual | high | 17 | ToM > non-ToM visual puns cartoons |
| | | | | | | | | ToM > non-ToM semantic cartoons |
| Schiffer et al. | 2013 | fMRI | Explicit | Non-verbal | visual | high | 22 | mental state > gender judgement |
| Schlaffke et al. | 2014 | | Explicit | Non-verbal | visual | high | 39 | correct (ToM) > jumbled (non-ToM) cartoon sequence |
| | | | | | | | | correct (ToM) > jumbled (non-ToM) cartoon sequence |
| Schmitgen et al. | 2016 | fMRI | Explicit | Non-verbal | visual | high | 21 | ToM > physical judgements |
| Schneider et al. | 2014 | fMRI | Explicit | Non-verbal | visual | high | 16 | false belief > false photograph |
| Shimada et al. | 2018 | fMRI | Explicit | Non-verbal | visual | high | 30 | ToM > gender judgement |
| Specht & Wigglesworth | 2018 | fMRI | Explicit | Non-verbal | visual | high | 18 | ToM > physical causality |
| Spunt & Adolphs | 2014 | fMRI | Explicit | Non-verbal | visual | high | 29 | why > how inferences |
| Spunt & Adolphs | 2014 | fMRI | Explicit | Non-verbal | visual | high | 21 | why > how inferences |
| Spunt & Lieberman | 2012b | fMRI | Explicit | Non-verbal | visual | high | 22 | why > how judgements |
| Spunt et al. | 2011 | fMRI | Explicit | Non-verbal | visual | high | 15 | why > how judgements |
| | | | | | | | | why > what judgements |
| Tholen et al. | 2020 | fMRI | Explicit | Non-verbal | visual | high | 130 | ToM > factual reasoning |
| Thye et al. | 2018 | fMRI | Explicit | Non-verbal | visual | high | 18 | mental state > gender judgement |
| | | | | | | | | ToM > physical causality |
| Van der Meer et al. | 2011 | fMRI | Explicit | Non-verbal | visual | high | 19 | mental state > physical inference |
| Vanderwal et al | 2008 | fMRI | Explicit | Non-verbal | visual | high | 17 | Intentional social > random movement |
| Vollm et al. | 2006 | fMRI | Explicit | Non-verbal | visual | high | 13 | ToM > physical causality |
| Walter et al. | 2004 | fMRI | Explicit | Non-verbal | visual | high | 13 | communicative intention > physical causality |
| | | | | | | | | private intention (one agent) > physical causality |
| | | | | | | | | private intention (two agents) > physical causality |
| Walter et al. | 2004 | fMRI | Explicit | Non-verbal | visual | high | 12 | private intention > physical causality |
| | | | | | | | | prospective intention > physical causality |
| | | | | | | | | communicative intention > physical causality |

| | | | | | | | | |
|---|---|---|---|---|---|---|---|---|
| Walter et al. | 2009 | fMRI | Explicit | Non-verbal | visual | high | 12 | private intention > physical causality |
| | | | | | | | | prospective social intention > physical causality |
| | | | | | | | | communicative intention > physical causality |
| Wang et al. | 2015 | fMRI | Explicit | Non-verbal | visual | high | 56 | ToM > physical causality |
| Willert et al. | 2015 | fMRI | Explicit | Non-verbal | visual | high | 81 | ToM > physical judgements |
| EXCLUDED CONTRASTS CONTAINING BOTH VERBAL AND NON VERBAL STIMULI. THE EXLUDED VERBAL EXPERIMENTS CAN BE SEEN IN Table M.2.1 ToM VERBAL | | | | | | | | |
| Sommer et al. | 2010 | fMRI | Explicit | BOTH | visual | high | 14 | ToM inference > physical judgement |
| Spunt & Lieberman | 2012a | fMRI | Explicit | BOTH | visual | high | 21 | why > how judgements (conjunction video & text) |
| Wolf et al. | 2010 | fMRI | Explicit | BOTH | BOTH | high | 18 | ToM > non-ToM scenes (movie watching phase) |
| | | | | | | | | ToM > physical inferences (response phase 1) |
| | | | | | | | | ToM > physical inferences (response phase 2) |

**The references to access the listed studies are listed at the end of the document*

***Table M2.3 ToM VISUAL*** *List of studies included in the* ***theory of mind VISUAL INPUT MODALITY*** *(N =106 ) meta-analysis after excluding AUDITORY contrasts.* ***Note:*** *The list of excluded studies with AUDITORY INPUT MODALITY and contrasts with BOTH VISUAL and AUDITORY stimuli that were also excluded are listed at the end of the document highlighted in grey. Note that it was not possible to run and independent ALE analysis on the ToM AUDITORY contrasts due to a very small number of these contrasts in the ToM data. ToM= Theory of Mind, N= Sample Size*

| *Authors | Year | Imaging Method | Instructional Cue | Stimulus Domain | Sensory Input Modality | Baseline Type | N | Contrast |
|---|---|---|---|---|---|---|---|---|
| Abraham et al. | 2008 | fMRI | Explicit | Verbal | visual | high | 17 | stories about mental states > control stories |
| Abraham et al. | 2010 | fMRI | Explicit | Verbal | visual | high | 22 | Mental state inference > syllogistic reasoning (belief and desire conjunction) |
| Adams et al. | 2010 | fMRI | Explicit | Non-verbal | visual | high | 28 | mental state > gender judgement |
| Aichhorn et al. | 2009 | fMRI | Explicit | Verbal | visual | high | 21 | false beliefs > false photographs (timepoint 1 story) |
| | | | | | | | | false beliefs > false photographs (timepoint 2 question) |
| Alderson-Day et al. | 2016 | fMRI | Explicit | Non-verbal | visual | high | 21 | ToM > physical causality |
| Bahnemann et al. | 2010 | fMRI | Explicit | Non-verbal | visual | high | 25 | mental state inference > gender judgements |
| Baron-Cohen et al. | 1999 | fMRI | Explicit | Non-verbal | visual | high | 12 | mental state > gender judgement |
| Bartholomeusz et al. | 2018 | fMRI | Explicit | Non-verbal | visual | high | 22 | ToM > physical causality |
| Bliksted et al. | 2019 | fMRI | Implicit | Non-verbal | visual | high | 17 | intentional > random movement |
| Bodden et al. | 2013 | fMRI | Explicit | Non-verbal | visual | high | 30 | affective ToM > physical |
| | | | | | | | | cognitive ToM > physical |
| Brune et al. | 2008 | fMRI | Explicit | Non-verbal | visual | high | 13 | correct (ToM) > jumbled (non-ToM) cartoon sequence |
| Brunet et al. | 2000 | PET | Explicit | Non-verbal | visual | high | 8 | ToM > physical causality |
| Canessa et al. | 2012 | fMRI | Implicit | Non-verbal | visual | high | 27 | cooperative social interactions > landscapes |
| | | | | | | | | affective social interactions > landscapes |
| Cassidy et al. | 2020 | fMRI | Explicit | Verbal | visual | high | 40 | false belief > false photograph |
| Castelli et al. | 2000 | PET | Explicit/Implicit | Non-verbal | visual | high | 6 | intentional > random movement |
| Castelli et al. | 2010 | fMRI | Explicit | Non-verbal | visual | high | 12 | mental state > gender judgement |
| Chakroff et al. | 2016 | fMRI | Explicit | Verbal | visual | high | 23 | false belief > false photograph |
| Cheung et al. | 2012 | fMRI | Explicit | Non-verbal/Verbal | visual | high | 20 | false belief > physical judgements |

| | | | | | | | | |
|---|---|---|---|---|---|---|---|---|
| | | | | | | | | true belief > physical judgements |
| | | | | | | | | false belief > physical judgements |
| | | | | | | | | true belief > physical judgements |
| Cole et al. | 2019 | fMRI | Explicit | Non-verbal | visual | high | 20 | ToM > non-ToM judgements |
| Contreras et al. | 2013 | fMRI | Explicit | Non-verbal | visual | high | 25 | ToM (group) > physical judgements |
| | | | | | | | | ToM(group member) > physical judgements |
| Contreras et al. | 2013 | fMRI | Explicit | Non-verbal/Verbal | visual | high | 13 | ToM > physical inference |
| | | | | | | | | false belief > false photograph |
| Corradi-Dell'Acqua et al., | 2014 | fMRI | Explicit | Verbal | visual | high | 46 | mental state > physical judgements |
| | | | | | | | | mental state > physical judgements |
| Das et al. | 2012 | fMRI | Implicit | Non-verbal | visual | high | 19 | ToM > random movement |
| de Achaval et al. | 2012 | fMRI | Explicit | Non-verbal | visual | high | 14 | mental state > gender judgement |
| | | | | | | | | mental state > gender judgement |
| Deuse et al. | 2016 | fMRI | Explicit | Non-verbal | visual | high | 38 | social > non-social scenes |
| Dodell-Feder et al. | 2011 | fMRI | Explicit | Verbal | visual | high | 62 | false beliefs > false photographs |
| Dodell-Feder et al. | 2014 | fMRI | Explicit | Verbal | visual | high | 18 | stories about thoughts > appearance |
| | | | | | | | | stories about thoughts > appearance |
| Dohnel et al. | 2012 | fMRI | Explicit | Non-verbal | visual | high | 22 | mental state inference > physical judgements |
| Fletcher et al. | 1995 | PET | Explicit | Verbal | visual | high | 6 | ToM > physical stories |
| | | | | | | | | ToM stories > unlinked sentences |
| Focquaert et al | 2010 | fMRI | Explicit | Non-verbal | visual | high | 12 | mental state > gender judgement |
| Focquaert et al. | 2010 | fMRI | Explicit | Non-verbal | visual | high | 12 | mental state > gender judgement |
| Gallagher et al. | 2000 | fMRI | Implicit | Non-verbal/Verbal | visual | high | 6 | ToM (attribution of false belief/ignorance) > non- ToM (no mental state attribution) cartoons |
| | | | | | | | | ToM > non-ToM stories |
| Geiger et al. | 2019 | fMRI | Explicit | Non-verbal | visual | high | 32 | mental state inference > movement identification |
| Gobbini et al. | 2007 | fMRI | Explicit | Non-verbal/Verbal | visual | high | 12 | Intentional social > random movement |
| | | | | | | | | false beliefs > physical stories |

| | | | | | | | | |
|---|---|---|---|---|---|---|---|---|
| Hartwright et al. | 2015 | fMRI | Explicit | Non-verbal | visual | high | 21 | false belief > false photograph |
| Herve et al. | 2013 | fMRI | Explicit | Verbal | visual | high | 42 | ToM > semantic judgements |
| Hooker et al. | 2008 | fMRI | Explicit | Non-verbal | visual | high | 20 | infer emotion > recognize emotion (false belief trials) |
| Hooker et al. | 2010 | fMRI | Explicit | Non-verbal | visual | high | 15 | change in the protagonist's mental state > no change |
| Jack & Pelphre | 2015 | fMRI | Implicit | Non-verbal | visual | high | 34 | ToM > random movement |
| Jacoby et al. | 2016 | fMRI | Explicit | Verbal/Nonverbal | visual | high | 17 | false belief > false photograph |
| | | | | | | | | ToM > pain events |
| Jenkins & Mitchell | 2010 | fMRI | Explicit | Verbal | visual | high | 15 | ToM > physical inference |
| Jenkins et al. | 2014 | | Implicit | Verbal | visual | high | 19 | ToM > nonToM control stories |
| | | | | | | | | false belief > false photograph |
| Jimura et al. | 2010 | fMRI | Explicit | Verbal | visual | high | 34 | ToM > factual judgements |
| Kana et al. | 2009 | fMRI | Explicit | Non-verbal | visual | high | 12 | Intentional social > random movement |
| Kanske et al. | 2015 | fMRI | Explicit | Non-verbal | visual | high | 25 | ToM > factual reasoning |
| | | | | | | | | false belief > false photograph |
| Kirkovski et al. | 2016 | fMRI | Explicit | Non-verbal | visual | high | 23 | ToM > random movement |
| | | | | | | | | ToM > goal-directed movement |
| Kliemann et al. | 2008 | fMRI | Explicit | Verbal | visual | high | 26 | false beliefs > false photographs |
| Kobayashi et al. | 2006 | fMRI | Explicit | Verbal | visual | high | 16 | false belief > physical causality |
| Kobayashi et al. | 2006 | fMRI | Explicit | Verbal | visual | high | 16 | false belief > physical causality |
| Koelkebeck et al. | 2011 | fMRI | Explicit | Non-verbal | visual | high | 15 | Intentional social > random movement |
| Lavoie et al. | 2016 | fMRI | Explicit | Verbal | visual | high | 19 | ToM > physical inference (action description phase) |
| | | | | | | | | ToM > physical inference (contextual information phase) |
| | | | | | | | | ToM > physical inference (response phase) |
| Lee & McCarthy | 2016 | fMRI | Explicit | Verbal | visual | high | 19 | false belief > false photograph |
| Lewis et al. | 2017 | fMRI | Explicit | Verbal | visual | high | 17 | ToM > factual judgements |
| Libero et al. | 2014 | fMRI | Explicit | Non-verbal | visual | high | 22 | intention > means judgement |
| Lin et al. | 2018 | fMRI | Explicit | Verbal | visual | high | 39 | false belief > false photograph |
| Malhi et al. | 2008 | fMRI | Implicit | Non-verbal | visual | high | 20 | Intentional social > random movement |
| Marjoram et al. | 2006 | fMRI | Explicit | Non-verbal | visual | high | 13 | ToM> non-ToM cartoons |
| Martin & Weisberg | 2003 | fMRI | Explicit | Non-verbal | visual | high | 12 | social interaction > mechanical movement |

| | | | | | | | | |
|---|---|---|---|---|---|---|---|---|
| McAdams & Krawczyk | 2013 | fMRI | Explicit | Non-verbal | visual | high | 17 | Intentional social > random movement |
| Mier et al. | 2010 | fMRI | Explicit | Non-verbal | visual | high | 16 | Mental state inference > emotion categorization |
| Mitchell | 2008 | fMRI | Explicit | Verbal | visual | high | 20 | false beliefs > false photographs |
| Modinos et al. | 2010 | fMRI | Explicit | Non-verbal | visual | high | 36 | ToM > physical |
| Moessnang et al. | 2016 | fMRI | Explicit | Non-verbal | visual | high | 46 | ToM > goal-directed movement |
| | | | | | | | | ToM > random movement |
| Mohnke et al. | 2016 | fMRI | Explicit | Non-verbal | visual | high | 297 | ToM > physical judgements |
| Moran et al. | 2012 | fMRI | Implicit | Non-verbal/Verbal | visual | high | 31 | Intentional social > random movement |
| | | | | | | | | false belief > false photograph |
| Naughtin et al. | 2017 | fMRI | Explicit | Verbal | visual | high | 22 | false belief > false photograph |
| Oliver et al. | 2018 | fMRI | Explicit | Verbal | visual | high | 35 | false belief > false photograph |
| Otsuka et al. | 2009 | fMRI | Explicit | Verbal | visual | high | 22 | mental state > tense judgement |
| Perner et al. | 2007 | fMRI | Explicit | Verbal | visual | high | 19 | false belief > false photograph |
| | | | | | | | | false belief > temporal change |
| Platek et al. | 2004 | fMRI | Explicit | Non-verbal | visual | low | 5 | face stimuli > checkerboard |
| Powell et al. | 2017 | fMRI | Explicit | Non-verbal | visual | high | 12 | ToM > physical causality |
| Roser et al. | 2012 | fMRI | Explicit | Non-verbal | visual | high | 14 | correct (ToM) > jumbled (non-ToM) cartoon sequence |
| Ross & Olson | 2010 | fMRI | Explicit | Non-verbal | visual | high | 15 | Intentional social > random movement |
| Russel et al | 2000 | fMRI | Explicit | Non-verbal | visual | high | 7 | mental state > gender judgement |
| Saft et al. | 2013 | fMRI | Explicit | Non-verbal | visual | high | 26 | correct (ToM) > jumbled (non-ToM) cartoon sequence |
| Samson et al. | 2008 | fMRI | Implicit | Non-verbal | visual | high | 17 | ToM > non-ToM visual puns cartoons |
| | | | | | | | | ToM > non-ToM semantic cartoons |
| Saxe et al. | 2006 | fMRI | Explicit | Verbal | visual | high | 12 | false belief > false photograph |
| Saxe & Kaniwisher | 2003 | fMRI | Implicit | Verbal | visual | high | 25 | false belief > mechanical inference stories |
| Saxe & Kaniwisher | 2003 | fMRI | Explicit | Verbal | visual | high | 28 | false beliefs > false photographs |
| Saxe & Powell | 2006 | fMRI | Explicit | Verbal | visual | high | 12 | false beliefs > false photographs |
| Schiffer et al. | 2013 | fMRI | Explicit | Non-verbal | visual | high | 22 | mental state > gender judgement |
| Schlaffke et al. | 2014 | | Explicit | Non-verbal | visual | high | 39 | correct (ToM) > jumbled (non-ToM) cartoon sequence |
| | | | | | | | | correct (ToM) > jumbled (non-ToM) cartoon sequence |
| Schmitgen et al. | 2016 | fMRI | Explicit | Non-verbal | visual | high | 21 | ToM > physical judgements |

| | | | | | | | | |
|---|---|---|---|---|---|---|---|---|
| Schneider et al. | 2014 | fMRI | Explicit | Non-verbal | visual | high | 16 | false belief > false photograph |
| Shimada et al. | 2018 | fMRI | Explicit | Non-verbal | visual | high | 30 | ToM > gender judgement |
| Sommer et al. | 2010 | fMRI | Explicit | BOTH | visual | high | 14 | ToM inference > physical judgement |
| Specht & Wigglesworth | 2018 | fMRI | Explicit | Non-verbal | visual | high | 18 | ToM > physical causality |
| Spunt & Adolphs | 2014 | fMRI | Explicit | Non-verbal | visual | high | 29 | why > how inferences |
| Spunt & Adolphs | 2014 | fMRI | Explicit | Non-verbal | visual | high | 21 | why > how inferences |
| Spunt & Lieberman | 2012a | fMRI | Explicit | BOTH | visual | high | 21 | why > how judgements (conjunction video & text) |
| Spunt & Lieberman | 2012b | fMRI | Explicit | Non-verbal | visual | high | 22 | why > how judgements |
| Spunt et al. | 2011 | fMRI | Explicit | Non-verbal | visual | high | 15 | why > how judgements |
| | | | | | | | | why > what judgements |
| Tholen et al. | 2020 | fMRI | Explicit | Non-verbal | visual | high | 130 | ToM > factual reasoning |
| Thye et al. | 2018 | fMRI | Explicit | Non-verbal | visual | high | 18 | mental state > gender judgement |
| | | | | | | | | ToM > physical causality |
| Van der Meer et al. | 2011 | fMRI | Explicit | Non-verbal | visual | high | 19 | mental state > physical inference |
| Van Hoeck et al. | 2014 | fMRI | Explicit | Verbal | visual | high | 19 | false belief > physical inference |
| Vanderwal et al | 2008 | fMRI | Explicit | Non-verbal | visual | high | 17 | Intentional social > random movement |
| Vogeley et al. | 2001 | fMRI | Explicit | Verbal | visual | high | 8 | ToM > non-ToM stories |
| | | | | | | | | ToM (self not in the story) > unlinked senentences |
| | | | | | | | | ToM (self as one of the agents in the story) > unlinked sentences |
| Vollm et al. | 2006 | fMRI | Explicit | Non-verbal | visual | high | 13 | ToM > physical causality |
| Walter et al. | 2004 | fMRI | Explicit | Non-verbal | visual | high | 13 | communicative intention > physical causality |
| | | | | | | | | private intention (one agent) > physical causality |
| | | | | | | | | private intention (two agents) > physical causality |
| Walter et al. | 2004 | fMRI | Explicit | Non-verbal | visual | high | 12 | private intention > physical causality |
| | | | | | | | | prospective intention > physical causality |
| | | | | | | | | communicative intention > physical causality |
| Walter et al. | 2009 | fMRI | Explicit | Non-verbal | visual | high | 12 | private intention > physical causality |
| | | | | | | | | prospective social intention > physical causality |
| | | | | | | | | communicative intention > physical causality |
| Wang et al. | 2015 | fMRI | Explicit | Non-verbal | visual | high | 56 | ToM > physical causality |
| Willert et al. | 2015 | fMRI | Explicit | Non-verbal | visual | high | 81 | ToM > physical judgements |

| | | | | | | | | |
|---|---|---|---|---|---|---|---|---|
| Young et al. | 2011 | fMRI | Explicit | Verbal | visual | high | 17 | false beliefs > false photographs |
| Zaitchik et al. | 2010 | fMRI | Explicit | Verbal | visual | high | 15 | stories about beliefs > non-ToM stories |
| | | | | | | | | stories about emotions > non-ToM stories |
| EXCLUDED CONTRASTS CONTAINING BOTH VISUAL AND AUDITORY STIMULI AND ALSO AUDITORY ONLY | | | | | | | | |
| Briend et al. | 2019 | fMRI | Explicit | Verbal | auditory | low | 28 | known (ToM) > unknown (non-ToM) language |
| Ferstl & von Cramon | 2002 | fMRI | Explicit | Verbal | auditory | high | 9 | ToM stories > pseudo-sentences |
| Gweon et al. | 2012 | fMRI | Explicit | Verbal | auditory | high | 8 | ToM > physical judgements |
| Kandylaki et al. | 2015 | fMRI | Explicit | Verbal | auditory | high | 20 | false belief > physical causality stories |
| | | | | | | | | ToM > physical judgements |
| Nieminen-von Wendt et al. | 2003 | PET | Explicit | Verbal | auditory | high | 8 | ToM > physical stories |
| Thye et al. | 2018 | fMRI | Explicit | Non-verbal | auditory | high | 18 | mental state > gender judgement |
| Dufour et al. | 2013 | fMRI | Explicit | Verbal | BOTH | high | 27 | false beliefs > false photographs |
| Wolf et al. | 2010 | fMRI | Explicit | BOTH | BOTH | high | 18 | ToM > non-ToM scenes (movie watching phase) |
| | | | Explicit | BOTH | BOTH | high | | ToM > physical inferences (response phase 1) |
| | | | Explicit | BOTH | BOTH | high | | ToM > physical inferences (response phase 2) |

**The references to access the listed studies are listed at the end of the document*

##### *Semantics References*

AbdulSabur NY, Xu Y, Liu S, Chow HM, Baxter M, Carson J, Braun AR. 2014. Neural correlates and network connectivity underlying narrative production and comprehension: A combined fMRI and PET study. Cortex. 57:107–127.

Abraham A, Pieritz K, Thybusch K, Rutter B, Kröger S, Schweckendiek J, Stark R, Windmann S, Hermann C. 2012. Creativity and the brain: Uncovering the neural signature of conceptual expansion. Neuropsychologia. 50:1906–1917.

Adank P, Davis MH, Hagoort P. 2012. Neural dissociation in processing noise and accent in spoken language comprehension. Neuropsychologia. 50:77–84.

Alain C, He Y, Grady C. 2008. The Contribution of the Inferior Parietal Lobe to Auditory Spatial Working Memory. J Cogn Neurosci. 20:285–295.

Assadollahi R, Meinzer M, Flaisch T, Obleser J, Rockstroh B. 2009. The representation of the verb's argument structure as disclosed by fMRI. BMC Neurosci. 10:3.

Axmacher N, Bialleck KA, Weber B, Helmstaedter C, Elger CE, Fell J. 2009. Working memory representation in atypical language dominance. Hum Brain Mapp. 30:2032–2043.

Bagga D, Singh N, Modi S, Kumar P, Bhattacharya D, Garg M, Khushu S. 2013. Assessment of lexical semantic judgment abilities in alcohol-dependent subjects: An fMRI study. J Biosci. 38:905–915.

Barrós-Loscertales A, González J, Pulvermüller F, Ventura-Campos N, Bustamante JC, Costumero V, Parcet MA, Ávila C. 2012. Reading Salt Activates Gustatory Brain Regions: fMRI Evidence for Semantic Grounding in a Novel Sensory Modality. Cereb Cortex. 22:2554–2563.

Baumgaertner A, Weiller C, Büchel C. 2002. Event-Related fMRI Reveals Cortical Sites Involved in Contextual Sentence Integration. Neuroimage. 16:736–745.

Baumgaertner A, Buccino G, Lange R, McNamara A, Binkofski F. 2007. Polymodal conceptual processing of human biological actions in the left inferior frontal lobe. Eur J Neurosci. 25:881–889.

Bautista A, Wilson SM. 2016. Neural responses to grammatically and lexically degraded speech. Lang Cogn Neurosci. 31:567–574.

Bick A, Goelman G, Frost R. 2008. Neural Correlates of Morphological Processes in Hebrew. J Cogn Neurosci. 20:406–420.

Binder JR, Frost JA, Hammeke TA, Bellgowan PSF, Rao SM, Cox RW. 1999. Conceptual Processing during the Conscious Resting State: A Functional MRI Study. J Cogn Neurosci. 11:80–93.

Binder JR, McKiernan KA, Parsons ME, Westbury CF, Possing ET, Kaufman JN, Buchanan L. 2003. Neural Correlates of Lexical Access during Visual Word Recognition. J Cogn Neurosci. 15:372–393.

Birn RM, Kenworthy L, Case L, Caravella R, Jones TB, Bandettini PA, Martin A. 2010. Neural systems supporting lexical search guided by letter and semantic category cues: A self-paced overt response fMRI study of verbal fluency. Neuroimage. 49:1099–1107.

Bonhage CE, Fiebach CJ, Bahlmann J, Mueller JL. 2014. Brain Signature of Working Memory for Sentence Structure: Enriched Encoding and Facilitated Maintenance. J Cogn Neurosci. 26:1654–1671.

Bonhage CE, Mueller JL, Friederici AD, Fiebach CJ. 2015. Combined eye tracking and fMRI reveals neural basis of linguistic predictions during sentence comprehension. Cortex. 68:33–47.

Booth JR, Lu D, Burman DD, Chou T-L, Jin Z, Peng D-L, Zhang L, Ding G-S, Deng Y, Liu L. 2006. Specialization of phonological and semantic processing in Chinese word reading. Brain Res. 1071:197–207.

Boulenger V, Hauk O, Pulvermuller F. 2009. Grasping Ideas with the Motor System: Semantic Somatotopy in Idiom Comprehension. Cereb Cortex. 19:1905–1914.

Bozic M, Marslen-Wilson W. 2013. Neurocognitive mechanisms for processing inflectional and derivational complexity in English. Psihologija. 46:439–454.

Brambati SM, Benoit S, Monetta L, Belleville S, Joubert S. 2010. The role of the left anterior temporal lobe in the semantic processing of famous faces. Neuroimage. 53:674–681.

Bruffaerts R, Dupont P, Peeters R, De Deyne S, Storms G, Vandenberghe R. 2013. Similarity of fMRI Activity Patterns in Left Perirhinal Cortex Reflects Semantic Similarity between Words. J Neurosci. 33:18597–18607.

Bulut T, Hung Y-H, Tzeng O, Wu DH. 2017. Neural correlates of processing sentences and compound words in Chinese. PLoS One. 12:e0188526.
Cai C, Kochiyama T, Osaka K, Wu J. 2007. Lexical/semantic processing in dorsal left inferior frontal gyrus. Neuroreport. 18:1147–1151.
Cao F, Peng D, Liu L, Jin Z, Fan N, Deng Y, Booth JR. 2009. Developmental differences of neurocognitive networks for phonological and semantic processing in Chinese word reading. Hum Brain Mapp. 30:797–809.
Cappa SF, Perani D, Schnur T, Tettamanti M, Fazio F. 1998. The Effects of Semantic Category and Knowledge Type on Lexical-Semantic Access: A PET Study. Neuroimage. 8:350–359.
Carota F, Kriegeskorte N, Nili H, Pulvermüller F. 2017. Representational Similarity Mapping of Distributional Semantics in Left Inferior Frontal, Middle Temporal, and Motor Cortex. Cereb Cortex. 27:294–309.
Carota F, Moseley R, Pulvermüller F. 2012. Body-part-specific Representations of Semantic Noun Categories. J Cogn Neurosci. 24:1492–1509.
Chan S, Tang S, Tang K, Lee W, Lo S, Kwong KK. 2009. Hierarchical coding of characters in the ventral and dorsal visual streams of Chinese language processing. Neuroimage. 48:423–435.
Chiao JY, Harada T, Oby ER, Li Z, Parrish T, Bridge DJ. 2009. Neural representations of social status hierarchy in human inferior parietal cortex. Neuropsychologia. 47:354–363.
Chou T-L, Chen C-W, Wu M-Y, Booth JR. 2009. The role of inferior frontal gyrus and inferior parietal lobule in semantic processing of Chinese characters. Exp Brain Res. 198:465–475.
Chouinard PA, Morrissey BF, Köhler S, Goodale MA. 2008. Repetition suppression in occipital–temporal visual areas is modulated by physical rather than semantic features of objects. Neuroimage. 41:130–144.
Chow HM, Kaup B, Raabe M, Greenlee MW. 2008. Evidence of fronto-temporal interactions for strategic inference processes during language comprehension. Neuroimage. 40:940–954.
Christensen TA, Antonucci SM, Lockwood JL, Kittleson M, Plante E. 2008. Cortical and subcortical contributions to the attentive processing of speech. Neuroreport. 19:1101–1105.
Clos M, Langner R, Meyer M, Oechslin MS, Zilles K, Eickhoff SB. 2014. Effects of prior information on decoding degraded speech: An fMRI study. Hum Brain Mapp. 35:61–74.
D'Arcy RCN, Bolster RB, Ryner L, Mazerolle EL, Grant J, Song X. 2007. A site directed fMRI approach for evaluating functional status in the anterolateral temporal lobes. Neurosci Res. 57:120–128.
Damasio H, Tranel D, Grabowski T, Adolphs R, Damasio A. 2004. Neural systems behind word and concept retrieval. Cognition. 92:179–229.
Damasio H, Grabowski TJ, Tranel D, Ponto LLB, Hichwa RD, Damasio AR. 2001. Neural Correlates of Naming Actions and of Naming Spatial Relations. Neuroimage. 13:1053–1064.
Dapretto M, Bookheimer SY. 1999. Form and Content. Neuron. 24:427–432.
Davis MH, Ford MA, Kherif F, Johnsrude IS. 2011. Does Semantic Context Benefit Speech Understanding through "Top–Down" Processes? Evidence from Time-resolved Sparse fMRI. J Cogn Neurosci. 23:3914–3932.
Davis MH, Meunier F, Marslen-Wilson WD. 2004. Neural responses to morphological, syntactic, and semantic properties of single words: An fMRI study☆. Brain Lang. 89:439–449.
Demonet, J. F., Choullet. F., Ramsay S., Cardebat, D., Nespoulous, J.L., Wise, R., Rascol, A., Frackowiak, L. 1992. The Anatomy of Phonological and Semantic Processing in Normal Subjects. Brain. 115:1753–1768.
Devlin J. 2002. Is there an anatomical basis for category-specificity? Semantic memory studies in PET and fMRI. Neuropsychologia. 40:54–75.
Devlin JT, Matthews PM, Rushworth MFS. 2003. Semantic Processing in the Left Inferior Prefrontal Cortex: A Combined Functional Magnetic Resonance Imaging and Transcranial Magnetic Stimulation Study. J Cogn Neurosci. 15:71–84.

Devlin JT, Russell RP, Davis MH, Price CJ, Wilson J, Moss HE, Matthews PM, Tyler LK. 2000. Susceptibility-Induced Loss of Signal: Comparing PET and fMRI on a Semantic Task. Neuroimage. 11:589–600.

Diaz MT, McCarthy G. 2009. A comparison of brain activity evoked by single content and function words: An fMRI investigation of implicit word processing. Brain Res. 1282:38–49.

Dreyer FR, Pulvermüller F. 2018. Abstract semantics in the motor system? – An event-related fMRI study on passive reading of semantic word categories carrying abstract emotional and mental meaning. Cortex. 100:52–70.

Ebisch SJH, Babiloni C, Del Gratta C, Ferretti A, Perrucci MG, Caulo M, Sitskoorn MM, Romani GL. 2007. Human Neural Systems for Conceptual Knowledge of Proper Object Use: A Functional Magnetic Resonance Imaging Study. Cereb Cortex. 17:2744–2751.

Elfgren C, van Westen D, Passant U, Larsson E-M, Mannfolk P, Fransson P. 2006. fMRI activity in the medial temporal lobe during famous face processing. Neuroimage. 30:609–616.

Emmorey K, Weisberg J, McCullough S, Petrich JAF. 2013. Mapping the reading circuitry for skilled deaf readers: An fMRI study of semantic and phonological processing. Brain Lang. 126:169–180.

Emmorey K, Xu J, Gannon P, Goldin-Meadow S, Braun A. 2010. CNS activation and regional connectivity during pantomime observation: No engagement of the mirror neuron system for deaf signers. Neuroimage. 49:994–1005.

Engelien A, Tüscher O, Hermans W, Isenberg N, Eidelberg D, Frith C, Stern E, Silbersweig D. 2006. Functional neuroanatomy of non-verbal semantic sound processing in humans. J Neural Transm. 113:599–608.

Erb J, Henry MJ, Eisner F, Obleser J. 2013. The Brain Dynamics of Rapid Perceptual Adaptation to Adverse Listening Conditions. J Neurosci. 33:10688–10697.

Europa E, Gitelman DR, Kiran S, Thompson CK. 2019. Neural Connectivity in Syntactic Movement Processing. Front Hum Neurosci. 13:1–15.

Foki T, Gartus A, Geissler A, Beisteiner R. 2008. Probing overtly spoken language at sentential level—A comprehensive high-field BOLD–fMRI protocol reflecting everyday language demands. Neuroimage. 39:1613–1624.

Friederici AD, Kotz SA, Scott SK, Obleser J. 2009. Disentangling syntax and intelligibility in auditory language comprehension. Hum Brain Mapp. 31:NA-NA.

Friese U, Rutschmann R, Raabe M, Schmalhofer F. 2008. Neural Indicators of Inference Processes in Text Comprehension: An Event-related Functional Magnetic Resonance Imaging Study. J Cogn Neurosci. 20:2110–2124.

Garbin G, Collina S, Tabossi P. 2012. Argument Structure and Morphological Factors in Noun and Verb Processing: An fMRI Study. PLoS One. 7:e45091.

Garn CL, Allen MD, Larsen JD. 2009. An fMRI study of sex differences in brain activation during object naming. Cortex. 45:610–618.

Geranmayeh F, Brownsett SLE, Leech R, Beckmann CF, Woodhead Z, Wise RJS. 2012. The contribution of the inferior parietal cortex to spoken language production. Brain Lang. 121:47–57.

Gerlach C, Law I, Gade A, Paulson OB. 1999. Perceptual differentiation and category effects in normal object recognition. Brain. 122:2159–2170.

Gesierich B, Jovicich J, Riello M, Adriani M, Monti A, Brentari V, Robinson SD, Wilson SM, Fairhall SL, Gorno-Tempini ML. 2012. Distinct neural substrates for semantic knowledge and naming in the temporoparietal network. Cereb Cortex. 22:2217–2226.

Giraud AL, Price CJ. 2001. The Constraints Functional Neuroimaging Places on Classical Models of Auditory Word Processing. J Cogn Neurosci. 13:754–765.

Giraud AL. 2004. Contributions of Sensory Input, Auditory Search and Verbal Comprehension to Cortical Activity during Speech Processing. Cereb Cortex. 14:247–255.

Gitelman DR, Nobre AC, Sonty S, Parrish TB, Mesulam M-M. 2005. Language network specializations: An analysis with parallel task designs and functional magnetic resonance imaging. Neuroimage. 26:975–985.

Gorno-Tempini, M. L., Price, C. J., Josephs, O., Vandenberghe, R., Cappa, S. F., Kapur, N., ... & Tempini, M. L. (1998). The neural systems sustaining face and proper-name processing. Brain: a journal of neurology, 121(11), 2103-2118.

Grabowski TJ, Damasio H, Tranel D, Ponto LLB, Hichwa RD, Damasio AR. 2001. A role for left temporal pole in the retrieval of words for unique entities. Hum Brain Mapp. 13:199–212.

Graves WW, Binder JR, Desai RH, Conant LL, Seidenberg MS. 2010. Neural correlates of implicit and explicit combinatorial semantic processing. Neuroimage. 53:638–646.

Grindrod CM, Garnett EO, Malyutina S, den Ouden DB. 2014. Effects of representational distance between meanings on the neural correlates of semantic ambiguity. Brain Lang. 139:23–35.

Grossman M, Koenig P, DeVita C, Glosser G, Alsop D, Detre J, Gee J. 2002. The Neural Basis for Category-Specific Knowledge: An fMRI Study. Neuroimage. 15:936–948.

Grossman M, Koenig P, DeVita C, Glosser G, Alsop D, Detre J, Gee J. 2002. Neural representation of verb meaning: An fMRI study. Hum Brain Mapp. 15:124–134.

Groussard M, Viader F, Hubert V, Landeau B, Abbas A, Desgranges B, Eustache F, Platel H. 2010. Musical and verbal semantic memory: Two distinct neural networks? Neuroimage. 49:2764–2773.

Guediche S, Reilly M, Santiago C, Laurent P, Blumstein SE. 2016. An fMRI study investigating effects of conceptually related sentences on the perception of degraded speech. Cortex. 79:57–74.

Gurd JM, Amunts K, Weiss PH, Zafiris O, Zilles K, Marshall JC, Fink GR. 2002. Posterior parietal cortex is implicated in continuous switching between verbal fluency tasks: an fMRI study with clinical implications. Brain. 125:1024–1038.

Häberling IS, Corballis PM, Corballis MC. 2016. Language, gesture, and handedness: Evidence for independent lateralized networks. Cortex. 82:72–85.

Hagoort P, Indefrey P, Brown C, Herzog H, Steinmetz H, Seitz RJ. 1999. The Neural Circuitry Involved in the Reading of German Words and Pseudowords: A PET Study. J Cogn Neurosci. 11:383–398.

Harrington GS, Farias D, Davis CH. 2009. The neural basis for simulated drawing and the semantic implications. Cortex. 45:386–393.

Hartung F, Hagoort P, Willems RM. 2017. Readers select a comprehension mode independent of pronoun: Evidence from fMRI during narrative comprehension. Brain Lang. 170:29–38.

Hauk O, Pulvermuller F. 2011. The lateralization of motor cortex activation to action-words. Front Hum Neurosci. 5:1–10.

Hayashi A, Okamoto Y, Yoshimura S, Yoshino A, Toki S, Yamashita H, Matsuda F, Yamawaki S. 2014. Visual imagery while reading concrete and abstract Japanese kanji words: An fMRI study. Neurosci Res. 79:61–66.

Heim S, Eickhoff SB, Amunts K. 2008. Specialisation in Broca's region for semantic, phonological, and syntactic fluency? Neuroimage. 40:1362–1368.

Henke K, Weber B, Kneifel S, Wieser HG, Buck A. 1999. Human hippocampus associates information in memory. Proc Natl Acad Sci. 96:5884–5889.

Herbster AN, Mintun MA, Nebes RD, Becker JT. 1997. Regional cerebral blood flow during word and nonword reading. Hum Brain Mapp. 5:84–92.

Hervais-Adelman AG, Carlyon RP, Johnsrude IS, Davis MH. 2012. Brain regions recruited for the effortful comprehension of noise-vocoded words. Lang Cogn Process. 27:1145–1166.

Higuchi H, Moriguchi Y, Murakami H, Katsunuma R, Mishima K, Uno A. 2015. Neural basis of hierarchical visual form processing of Japanese Kanji characters. Brain Behav. 5:n/a-n/a.

Hocking J, McMahon KL, de Zubicaray GI. 2011. Cortical organization of environmental sounds by attribute. Hum Brain Mapp. 32:688–698.

Holle H, Gunter TC, Rüschemeyer S-A, Hennenlotter A, Iacoboni M. 2008. Neural correlates of the processing of co-speech gestures. Neuroimage. 39:2010–2024.

Homae F, Yahata N, Sakai KL. 2003. Selective enhancement of functional connectivity in the left prefrontal cortex during sentence processing. Neuroimage. 20:578–586.

Husain FT, Patkin DJ, Kim J, Braun AR, Horwitz B. 2012. Dissociating neural correlates of meaningful emblems from meaningless gestures in deaf signers and hearing non-signers. Brain Res. 1478:24–35.

Hwang K, Palmer ED, Basho S, Zadra JR, Müller R-A. 2009. Category-specific activations during word generation reflect experiential sensorimotor modalities. Neuroimage. 48:717–725.

Ikuta S., Sugiura M., Sassa Y., Watanebe J., Akitsuki Y., Iwata K., Miura N., Okamotho H., Watanabe Y., Sato S. 2006. Brain activation during the course of sentence comprehension. Brain Lang. 97:154–161.

Jackson RL, Hoffman P, Pobric G, Lambon Ralph MA. 2015. The Nature and Neural Correlates of Semantic Association versus Conceptual Similarity. Cereb Cortex. 25:4319–4333.

Jensen EJ, Hargreaves I, Bass A, Pexman P, Goodyear BG, Federico P. 2011. Cortical reorganization and reduced efficiency of visual word recognition in right temporal lobe epilepsy: A functional MRI study. Epilepsy Res. 93:155–163.
Jeon H-A, Lee K-M, Kim Y-B, Cho Z-H. 2009. Neural substrates of semantic relationships: Common and distinct left-frontal activities for generation of synonyms vs. antonyms. Neuroimage. 48:449–457.
Joubert S, Beauregard M, Walter N, Bourgouin P, Beaudoin G, Leroux J-M, Karama S, Lecours AR. 2004. Neural correlates of lexical and sublexical processes in reading. Brain Lang. 89:9–20.
Kang E, Lee DS, Kang H, Hwang CH, Oh S-H, Kim C-S, Chung J-K, Lee MC. 2006. The neural correlates of cross-modal interaction in speech perception during a semantic decision task on sentences: A PET study. Neuroimage. 32:423–431.
Khader PH, Jost K, Mertens M, Bien S, Rösler F. 2010. Neural correlates of generating visual nouns and motor verbs in a minimal phrase context. Brain Res. 1318:122–132.
Kim J, Koizumi M, Ikuta N, Fukumitsu Y, Kimura N, Iwata K, Watanabe J, Yokoyama S, Sato S, Horie K, Kawashima R. 2009. Scrambling effects on the processing of Japanese sentences: An fMRI study. J Neurolinguistics. 22:151–166.
Kinno R, Kawamura M, Shioda S, Sakai KL. 2008. Neural correlates of noncanonical syntactic processing revealed by a picture-sentence matching task. Hum Brain Mapp. 29:1015–1027.
Kotz S. 2002. Modulation of the Lexical–Semantic Network by Auditory Semantic Priming: An Event-Related Functional MRI Study. Neuroimage. 17:1761–1772.
Kuchinke L, Jacobs AM, Grubich C, Võ MLH, Conrad M, Herrmann M. 2005. Incidental effects of emotional valence in single word processing: An fMRI study. Neuroimage. 28:1022–1032.
Kumar U. 2016. Neural dichotomy of word concreteness: a view from functional neuroimaging. Cogn Process. 17:39–48.
Kuperberg GR, McGuire PK, Bullmore ET, Brammer MJ, Rabe-Hesketh S, Wright IC, Lythgoe DJ, Williams SCR, David AS. 2000. Common and Distinct Neural Substrates for Pragmatic, Semantic, and Syntactic Processing of Spoken Sentences: An fMRI Study. J Cogn Neurosci. 12:321–341.
Kyong JS, Scott SK, Rosen S, Howe TB, Agnew ZK, McGettigan C. 2014. Exploring the Roles of Spectral Detail and Intonation Contour in Speech Intelligibility: An fMRI Study. J Cogn Neurosci. 26:1748–1763.
Leff AP, Schofield TM, Stephan KE, Crinion JT, Friston KJ, Price CJ. 2008. The Cortical Dynamics of Intelligible Speech. J Neurosci. 28:13209–13215.
Leung AWS, Alain C. 2011. Working memory load modulates the auditory "What" and "Where" neural networks. Neuroimage. 55:1260–1269.
Leveroni CL, Seidenberg M, Mayer AR, Mead LA, Binder JR, Rao SM. 2000. Neural Systems Underlying the Recognition of Familiar and Newly Learned Faces. J Neurosci. 20:878–886.
Lin N, Wang X, Zhao Y, Liu Y, Li X, Bi Y. 2015. Premotor Cortex Activation Elicited during Word Comprehension Relies on Access of Specific Action Concepts. J Cogn Neurosci. 27:2051–2062.
Liu L, Deng X, Peng D, Cao F, Ding G, Jin Z, Zeng Y, Li K, Zhu L, Fan N, Deng Y, Bolger DJ, Booth JR. 2009. Modality- and Task-specific Brain Regions Involved in Chinese Lexical Processing. J Cogn Neurosci. 21:1473–1487.
Liuzzi AG, Bruffaerts R, Peeters R, Adamczuk K, Keuleers E, De Deyne S, Storms G, Dupont P, Vandenberghe R. 2017. Cross-modal representation of spoken and written word meaning in left pars triangularis. Neuroimage. 150:292–307.
Ludersdorfer P, Schurz M, Richlan F, Kronbichler M, Wimmer H. 2013. Opposite effects of visual and auditory word-likeness on activity in the visual word form area. Front Hum Neurosci. 7:1–10.
Ludersdorfer P, Wimmer H, Richlan F, Schurz M, Hutzler F, Kronbichler M. 2016. Left ventral occipitotemporal activation during orthographic and semantic processing of auditory words. Neuroimage. 124:834–842.
Malins JG, Gumkowski N, Buis B, Molfese P, Rueckl JG, Frost SJ, Pugh KR, Morris R, Mencl WE. 2016. Dough, tough, cough, rough: A "fast" fMRI localizer of component processes in reading. Neuropsychologia. 91:394–406.
Marques JF, Canessa N, Cappa S. 2009. Neural differences in the processing of true and false sentences: Insights into the nature of "truth" in language comprehension. Cortex. 45:759–768.

Marques JF, Canessa N, Siri S, Catricalà E, Cappa S. 2008. Conceptual knowledge in the brain: fMRI evidence for a featural organization. Brain Res. 1194:90–99.
Mashal N, Vishne T, Laor N, Titone D. 2013. Enhanced left frontal involvement during novel metaphor comprehension in schizophrenia: Evidence from functional neuroimaging. Brain Lang. 124:66–74.
Matchin W, Hammerly C, Lau E. 2017. The role of the IFG and pSTS in syntactic prediction: Evidence from a parametric study of hierarchical structure in fMRI. Cortex. 88:106–123.
Matchin W, Liao C-H, Gaston P, Lau E. 2019. Same words, different structures: An fMRI investigation of argument relations and the angular gyrus. Neuropsychologia. 125:116–128.
Mellem MS, Jasmin KM, Peng C, Martin A. 2016. Sentence processing in anterior superior temporal cortex shows a social-emotional bias. Neuropsychologia. 89:217–224.
Menz MM, Blangero A, Kunze D, Binkofski F. 2010. Got it! Understanding the concept of a tool. Neuroimage. 51:1438–1444.
Metz-Lutz. 2010. What physiological changes and cerebral traces tell us about adhesion to fiction during theater-watching? Front Hum Neurosci. 4:1–10.
Meyer M, Alter K, Friederici AD, Lohmann G, von Cramon DY. 2002. FMRI reveals brain regions mediating slow prosodic modulations in spoken sentences. Hum Brain Mapp. 17:73–88.
Moberget T, Gullesen EH, Andersson S, Ivry RB, Endestad T. 2014. Generalized role for the cerebellum in encoding internal models: Evidence from semantic processing. J Neurosci. 34:2871–2878.
Moseley R, Carota F, Hauk O, Mohr B, Pulvermüller F. 2012. A Role for the Motor System in Binding Abstract Emotional Meaning. Cereb Cortex. 22:1634–1647.
Mummery CJ, Patterson K, Hodges JR, Price CJ. 1998. Functional Neuroanatomy of the Semantic System: Divisible by What? J Cogn Neurosci. 10:766–777.
Nakamura K. 2000. Functional delineation of the human occipito-temporal areas related to face and scene processing: A PET study. Brain. 123:1903–1912.
Nakamura K, Kawashima R, Sugiura M, Kato T, Nakamura A, Hatano K, Nagumo S, Kubota K, Fukuda H, Ito K, Kojima S. 2001. Neural substrates for recognition of familiar voices: a PET study. Neuropsychologia. 39:1047–1054.
Nichelli P, Grafman J, Pietrini P, Clark K, Lee KY, Miletich R. 1995. Where the brain appreciates the moral of a story. Neuroreport. 6:2309–2313.
Nielson KA, Seidenberg M, Woodard JL, Durgerian S, Zhang Q, Gross WL, Gander A, Guidotti LM, Antuono P, Rao SM. 2010. Common neural systems associated with the recognition of famous faces and names: An event-related fMRI study. Brain Cogn. 72:491–498.
Noppeney U, Price CJ. 2003. Functional imaging of the semantic system: Retrieval of sensory-experienced and verbally learned knowledge. Brain Lang. 84:120–133.
Orfanidou E, Marslen-Wilson WD, Davis MH. 2006. Neural response suppression predicts repetition priming of spoken words and pseudowords. J Cogn Neurosci. 18:1237–1252.
Pallier C, Devauchelle A-D, Dehaene S. 2011. Cortical representation of the constituent structure of sentences. Proc Natl Acad Sci. 108:2522–2527.
Peelle JE, Eason RJ, Schmitter S, Schwarzbauer C, Davis MH. 2010. Evaluating an acoustically quiet EPI sequence for use in fMRI studies of speech and auditory processing. Neuroimage. 52:1410–1419.
Perani D, Schnur T, Tettamanti M, Italy, Cappa SF, Fazio F. 1999. Word and picture matching: a PET study of semantic category effects. Neuropsychologia. 37:293–306.
Perrone-Bertolotti M, Kauffmann L, Pichat C, Vidal JR, Baciu M. 2017. Effective Connectivity between Ventral Occipito-Temporal and Ventral Inferior Frontal Cortex during Lexico-Semantic Processing. A Dynamic Causal Modeling Study. Front Hum Neurosci. 11:1–13.
Pilgrim LK, Fadili J, Fletcher P, Tyler LK. 2002. Overcoming Confounds of Stimulus Blocking: An Event-Related fMRI Design of Semantic Processing. Neuroimage. 16:713–723.
Price CJ, Moore CJ, Humphreys GW, Wise RJS. 1997. Segregating Semantic from Phonological Processes during Reading. J Cogn Neurosci. 9:727–733.
Pulvermüller F, Cook C, Hauk O. 2012. Inflection in action: Semantic motor system activation to noun- and verb-containing phrases is modulated by the presence of overt grammatical markers. Neuroimage. 60:1367–1379.
Raettig T, Kotz SA. 2008. Auditory processing of different types of pseudo-words: An event-related fMRI study. Neuroimage. 39:1420–1428.
Raposo A, Frade S, Alves M. 2016. Framing memories: How the retrieval query format shapes the neural bases of remembering. Neuropsychologia. 89:309–319.

Raposo A, Moss HE, Stamatakis EA, Tyler LK. 2009. Modulation of motor and premotor cortices by actions, action words and action sentences. Neuropsychologia. 47:388–396.

Rapp B, Lipka K. 2011. The Literate Brain: The Relationship between Spelling and Reading. J Cogn Neurosci. 23:1180–1197.

Redcay E, Velnoskey KR, Rowe ML. 2016. Perceived communicative intent in gesture and language modulates the superior temporal sulcus. Hum Brain Mapp. 37:3444–3461.

Rissman J, Eliassen JC, Blumstein SE. 2003. An Event-Related fMRI Investigation of Implicit Semantic Priming. J Cogn Neurosci. 15:1160–1175.

Robertson DA, Gernsbacher MA, Guidotti SJ, Robertson RRW, Irwin W, Mock BJ, Campana ME. 2000. Functional neuroanatomy of the cognitive process of mapping during discourse comprehension. Psychol Sci. 11:255–260.

Rodd JM, Johnsrude IS, Davis MH. 2012. Dissociating Frontotemporal Contributions to Semantic Ambiguity Resolution in Spoken Sentences. Cereb Cortex. 22:1761–1773.

Rodd JM, Longe OA, Randall B, Tyler LK. 2010. The functional organisation of the fronto-temporal language system: Evidence from syntactic and semantic ambiguity. Neuropsychologia. 48:1324–1335.

Rogalsky C, Almeida D, Sprouse J, Hickok G. 2015. Sentence processing selectivity in Broca’s area: evident for structure but not syntactic movement. Lang Cogn Neurosci. 30:1326–1338.

Rogalsky C, Hickok G. 2009. Selective Attention to Semantic and Syntactic Features Modulates Sentence Processing Networks in Anterior Temporal Cortex. Cereb Cortex. 19:786–796.

Rogers T.T, Hocking J., Noppeney U., Mechelli A., Gorno-Tempini M., Patterson K., Price C.J. 2006. Anterior temporal cortex and semantic memory: Reconciling findings from neuropsychology and functional imaging. Cogn Affect Behav Neurosci. 6:201–213.

Roskies AL, Fiez JA, Balota DA, Raichle ME, Petersen SE. 2001. Task-dependent modulation of regions in the left inferior frontal cortex during semantic processing. J Cogn Neurosci. 13:829–843.

Ross LA, Olson IR. 2012. What’s unique about unique entities? An fMRI investigation of the semantics of famous faces and landmarks. Cereb Cortex. 22:2005–2015.

Roxbury T, McMahon K, Coulthard A, Copland DA. 2016. An fMRI Study of Concreteness Effects during Spoken Word Recognition in Aging. Preservation or Attenuation? Front Aging Neurosci. 7:5–7.

Ryan L, Cox C, Hayes SM, Nadel L. 2008. Hippocampal activation during episodic and semantic memory retrieval: Comparing category production and category cued recall. Neuropsychologia. 46:2109–2121.

Ryan L, Lin CY, Ketcham K, Nadel L. 2010. The role of medial temporal lobe in retrieving spatial and nonspatial relations from episodic and semantic memory. Hippocampus. 20:11–18.

Sabri M, Binder JR, Desai R, Medler DA, Leitl MD, Liebenthal E. 2008. Attentional and linguistic interactions in speech perception. Neuroimage. 39:1444–1456.

Sachs O, Weis S, Krings T, Huber W, Kircher T. 2008. Categorical and thematic knowledge representation in the brain: Neural correlates of taxonomic and thematic conceptual relations. Neuropsychologia. 46:409–418.

Saur D, Kreher BW, Schnell S, Kummerer D, Kellmeyer P, Vry M-S, Umarova R, Musso M, Glauche V, Abel S, Huber W, Rijntjes M, Hennig J, Weiller C. 2008. Ventral and dorsal pathways for language. Proc Natl Acad Sci. 105:18035–18040.

Schell M, Zaccarella E, Friederici AD. 2017. Differential cortical contribution of syntax and semantics: An fMRI study on two-word phrasal processing. Cortex. 96:105–120.

Schmitt JM, Auer P, Ferstl EC. 2019. Understanding fairy tales spoken in dialect: an fMRI study. Lang Cogn Neurosci. 34:440–456.

Schuil KDI, Smits M, Zwaan RA. 2013. Sentential Context Modulates the Involvement of the Motor Cortex in Action Language Processing: An fMRI Study. Front Hum Neurosci. 7:1–13.

Scott SK. 2000. Identification of a pathway for intelligible speech in the left temporal lobe. Brain. 123:2400–2406.

Segal E, Petrides M. 2012. The anterior superior parietal lobule and its interactions with language and motor areas during writing. Eur J Neurosci. 35:309–322.

Seghier ML, Josse G, Leff AP, Price CJ. 2011. Lateralization is Predicted by Reduced Coupling from the Left to Right Prefrontal Cortex during Semantic Decisions on Written Words. Cereb Cortex. 21:1519–1531.

Sergent J., Ohta S., Macdonald J. 1992. Functional Neuroanatomy of Face and Object Processing. Brain. 115:15–36.
Sheldon S, McAndrews MP, Pruessner J, Moscovitch M. 2016. Dissociating patterns of anterior and posterior hippocampal activity and connectivity during distinct forms of category fluency. Neuropsychologia. 90:148–158.
Simard F, Monetta L, Nagano-Saito A, Monchi O. 2013. A new lexical card-sorting task for studying fronto-striatal contribution to processing language rules. Brain Lang. 125:295–306.
Slioussar N, Kireev M V., Chernigovskaya T V., Kataeva G V., Korotkov AD, Medvedev S V. 2014. An ER-fMRI study of Russian inflectional morphology. Brain Lang. 130:33–41.
Smith EE, Myers N, Sethi U, Pantazatos S, Yanagihara T, Hirsch J. 2012. Conceptual representations of perceptual knowledge. Cogn Neuropsychol. 29:237–248.
Snijders TM, Vosse T, Kempen G, Van Berkum JJA, Petersson KM, Hagoort P. 2009. Retrieval and Unification of Syntactic Structure in Sentence Comprehension: an fMRI Study Using Word-Category Ambiguity. Cereb Cortex. 19:1493–1503.
Stove L.A., Paans A.M.J., Wijers A.A, Zwarts F., Mulder F., Vaalburg D. 1999. Sentence comprehension and word repetition: A positron emission tomography investigation. Psychophysiology. 36:S0048577299980150.
Straube B, Green A, Weis S, Kircher T. 2012. A Supramodal Neural Network for Speech and Gesture Semantics: An fMRI Study. PLoS One. 7:e51207.
Stringaris AK, Medford NC, Giampietro V, Brammer MJ, David AS. 2007. Deriving meaning: Distinct neural mechanisms for metaphoric, literal, and non-meaningful sentences. Brain Lang. 100:150–162.
Sugiura M, Kawashima R, Nakamura K, Sato N, Nakamura A, Kato T, Hatano K, Schormann T, Zilles K, Sato K, Ito K, Fukuda H. 2001. Activation reduction in anterior temporal cortices during repeated recognition of faces of personal acquaintances. Neuroimage. 13:877–890.
Sugiura M, Sassa Y, Watanabe J, Akitsuki Y, Maeda Y, Matsue Y, Fukuda H, Kawashima R. 2006. Cortical mechanisms of person representation: Recognition of famous and personally familiar names. Neuroimage. 31:853–860.
Sugiura M, Sassa Y, Watanabe J, Akitsuki Y, Maeda Y, Matsue Y, Kawashima R. 2008. Anatomical Segregation of Representations of Personally Familiar and Famous People in the Temporal and Parietal Cortices. J Cogn Neurosci. 21:1855–1868.
Sun K, Xue R, Zhang P, Zuo Z, Chen Z, Wang B, Martin T, Wang Y, Chen L, He S, Wang DJJ. 2017. Integrated SSFP for functional brain mapping at 7 T with reduced susceptibility artifact. J Magn Reson. 276:22–30.
Szlachta Z, Bozic M, Jelowicka A, Marslen-Wilson WD. 2012. Neurocognitive dimensions of lexical complexity in Polish. Brain Lang. 121:219–225.
Takeichi H, Koyama S, Terao A, Takeuchi F, Toyosawa Y, Murohashi H. 2010. Comprehension of degraded speech sounds with m-sequence modulation: An fMRI study. Neuroimage. 49:2697–2706.
Taminato T, Miura N, Sugiura M, Kawashima R. 2014. Neuronal substrates characterizing two stages in visual object recognition. Neurosci Res. 89:61–68.
Taylor MJ, Arsalidou M, Bayless SJ, Morris D, Evans JW, Barbeau EJ. 2009. Neural correlates of personally familiar faces: Parents, partner and own faces. Hum Brain Mapp. 30:2008–2020.
Thierry G, Price CJ. 2006. Dissociating Verbal and Nonverbal Conceptual Processing in the Human Brain. J Cogn Neurosci. 18:1018–1028.
Tieleman A, Seurinck R, Deblaere K, Vandemaele P, Vingerhoets G, Achten E. 2005. Stimulus pacing affects the activation of the medial temporal lobe during a semantic classification task: An fMRI study. Neuroimage. 26:565–572.
Tyler L., Stamatakis E., Dick E, Bright P, Fletcher P, Moss H. 2003. Objects and their actions: evidence for a neurally distributed semantic system. Neuroimage. 18:542–557.
Vagharchakian L, Dehaene-Lambertz G, Pallier C, Dehaene S. 2012. A Temporal Bottleneck in the Language Comprehension Network. J Neurosci. 32:9089–9102.
Van Ettinger-Veenstra H, McAllister A, Lundberg P, Karlsson T, Engström M. 2016. Higher Language Ability is Related to Angular Gyrus Activation Increase During Semantic Processing, Independent of Sentence Incongruency. Front Hum Neurosci. 10:1–9.
van Leeuwen TM, Lamers MJA, Petersson KM, Gussenhoven C, Rietveld T, Poser B, Hagoort P. 2014. Phonological markers of information structure: An fMRI study. Neuropsychologia. 58:64–74.
Vignali L, Hawelka S, Hutzler F, Richlan F. 2019. Processing of parafoveally presented words. An fMRI study. Neuroimage. 184:1–9.

Vingerhoets G. 2008. Knowing about tools: Neural correlates of tool familiarity and experience. Neuroimage. 40:1380–1391.
Visser M, Jefferies E, Embleton K V., Lambon Ralph MA. 2012. Both the Middle Temporal Gyrus and the Ventral Anterior Temporal Area Are Crucial for Multimodal Semantic Processing: Distortion-corrected fMRI Evidence for a Double Gradient of Information Convergence in the Temporal Lobes. J Cogn Neurosci. 24:1766–1778.
Vitello S, Warren JE, Devlin JT, Rodd JM. 2014. Roles of frontal and temporal regions in reinterpreting semantically ambiguous sentences. Front Hum Neurosci. 8:1–14.
von Kriegstein K, Eger E, Kleinschmidt A, Giraud AL. 2003. Modulation of neural responses to speech by directing attention to voices or verbal content. Cogn Brain Res. 17:48–55.
Wang X, Zhao R, Zevin JD, Yang J. 2016. The Neural Correlates of the Interaction between Semantic and Phonological Processing for Chinese Character Reading. Front Psychol. 7:1–14.
Weiss Y, Katzir T, Bitan T. 2015. Many ways to read your vowels—Neural processing of diacritics and vowel letters in Hebrew. Neuroimage. 121:10–19.
Welcome SE, Joanisse MF. 2012. Individual differences in skilled adult readers reveal dissociable patterns of neural activity associated with component processes of reading. Brain Lang. 120:360–371.
Wende KC, Straube B, Stratmann M, Sommer J, Kircher T, Nagels A. 2012. Neural correlates of continuous causal word generation. Neuroimage. 62:1399–1407.
Wirth M, Jann K, Dierks T, Federspiel A, Wiest R, Horn H. 2011. Semantic memory involvement in the default mode network: A functional neuroimaging study using independent component analysis. Neuroimage. 54:3057–3066.
Wright ND, Mechelli A, Noppeney U, Veltman DJ, Rombouts SARB, Glensman J, Haynes J-D, Price CJ. 2008. Selective activation around the left occipito-temporal sulcus for words relative to pictures: Individual variability or false positives? Hum Brain Mapp. 29:986–1000.
Wright P, Randall B, Marslen-Wilson WD, Tyler LK. 2011. Dissociating Linguistic and Task-related Activity in the Left Inferior Frontal Gyrus. J Cogn Neurosci. 23:404–413.
Wu H, Mai X, Tang H, Ge Y, Luo Y-J, Liu C. 2013. Dissociable Somatotopic Representations of Chinese Action Verbs in the Motor and Premotor Cortex. Sci Rep. 3:2049.
Xiao Z, Zhang JX, Wang X, Wu R, Hu X, Weng X, Tan LH. 2005. Differential activity in left inferior frontal gyrus for pseudowords and real words: An event-related fMRI study on auditory lexical decision. Hum Brain Mapp. 25:212–221.
Yang J, Li P, Fang X, Shu H, Liu Y, Chen L. 2016. Hemispheric involvement in the processing of Chinese idioms: An fMRI study. Neuropsychologia. 87:12–24.
Zaccarella E, Friederici AD. 2015. Merge in the Human Brain: A Sub-Region Based Functional Investigation in the Left Pars Opercularis. Front Psychol. 6:1–9.
Zhang H, Liu J, Zhang Q. 2014. Neural representations for the generation of inventive conceptions inspired by adaptive feature optimization of biological species. Cortex. 50:162–173.
Zhang JX, Xiao Z, Weng X. 2012. Neural evidence for direct meaning access from orthography in Chinese word reading. Int J Psychophysiol. 84:240–245.
Zhuang J, Devereux BJ. 2017. Phonological and syntactic competition effects in spoken word recognition: evidence from corpus-based statistics. Lang Cogn Neurosci. 32:221–235.
Zou L, Packard JL, Xia Z, Liu Y, Shu H. 2016. Neural correlates of morphological processing: Evidence from Chinese. Front Hum Neurosci. 9:1–12.
Zvyagintsev M, Clemens B, Chechko N, Mathiak KA, Sack AT, Mathiak K. 2013. Brain networks underlying mental imagery of auditory and visual information. Eur J Neurosci. 37:1421–1434.

#### *Theory of Mind References*

Abraham, A., Rakoczy, H., Werning, M., von Cramon, D. Y., & Schubotz, R. I. (2010). Matching mind to world and vice versa: Functional dissociations between belief and desire mental state processing. *Social Neuroscience*, *5*(1), 1–18. https://doi.org/10.1080/17470910903166853

Abraham, A., Werning, M., Rakoczy, H., von Cramon, D. Y., & Schubotz, R. I. (2008). Minds, persons, and space: An fMRI investigation into the relational complexity of higher-order inteAbraham, A., Werning, M., Rakoczy, H., von Cramon, D. Y., & Schubotz, R. I. (2008). Minds, persons, and space: An fMRI investigation into the relational co. *Consciousness and Cognition*, *17*(2), 438–450. https://doi.org/10.1016/j.concog.2008.03.011

Adams, R. B., Rule, N. O., Franklin, R. G., Wang, E., Stevenson, M. T., Yoshikawa, S., Nomura, M., Sato, W., Kveraga, K., & Ambady, N. (2010). Cross-cultural reading the mind in the eyes: An fMRI investigation. *Journal of Cognitive Neuroscience*, *22*(1), 97–108. https://doi.org/10.1162/jocn.2009.21187

Aichhorn, M., Perner, J., Weiss, B., Kronbichler, M., Staffen, W., & Ladurner, G. (2009). Temporo-parietal junction activity in theory-of-mind tasks: Falseness, beliefs, or attention. *Journal of Cognitive Neuroscience*, *21*(6), 1179–1192. https://doi.org/10.1162/jocn.2009.21082

Alderson-Day, B., Weis, S., Mccarthy-Jones, S., Moseley, P., Smailes, D., & Fernyhough, C. (2016). The brain's conversation with itself: neural substrates of dialogic inner speech. *Social Cognitive and Affective Neuroscience*, *11*(1), 110–120. https://doi.org/10.1093/scan/nsv094

Bahnemann, M., Dziobek, I., Prehn, K., … I. W.-S. cognitive and, & 2010, U. (2010). Sociotopy in the temporoparietal cortex: common versus distinct processes. *Social Cognitive and Affective Neuroscience*, *5*(1), 48–58. https://doi.org/10.1093/scan/nsp045

Baron-Cohen, S., Ring, H. A., Wheelwright, S., Bullmore, E. T., Brammer, M. J., Simmons, A., & Williams, S. C. R. (1999). Social intelligence in the normal and autistic brain: An fMRI study. *European Journal of Neuroscience*, *11*(6), 1891–1898. https://doi.org/10.1046/j.1460-9568.1999.00621.x

Bartholomeusz, C. F., Ganella, E. P., Whittle, S., Allott, K., Thompson, A., Abu-Akel, A., Walter, H., McGorry, P., Killackey, E., Pantelis, C., & Wood, S. J. (2018). An fMRI study of theory of mind in individuals with first episode psychosis. *Psychiatry Research - Neuroimaging*, *281*, 1–11. https://doi.org/10.1016/j.pscychresns.2018.08.011

Bliksted, V., Frith, C., Videbech, P., Fagerlund, B., Emborg, C., Simonsen, A., Roepstorff, A., & Campbell-Meiklejohn, D. (2019). Hyper-and Hypomentalizing in Patients with First-Episode Schizophrenia: fMRI and Behavioral Studies. *Schizophrenia Bulletin*, *45*(2), 377–385. https://doi.org/10.1093/schbul/sby027

Bodden, M. E., Kübler, D., Knake, S., Menzler, K., Heverhagen, J. T., Sommer, J., Kalbe, E., Krach, S., & Dodel, R. (2013). Comparing the neural correlates of affective and cognitive theory of mind using fMRI: Involvement of the basal ganglia in affective theory of mind. *Advances in Cognitive Psychology*, *9*(1), 32–43. https://doi.org/10.2478/v10053-008-0129-6

Briend, F., Marzloff, V., Brazo, P., Lecardeur, L., Leroux, E., Razafimandimby, A., & Dollfus, S. (2019). Social cognition in schizophrenia: Validation of an ecological fMRI task. *Psychiatry Research - Neuroimaging*, *286*, 60–68. https://doi.org/10.1016/j.pscychresns.2019.03.004

Brüne, M., Lissek, S., Fuchs, N., Witthaus, H., Peters, S., Nicolas, V., Juckel, G., & Tegenthoff, M. (2008). An fMRI study of theory of mind in schizophrenic patients with "passivity" symptoms. *Neuropsychologia*, *46*(7), 1992–2001. https://doi.org/10.1016/j.neuropsychologia.2008.01.023

Brunet, E., Sarfati, Y., Hardy-Baylé, M. C., & Decety, J. (2000). A PET investigation of the attribution of intentions with a nonverbal task. *NeuroImage*, *11*(2), 157–166. https://doi.org/10.1006/nimg.1999.0525

Canessa, N., Alemanno, F., Riva, F., Zani, A., Proverbio, A. M., Mannara, N., Perani, D., & Cappa, S. F. (2012). The neural bases of social intention understanding: The role of interaction goals. *PLoS ONE*, *7*(7), e42347. https://doi.org/10.1371/journal.pone.0042347

Cassidy, B. S., Hughes, C., & Krendl, A. C. (2020). Age differences in neural activity related to mentalizing during person perception. *Aging, Neuropsychology, and Cognition*, 1–18. https://doi.org/10.1080/13825585.2020.1718060

Castelli, F., Happé, F., Frith, U., & Frith, C. (2000). Movement and mind: A functional imaging study of perception and interpretation of complex intentional movement patterns. *NeuroImage*, *12*(3), 314–325. https://doi.org/10.1006/nimg.2000.0612

Castelli, I., Baglio, F., Blasi, V., Alberoni, M., Falini, A., Liverta-Sempio, O., Nemni, R., & Marchetti, A. (2010). Effects of aging on mindreading ability through the eyes: An fMRI study. *Neuropsychologia*, *48*(9), 2586–2594. https://doi.org/10.1016/j.neuropsychologia.2010.05.005

Chakroff, A., Dungan, J., … J. K.-H.-S. cognitive and, & 2016, U. (2016). When minds matter for moral judgment: intent information is neurally encoded for harmful but not impure acts. *Social Cognitive and Affective Neuroscience*, *11*(3), 476–484. https://doi.org/10.1093/scan/nsv131

Cheung, H., Chen, L., Szeto, C. Y., Feng, G., Lu, G., Zhang, Z., Zhu, Z., & Wang, S. (2012). False belief and verb non-factivity: A common neural basis? *International Journal of Psychophysiology*, *83*(3), 357–364. https://doi.org/10.1016/j.ijpsycho.2011.12.002

Cole, E. J., Barraclough, N. E., & Andrews, T. J. (2019). Reduced connectivity between mentalizing and mirror systems in autism spectrum condition. *Neuropsychologia*, *122*, 88–97. https://doi.org/10.1016/j.neuropsychologia.2018.11.008

Contreras, J. M., Schirmer, J., Banaji, M. R., & Mitchell, J. P. (2013). Common Brain Regions with Distinct Patterns of Neural Responses during Mentalizing about Groups and Individuals. *Journal of Cognitive Neuroscience*, *25*(9), 1406–1417. https://doi.org/10.1162/jocn_a_00403

Corradi-Dell'Acqua, C., … C. H.-S. C. and, & 2014, U. (2014). Cognitive and affective theory of mind share the same local patterns of activity in posterior temporal but not medial prefrontal cortex. *Social Cognitive and Affective Neuroscience*, *9*(8), 1175–1184. https://doi.org/10.1093/scan/nst097

Das, P., Lagopoulos, J., Coulston, C. M., Henderson, A. F., & Malhi, G. S. (2012). Mentalizing impairment in schizophrenia: A functional MRI study. *Schizophrenia Research*, *134*(2–3), 158–164. https://doi.org/10.1016/j.schres.2011.08.019

de Achával, D., Villarreal, M. F., Costanzo, E. Y., Douer, J., Castro, M. N., Mora, M. C., Nemeroff, C. B., Chu, E., Bär, K. J., & Guinjoan, S. M. (2012). Decreased activity in right-hemisphere structures involved in social cognition in siblings discordant for schizophrenia. *Schizophrenia Research*, *134*(2–3), 171–179. https://doi.org/10.1016/j.schres.2011.11.010

Deuse, L., Rademacher, L., … L. W.-S. cognitive and, & 2016, U. (2016). Neural correlates of naturalistic social cognition: brain-behavior relationships in healthy adults. *Social Cognitive and Affective Neuroscience*, *11*(11), 1741–1751. https://doi.org/10.1093/scan/nsw094

Dodell-Feder, D, … L. D.-S. cognitive and, & 2014, U. (2014). Neural disruption to theory of mind predicts daily social functioning in individuals at familial high-risk for schizophrenia. *Social Cognitive and Affective Neuroscience*, *9*(12), 1914–1925. https://doi.org/10.1093/scan/nst186

Dodell-Feder, David, Koster-Hale, J., Bedny, M., & Saxe, R. (2011). FMRI item analysis in a theory of mind task. *NeuroImage*, *55*(2), 705–712. https://doi.org/10.1016/j.neuroimage.2010.12.040

Döhnel, K., Schuwerk, T., Meinhardt, J., Sodian, B., Hajak, G., & Sommer, M. (2012). Functional activity of the right temporo-parietal junction and of the medial prefrontal cortex associated with true and false belief reasoning. *NeuroImage*, *60*(3), 1652–1661. https://doi.org/10.1016/j.neuroimage.2012.01.073

Dufour, N., Redcay, E., Young, L., Mavros, P., One, J. M.-P., & 2013, U. (2013). Similar brain activation during false belief tasks in a large sample of adults with and without autism. *PLoS One*, *8*(9), e75468. https://doi.org/10.1371/journal.pone.0075468

Ferstl, E. C., & Von Cramon, D. Y. (2002). What does the frontomedian cortex contribute to language processing: Coherence or theory of mind? *NeuroImage*, *17*(3), 1599–1612. https://doi.org/10.1006/nimg.2002.1247

Fletcher, P. C., Happé, F., Frith, U., Baker, S. C., Dolan, R. J., Frackowiak, R. S. J., & Frith, C. D. (1995). Other minds in the brain: a functional imaging study of "theory of mind" in story comprehension. *Cognition*, *57*(2), 109–128. https://doi.org/10.1016/0010-0277(95)00692-R

Focquaert, F., Steven-Wheeler, M. S., Vanneste, S., Doron, K. W., & Platek, S. M. (2010). Mindreading in individuals with an empathizing versus systemizing cognitive style: An fMRI study. *Brain Research Bulletin*, *83*(5), 214–222. https://doi.org/10.1016/j.brainresbull.2010.08.008

Gallagher, H. L., Happé, F., Brunswick, N., Fletcher, P. C., Frith, U., & Frith, C. D. (2000). Reading the mind in cartoons and stories: An fMRI study of "theory of mind" in verbal and nonverbal tasks. *Neuropsychologia*, *38*(1), 11–21. https://doi.org/10.1016/S0028-3932(99)00053-6

Geiger, A., Bente, G., Lammers, S., Tepest, R., Roth, D., Bzdok, D., & Vogeley, K. (2019). Distinct functional roles of the mirror neuron system and the mentalizing system. *NeuroImage*, *202*, 116102. https://doi.org/10.1016/j.neuroimage.2019.116102

Gobbini, M. I., Koralek, A. C., Bryan, R. E., Montgomery, K. J., & Haxby, J. V. (2007). Two takes on the social brain: A comparison of theory of mind tasks. *Journal of Cognitive Neuroscience*, *19*(11), 1803–1814. https://doi.org/10.1162/jocn.2007.19.11.1803

Gweon, H., Dodell-Feder, D., Bedny, M., & Saxe, R. (2012). Theory of Mind Performance in Children Correlates With Functional Specialization of a Brain Region for Thinking About Thoughts. *Child Development*, *83*(6), 1853–1868. https://doi.org/10.1111/j.1467-8624.2012.01829.x

Hartwright, C. E., Apperly, I. A., & Hansen, P. C. (2015). The special case of self-perspective inhibition in mental, but not non-mental, representation. *Neuropsychologia*, *67*, 183–192. https://doi.org/10.1016/j.neuropsychologia.2014.12.015

Hervé, P. Y., Razafimandimby, A., Jobard, G., & Tzourio-Mazoyer, N. (2013). A Shared Neural Substrate for Mentalizing and the Affective Component of Sentence Comprehension. *PLoS ONE*, *8*(1). https://doi.org/10.1371/journal.pone.0054400

Hooker, C. I., Verosky, S. C., Germine, L. T., Knight, R. T., & D'Esposito, M. (2010). Neural activity during social signal perception correlates with self-reported empathy. *Brain Research*, *1308*, 100–113. https://doi.org/10.1016/j.brainres.2009.10.006

Hooker, C., Verosky, S., … L. G.-S. cognitive and, & 2008, U. (2008). Mentalizing about emotion and its relationship to empathy. *Social Cognitive and Affective Neuroscience*, *3*(3), 204–217. https://doi.org/10.1093/scan/nsn019

Jack, A., Cortex, K. P.-C., & 2015, U. (2015). Neural correlates of animacy attribution include neocerebellum in healthy adults. *Cerebral Cortex*, *25*(11), 4240–4247. https://doi.org/10.1093/cercor/bhu146

Jacoby, N., Bruneau, E., Koster-Hale, J., & Saxe, R. (2016). Localizing Pain Matrix and Theory of Mind networks with both verbal and non-verbal stimuli. *NeuroImage*, *126*, 39–48. https://doi.org/10.1016/j.neuroimage.2015.11.025

Jenkins, A. C., Dodell-Feder, D., Saxe, R., & Knobe, J. (2014). The neural bases of directed and spontaneous mental state attributions to group agents. *PLoS ONE*, *9*(8), e105341. https://doi.org/10.1371/journal.pone.0105341

Jenkins, A. C., & Mitchell, J. P. (2010). Mentalizing under Uncertainty: Dissociated Neural Responses to Ambiguous and Unambiguous Mental State Inferences. *Cerebral Cortex*, *20*, 404–410. https://doi.org/10.1093/cercor/bhp109

Jimura, K., Konishi, S., Asari, T., & Miyashita, Y. (2010). Temporal pole activity during understanding other persons' mental states correlates with neuroticism trait. *Brain Research*, *1328*, 104–112. https://doi.org/10.1016/j.brainres.2010.03.016

Kana, R. K., Keller, T. A., Cherkassky, V. L., Minshew, N. J., & Just, M. A. (2009). Atypical frontal-posterior synchronization of Theory of Mind regions in autism during mental state attribution. *Social Neuroscience*, *4*(2), 135–152. https://doi.org/10.1080/17470910802198510

Kandylaki, K. D., Nagels, A., Tune, S., Wiese, R., Bornkessel-Schlesewsky, I., & Kircher, T. (2015). Processing of false belief passages during natural story comprehension: An fMRI study. *Human Brain Mapping*, *36*(11), 4231–4246. https://doi.org/10.1002/hbm.22907

Kanske, P., Böckler, A., Trautwein, F. M., & Singer, T. (2015). Dissecting the social brain: Introducing the EmpaToM to reveal distinct neural networks and brain-behavior relations for empathy and Theory of Mind. *NeuroImage*, *122*, 6–19. https://doi.org/10.1016/j.neuroimage.2015.07.082

Kirkovski, M., Enticott, P. G., Hughes, M. E., Rossell, S. L., & Fitzgerald, P. B. (2016). Atypical Neural Activity in Males But Not Females with Autism Spectrum Disorder. *Journal of Autism and Developmental Disorders*, *46*(3), 954–963. https://doi.org/10.1007/s10803-015-2639-7

Kliemann, D., Young, L., Scholz, J., & Saxe, R. (2008). The influence of prior record on moral judgment. *Neuropsychologia*, *46*(12), 2949–2957. https://doi.org/10.1016/j.neuropsychologia.2008.06.010

Kobayashi, C., Glover, G. H., & Temple, E. (2006). Cultural and linguistic influence on neural bases of "Theory of Mind": An fMRI study with Japanese bilinguals. *Brain and Language*, *98*(2), 210–220. https://doi.org/10.1016/j.bandl.2006.04.013

Kobayashi, C., Glover, G. H., & Temple, E. (2007a). Children's and adults' neural bases of verbal and nonverbal 'theory of mind.' *Neuropsychologia*, *45*(7), 1522–1532. https://doi.org/10.1016/j.neuropsychologia.2006.11.017

Kobayashi, C., Glover, G. H., & Temple, E. (2007b). Cultural and linguistic effects on neural bases of 'Theory of Mind' in American and Japanese children. *Brain Research*, *1164*(1), 95–107. https://doi.org/10.1016/j.brainres.2007.06.022

Koelkebeck, K., Hirao, K., Kawada, R., Miyata, J., Saze, T., Ubukata, S., Itakura, S., Kanakogi, Y., Ohrmann, P., Bauer, J., Pedersen, A., Sawamoto, N., Fukuyama, H., Takahashi, H., & Murai, T. (2011). Transcultural differences in brain activation patterns during theory of mind (ToM) task performance in Japanese and Caucasian participants. *Social Neuroscience*, *6*(5–6), 615–626. https://doi.org/10.1080/17470919.2011.620763

Lavoie, M. A., Vistoli, D., Sutliff, S., Jackson, P. L., & Achim, A. M. (2016). Social representations and contextual adjustments as two distinct components of the Theory of Mind brain network: Evidence from the REMICS task. *Cortex*, *81*, 176–191. https://doi.org/10.1016/j.cortex.2016.04.017

Lee, S., Cortex, G. M.-C., & 2016, U. (2016). Functional heterogeneity and convergence in the right temporoparietal junction. *Cerebral Cortex*, *26*(3), 1108–1116. https://doi.org/10.1093/cercor/bhu292

Lewis, P., Birch, A., … A. H.-S. cognitive and, & 2017, U. (2017). Higher order intentionality tasks are cognitively more demanding. *Social Cognitive and Affective Neuroscience*, *12*(7), 1063–1071. https://doi.org/10.1093/scan/nsx034

Libero, L. E., Maximo, J. O., Deshpande, H. D., Klinger, L. G., Klinger, M. K., & Kana, R. K. (2014). The role of mirroring and mentalizing networks in mediating action intentions in autism. *Molecular Autism*, *5*(1), 50. https://doi.org/10.1186/2040-2392-5-50

Lin, N., Yang, X., Li, J., Wang, S., Hua, H., Ma, Y., & Li, X. (2018). Neural correlates of three cognitive processes involved in theory of mind and discourse comprehension. *Cognitive, Affective and Behavioral Neuroscience*, *18*(2), 273–283. https://doi.org/10.3758/s13415-018-0568-6

Malhi, G. S., Lagopoulos, J., Das, P., Moss, K., Berk, M., & Coulston, C. M. (2008). A functional MRI study of Theory of Mind in euthymic bipolar disorder patients. *Bipolar Disorders*, *10*(8), 943–956. https://doi.org/10.1111/j.1399-5618.2008.00643.x

Marjoram, D., Job, D. E., Whalley, H. C., Gountouna, V. E., McIntosh, A. M., Simonotto, E., Cunningham-Owens, D., Johnstone, E. C., & Lawrie, S. (2006). A visual joke fMRI investigation into Theory of Mind and enhanced risk of schizophrenia. *NeuroImage*, *31*(4), 1850–1858. https://doi.org/10.1016/j.neuroimage.2006.02.011

Martin, A., & Weisberg, J. (2003). Neural foundations for understanding social and mechanical concepts. *Cognitive Neuropsychology*, *20*(3–6), 575–587. https://doi.org/10.1080/02643290342000005

Mason, R. A., Williams, D. L., Kana, R. K., Minshew, N., & Just, M. A. (2008). Theory of Mind disruption and recruitment of the right hemisphere during narrative comprehension in autism. *Neuropsychologia*, *46*(1), 269–280. https://doi.org/10.1016/j.neuropsychologia.2007.07.018

McAdams, C. J., & Krawczyk, D. C. (2013). Neural responses during social and self-knowledge tasks in Bulimia nervosa. *Frontiers in Psychiatry*, *4*, 103. https://doi.org/10.3389/fpsyt.2013.00103

Mier, D., Sauer, C., Lis, S., Esslinger, C., Wilhelm, J., Gallhofer, B., & Kirsch, P. (2020). Neuronal correlates of affective theory of mind in schizophrenia out-patients: evidence for a baseline deficit. *Psychological Medicine*, *40*(10), 1607–1617. https://doi.org/10.1017/S0033291709992133

Mitchell, J. P. (2008). Activity in Right Temporo-Parietal Junction is Not Selective for Theory-of-Mind. *Cerebral Cortex*, *18*(2), 262–271. https://doi.org/10.1093/cercor/bhm051

Modinos, G., Renken, R., Shamay-Tsoory, S. G., Ormel, J., & Aleman, A. (2010). Neurobiological correlates of theory of mind in psychosis proneness. *Neuropsychologia*, *48*(13), 3715–3724. https://doi.org/10.1016/j.neuropsychologia.2010.09.030

Moessnang, C., Schäfer, A., … E. B.-S. cognitive and, & 2016, U. (2016). Specificity, reliability and sensitivity of social brain responses during spontaneous mentalizing. *Social Cognitive and Affective Neuroscience*, *11*(11), 1687–1697. https://doi.org/10.1093/scan/nsw098

Mohnke, S., Erk, S., … K. S.-S. cognitive and, & 2016, U. (2016). Theory of mind network activity is altered in subjects with familial liability for schizophrenia. *Social Cognitive and Affective Neuroscience*, *11*(2), 299–307. https://doi.org/10.1093/scan/nsv111

Moran, J. M., Jolly, E., & Mitchell, J. P. (2012). Social-cognitive deficits in normal aging. *Journal of Neuroscience*, *32*(16), 5553–5561. https://doi.org/10.1523/jneurosci.5511-11.2012

Naughtin, C. K., Horne, K., Schneider, D., Venini, D., York, A., & Dux, P. E. (2017). Do implicit and explicit belief processing share neural substrates? *Human Brain Mapping*, *38*(9), 4760–4772. https://doi.org/10.1002/hbm.23700

Nieminen-Von Wendt, T., Metsähonkala, L., Kulomäki, T., Aalto, S., Raija, T. A., Lennart Von Wendt, V., Metsähonkala, L., Aalto, · S, & Autti, T. (2003). Changes in cerebral blood flow in Asperger syndrome during theory of mind tasks presented by the auditory route. *European Child & Adolescent Psychiatry*, *12*(4), 178–189. https://doi.org/10.1007/s00787-003-0337-z

Oliver, L., Vieira, J., … R. N.-S. cognitive and, & 2018, U. (2018). Greater involvement of action simulation mechanisms in emotional vs cognitive empathy. *Social Cognitive and Affective Neuroscience*, *13*(4), 367–380. https://doi.org/10.1093/scan/nsy013

Otsuka, Y., Osaka, N., Ikeda, T., & Osaka, M. (2009). Individual differences in the theory of mind and superior temporal sulcus. *Neuroscience Letters*, *463*(2), 150–153. https://doi.org/10.1016/j.neulet.2009.07.064

Otti, A., Wohlschlaeger, A. M., & Noll-Hussong, M. (2015). Is the medial prefrontal cortex necessary for theory of mind? *PLoS ONE*, *10*(8). https://doi.org/10.1371/journal.pone.0135912

Overgaauw, S., … A. van D.-S. C. and, & 2015, U. (2015). A longitudinal analysis of neural regions involved in reading the mind in the eyes. *Social Cognitive and Affective Neuroscience*, *10*(5), 619–627. https://doi.org/10.1093/scan/nsu095

Perner, J., Aichhorn, M., Kronbichler, M., Staffen, W., & Ladurner, G. (2006). Thinking of mental and other representations: the roles of left and right temporo-parietal junction. *Social Neuroscience*, *1*(3–4), 245–258. https://doi.org/10.1080/17470910600989896

Platek, S. M., Keenan, J. P., Gallup, G. G., & Mohamed, F. B. (2004). Where am I? The neurological correlates of self and other. *Cognitive Brain Research*, *19*(2), 114–122. https://doi.org/10.1016/j.cogbrainres.2003.11.014

Powell, J. L., Grossi, D., Corcoran, R., Gobet, F., & García-Fiñana, M. (2017). The neural correlates of theory of mind and their role during empathy and the game of chess: A functional magnetic resonance imaging study. *Neuroscience*, *355*, 149–160. https://doi.org/10.1016/j.neuroscience.2017.04.042

Roser, P., Lissek, S., Tegenthoff, M., Nicolas, V., Juckel, G., & Brüne, M. (2012). Alterations of theory of mind network activation in chronic cannabis users. *Schizophrenia Research*, *139*(1–3), 19–26. https://doi.org/10.1016/j.schres.2012.05.020

Ross, L. A., & Olson, I. R. (2010). Social cognition and the anterior temporal lobes. *NeuroImage*, *49*(4), 3452–3462. https://doi.org/10.1016/j.neuroimage.2009.11.012

Russell, T. A., Rubia, K., Bullmore, E. T., Soni, W., Suckling, J., Brammer, M. J., Simmons, A., Williams, S. C. R., & Sharma, T. (2000). Exploring the social brain in schizophrenia: Left prefrontal underactivation during mental state attribution. *American Journal of Psychiatry*, *157*(12), 2040–2042. https://doi.org/10.1176/appi.ajp.157.12.2040

Saft, C., Lissek, S., Hoffmann, R., Nicolas, V., Tegenthoff, M., Juckel, G., & Brüne, M. (2013). Mentalizing in preclinical Huntington's disease: an fMRI study using cartoon picture stories. *Brain Imaging and Behavior*, *7*(2), 154–162. https://doi.org/10.1007/s11682-012-9209-9

Samson, A. C., Zysset, S., & Huber, O. (2008). Cognitive humor processing: Different logical mechanisms in nonverbal cartoons - An fMRI study. *Social Neuroscience*, *3*(2), 125–140. https://doi.org/10.1080/17470910701745858

Saxe, R., & Kanwisher, N. (2003). People thinking about thinking people: The role of the temporo-parietal junction in "theory of mind." *NeuroImage*, *19*(4), 1835–1842. https://doi.org/10.1016/S1053-8119(03)00230-1

Saxe, Rebecca, & Powell, L. J. (2006). It's the thought that counts: Specific brain regions for one component of theory of mind. *Psychological Science*, *17*(8), 692–699. https://doi.org/10.1111/j.1467-9280.2006.01768.x

Saxe, Rebecca, Schulz, L. E., & Jiang, Y. V. (2006). Reading minds versus following rules: dissociating theory of mind and executive control in the brain. *Social Neuroscience*, *1*(3–4), 284–298. https://doi.org/10.1080/17470910601000446

Schiffer, B., Pawliczek, C., Müller, B. W., Gizewski, E. R., & Walter, H. (2013). Why Don't Men Understand Women? Altered Neural Networks for Reading the Language of Male and Female Eyes. *PLoS ONE*, *8*(4), e60278. https://doi.org/10.1371/journal.pone.0060278

Schlaffke, L., Lissek, S., Lenz, M., Juckel, G., Schultz, T., Tegenthoff, M., Schmidt-Wilcke, T., & Brune, M. (2015). Shared and nonshared neural networks of cognitive and affective theory-of-mind: A neuroimaging study using cartoon picture stories. *Human Brain Mapping*, *36*(1), 29–39. https://doi.org/10.1002/hbm.22610

Schmitgen, M. M., Walter, H., Drost, S., Rückl, S., & Schnell, K. (2016). Stimulus-dependent amygdala involvement in affective theory of mind generation. *NeuroImage*, *129*, 450–459. https://doi.org/10.1016/j.neuroimage.2016.01.029

Schneider, D., Slaughter, V. P., Becker, S. I., & Dux, P. E. (2014). Implicit false-belief processing in the human brain. *NeuroImage*, *101*, 268–275. https://doi.org/10.1016/j.neuroimage.2014.07.014

Shimada, K., Kasaba, R., Fujisawa, T. X., Sakakibara, N., Takiguchi, S., & Tomoda, A. (2018). Subclinical maternal depressive symptoms modulate right inferior frontal response to inferring affective mental states of adults but not of infants. *Journal of Affective Disorders*, *229*, 32–40. https://doi.org/10.1016/j.jad.2017.12.031

Sommer, M., Sodian, B., Döhnel, K., Schwerdtner, J., Meinhardt, J., & Hajak, G. (2010). In psychopathic patients emotion attribution modulates activity in outcome-related brain areas. *Psychiatry Research - Neuroimaging*, *182*(2), 88–95. https://doi.org/10.1016/j.pscychresns.2010.01.007

Specht, K., & Wigglesworth, P. (2018). The functional and structural asymmetries of the superior temporal sulcus. *Scandinavian Journal of Psychology*, *59*(1), 74–82. https://doi.org/10.1111/sjop.12410

Spunt, R. P., & Lieberman, M. D. (2012). Dissociating Modality-Specific and Supramodal Neural Systems for Action Understanding. *Journal of Neuroscience*, *32*(10), 3575–3583. https://doi.org/10.1523/JNEUROSCI.5715-11.2012

Spunt, Robert P., & Adolphs, R. (2014). Validating the Why/How contrast for functional MRI studies of Theory of Mind. *NeuroImage*, *99*, 301–311.

https://doi.org/10.1016/j.neuroimage.2014.05.023

Spunt, Robert P., & Lieberman, M. D. (2012). An integrative model of the neural systems supporting the comprehension of observed emotional behavior. *NeuroImage*, *59*(3), 3050–3059. https://doi.org/10.1016/j.neuroimage.2011.10.005

Spunt, Robert P., Satpute, A. B., & Lieberman, M. D. (2011). Identifying the What, Why, and How of an Observed Action: An fMRI Study of Mentalizing and Mechanizing during Action Observation. *Journal of Cognitive Neuroscience*, *23*(1), 63–74. https://doi.org/10.1162/jocn.2010.21446

Tholen, M. G., Trautwein, F. M., Böckler, A., Singer, T., & Kanske, P. (2020). Functional magnetic resonance imaging (fMRI) item analysis of empathy and theory of mind. *Human Brain Mapping*, 1–18. https://doi.org/10.1002/hbm.24966

Thye, M. D., Murdaugh, D. L., & Kana, R. K. (2018). Brain Mechanisms Underlying Reading the Mind from Eyes, Voice, and Actions. *Neuroscience*, *374*, 172–186. https://doi.org/10.1016/j.neuroscience.2018.01.045

Van der Meer, L., Groenewold, N. A., Nolen, W. A., Pijnenborg, M., & Aleman, A. (2011). Inhibit yourself and understand the other: Neural basis of distinct processes underlying Theory of Mind. *NeuroImage*, *56*(4), 2364–2374. https://doi.org/10.1016/j.neuroimage.2011.03.053

Van Hoeck, N., Begtas, E., Steen, J., Kestemont, J., Vandekerckhove, M., & Van Overwalle, F. (2014). False belief and counterfactual reasoning in a social environment. *NeuroImage*, *90*, 315–325. https://doi.org/10.1016/j.neuroimage.2013.12.043

Vanderwal, T., Hunyadi, E., Grupe, D. W., Connors, C. M., & Schultz, R. T. (2008). Self, mother and abstract other: An fMRI study of reflective social processing. *NeuroImage*, *41*(4), 1437–1446. https://doi.org/10.1016/j.neuroimage.2008.03.058

Vogeley, K., Bussfeld, P., Newen, A., Herrmann, S., Happé, F., Falkai, P., Maier, W., Shah, N. J., Fink, G. R., & Zilles, K. (2001). Mind Reading: Neural Mechanisms of Theory of Mind and Self-Perspective. *NeuroImage*, *14*(1), 170–181. https://doi.org/10.1006/nimg.2001.0789

Völlm, B. A., Taylor, A. N. W., Richardson, P., Corcoran, R., Stirling, J., McKie, S., Deakin, J. F. W., & Elliott, R. (2006). Neuronal correlates of theory of mind and empathy: A functional magnetic resonance imaging study in a nonverbal task. *NeuroImage*, *29*(1), 90–98. https://doi.org/10.1016/j.neuroimage.2005.07.022

Walter, H, Ciaramidaro, A., … M. A.-S. cognitive and, & 2009, U. (2009). Dysfunction of the social brain in schizophrenia is modulated by intention type: an fMRI study. *Social Cognitive and Affective Neuroscience*, *4*(2), 166–176. https://doi.org/10.1093/scan/nsn047

Walter, Henrik, Adenzato, M., Ciaramidaro, A., Enrici, I., Pia, L., & Bara, B. G. (2004). Understanding Intentions in Social Interaction: The Role of the Anterior Paracingulate Cortex. *Journal of Cognitive Neuroscience*, *16*(10), 1854–1863. https://doi.org/10.1162/0898929042947838

Wang, Y., Liu, W. H., Li, Z., Wei, X. H., Jiang, X. Q., Neumann, D. L., Shum, D. H. K., Cheung, E. F. C., & Chan, R. C. K. (2015). Dimensional schizotypy and social cognition: An fMRI imaging study. *Frontiers in Behavioral Neuroscience*, *9*, 133. https://doi.org/10.3389/fnbeh.2015.00133

Willert, A., Mohnke, S., Erk, S., Schnell, K., Romanczuk-Seiferth, N., Quinlivan, E., Schreiter, S., Spengler, S., Herold, D., Wackerhagen, C., Romund, L., Garbusow, M., Lett, T., Stamm, T., Adli, M., Heinz, A., Bermpohl, F., & Walter, H. (2015). Alterations in neural Theory of Mind processing in euthymic patients with bipolar disorder and unaffected relatives. *Bipolar Disorders*, *17*(8), 880–891. https://doi.org/10.1111/bdi.12352

Wolf, I., Dziobek, I., & Heekeren, H. R. (2010). Neural correlates of social cognition in naturalistic settings: A model-free analysis approach. *NeuroImage*, *49*(1), 894–904. https://doi.org/10.1016/j.neuroimage.2009.08.060

Young, L., Scholz, J., & Saxe, R. (2011). Neural evidence for “intuitive prosecution”: The use of mental state information for negative moral verdicts. *Social Neuroscience*, *6*(3), 302–315. https://doi.org/10.1080/17470919.2010.529712

Zaitchik, D., Walker, C., Miller, S., LaViolette, P., Feczko, E., & Dickerson, B. C. (2010). Mental state attribution and the temporoparietal junction: An fMRI study comparing belief, emotion, and perception. *Neuropsychologia*, *48*(9), 2528–2536. https://doi.org/10.1016/j.neuropsychologia.2010.04.031
