## Supplementary_Information_No.2_Results for "Overlapping Neural Correlates Underpin Theory of Mind and Semantic Cognition: Evidence from a Meta-Analysis of 344 Functional Neuroimaging Studies"

for

### Investigating the Similarities in Neural Networks Underpinning Theory of Mind and Semantic Cognition: A Meta-Analysis

This document contains the results figures and tables for all analyses conducted and not listed in the main text. The input files and outputs of all analyses can be accessed via OSF (https://osf.io/ydnxh/).

*Supplementary Figures:*

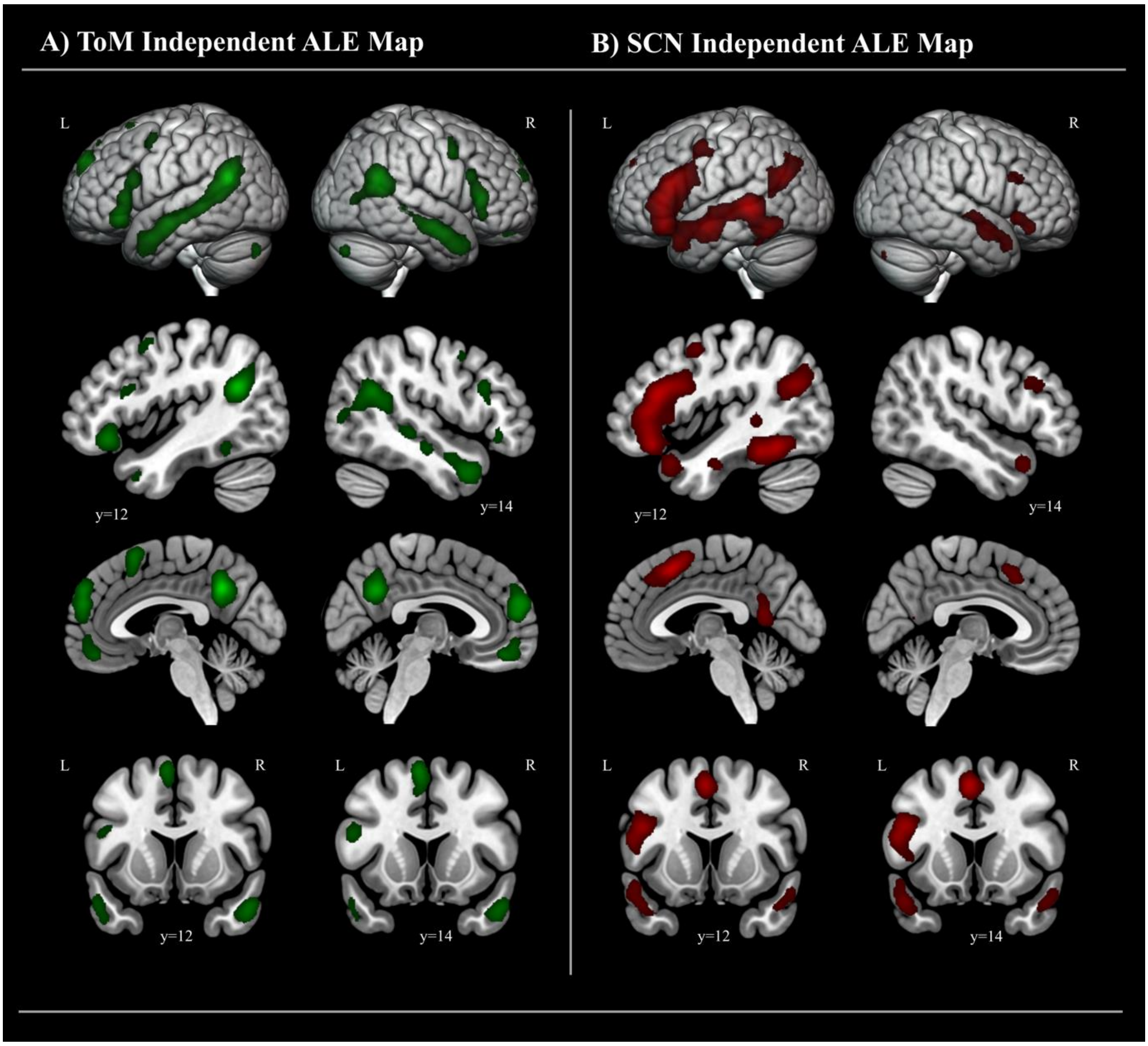

***Supplementary Figure R1*** *Independent ALE activation maps for ToM (N= 113) and SC (N=211); The maps were treated to a cluster forming threshold at p<.001 and an FWE corrected cluster-extent threshold at p<.05. The sagittal and coronal sections are chosen as representative slices positioned over peak coordinates at which there is the greatest conjunction in the bilateral anterior temporal lobes (left y= 12; right y= 14).*

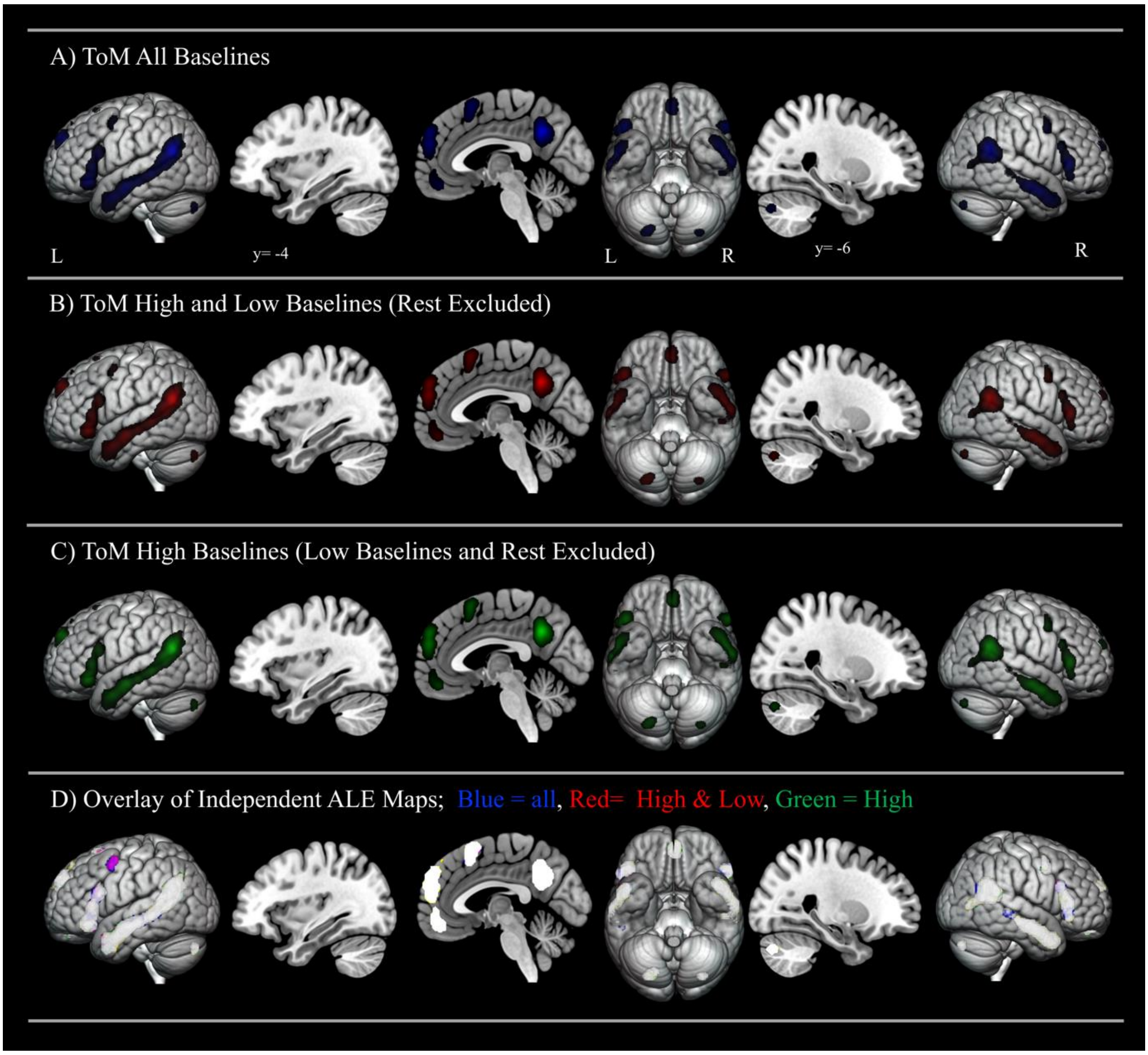

***Supplementary Figure R2a Panels A-C**: Independent ALE activation maps for three ToM data subsets (all N= 114; high & low N= 113; high N = 111) with experiments using different baseline matching levels. The maps were treated to a cluster forming threshold at p<.001 and an FWE corrected cluster-extent threshold at p<.05. **Panel D**: Binarized and overlayed independent ALE maps showing overlap and capturing the similarities and differences across the ToM data subsets. The sagittal and coronal sections (y= 4) correspond to peak ALE coordinates representing the most notable differences in areas of interest located in the medial and anterior temporal lobes in the SC data.*

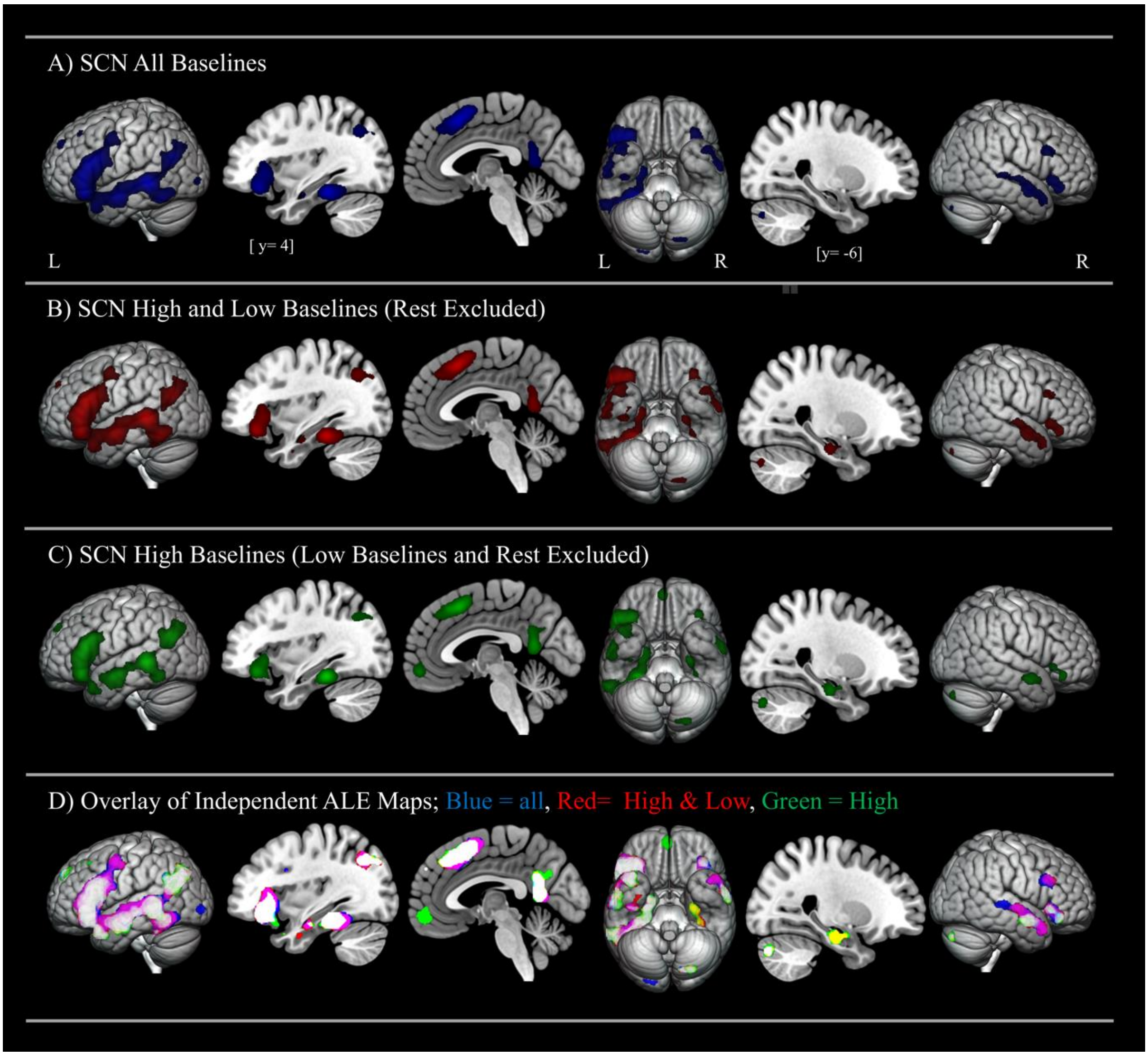

***Supplementary Figure R2b Panels A-C****: Independent ALE activation maps for three SC data subsets (all N= 214; high & low N= 211; high N = 170) with experiments using different baseline matching levels. The maps were treated to a cluster forming threshold at p<.001 and an FWE corrected cluster-extent threshold at p<.05.* ***Panel D****: Binarized and overlayed independent ALE maps showing overlap and capturing the similarities and differences across the SC data subsets. The sagittal and coronal sections (y= 4) correspond to peak ALE coordinates representing the most notable differences in areas of interest in the medial and anterior temporal lobes.*

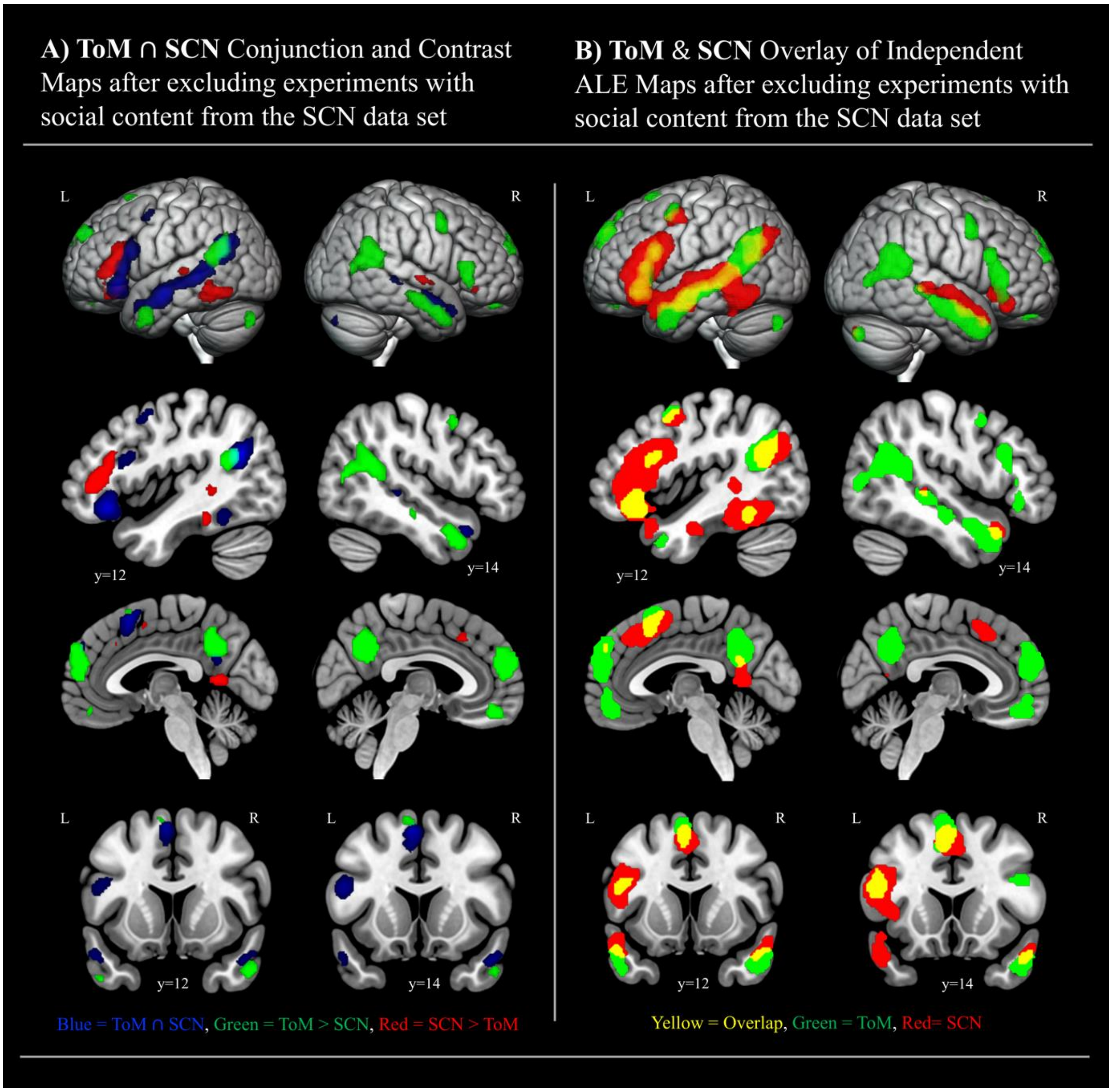

***Supplementary Figure R3** Formal and overlay conjunction and contrast analyses showing common converging and differential activation for Non-Social SC (N=193) and ToM (N= 113) networks after controlling for stimuli with a degree of social content in the SC data. The initial ALE maps were treated to a cluster-forming threshold at p<.001 and an FWE-corrected cluster-extent threshold at p<.05 prior to the conjunction and contrast analysis. The contrast maps in **Panel A** were additionally thresholded with a cluster-forming threshold at p<.001 and a minimum cluster size of 100mm$^3$. **Panel A** displays the conjunction alongside side statistically significant differences. In **Panel B,** we have overlaid the binarised versions of the complete*

*ALE maps resulting from independent analysis of Non-Social SC and ToM studies. This allows for full visualisation of the topography of the associated networks. The sagittal and coronal sections are chosen as representative slices positioned over peak coordinates at which there is the greatest conjunction in the bilateral anterior temporal lobes (left y= 12; right y= 14).*

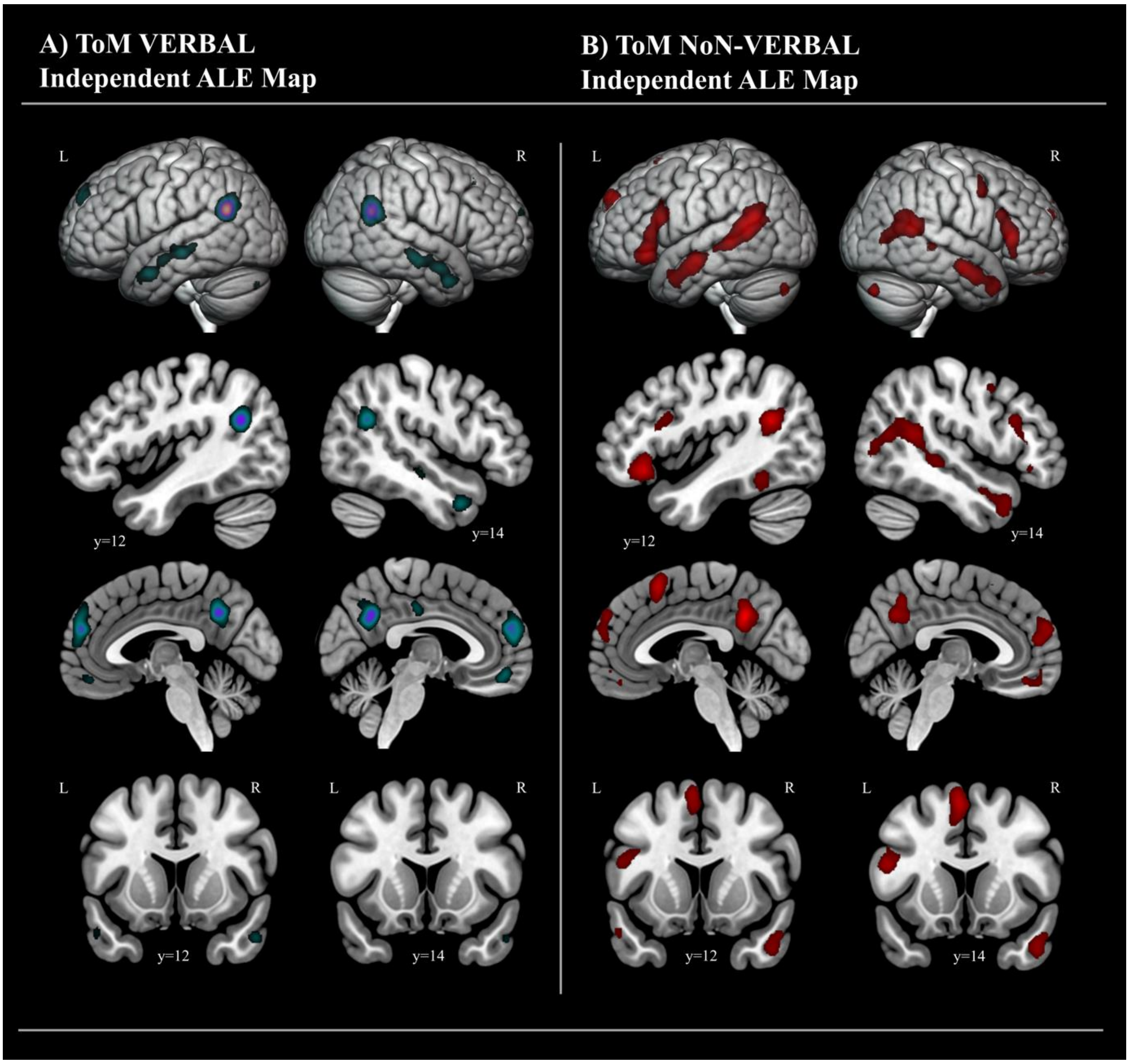

***Supplementary Figure R4*** *Independent ALE activation maps for VERBAL ToM (N= 46) and NoN-VERBAL ToM (N=71); The maps were treated to a cluster forming threshold at p<.001 and an FWE corrected cluster-extent threshold at p<.05. The sagittal and coronal sections are chosen as representative slices positioned over peak coordinates at which there is the greatest conjunction in the bilateral anterior temporal lobes (left y= 12; right y= 14).*

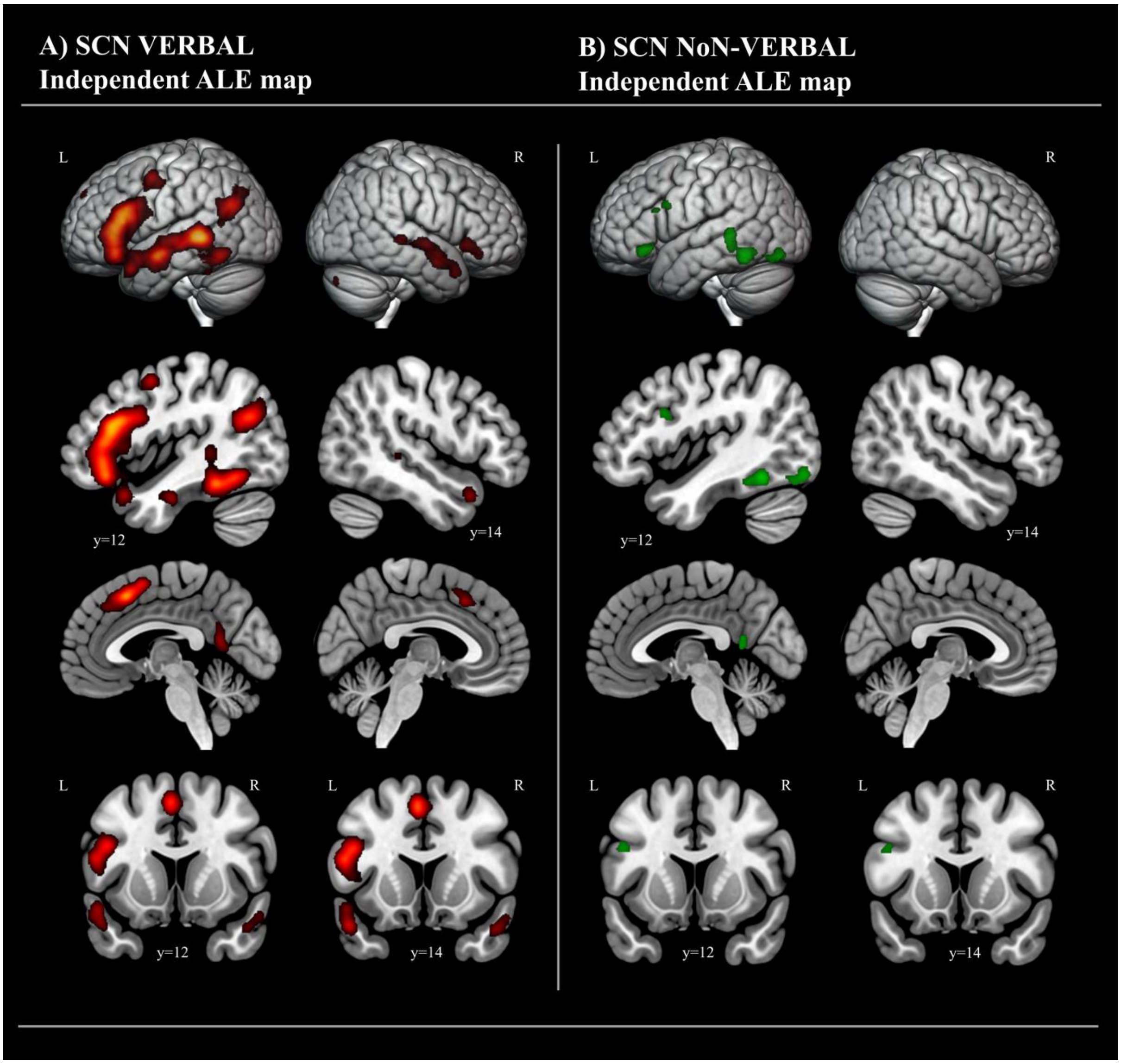

***Supplementary Figure R5*** *Independent ALE activation maps for VERBAL SC (N= 175) and NON-VERBAL SC (N=37); The maps were treated to a cluster forming threshold at p<.001 and an FWE corrected cluster-extent threshold at p<.05. The sagittal and coronal sections are chosen as representative slices positioned over peak coordinates at which there is the greatest conjunction in the bilateral anterior temporal lobes (left y= 12; right y= 14).*

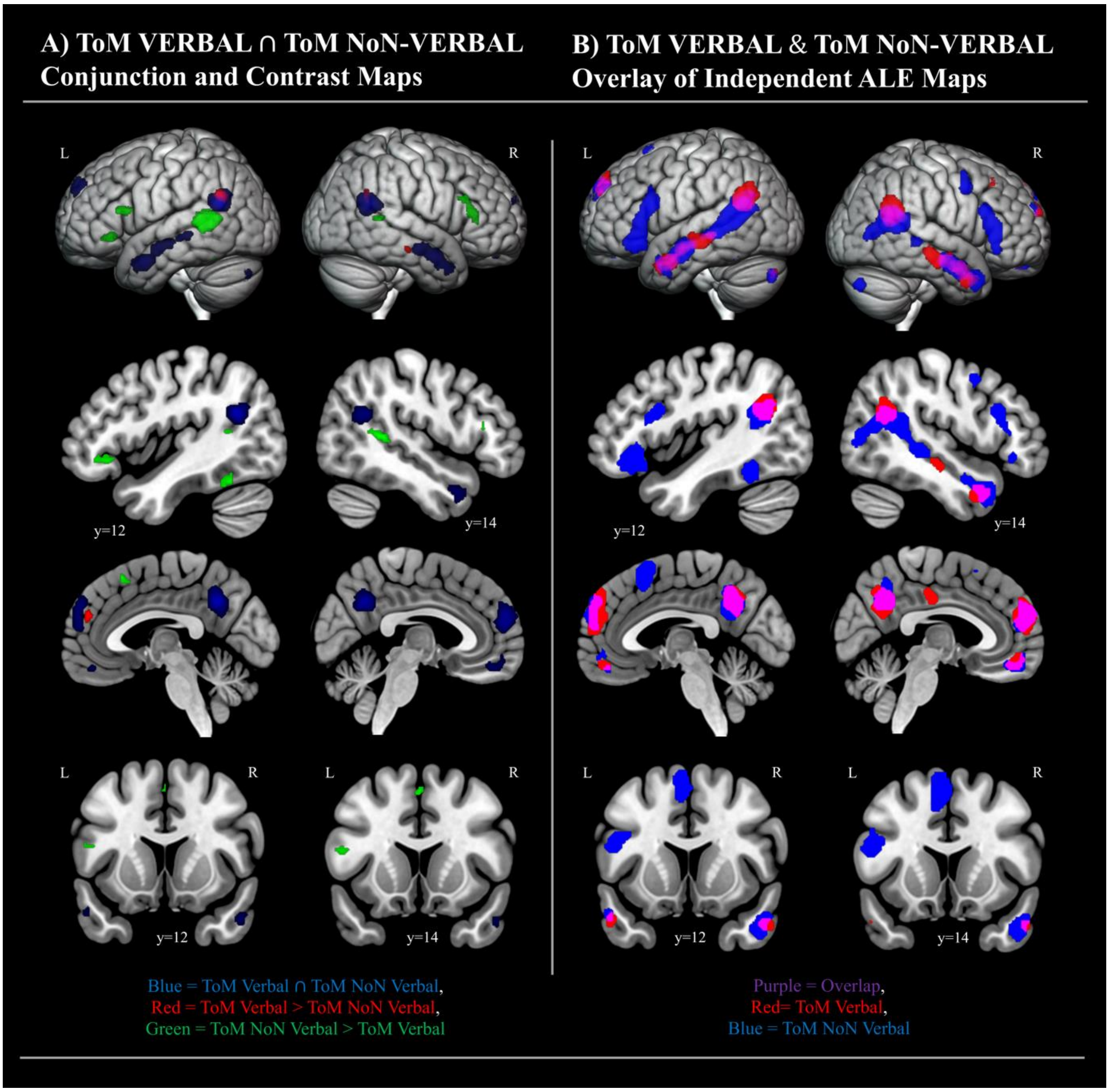

***Supplementary Figure R6*** *Formal and overlay conjunction and contrast maps showing common converging and differential activation for VERBAL ToM (N=46) and NON-VERBAL ToM (N= 71) experiments. The initial ALE maps were treated to a cluster-forming threshold at p<.001 and an FWE-corrected cluster-extent threshold at p<.05 prior to the conjunction and contrast analysis. The contrast maps in* ***Panel A*** *were additionally thresholded with a cluster forming threshold at p<.001 and a minimum cluster size of 100mm*$^{3}$*.* ***Panel A*** *displays the conjunction alongside side statistically significant differences. In* ***Panel B,*** *we have overlaid the binarised versions of the complete ALE maps resulting from independent analysis*

*of VERBAL ToM and NON-VERBAL ToM studies. This allows for full visualisation of the topography of the associated networks. The sagittal and coronal sections are chosen as representative slices positioned over peak coordinates at which there is the greatest conjunction in the bilateral anterior temporal lobes (left y= 12; right y= 14).*

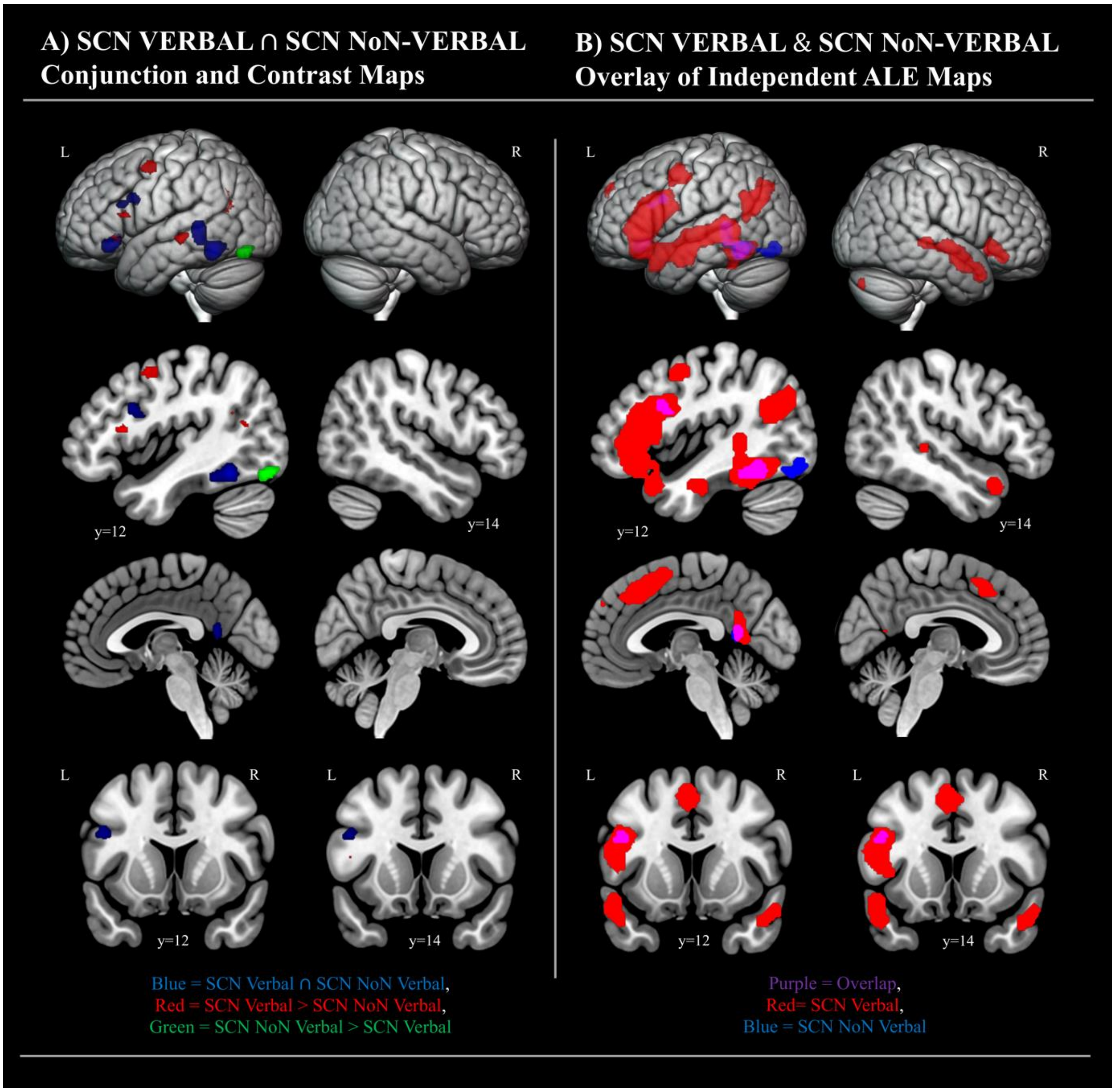

***Supplementary Figure R7*** *Formal and overlay conjunction and contrast maps showing common converging and differential activation for VERBAL SC (N=175) and NON-VERBAL SC (N= 37) experiments. The initial ALE maps were treated to a cluster-forming threshold at p<.001 and an FWE-corrected cluster-extent threshold at p<.05 prior to the conjunction and contrast analysis. The contrast maps in* ***Panel A*** *were additionally thresholded with a cluster-*

*forming threshold at p<.001 and a minimum cluster size of 100mm³.* ***Panel A*** *displays the conjunction alongside side statistically significant differences. In* ***Panel B,*** *we have overlaid the binarised versions of the complete ALE maps resulting from independent analysis of VERBAL SC and NON-VERBAL SC studies. This allows for full visualisation of the topography of the associated networks. The sagittal and coronal sections are chosen as representative slices positioned over peak coordinates at which there is the greatest conjunction in the bilateral anterior temporal lobes (left y= 12; right y= 14)*

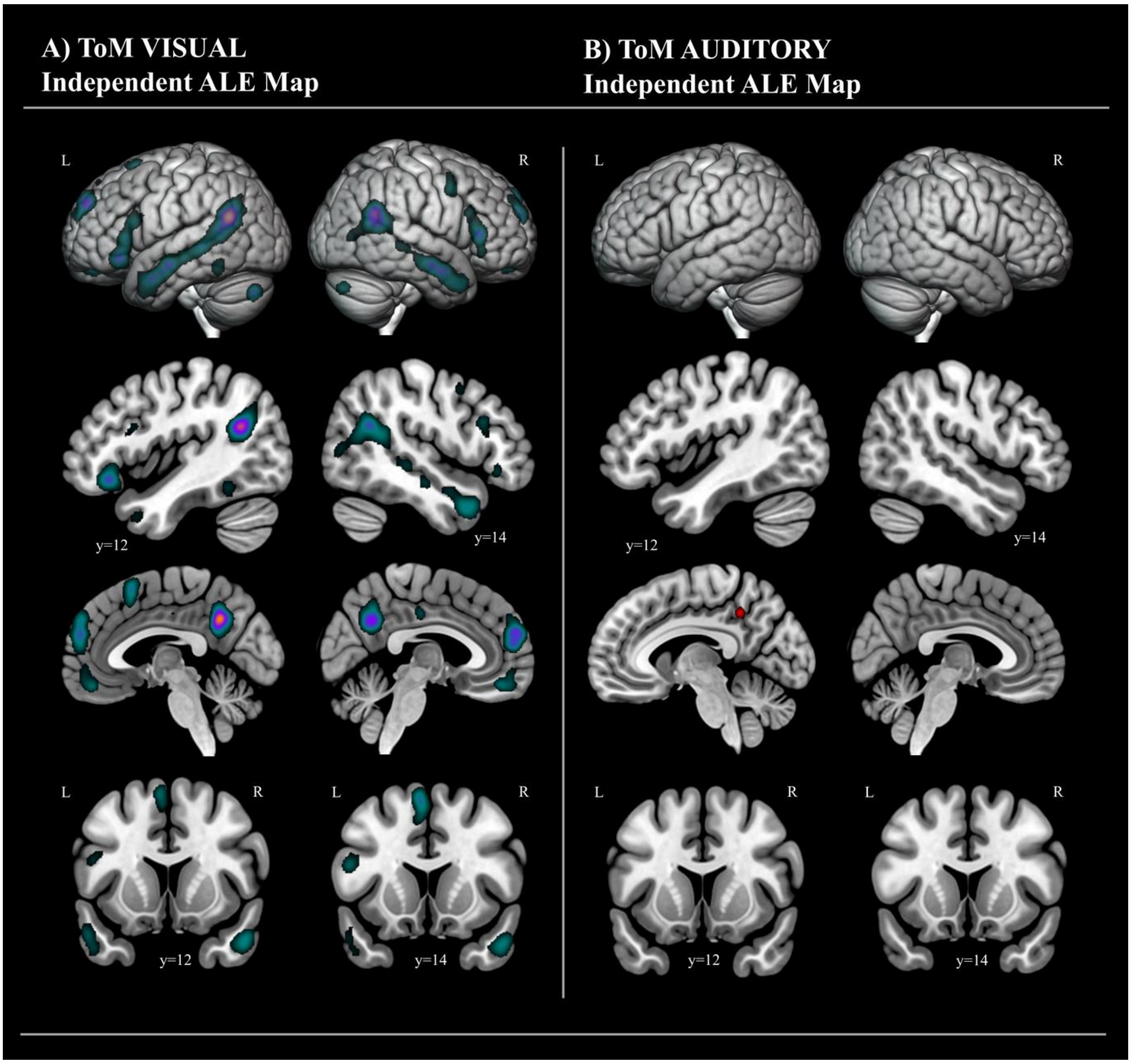

***Supplementary Figure R8*** *Independent ALE activation maps for VISUAL ToM (N= 106) and AUDITORY ToM (N=6); The maps were treated to a cluster forming threshold at p<.001 and*

*an FWE corrected cluster-extent threshold at p<.05. The sagittal and coronal sections are chosen as representative slices positioned over peak coordinates at which there is the greatest conjunction in the bilateral anterior temporal lobes (left y= 12; right y= 14).*

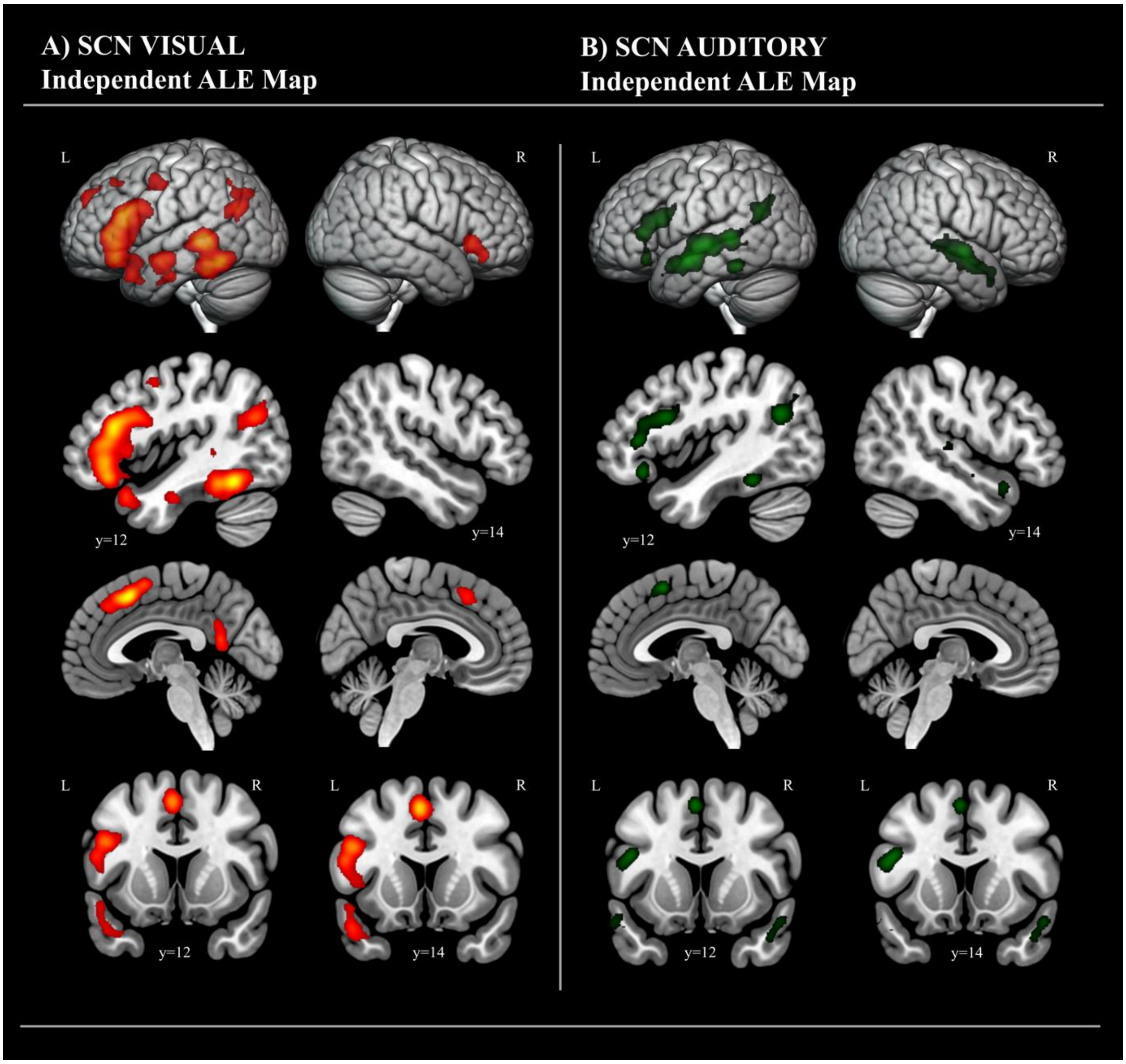

***Supplementary Figure R9*** *Independent ALE activation maps for VISUAL SC (N= 152) and AUDITORY SC (N=60); The maps were treated to a cluster forming threshold at p<.001 and an FWE corrected cluster-extent threshold at p<.05. The sagittal and coronal sections are chosen as representative slices positioned over peak coordinates at which there is the greatest conjunction in the bilateral anterior temporal lobes (left y= 12; right y= 14).*

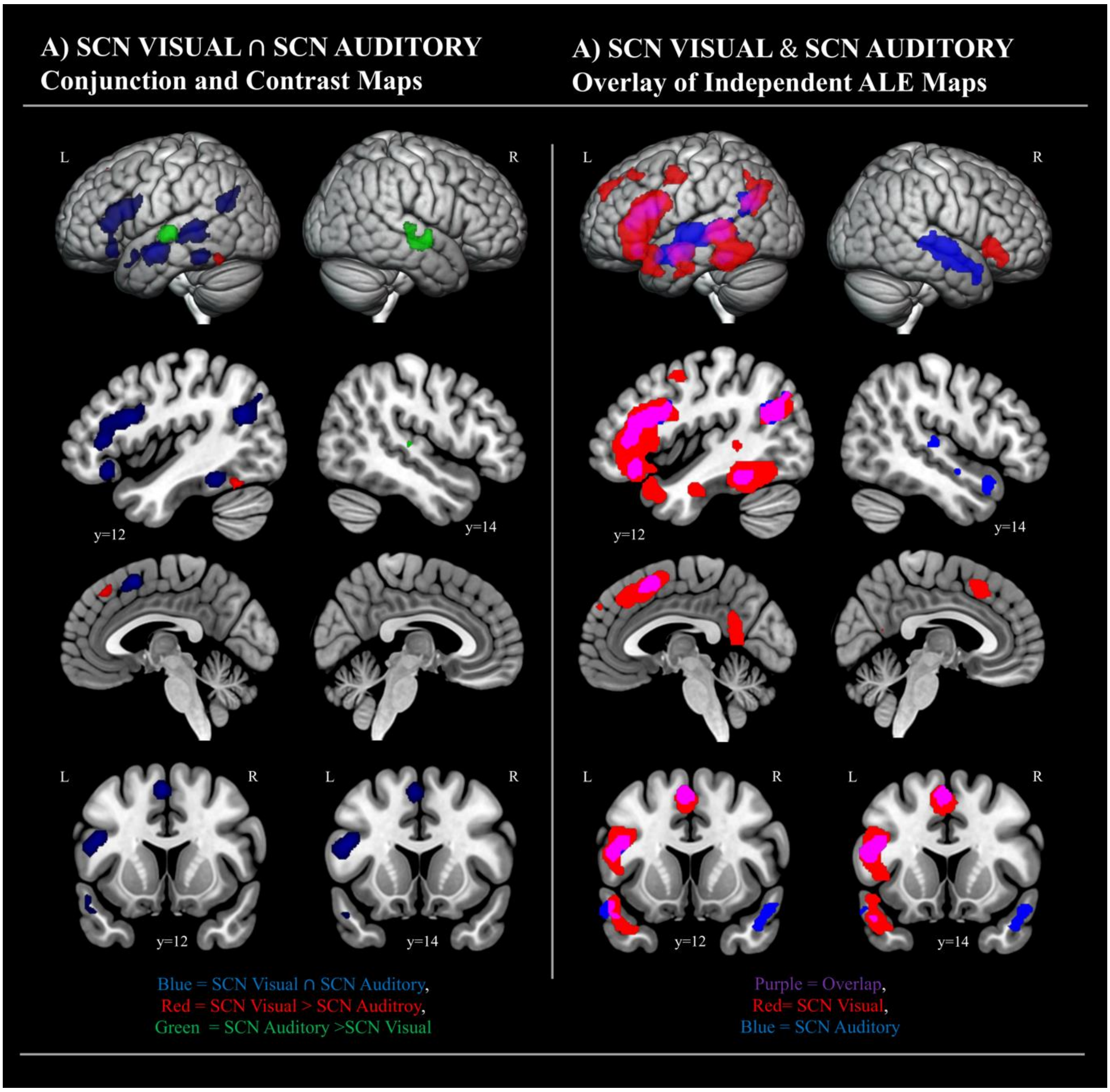

***Supplementary Figure R10*** *Formal and overlay conjunction and contrast maps showing common converging and differential activation for VISUAL SC (N=152) and AUDITORY SC (N= 60) experiments. The initial ALE maps were treated to a cluster-forming threshold at p<.001 and an FWE-corrected cluster-extent threshold at p<.05 prior to the conjunction and contrast analysis. The contrast maps in* ***Panel A*** *were additionally thresholded with a cluster forming threshold at p<.001 and a minimum cluster size of 100mm$^{3}$.* ***Panel A*** *displays the conjunction alongside side statistically significant differences. In* ***Panel B,*** *we have overlaid the binarised versions of the complete ALE maps resulting from independent analysis of VISUAL SC and AUDITORY SC studies. This allows for full visualisation of the topography of*

*the associated networks. The sagittal and coronal sections are chosen as representative slices positioned over peak coordinates at which there is the greatest conjunction in the bilateral anterior temporal lobes (left y= 12; right y= 14)*

#### *Supplementary Tables:*

***Supplementary Table R1:*** *The results of the independent ALE analyses across subsets of the ToM data (cluster forming threshold p<.001; cluster-extent FWE p<.05). Note that all subsequent analyses were conducted on the ToM data containing experiments with high and low baselines (SECTION B).*

| Cluster Size | Region of Activation | Peak MNI Co-ordinates X | Y | Z | ALE Value | Z Value |
|---|---|---|---|---|---|---|
| | | **SECTION A: ToM All Baselines; (N= 114)** | | | | |
| 24400 | Left MTG | -50 | -58 | 22 | 0.15 | 13.04 |
| | Left MTG | -58 | -10 | -14 | 0.08 | 8.37 |
| | Left MTG | -54 | 2 | -26 | 0.08 | 7.71 |
| | Left MTG | -56 | -38 | 0 | 0.07 | 7.42 |
| | Left MTG | -60 | -22 | -8 | 0.07 | 6.98 |
| 11840 | Right MTG | 54 | 0 | -22 | 0.09 | 8.55 |
| | Right MTG | 60 | -8 | -18 | 0.08 | 8.30 |
| | Right MTG | 52 | 6 | -28 | 0.07 | 7.14 |
| | Right MTG | 50 | -30 | -4 | 0.05 | 5.27 |
| 11688 | Left Medial SFG | -10 | 54 | 34 | 0.11 | 10.24 |
| | Right Medial SFG | 4 | 56 | 24 | 0.08 | 7.95 |
| | Left Medial SFG | -4 | 56 | 24 | 0.07 | 7.50 |
| | Left Medial SFG | -8 | 44 | 50 | 0.03 | 3.70 |
| 9992 | Left Precuneus | -2 | -54 | 36 | 0.13 | 11.52 |
| 9560 | Right STG | 54 | -54 | 24 | 0.11 | 10.37 |
| | Right MTG | 50 | -72 | 6 | 0.04 | 4.12 |
| 7880 | Left IFG (pars orbitalis) | -48 | 28 | -10 | 0.09 | 8.65 |
| | Left IFG (pars triangularis) | -52 | 24 | 6 | 0.07 | 7.25 |
| | Left IFG (pars opercularis) | -50 | 18 | 20 | 0.07 | 6.87 |
| 6208 | Right IFG (pars triangularis) | 56 | 28 | 8 | 0.09 | 8.54 |
| | Right IFG (pars triangularis) | 46 | 20 | 24 | 0.05 | 5.27 |
| 4120 | Left Gyrus Rectus | 0 | 46 | -18 | 0.07 | 7.10 |
| | Left Anterior Cingulum | -8 | 50 | 0 | 0.04 | 4.30 |
| 3568 | Left SMA | -6 | 14 | 60 | 0.07 | 7.37 |
| 2248 | Left Cerebellum | -24 | -78 | -36 | 0.08 | 7.86 |
| 1736 | Right Precentral Gyrus | 42 | 6 | 42 | 0.05 | 5.85 |
| 1272 | Left Precentral Gyrus | -42 | 2 | 54 | 0.04 | 4.61 |
| 1176 | Left Fusiform Gyrus | -42 | -50 | -16 | 0.05 | 5.48 |
| 1048 | Right Cerebellum | 28 | -78 | -34 | 0.06 | 6.31 |
| | | **SECTION B: ToM High & Low Baselines (rest excluded); (N= 113)** | | | | |

| | | | | | | |
|---|---|---|---|---|---|---|
| 24328 | Left MTG | -50 | -58 | 22 | 0.15 | 12.88 |
| | Left MTG | -58 | -10 | -14 | 0.08 | 8.44 |
| | Left MTG | -54 | 2 | -26 | 0.08 | 7.78 |
| | Left MTG | -56 | -38 | 0 | 0.07 | 7.43 |
| | Left MTG | -60 | -22 | -10 | 0.06 | 6.62 |
| 11968 | Left Medial SFG | -10 | 54 | 34 | 0.10 | 9.90 |
| | Right Medial SFG | 4 | 56 | 24 | 0.08 | 8.02 |
| | Left Medial SFG | -4 | 56 | 24 | 0.07 | 7.56 |
| | Left Medial SFG | -8 | 44 | 50 | 0.03 | 3.76 |
| 11920 | Right MTG | 54 | 0 | -22 | 0.09 | 8.61 |
| | Right MTG | 60 | -8 | -18 | 0.08 | 8.36 |
| | Right MTG | 52 | 6 | -28 | 0.07 | 7.20 |
| | Right MTG | 52 | -18 | -14 | 0.06 | 6.79 |
| | Right MTG | 50 | -30 | -4 | 0.04 | 4.80 |
| | Right MTG | 52 | -34 | -2 | 0.04 | 4.59 |
| 10112 | Left Precuneus | -2 | -54 | 36 | 0.12 | 11.34 |
| 9648 | Right STG | 54 | -54 | 24 | 0.11 | 10.44 |
| | Right MTG | 50 | -72 | 6 | 0.04 | 4.18 |
| 7616 | Left IFG (pars orbitalis) | -48 | 28 | -10 | 0.09 | 8.60 |
| | Left IFG (pars triangularis) | -52 | 24 | 6 | 0.07 | 7.29 |
| | Left IFG (pars opercularis) | -50 | 18 | 20 | 0.06 | 6.72 |
| 5904 | Right IFG (pars triangularis) | 56 | 28 | 8 | 0.09 | 8.51 |
| | Right IFG (pars triangularis) | 46 | 20 | 24 | 0.05 | 5.31 |
| | Right IFG (pars orbitalis) | 52 | 30 | -6 | 0.05 | 5.30 |
| 4288 | Left Gyrus Rectus | 0 | 46 | -18 | 0.07 | 7.17 |
| | Left Anterior Cingulum | -8 | 50 | 0 | 0.04 | 4.36 |
| 3368 | Left SMA | -6 | 14 | 62 | 0.07 | 7.17 |
| 2304 | Left Cerebellum | -24 | -78 | -36 | 0.08 | 7.92 |
| 1744 | Right Precentral Gyrus | 42 | 6 | 42 | 0.05 | 5.91 |
| 1080 | Right Cerebellum | 28 | -78 | -34 | 0.06 | 6.37 |
| 1048 | Left Precentral Gyrus | -42 | 0 | 54 | 0.04 | 4.53 |
| 976 | Left Fusiform Gyrus | -42 | -50 | -16 | 0.04 | 4.97 |

**SECTION C: ToM High Baselines (low baselines and rest excluded); (N= 111)**

| | | | | | | |
|---|---|---|---|---|---|---|
| 24440 | Left MTG | -50 | -58 | 22 | 0.15 | 12.91 |
| | Left MTG | -58 | -10 | -14 | 0.08 | 8.46 |
| | Left MTG | -54 | 2 | -26 | 0.08 | 7.80 |
| | Left MTG | -56 | -38 | 0 | 0.07 | 7.45 |
| | Left MTG | -60 | -22 | -10 | 0.06 | 6.64 |
| 12008 | Left Medial SFG | -10 | 54 | 34 | 0.10 | 9.93 |
| | Right Medial SFG | 4 | 56 | 24 | 0.08 | 8.04 |
| | Left Medial SFG | -4 | 56 | 24 | 0.07 | 7.58 |
| | Left Medial SFG | -8 | 44 | 50 | 0.03 | 3.78 |
| 11992 | Right MTG | 54 | 0 | -22 | 0.09 | 8.64 |
| | Right MTG | 60 | -8 | -18 | 0.08 | 8.38 |
| | Right MTG | 52 | 6 | -28 | 0.07 | 7.22 |
| | Right MTG | 52 | -18 | -14 | 0.06 | 6.81 |

| | | | | | | |
|---|---|---|---|---|---|---|
| | Right MTG | 50 | -30 | -4 | 0.04 | 4.82 |
| | Right MTG | 52 | -34 | -2 | 0.04 | 4.61 |
| 10080 | Left Precuneus | -2 | -54 | 36 | 0.12 | 11.36 |
| 9720 | Right STG | 54 | -54 | 24 | 0.11 | 10.46 |
| | Right MTG | 50 | -72 | 6 | 0.04 | 4.19 |
| 7448 | Left IFG (pars orbitalis) | -48 | 28 | -10 | 0.09 | 8.63 |
| | Left IFG (pars triangularis) | -52 | 24 | 6 | 0.07 | 7.30 |
| | Left IFG (Pars opercularis) | -50 | 18 | 20 | 0.06 | 6.58 |
| 5840 | Right IFG (pars triangularis) | 56 | 28 | 8 | 0.09 | 8.53 |
| | Right IFG (pars orbitalis) | 52 | 30 | -6 | 0.05 | 5.31 |
| | Right IFG (pars triangularis) | 46 | 20 | 24 | 0.05 | 5.22 |
| 4344 | Left Gyrus Rectus | 0 | 46 | -18 | 0.07 | 7.19 |
| | Left Anterior Cingulum | -8 | 50 | 0 | 0.04 | 4.38 |
| 3200 | Left SMA | -6 | 16 | 62 | 0.06 | 6.76 |
| 2328 | Left Cerebellum | -24 | -78 | -36 | 0.08 | 7.94 |
| 1760 | Right Precentral Gyrus | 42 | 6 | 42 | 0.05 | 5.92 |
| 1064 | Right Cerebellum | 28 | -78 | -34 | 0.06 | 6.38 |
| 984 | Left Fusiform Gyrus | -42 | -50 | -16 | 0.04 | 4.99 |

*Anatomical labels are derived from the Automatic Anatomical Labelling Atlas. MTG = middle temporal gyrus; TP = temporal pole; MOG = middle occipital gyrus; ITG = inferior temporal gyrus; SMA = supplementary motor area; SFG = superior frontal gyrus; STG = superior temporal gyrus; IOG = inferior occipital gyrus; OFG = orbitofrontal gyrus*

***Supplementary Table R2:*** *Significant clusters of converging activation likelihood as given by the independent ALE analyses across subsets of the SC data (cluster forming threshold p<.001; cluster-extent FWE p<.05). Note that all analyses were conducted on the SC data containing experiments with high and low baselines (SECTION B).*

| **Cluster Size** | **Region of Activation** | **Peak MNI Co-ordinates** | | | **ALE Value** | **Z Value** |
|---|---|---|---|---|---|---|
| | | **X** | **Y** | **Z** | | |
| | **SECTION A: SC All Baselines; (N= 214)** | | | | | |
| 87128 | Left MTG | -56 | -38 | 2 | 0.17 | 12.99 |
| | Left IFG (pars triangularis) | -50 | 30 | 6 | 0.14 | 11.19 |
| | Left Fusiform | -30 | -34 | -20 | 0.14 | 10.95 |
| | Left IFG (pars triangularis) | -48 | 24 | 18 | 0.14 | 10.90 |
| | Left Fusiform | -44 | -54 | -14 | 0.13 | 10.22 |
| | Left Fusiform | -40 | -44 | -18 | 0.13 | 10.09 |
| | Left MTG | -56 | -6 | -14 | 0.13 | 10.08 |
| | Left IFG (pars orbitalis) | -40 | 30 | -12 | 0.12 | 9.88 |
| | Left Angular Gyrus | -46 | -66 | 26 | 0.10 | 8.24 |
| | Left Hippocampus | -22 | -8 | -16 | 0.09 | 7.50 |
| | Left MTG | -60 | -24 | -4 | 0.08 | 7.11 |
| | Left Middle TP | -46 | 16 | -26 | 0.08 | 6.76 |
| | Left Insula | -32 | 26 | -2 | 0.08 | 6.55 |
| | Left Precentral Gyrus | -48 | 0 | 50 | 0.07 | 6.34 |
| | Left Superior TP | -52 | 8 | -18 | 0.07 | 5.82 |
| | Left MOG | -34 | -66 | 40 | 0.06 | 5.42 |
| | Left ITG | -42 | -14 | -26 | 0.06 | 5.06 |
| | Left MOG | -38 | -76 | 38 | 0.06 | 4.82 |
| | Left Postcentral Gyrus | -52 | -8 | 44 | 0.05 | 4.42 |
| | Left Angular Gyrus | -54 | -54 | 32 | 0.05 | 3.96 |
| | Left Superior Parietal Lobule | -30 | -60 | 48 | 0.04 | 3.77 |
| 11088 | Left SMA | -4 | 18 | 50 | 0.13 | 10.17 |
| | Left SMA | -4 | 8 | 58 | 0.11 | 8.74 |
| | Left Medial SFG | -8 | 52 | 34 | 0.06 | 5.30 |
| | Left SFG | -16 | 32 | 46 | 0.06 | 5.01 |
| 6736 | Right MTG | 60 | 0 | -14 | 0.07 | 6.05 |
| | Right Superior TP | 52 | 12 | -20 | 0.06 | 5.06 |
| | Right Middle TP | 48 | 16 | -26 | 0.06 | 5.01 |
| | Right MTG | 56 | -28 | 2 | 0.05 | 4.76 |
| | Right STG | 62 | -8 | -4 | 0.05 | 4.66 |
| 4360 | Right Insula | 36 | 24 | -2 | 0.08 | 6.68 |

| | | | | | | |
|---|---|---|---|---|---|---|
| | Right IFG (pars orbitalis) | 36 | 34 | -12 | 0.07 | 6.10 |
| 3288 | Left Precuneus | -6 | -56 | 14 | 0.08 | 6.49 |
| | Left Precuneus | -6 | -56 | 10 | 0.07 | 6.38 |
| | Right Calcarine | 8 | -58 | 12 | 0.04 | 3.11 |
| 1688 | Right IFG (pars opercularis) | 48 | 18 | 28 | 0.06 | 5.44 |
| 1224 | Left IOG | -24 | -92 | -2 | 0.06 | 5.28 |
| 1136 | Right Cerebellum | 22 | -80 | -32 | 0.06 | 5.12 |
| 1136 | Left Putamen | -18 | 4 | 10 | 0.06 | 4.85 |

**SECTION B: SC High & Low Baselines (rest excluded); (N= 211)**

| | | | | | | |
|---|---|---|---|---|---|---|
| 86104 | Left MTG | -56 | -38 | 2 | 0.16 | 12.79 |
| | Left IFG (pars triangularis) | -50 | 30 | 6 | 0.14 | 11.39 |
| | Left Fusiform | -30 | -34 | -20 | 0.14 | 11.18 |
| | Left IFG (pars triangularis) | -48 | 24 | 18 | 0.14 | 11.06 |
| | Left MTG | -56 | -6 | -14 | 0.12 | 1.00 |
| | Left Fusiform | -40 | -42 | -20 | 0.12 | 9.96 |
| | Left Fusiform | -46 | -52 | -16 | 0.12 | 9.58 |
| | Left IFG (pars orbitalis) | -40 | 30 | -12 | 0.11 | 9.52 |
| | Left Angular Gyrus | -46 | -66 | 26 | 0.10 | 8.43 |
| | Left Hippocampus | -22 | -8 | -16 | 0.09 | 7.73 |
| | Left Middle TP | -46 | 16 | -26 | 0.08 | 6.98 |
| | Left MTG | -62 | -20 | -2 | 0.07 | 6.51 |
| | Left MTG | -60 | -24 | -4 | 0.07 | 6.51 |
| | Left Insula | -34 | 26 | -2 | 0.07 | 6.26 |
| | Left Precentral Gyrus | -48 | 0 | 50 | 0.07 | 6.22 |
| | Left Superior TP | -52 | 8 | -18 | 0.06 | 5.74 |
| | Left MOG | -34 | -66 | 40 | 0.06 | 5.65 |
| | Left ITG | -42 | -14 | -26 | 0.06 | 5.27 |
| | Left MOG | -38 | -76 | 38 | 0.06 | 5.04 |
| | Left Postcentral Gyrus | -52 | -8 | 44 | 0.05 | 4.56 |
| | Left ITG | -34 | -4 | -34 | 0.04 | 3.60 |
| 11128 | Left SMA | -4 | 18 | 50 | 0.13 | 10.33 |
| | Left Medial SFG | -8 | 52 | 34 | 0.06 | 5.36 |
| | Left SFG | -16 | 32 | 46 | 0.06 | 5.24 |
| 5616 | Right MTG | 58 | 0 | -16 | 0.07 | 6.15 |
| | Right Middle TP | 48 | 16 | -26 | 0.06 | 5.20 |
| | Right STG | 62 | -10 | -2 | 0.05 | 4.77 |
| | Right Middle TP | 54 | 10 | -18 | 0.05 | 4.56 |
| 3816 | Right Insula | 36 | 24 | -2 | 0.07 | 6.36 |
| | Right IFG (pars orbitalis) | 36 | 34 | -12 | 0.07 | 6.02 |
| 3600 | Left Precuneus | -6 | -56 | 14 | 0.08 | 6.72 |
| | Left Precuneus | -6 | -56 | 10 | 0.07 | 6.59 |
| 1960 | Right Para Hippocampal Gyrus | 26 | -16 | -22 | 0.06 | 5.32 |
| | Right Fusiform | 34 | -36 | -20 | 0.05 | 4.26 |
| | Right Hippocampus | 26 | -6 | -18 | 0.04 | 3.31 |
| 1272 | Right Cerebellum | 22 | -80 | -32 | 0.06 | 5.31 |
| 1168 | Right IFG (pars opercularis) | 46 | 20 | 28 | 0.05 | 4.74 |

**SECTION C: SC High Baselines (low baselines and rest excluded); (N= 170)**

| | | | | | | |
|---|---|---|---|---|---|---|
| 31376 | Left MTG | -56 | -40 | 0 | 0.13 | 11.29 |
| | Left Fusiform | -30 | -34 | -18 | 0.12 | 11.00 |
| | Left MTG | -56 | -6 | -16 | 0.08 | 8.02 |
| | Left Amygdala | -22 | -6 | -16 | 0.07 | 6.86 |
| | Left ITG | -48 | -52 | -16 | 0.06 | 6.48 |
| | Left MTG | -62 | -22 | -6 | 0.05 | 4.88 |
| | Left ITG | -42 | -16 | -28 | 0.03 | 3.53 |
| | Left Fusiform | -36 | -16 | -24 | 0.03 | 3.40 |
| 25088 | Left IFG (pars triangularis) | -50 | 28 | 6 | 0.10 | 9.42 |
| | Left IFG (pars triangularis) | -50 | 24 | 14 | 0.10 | 9.41 |
| | Left IFG (pars orbitalis) | -42 | 28 | -14 | 0.09 | 8.42 |
| | Left Middle TP | -46 | 16 | -26 | 0.05 | 5.73 |
| | Left Insula | -34 | 24 | -4 | 0.05 | 5.25 |
| | Left Insula | -42 | 18 | 4 | 0.04 | 4.43 |
| 10160 | Left SMA | -4 | 18 | 50 | 0.09 | 8.43 |
| | Left SFG | -16 | 32 | 46 | 0.05 | 5.38 |
| | Left Medial SFG | -8 | 52 | 34 | 0.05 | 5.36 |
| | Left MFG | -24 | 26 | 46 | 0.04 | 3.83 |
| 9632 | Left Angular Gyrus | -46 | -66 | 26 | 0.09 | 8.90 |
| | Left MOG | -38 | -76 | 38 | 0.05 | 5.38 |
| | Left Angular Gyrus | -36 | -64 | 38 | 0.04 | 4.36 |
| 4768 | Left Precuneus | -6 | -56 | 14 | 0.07 | 7.31 |
| | Left Precuneus | -4 | -66 | 30 | 0.03 | 3.64 |
| 2752 | Right MTG | 58 | 0 | -16 | 0.06 | 6.34 |
| 2384 | Right IFG (pars orbitalis) | 36 | 34 | -12 | 0.05 | 5.71 |
| | Right Insula | 34 | 22 | -2 | 0.05 | 5.12 |
| 2032 | Right Para Hippocampal Gyrus | 26 | -16 | -22 | 0.05 | 5.63 |
| | Right Fusiform | 34 | -32 | -20 | 0.04 | 3.93 |
| | Right Hippocampus | 26 | -6 | -18 | 0.03 | 3.79 |
| 1736 | Right Cerebellum | 22 | -80 | -32 | 0.06 | 6.02 |
| 1712 | Left Medial OFG | -2 | 54 | -12 | 0.06 | 5.83 |

*Anatomical labels are derived from the Automatic Anatomical Labelling Atlas. MTG = middle temporal gyrus; TP = temporal pole; MOG = middle occipital gyrus; ITG = inferior temporal gyrus; SMA = supplementary motor area; SFG = superior frontal gyrus; STG = superior temporal gyrus; IOG = inferior occipital gyrus; OFG = orbitofrontal gyrus*

***Supplementary Table R3*** *Formal conjunction and contrast analyses of the ToM (N= 113) and NON-SOCIAL SC (N= 193) experiments after excluding experiments with a degree of social content from the SC data; (cluster forming threshold p<.001; FWE cluster-extent corrected at p<.05). The contrast analyses were further thresholded with a cluster forming threshold at p<a .001 and minimum cluster size of 100mm3.*

| **Cluster Size** | **Region of Activation** | **Peak MNI Co-ordinates** | | | **ALE Value** | **Z Value** |
|---|---|---|---|---|---|---|
| | | **X** | **Y** | **Z** | | |
| | **ToM ∩ SC NON-SOCIAL CONJUNCTION** | | | | | |
| 9968 | Left MTG | -58 | -10 | -14 | 0.08 | |
| | Left MTG | -56 | -38 | 0 | 0.07 | |
| | Left MTG | -58 | -24 | -6 | 0.05 | |
| | Left MTG | -54 | 6 | -22 | 0.05 | |
| 7248 | Left IFG (pars orbitalis) | -48 | 28 | -10 | 0.09 | |
| | Left IFG (pars triangularis) | -52 | 24 | 6 | 0.07 | |
| | Left IFG (pars opercularis) | -50 | 18 | 20 | 0.06 | |
| 4600 | Left Angular Gyrus | -46 | -62 | 26 | 0.08 | |
| 2096 | Left SMA | -4 | 14 | 58 | 0.06 | |
| 1816 | Right MTG | 56 | 0 | -18 | 0.06 | |
| | Right Middle TP | 48 | 14 | -26 | 0.05 | |
| 1192 | Left Medial SFG | -8 | 52 | 34 | 0.06 | |
| 976 | Left Fusiform | -42 | -50 | -16 | 0.04 | |
| 424 | Left Precuneus | -6 | -54 | 24 | 0.04 | |
| 416 | Left Precentral | -42 | 2 | 52 | 0.04 | |
| 400 | Right Cerebellum | 26 | -80 | -34 | 0.05 | |
| 384 | Right MTG | 50 | -32 | -2 | 0.04 | |
| 8 | Left Angular Gyrus | -46 | -70 | 38 | 0.03 | |
| | **ToM > SC NON-SOCIAL CONTRAST** | | | | | |
| 8480 | Left Medial SFG | 1.2 | 56.2 | 22.3 | | 3.89 |
| | Right Medial SFG | 12 | 62 | 26 | | 3.72 |
| 7816 | Right Precuneus | 1 | -55 | 35.9 | | 3.89 |
| 7296 | Right MTG | 55.2 | -54 | 19.3 | | 3.89 |
| | Right MTG | 48 | -42 | 8 | | 3.54 |
| 4968 | Right MTG | 54.3 | -0.6 | -26 | | 3.89 |
| | Right MTG | 52 | -22 | -13 | | 3.54 |
| 4832 | Left MTG | -53 | -55 | 20.9 | | 3.89 |
| 2256 | Right IFG (pars triangularis) | 55.9 | 25.8 | 7 | | 3.89 |

| | | | | | |
|---|---|---|---|---|---|
| | Right IFG (pars orbitalis) | 53 | 28 | -4 | 3.72 |
| 2192 | Left Cerebellum | -24 | -78 | -36 | 3.89 |
| 1616 | Right Precentral Gyrus | 42.3 | 6 | 45.3 | 3.89 |
| 1544 | Left ITG | -51 | 5 | -34 | 3.89 |
| 1208 | Right Gyrus Rectus | 3.9 | 47.7 | -18 | 3.89 |
| 424 | Left SMA | -9.3 | 18.7 | 64.7 | 3.89 |
| | Left SMA | -6 | 14 | 64 | 3.72 |
| 160 | Left Anterior Cingulum | -10 | 54 | 2 | 3.89 |
| 136 | Right IFG (pars triangularis) | 37 | 18.3 | 23.7 | 3.72 |
| | **SC NON-SOCIAL > ToM CONTRAST** | | | | |
| 6936 | Left IFG (pars triangularis) | -42 | 31.9 | 6.5 | 3.89 |
| 5704 | Left Fusiform | -41 | -42 | -19 | 0.00 |
| | Left ITG | -54 | -42 | -13 | 3.54 |
| | Left PHG | -25 | -24 | -21 | 3.09 |
| 872 | Left Calcarine | -3.8 | -57 | 6.6 | 3.89 |
| 552 | Left Fusiform | -38 | -18 | -25 | 3.89 |
| | Left Fusiform | -36 | -14 | -28 | 3.72 |
| | Left ITG | -40 | -12 | -30 | 3.54 |
| 544 | Right STG | 62.3 | -8.6 | 0.4 | 3.89 |
| 424 | Right Middle Cingulum | 3.3 | 23.3 | 40.7 | 3.54 |
| | Right Middle Cingulum | 7.3 | 20 | 42 | 3.54 |
| 400 | Right IFG (pars orbitalis) | 33.3 | 36 | -6.7 | 3.89 |
| | Right IFG (pars orbitalis) | 36 | 32 | -8 | 3.72 |
| 272 | Left STG | -63 | -25 | 5.3 | 3.89 |
| 184 | Left MTG | -44 | -42 | 0 | 3.54 |
| 136 | Left SMA | -6 | 2 | 52 | 3.43 |
| 112 | Left MOG | -34 | -66 | 34 | 3.16 |

*Anatomical labels are derived from the Automatic Anatomical Labelling Atlas. MTG = middle temporal gyrus, IFG - inferior frontal gyrus, AG = angular gyrus, TP = temporal pole, MOG = middle occipital gyrus, ITG = inferior temporal gyrus, SMA = supplementary motor area, SFG = superior frontal gyrus, STG = superior temporal gyrus, PHG= parahippocampalpal gyrus.*

***Supplementary Table R4** Formal conjunction and contrast analyses of the VERBAL ToM (N= 46) and VERBAL SC (N= 175) experiments after excluding experiments with NON-VERBAL stimuli; (cluster forming threshold p<.001; FWE cluster-extent corrected at* *p<.05). The contrast analyses were further thresholded with a cluster forming threshold at p<.001 and minimum cluster size of 100mm3*

| Cluster Size | Region of Activation | Peak MNI Co-ordinates X | Y | Z | ALE Value | Z Value |
|---|---|---|---|---|---|---|
| | **VERBAL ToM ∩ VERBAL SC CONJUNCTION** | | | | | |
| 4032 | Left MTG | -60 | -24 | -10 | 0.04 | |
| | Left MTG | -54 | 2 | -24 | 0.04 | |
| | Left MTG | -60 | -10 | -14 | 0.03 | |
| 3632 | Left AG | -48 | -62 | 26 | 0.07 | |
| 1072 | Right MTG | 54 | 0 | -20 | 0.03 | |
| | Right MTG | 60 | -8 | -14 | 0.02 | |
| 1032 | Left Medial SFG | -8 | 54 | 32 | 0.04 | |
| 168 | Left Precuneus | -2 | -56 | 28 | 0.03 | |
| 88 | Right Middle TP | 48 | 12 | -30 | 0.02 | |
| 24 | Right MTG | 56 | -24 | -4 | 0.02 | |
| 8 | Right Middle TP | 44 | 10 | -30 | 0.02 | |
| 8 | Right Middle TP | 46 | 10 | -28 | 0.02 | |
| 8 | Right Middle TP | 48 | 10 | -26 | 0.02 | |
| 8 | Right MTG | 64 | -10 | -16 | 0.02 | |
| | **VERBAL ToM > VERBAL SC CONTRAST** | | | | | |
| 4552 | Right Medial SFG | 1.1 | 55.3 | 22.2 | | 3.72 |
| 4296 | Right Precuneus | 1.8 | -55 | 35.3 | | 3.72 |
| 3960 | Right STG | 55.6 | -53 | 24.2 | | 3.72 |
| 2664 | Left AG | -53 | -58 | 25 | | 3.72 |
| 1048 | Left Cerebellum | -24 | -78 | -36 | | 3.72 |
| 824 | Right Middle TP | 50 | 5 | -33 | | 3.72 |
| 648 | Right MTG | 54.8 | -22 | -12 | | 3.72 |
| | Right MTG | 60 | -18 | -18 | | 3.54 |
| 536 | Right Middle Cingulum | 2 | -19 | 36 | | 3.72 |
| 512 | Right MFG | 24.3 | 27 | 43.5 | | 3.54 |
| 440 | Right Gyrus Rectus | 4.6 | 48.9 | -17 | | 3.54 |
| | Right Gyrus Rectus | 4 | 52 | -20 | | 3.54 |
| 352 | Left MTG | -53 | 6.3 | -30 | | 3.72 |
| 136 | Left Medial SFG | -1.3 | 50 | 44.7 | | 3.35 |
| | Left Medial SFG | -6 | 50 | 46 | | 3.54 |
| | **VERBAL SC > VERBAL ToM CONTRAST** | | | | | |

| | | | | | |
|---|---|---|---|---|---|
| 10920 | Left IFG (pars triangularis) | -40 | 29.3 | 2.9 | 3.72 |
| | Left IFG (pars opercularis) | -52 | 8 | 20 | 3.54 |
| 5920 | Left Fusiform | -40 | -46 | -21 | 3.72 |
| | Left ITG | -52 | -55 | -20 | 3.54 |
| 5136 | Left MTG | -55 | -43 | 3.7 | 3.72 |
| 912 | Right Insula | 32.7 | 22 | 4 | 3.35 |
| | Right Insula | 32.5 | 28.8 | -1.2 | 3.16 |
| | Right IFG (pars orbitalis) | 30 | 26 | -4 | 3.16 |
| 448 | Left Hippocampus | -25 | -14 | -9 | 3.54 |
| | Left Hippocampus | -20 | -14 | -12 | 3.35 |
| 416 | Left SMA | -0.8 | 16.8 | 52.4 | 3.35 |
| | Left SMA | -2.5 | 9.5 | 49.5 | 3.35 |
| 408 | Left ITG | -41 | -17 | -30 | 3.72 |
| | Left ITG | -39 | -8 | -30 | 3.09 |
| 192 | Left Medial SFG | -4 | 32 | 46 | 3.72 |
| | Left Medial SFG | -8 | 30 | 45 | 3.54 |

*Anatomical labels are derived from the Automatic Anatomical Labelling Atlas. MTG = middle temporal gyrus, IFG - inferior frontal gyrus, AG = angular gyrus, TP = temporal pole, MOG = middle occipital gyrus, ITG = inferior temporal gyrus, SMA = supplementary motor area, SFG = superior frontal gyrus, STG = superior temporal gyrus, PHG = para hippocampal gyrus, OFG = orbitofrontal gyrus.*

***Supplementary Table R5*** *Formal conjunction and contrast analyses of the NON-VERBAL ToM (N= 71) and NON-VERBAL SC (N= 37) experiments after excluding experiments with VERBAL stimuli; (cluster forming threshold p<.001; FWE cluster-extent corrected at p<.05). The contrast analyses were further thresholded with a cluster forming threshold at p<.001 and minimum cluster size of 100mm3*

| Cluster Size | Region of Activation | Peak MNI Co-ordinates X | Y | Z | ALE Value | Z Value |
|---|---|---|---|---|---|---|
| | **NON-VERBAL ToM ∩ NON-VERBAL SC CONJUNCTION** | | | | | |
| 1208 | Left MTG | -56 | -40 | 2 | 0.03 | |
| 880 | Left Fusiform | -44 | -52 | -16 | 0.03 | |
| | Left Fusiform | -36 | -50 | -16 | 0.02 | |
| | Left Fusiform | -40 | -42 | -18 | 0.02 | |
| 632 | Left IFG (pars opercularis) | -46 | 12 | 26 | 0.02 | |
| | Left IFG (pars triangularis) | -54 | 20 | 22 | 0.02 | |
| 144 | Left IFG (pars orbitalis) | -40 | 32 | -12 | 0.02 | |
| | **NON-VERBAL ToM > NON-VERBAL SC CONTRAST** | | | | | |
| 4504 | Left MTG | -50.5 | -54.7 | 18.3 | | 3.72 |
| 3896 | Right MTG | 53.6 | -50.2 | 16.4 | | 3.72 |
| | Right STG | 51 | -54.2 | 24.4 | | 3.54 |
| | Right MTG | 62 | -56 | 10 | | 3.35 |
| 1568 | Right Precuneus | 7.1 | -53 | 40 | | 3.54 |
| | Right Precuneus | 11 | -54.7 | 41.7 | | 3.54 |
| | Left Precuneus | -6 | -50 | 38 | | 3.35 |
| | Left Precuneus | -7 | -57 | 38 | | 3.09 |
| 1040 | Right MTG | 56.4 | -14.8 | -14.7 | | 3.72 |
| 1008 | Left Cerebellum | -20.5 | -79.2 | -35.2 | | 3.72 |
| | Left Cerebellum | -24 | -74 | -36 | | 3.54 |
| 648 | Left Medial SFG | -12 | 60 | 32 | | 3.72 |
| | Left Medial SFG | -8.9 | 58.6 | 35.1 | | 3.54 |
| 632 | Right Medial SFG | 8.6 | 55.6 | 19.3 | | 3.72 |
| 616 | Left IFG (pars orbitalis) | -50.5 | 24.4 | -6.2 | | 3.72 |
| 216 | Right IFG (pars triangularis) | 52.7 | 24.7 | 0.7 | | 0.00 |
| | Right IFG (pars orbitalis) | 52.7 | 26 | -4.7 | | 3.54 |
| | **NON-VERBAL SC > NON-VERBAL ToM CONTRAST** | | | | | |

| | | | | | |
|---|---|---|---|---|---|
| 896 | Left ITG | -53.2 | -49.7 | -12.9 | 3.72 |
| 240 | Left IOG | -44.7 | -82.7 | -12 | 3.54 |
| | Left Fusiform | -44 | -80.7 | -14.7 | 3.35 |

*Anatomical labels are derived from the Automatic Anatomical Labelling Atlas. MTG = middle temporal gyrus, IFG - inferior frontal gyrus, AG = angular gyrus, TP = temporal pole, MOG = middle occipital gyrus, ITG = inferior temporal gyrus, SMA = supplementary motor area, SFG = superior frontal gyrus, STG = superior temporal gyrus, PHG = para hippocampal gyrus, OFG = orbitofrontal gyrus.*

***Supplementary Table R6*** *The results of the independent ALE analyses for the VERBAL ToM (N= 46) and NON-VERBAL ToM (N= 71) experiments (cluster forming threshold p<.001; cluster-extent FWE p<.05).*

| **Cluster Size** | **Region of Activation** | **Peak MNI Coordinates** | | | **ALE Value** | **Z Value** |
|---|---|---|---|---|---|---|
| | | **X** | **Y** | **Z** | | |
| | | **VERBAL ToM** | | | | |
| 8728 | Left Medial SFG | 0 | 54 | 24 | 0.05 | 7.58 |
| | Left Medial SFG | -8 | 54 | 32 | 0.04 | 6.38 |
| 6616 | Left Precuneus | 0 | -54 | 36 | 0.06 | 8.59 |
| 6512 | Right ITG | 52 | 6 | -32 | 0.04 | 5.88 |
| | Right MTG | 56 | -18 | -14 | 0.04 | 5.83 |
| | Right MTG | 54 | 0 | -20 | 0.03 | 5.41 |
| | Right MTG | 60 | -10 | -14 | 0.02 | 4.22 |
| 5784 | Left MTG | -50 | -58 | 24 | 0.10 | 11.59 |
| 5472 | Left MTG | -54 | 2 | -26 | 0.04 | 6.00 |
| | Left MTG | -60 | -24 | -10 | 0.04 | 5.97 |
| | Left MTG | -60 | -10 | -14 | 0.03 | 4.75 |
| 5104 | Right STG | 54 | -54 | 24 | 0.08 | 9.93 |
| 1688 | Right Gyrus Rectus | 2 | 48 | -16 | 0.03 | 5.57 |
| 1104 | Left Cerebellum | -24 | -78 | -36 | 0.03 | 5.21 |
| 1000 | Right Middle Cingulum | 2 | -18 | 38 | 0.03 | 5.27 |
| 792 | Right MFG | 24 | 26 | 44 | 0.03 | 5.38 |
| | | **NON-VERBAL ToM** | | | | |
| 8728 | Left Medial SFG | 0 | 54 | 24 | 0.05 | 7.58 |
| | Left Medial SFG | -8 | 54 | 32 | 0.04 | 6.38 |
| 6616 | Left Precuneus | 0 | -54 | 36 | 0.06 | 8.59 |
| 6512 | Right ITG | 52 | 6 | -32 | 0.04 | 5.88 |
| | Right MTG | 56 | -18 | -14 | 0.04 | 5.83 |
| | Right MTG | 54 | 0 | -20 | 0.03 | 5.41 |
| | Right MTG | 60 | -10 | -14 | 0.02 | 4.22 |
| 5784 | Left MTG | -50 | -58 | 24 | 0.10 | 11.59 |
| 5472 | Left MTG | -54 | 2 | -26 | 0.04 | 6.00 |
| | Left MTG | -60 | -24 | -10 | 0.04 | 5.97 |
| | Left MTG | -60 | -10 | -14 | 0.03 | 4.75 |
| 5104 | Right STG | 54 | -54 | 24 | 0.08 | 9.93 |
| 1688 | Right Gyrus Rectus | 2 | 48 | -16 | 0.03 | 5.57 |

| | | | | | | |
|---|---|---|---|---|---|---|
| 1104 | Left Cerebellum | -24 | -78 | -36 | 0.03 | 5.21 |
| 1000 | Right Middle Cingulum | 2 | -18 | 38 | 0.03 | 5.27 |
| 792 | Right MFG | 24 | 26 | 44 | 0.03 | 5.38 |

*Anatomical labels are derived from the Automatic Anatomical Labelling Atlas. MTG = middle temporal gyrus, IFG - inferior frontal gyrus, AG = angular gyrus, TP = temporal pole, MOG = middle occipital gyrus, ITG = inferior temporal gyrus, SMA = supplementary motor area, SFG = superior frontal gyrus, STG = superior temporal gyrus, PHG = para hippocampal gyrus, OFG = orbitofrontal gyrus.*

***Supplementary Table R7*** *The results of the independent ALE analyses for the VERBAL SC (N= 175) and NON-VERBAL SC (N= 37) experiments (cluster forming threshold p<.001; cluster-extent FWE p<.05).*

| **Cluster Size** | **Region of Activation** | **Peak MNI Co-ordinates X** | **Y** | **Z** | **ALE Value** | **Z Value** |
|---|---|---|---|---|---|---|
| | | **VERBAL SC** | | | | |
| 68976 | Left MTG | -56 | -38 | 2 | 0.14 | 11.58 |
| | Left IFG (pars triangularis) | -48 | 24 | 16 | 0.12 | 10.73 |
| | Left IFG (pars triangularis) | -50 | 30 | 6 | 0.12 | 10.67 |
| | Left MTG | -56 | -6 | -14 | 0.11 | 9.75 |
| | Left Fusiform Gyrus | -30 | -36 | -18 | 0.10 | 9.22 |
| | Left Fusiform Gyrus | -42 | -44 | -18 | 0.09 | 8.74 |
| | Left IFG (pars orbitalis) | -42 | 30 | -12 | 0.09 | 8.56 |
| | Left Fusiform Gyrus | -46 | -56 | -14 | 0.08 | 7.88 |
| | Left Hippocampus | -24 | -10 | -16 | 0.08 | 7.48 |
| | Left MTG | -60 | -24 | -4 | 0.07 | 7.03 |
| | Left Superior TP | -48 | 16 | -24 | 0.07 | 6.96 |
| | Left ITG | -42 | -14 | -26 | 0.06 | 5.73 |
| | Left Superior TP | -50 | 10 | -16 | 0.06 | 5.59 |
| | Left Insula | -32 | 26 | -2 | 0.06 | 5.58 |
| | Left PHG | -24 | -24 | -18 | 0.05 | 4.83 |
| | Left Insula | -38 | 20 | 2 | 0.05 | 4.63 |
| | Left Precentral Gyrus | -54 | 2 | 20 | 0.04 | 4.06 |
| 9152 | Left Angular Gyrus | -46 | -64 | 26 | 0.09 | 8.05 |
| | Left MOG | -34 | -64 | 38 | 0.05 | 5.35 |
| | Left MOG | -38 | -76 | 38 | 0.04 | 4.56 |
| | Left SPL | -30 | -60 | 46 | 0.04 | 3.92 |
| 8712 | Left SMA | -4 | 18 | 50 | 0.10 | 8.88 |
| | Left SMA | -4 | 6 | 58 | 0.07 | 6.71 |
| | Right Middle Cingulum | 4 | 20 | 44 | 0.05 | 5.48 |
| | Left SFG | -16 | 30 | 46 | 0.04 | 4.42 |
| 6912 | Right MTG | 56 | 0 | -18 | 0.06 | 5.55 |
| | Right STG | 62 | -10 | -2 | 0.05 | 5.28 |
| | Right Middle TP | 46 | 14 | -26 | 0.05 | 5.08 |
| | Right MTG | 54 | -30 | 0 | 0.05 | 4.87 |
| | Right MTG | 66 | -28 | -2 | 0.03 | 3.31 |
| 3488 | Right Insula | 36 | 24 | -2 | 0.06 | 5.82 |
| | Right IFG (pars orbitalis) | 36 | 34 | -10 | 0.05 | 4.64 |
| 2936 | Left Precentral Gyrus | -48 | 0 | 50 | 0.07 | 6.63 |

| | | | | | | |
|---|---|---|---|---|---|---|
| | Left Postcentral Gyrus | -52 | -8 | 44 | 0.05 | 5.07 |
| 2712 | Left Precuneus | -6 | -56 | 16 | 0.05 | 5.43 |
| | Right Calcarine | 6 | -58 | 12 | 0.03 | 3.35 |
| 1128 | Right Cerebellum | 22 | -80 | -34 | 0.05 | 5.49 |
| 1064 | Left Medial SFG | -8 | 54 | 34 | 0.05 | 5.03 |
| | **NON-VERBAL SC** | | | | | |
| 8264 | Left ITG | -48 | -52 | -16 | 0.04 | 6.29 |
| | Left Fusiform Gyrus | -30 | -34 | -20 | 0.03 | 5.69 |
| | Left MTG | -56 | -40 | 2 | 0.03 | 5.03 |
| | Left Fusiform Gyrus | -42 | -80 | -14 | 0.03 | 4.83 |
| | Left Fusiform Gyrus | -36 | -50 | -16 | 0.02 | 4.47 |
| | Left Cerebellum | -42 | -74 | -18 | 0.02 | 4.20 |
| | Left Fusiform Gyrus | -40 | -42 | -18 | 0.02 | 4.14 |
| 1480 | Left IFG (pars orbitalis) | -36 | 32 | -10 | 0.03 | 4.97 |
| 1072 | Left Precuneus | -8 | -56 | 14 | 0.02 | 4.44 |
| | Left Precuneus | -6 | -54 | 10 | 0.02 | 4.36 |
| 888 | Left IFG (pars opercularis) | -46 | 12 | 26 | 0.02 | 4.09 |
| | Left IFG (pars triangularis) | -54 | 20 | 22 | 0.02 | 3.81 |

*Anatomical labels are derived from the Automatic Anatomical Labelling Atlas. MTG = middle temporal gyrus, IFG - inferior frontal gyrus, AG = angular gyrus, TP = temporal pole, MOG = middle occipital gyrus, ITG = inferior temporal gyrus, SMA = supplementary motor area, SFG = superior frontal gyrus, STG = superior temporal gyrus, PHG = para hippocampal gyrus, OFG = orbitofrontal gyrus.*

***Supplementary Table R8*** *Formal conjunction and contrast analyses comparing the within domain differences of the VERBAL ToM (N= 46) and NON-VERBAL ToM (N= 71) experiments (cluster forming threshold p<.001; FWE cluster-extent corrected at p<.05). The contrast analyses were further thresholded with a cluster forming threshold at p<.001 and minimum cluster size of 100mm3*

| Cluster Size | Region of Activation | Peak MNI Co-ordinates | | | ALE Value | Z Value |
|---|---|---|---|---|---|---|
| | | X | Y | Z | | |
| | | **VERBAL ToM ∩ NON-VERBAL ToM CONJUNCTION** | | | | |
| 5632 | Left Medial SFG | -8 | 54 | 32 | 0.04 | |
| | Right Medial SFG | 4 | 56 | 24 | 0.04 | |
| | Right Medial SFG | 4 | 56 | 18 | 0.04 | |
| | Left Medial SFG | -6 | 56 | 24 | 0.03 | |
| 4880 | Left Precuneus | -2 | -54 | 36 | 0.06 | |
| 3656 | Left MTG | -50 | -56 | 22 | 0.07 | |
| 3496 | Right MTG | 56 | -52 | 18 | 0.05 | |
| 3376 | Right MTG | 54 | 0 | -20 | 0.03 | |
| | Right MTG | 54 | -16 | -16 | 0.03 | |
| | Right MTG | 52 | 6 | -28 | 0.03 | |
| | Right Middle TP | 48 | 10 | -32 | 0.03 | |
| | Right MTG | 60 | -10 | -14 | 0.02 | |
| 2912 | Left MTG | -54 | 2 | -26 | 0.04 | |
| | Left MTG | -60 | -10 | -14 | 0.03 | |
| | Left MTG | -62 | -20 | -10 | 0.03 | |
| 1056 | Right Gyrus Rectus | 2 | 46 | -18 | 0.03 | |
| 904 | Left Cerebellum | -24 | -78 | -36 | 0.03 | |
| 560 | Left MTG | -54 | -30 | -6 | 0.03 | |
| 8 | Right MTG | 50 | -28 | -6 | 0.02 | |
| | | **VERBAL ToM > NON-VERBAL ToM CONTRAST** | | | | |
| 752 | Left Angular | -51.8 | -59.6 | 27.8 | | 3.72 |
| 384 | Left Medial SFG | -3.8 | 48 | 25.7 | | 3.72 |
| | Left Medial SFG | 0 | 50 | 22 | | 3.54 |
| | Left Medial SFG | 0 | 54 | 24 | | 3.35 |
| 208 | Right Angular | 56 | -55 | 26 | | 3.72 |
| 112 | Right ITG | 59 | -23 | -16 | | 3.54 |
| | | **NON-VERBAL ToM > VERBAL ToM CONTRAST** | | | | |

| | | | | | |
|---|---|---|---|---|---|
| 3816 | Left MTG | -54.9 | -47.9 | 7.7 | 3.72 |
| | Left MTG | -60 | -44 | -2 | 3.54 |
| 1088 | Left Fusiform | -38.9 | -51 | -17.7 | 3.72 |
| | Left Fusiform | -42 | -48 | -18 | 3.54 |
| 768 | Right IFG (pars triangularis) | 52 | 27 | 19 | 0.00 |
| | Right IFG (pars triangularis) | 52.3 | 30.3 | 15 | 3.54 |
| | Right IFG (pars triangularis) | 54.3 | 25.8 | 21 | 3.35 |
| 568 | Left IFG (pars triangularis) | -50.7 | 21.8 | 15.5 | 3.72 |
| | Left IFG (pars opercularis) | -49.6 | 15.6 | 14.4 | 3.54 |
| 552 | Right MTG | 48.6 | -42.1 | 7.2 | 3.72 |
| | Right MTG | 47 | -48 | 10 | 3.54 |
| 344 | Left SMA | -2.7 | 22 | 55.3 | 3.72 |
| | Left SMA | -2.5 | 15.5 | 53 | 3.24 |
| 320 | Left OFG | -43 | 28 | -8 | 3.72 |
| | Left OFG | -42 | 30.3 | -4.7 | 3.54 |

*Anatomical labels are derived from the Automatic Anatomical Labelling Atlas. MTG = middle temporal gyrus, IFG - inferior frontal gyrus, AG = angular gyrus, TP = temporal pole, MOG = middle occipital gyrus, ITG = inferior temporal gyrus, SMA = supplementary motor area, SFG = superior frontal gyrus, STG = superior temporal gyrus, PHG = para hippocampal gyrus, OFG = orbitofrontal gyrus.*

***Supplementary Table R9*** *Formal conjunction and contrast analyses comparing the within domain differences of the VERBAL SC (N= 175) and NON-VERBAL SC (N= 37) experiments (cluster forming threshold p<.001; FWE cluster-extent corrected at p<.05). The contrast analyses were further thresholded with a cluster forming threshold at p<.001 and minimum cluster size of 100mm3.*

| **Cluster Size** | **Region of Activation** | **Peak MNI Co-ordinates X** | **Y** | **Z** | **ALE Value** | **Z Value** |
|---|---|---|---|---|---|---|
| | **VERBAL SC ∩ NON-VERBAL SC CONJUNCTION** | | | | | |
| 6248 | Left ITG | -48 | -52 | -16 | 0.04 | |
| | Left Fusiform Gyrus | -30 | -34 | -20 | 0.03 | |
| | Left MTG | -56 | -40 | 2 | 0.03 | |
| | Left Fusiform Gyrus | -36 | -50 | -16 | 0.02 | |
| | Left Fusiform Gyrus | -40 | -42 | -18 | 0.02 | |
| 1464 | Left IFG (pars orbitalis) | -36 | 32 | -10 | 0.03 | |
| 888 | Left IFG (pars opercularis) | -46 | 12 | 26 | 0.02 | |
| | Left IFG (pars triangularis) | -54 | 20 | 22 | 0.02 | |
| 792 | Left Precuneus | -8 | -56 | 14 | 0.02 | |
| | Left Precuneus | -6 | -54 | 10 | 0.02 | |
| 8 | Left IOG | -48 | -64 | -14 | 0.01 | |
| | **VERBAL SC > NON-VERBAL SC CONTRAST** | | | | | |
| 1016 | Left Angular Gyrus | -38.6 | -61 | 27 | | 3.72 |
| | Left MTG | -39.5 | -60 | 19 | | 3.35 |
| 864 | Left Precentral Gyrus | -45.6 | 3 | 51.4 | | 3.35 |
| | Left Precentral Gyrus | -47 | -2.9 | 53.5 | | 3.24 |
| | Left Precentral Gyrus | -46 | -2 | 46 | | 3.09 |
| 688 | Left MTG | -61.5 | -27.6 | -5.4 | | 3.72 |
| | Left MTG | -50 | -26 | -4 | | 3.54 |
| | Left MTG | -50 | -22 | -4 | | 3.16 |
| 256 | Left IFG (pars opercularis) | -46 | 18.7 | 13.7 | | 3.72 |
| 136 | Left IFG (pars orbitalis) | -56 | 24 | -6 | | 3.54 |
| | **NON-VERBAL SC > VERBAL SC CONTRAST** | | | | | |
| 768 | Left Fusiform Gyrus | -42.5 | -77.5 | -16.4 | | 3.72 |

*Anatomical labels are derived from the Automatic Anatomical Labelling Atlas. MTG = middle temporal gyrus, IFG - inferior frontal gyrus, AG = angular gyrus, TP = temporal pole, MOG = middle occipital gyrus, ITG = inferior temporal gyrus, SMA = supplementary motor area, SFG = superior frontal gyrus, STG = superior temporal gyrus, PHG = para hippocampal gyrus, OFG = orbitofrontal gyrus.*

***Supplementary Table R10*** *Formal conjunction and contrast analyses of the VISUAL ToM (N= 46) and VISUAL SC (N= 175) experiments after excluding experiments with AUDITORY stimuli; (cluster forming threshold p<.001; FWE cluster-extent corrected at p<.05). The contrast analyses were further thresholded with a cluster forming threshold at p<.001 and minimum cluster size of 100mm3*

| Cluster Size | Region of Activation | Peak MNI Co-ordinates X | Y | Z | ALE Value | Z Value |
|---|---|---|---|---|---|---|
| | **VISUAL ToM ∩ VISAUL SC CONJUNCTION** | | | | | |
| 6992 | Left IFG (pars orbitalis) | -46 | 30 | -10 | 0.08 | |
| | Left IFG (pars triangularis) | -52 | 24 | 6 | 0.07 | |
| | Left IFG (pars opercularis) | -50 | 18 | 18 | 0.06 | |
| 3808 | Left MTG | -56 | -38 | 0 | 0.07 | |
| | Left MTG | -58 | -46 | 6 | 0.05 | |
| 3152 | Left MTG | -56 | -8 | -16 | 0.07 | |
| 2560 | Left AG | -48 | -64 | 26 | 0.07 | |
| 1808 | Left SMA | -4 | 16 | 56 | 0.06 | |
| 1112 | Left Medial SFG | -8 | 52 | 36 | 0.05 | |
| 1064 | Left Fusiform | -42 | -50 | -16 | 0.04 | |
| 808 | Left Precuneus | -4 | -54 | 24 | 0.05 | |
| 504 | Left Middle TP | -48 | 12 | -28 | 0.04 | |
| | Left Middle TP | -50 | 12 | -24 | 0.04 | |
| | **VISUAL ToM > VISUAL SC CONTRAST** | | | | | |
| 6744 | Right MTG | 54.1 | -53.1 | 19.7 | | 3.72 |
| 5952 | Right Medial SFG | 3.3 | 56.6 | 22.7 | | 3.72 |
| | Left SFG | -12.9 | 54.5 | 34 | | 3.54 |
| 5592 | Right Precuneus | 2.3 | -55.2 | 36.7 | | 3.72 |
| 5000 | | 56.9 | -7.4 | -21.1 | | 3.72 |
| | Right ITG | 54 | -20 | -20 | | 3.54 |
| 4592 | Left MTG | -51.6 | -54.6 | 20.4 | | 3.72 |
| 2152 | Left Cerebellum | -24.2 | -77.5 | -36.4 | | 3.72 |
| 1752 | Right IFG (pars triangularis) | 56 | 25.9 | 7.2 | | 3.72 |
| 1608 | Right Precentral Gyrus | 42.3 | 6.4 | 43.8 | | 3.72 |
| | Right MFG | 42 | 5.3 | 53.7 | | 3.54 |
| 1000 | Left MTG | -52.2 | 5.3 | -32 | | 3.72 |

| | | | | | |
|---|---|---|---|---|---|
| | Left ITG | -49 | 12 | -40 | 3.54 |
| 528 | Right Gyrus Rectus | 3.3 | 49.3 | -19.4 | 3.72 |
| 344 | Left Middle Cingulum | 0.5 | -18.5 | 38.3 | 3.72 |
| 192 | Left MTG | -63.2 | -11.6 | -13.6 | 3.54 |
| | **VISUAL SC > VISUAL ToM CONTRAST** | | | | |
| 7440 | Left IFG (pars triangularis) | -41.5 | 30.7 | 5.2 | 3.72 |
| 5528 | Left ITG | -46.2 | -46.3 | -16.3 | 0.00 |
| | Left PHG | -25.1 | -25.9 | -18.1 | 3.24 |
| | Left Hippocampus | -26 | -22 | -16 | 3.35 |
| 1080 | Right Middle Cingulum | 3.9 | 21.8 | 41.8 | 3.72 |
| | Left Medial SFG | -9.2 | 28.4 | 44.4 | 3.54 |
| 680 | Left ITG | -39.2 | -14.1 | -26.4 | 3.72 |
| 648 | Left Calcarine | -3.7 | -57.5 | 8.3 | 3.72 |
| 472 | Right OFG | 35.2 | 33.7 | -6.7 | 3.72 |
| | Right OFG | 34 | 36 | -12 | 3.54 |
| 464 | Right PHG | 30 | -16 | -24.7 | 3.54 |
| | Right Hippocampus | 27 | -12 | -21 | 3.54 |
| | Right PHG | 22 | -14 | -24 | 3.35 |
| 208 | Left MOG | -34 | -66 | 40 | 3.72 |

*Anatomical labels are derived from the Automatic Anatomical Labelling Atlas. MTG = middle temporal gyrus, IFG - inferior frontal gyrus, AG = angular gyrus, TP = temporal pole, MOG = middle occipital gyrus, ITG = inferior temporal gyrus, SMA = supplementary motor area, SFG = superior frontal gyrus, STG = superior temporal gyrus, PHG = para hippocampal gyrus, OFG = orbitofrontal gyrus.*

***Supplementary Table R11*** *The results of the independent ALE analyses for the VISUAL ToM (N= 106) and AUDITORY ToM (N= 6) experiments (cluster forming threshold p<.001; cluster-extent FWE p<.05).*

| Cluster Size | Region of Activation | Peak MNI Co-ordinates | | | ALE Value | Z Value |
|---|---|---|---|---|---|---|
| | | X | Y | Z | | |
| | **VISUAL ToM** | | | | | |
| 23832 | Left MTG | -50 | -58 | 22 | 0.14 | 12.53 |
| | Left MTG | -60 | -10 | -14 | 0.08 | 8.16 |
| | Left MTG | -54 | 2 | -26 | 0.08 | 7.89 |
| | Left MTG | -56 | -38 | 0 | 0.07 | 7.42 |
| | Left MTG | -58 | -24 | -8 | 0.05 | 6.05 |
| 15736 | Left Medial SFG | -10 | 54 | 34 | 0.10 | 9.63 |
| | Right Medial SFG | 4 | 56 | 24 | 0.08 | 7.86 |
| | Left Gyrus Rectus | 0 | 48 | -18 | 0.06 | 6.76 |
| | Left Anterior Cingulum | -8 | 50 | 0 | 0.04 | 4.46 |
| | Left Medial SFG | -8 | 44 | 50 | 0.03 | 3.84 |
| 11192 | Right MTG | 54 | 0 | -22 | 0.08 | 8.44 |
| | Right MTG | 60 | -8 | -18 | 0.08 | 8.30 |
| | Right MTG | 52 | -34 | -2 | 0.04 | 4.63 |
| | Right MTG | 50 | -30 | -4 | 0.04 | 4.59 |
| 9816 | Right STG | 54 | -54 | 24 | 0.11 | 10.17 |
| | Right MTG | 50 | -72 | 6 | 0.04 | 4.25 |
| 9384 | Left Precuneus | -2 | -54 | 36 | 0.12 | 10.88 |
| 7312 | Left IFG (pars orbitalis) | -48 | 28 | -10 | 0.08 | 8.39 |
| | Left IFG (pars triangularis) | -52 | 24 | 6 | 0.07 | 7.05 |
| | Left IFG (pars opercularis) | -50 | 18 | 18 | 0.06 | 6.36 |
| 5616 | Right IFG (pars triangularis) | 56 | 28 | 8 | 0.08 | 8.15 |
| | Right OFG | 52 | 30 | -6 | 0.04 | 5.15 |
| | Right IFG (pars triangularis) | 44 | 20 | 24 | 0.04 | 5.09 |
| 3120 | Left SMA | -6 | 16 | 62 | 0.06 | 6.78 |
| 2264 | Left Cerebellum | -24 | -78 | -36 | 0.07 | 7.71 |
| 1896 | Right Precentral Gyrus | 42 | 6 | 42 | 0.05 | 6.01 |
| 1120 | Right Cerebellum | 28 | -78 | -34 | 0.06 | 6.47 |
| 1064 | Left Fusiform | -42 | -50 | -16 | 0.04 | 5.07 |
| 848 | Left Middle Cingulum | 0 | -16 | 38 | 0.05 | 5.62 |
| | **AUDITORY ToM** | | | | | |

NA

---

*Anatomical labels are derived from the Automatic Anatomical Labelling Atlas. MTG = middle temporal gyrus, IFG - inferior frontal gyrus, AG = angular gyrus, TP = temporal pole, MOG = middle occipital gyrus, ITG = inferior temporal gyrus, SMA = supplementary motor area, SFG = superior frontal gyrus, STG = superior temporal gyrus, PHG = para hippocampal gyrus, OFG = orbitofrontal gyrus.*

***Supplementary Table R12** The results of the independent ALE analyses for the VISUAL SC (N= 152) and AUDITORY SC (N= 60) experiments (cluster forming threshold p<.001; cluster-extent FWE p<.05).*

| Cluster Size | Region of Activation | Peak MNI Co-ordinates X | Y | Z | ALE Value | Z Value |
|---|---|---|---|---|---|---|
| | | **VISUAL SC** | | | | |
| 30792 | Left IFG (pars triangularis) | -50 | 30 | 6 | 0.11 | 10.10 |
| | Left IFG (pars triangularis) | -48 | 24 | 16 | 0.11 | 9.68 |
| | Left IFG (pars orbitalis) | -36 | 32 | -14 | 0.09 | 8.32 |
| | Left IFG (pars orbitalis) | -40 | 30 | -12 | 0.09 | 8.28 |
| | Left Insula | -34 | 26 | -2 | 0.07 | 6.48 |
| | Left Middle TP | -46 | 16 | -26 | 0.06 | 6.32 |
| | Left Insula | -42 | 18 | 4 | 0.05 | 4.78 |
| | Left Superior TP | -48 | 12 | -14 | 0.04 | 4.14 |
| 25792 | Left MTG | -56 | -38 | 2 | 0.13 | 11.03 |
| | Left Fusiform | -46 | -54 | -16 | 0.10 | 9.29 |
| | Left Fusiform | -30 | -34 | -20 | 0.10 | 9.01 |
| | Left Fusiform | -38 | -42 | -20 | 0.09 | 8.22 |
| | Left Hippocampus | -22 | -8 | -16 | 0.08 | 7.25 |
| 10584 | Left SMA | -4 | 18 | 50 | 0.10 | 9.38 |
| | Left SMA | -4 | 6 | 58 | 0.06 | 6.15 |
| | Left Medial SFG | -2 | 32 | 42 | 0.05 | 5.37 |
| | Left Medial SFG | -8 | 52 | 36 | 0.05 | 5.22 |
| | Left SFG | -14 | 30 | 46 | 0.04 | 4.33 |
| | Left MFG | -24 | 26 | 46 | 0.04 | 3.73 |
| 5872 | Left AG | -48 | -66 | 26 | 0.07 | 6.88 |
| | Left MOG | -38 | -76 | 38 | 0.05 | 4.92 |
| | Left MOG | -32 | -64 | 36 | 0.04 | 4.32 |
| | Left SPL | -30 | -60 | 48 | 0.04 | 3.89 |
| 5352 | Left MTG | -56 | -6 | -16 | 0.08 | 7.23 |
| | Left ITG | -42 | -14 | -28 | 0.05 | 4.91 |
| | Left ITG | -36 | -4 | -32 | 0.04 | 3.60 |
| 3752 | Right Insula | 36 | 24 | -2 | 0.07 | 6.42 |
| | Right OFG | 36 | 34 | -12 | 0.06 | 5.85 |
| 3096 | Left Precuneus | -6 | -56 | 14 | 0.06 | 6.12 |
| | Left Precuneus | -6 | -56 | 10 | 0.06 | 5.97 |
| | Right Calcarine | 8 | -58 | 12 | 0.03 | 3.44 |
| 1984 | Left Precentral Gyrus | -48 | -2 | 50 | 0.05 | 5.29 |
| | Left Postcentral Gyrus | -52 | -8 | 44 | 0.05 | 4.99 |

| | | | | | | |
|---|---|---|---|---|---|---|
| 1376 | Right PHG | 26 | -16 | -22 | 0.05 | 5.45 |
| | Right Hippocampus | 26 | -6 | -18 | 0.03 | 3.54 |
| | **AUDITORY SC** | | | | | |
| 16160 | Left MTG | -56 | -6 | -14 | 0.05 | 7.72 |
| | Left MTG | -60 | -18 | -2 | 0.05 | 7.40 |
| | Left MTG | -56 | -38 | 4 | 0.04 | 6.82 |
| | Left MTG | -64 | -52 | 0 | 0.02 | 3.58 |
| | Left Middle TP | -48 | 14 | -24 | 0.02 | 3.29 |
| 10336 | Right STG | 62 | -8 | -4 | 0.04 | 6.94 |
| | Right Superior TP | 50 | 14 | -20 | 0.02 | 4.50 |
| | Right STG | 54 | -28 | 2 | 0.02 | 4.30 |
| | Right Middle TP | 44 | 12 | -28 | 0.02 | 4.17 |
| | Right STG | 46 | -22 | 8 | 0.02 | 3.76 |
| 7928 | Left IFG (pars opercularis) | -50 | 18 | 18 | 0.04 | 6.19 |
| | Left IFG (pars triangularis) | -48 | 30 | 8 | 0.04 | 5.97 |
| | Left IFG (pars orbitalis) | -44 | 28 | -12 | 0.03 | 5.17 |
| 3464 | Left AG | -46 | -62 | 26 | 0.03 | 5.58 |
| 3280 | Left Fusiform | -28 | -36 | -18 | 0.04 | 6.20 |
| | Left Fusiform | -40 | -42 | -18 | 0.04 | 6.14 |
| 1520 | Left SMA | -4 | 10 | 56 | 0.03 | 5.42 |

*Anatomical labels are derived from the Automatic Anatomical Labelling Atlas. MTG = middle temporal gyrus, IFG - inferior frontal gyrus, AG = angular gyrus, TP = temporal pole, MOG = middle occipital gyrus, ITG = inferior temporal gyrus, SMA = supplementary motor area, SFG = superior frontal gyrus, STG = superior temporal gyrus, PHG = para hippocampal gyrus, OFG = orbitofrontal gyrus.*

***Supplementary Table R13** Formal conjunction and contrast analyses comparing the within domain differences of the VISUAL SC (N= 152) and AUDITORY SC (N= 60) experiments (cluster forming threshold p<.001; FWE cluster-extent corrected at p<.05). The contrast* *analyses were further thresholded with a cluster forming threshold at p<.001 and minimum cluster size of 100mm3.*

| Cluster Size | Region of Activation | Peak MNI Co-ordinates | | | ALE Value | Z Value |
|---|---|---|---|---|---|---|
| | | X | Y | Z | | |
| | **VISUAL SC ∩ AUDITORY SC CONJUNCTION** | | | | | |
| 7872 | Left IFG (pars opercularis) | -50 | 18 | 18 | 0.04 | |
| | Left IFG (pars triangularis) | -48 | 30 | 8 | 0.04 | |
| | Left IFG (pars orbitalis) | -44 | 28 | -12 | 0.03 | |
| 4848 | Left MTG | -56 | -38 | 4 | 0.04 | |
| | Left MTG | -56 | -26 | -4 | 0.03 | |
| | Left MTG | -62 | -50 | 0 | 0.02 | |
| 3248 | Left Fusiform | -28 | -36 | -18 | 0.04 | |
| | Left Fusiform | -40 | -42 | -18 | 0.04 | |
| 2960 | Left MTG | -56 | -6 | -14 | 0.05 | |
| 2240 | Left AG | -46 | -62 | 26 | 0.03 | |
| 1520 | Left SMA | -4 | 10 | 56 | 0.03 | |
| 360 | Left Superior TP | -52 | 8 | -18 | 0.02 | |
| | Left Middle TP | -48 | 14 | -24 | 0.02 | |
| | **VISUAL SC > AUDITORY SC CONTRAST** | | | | | |
| 576 | Left Fusiform | -41.6 | -57.7 | -21 | | 3.72 |
| 351 | Left Medial SFG | -4 | 34 | 44 | | 3.72 |
| | Left Medial SFG | -2 | 34 | 48 | | 3.54 |
| | Left Medial SFG | -6 | 36 | 48 | | 3.16 |
| | **AUDITORY SC > VISUAL SC CONTRAST** | | | | | |
| 2560 | Right STG | 62.6 | -9.7 | -4.3 | | 3.72 |
| | Right MTG | 65.9 | -17.1 | -6.9 | | 3.54 |
| 1688 | Left STG | -63 | -17.4 | 0.9 | | 3.72 |
| 104 | Right STG | 50 | -23 | 7 | | 3.72 |

*Anatomical labels are derived from the Automatic Anatomical Labelling Atlas. MTG = middle temporal gyrus, IFG - inferior frontal gyrus, AG = angular gyrus, TP = temporal pole, MOG = middle occipital gyrus, ITG = inferior temporal gyrus, SMA = supplementary motor area, SFG = superior frontal gyrus, STG = superior temporal gyrus, PHG = para hippocampal gyrus, OFG = orbitofrontal gyrus.*

#### Cluster Analyses

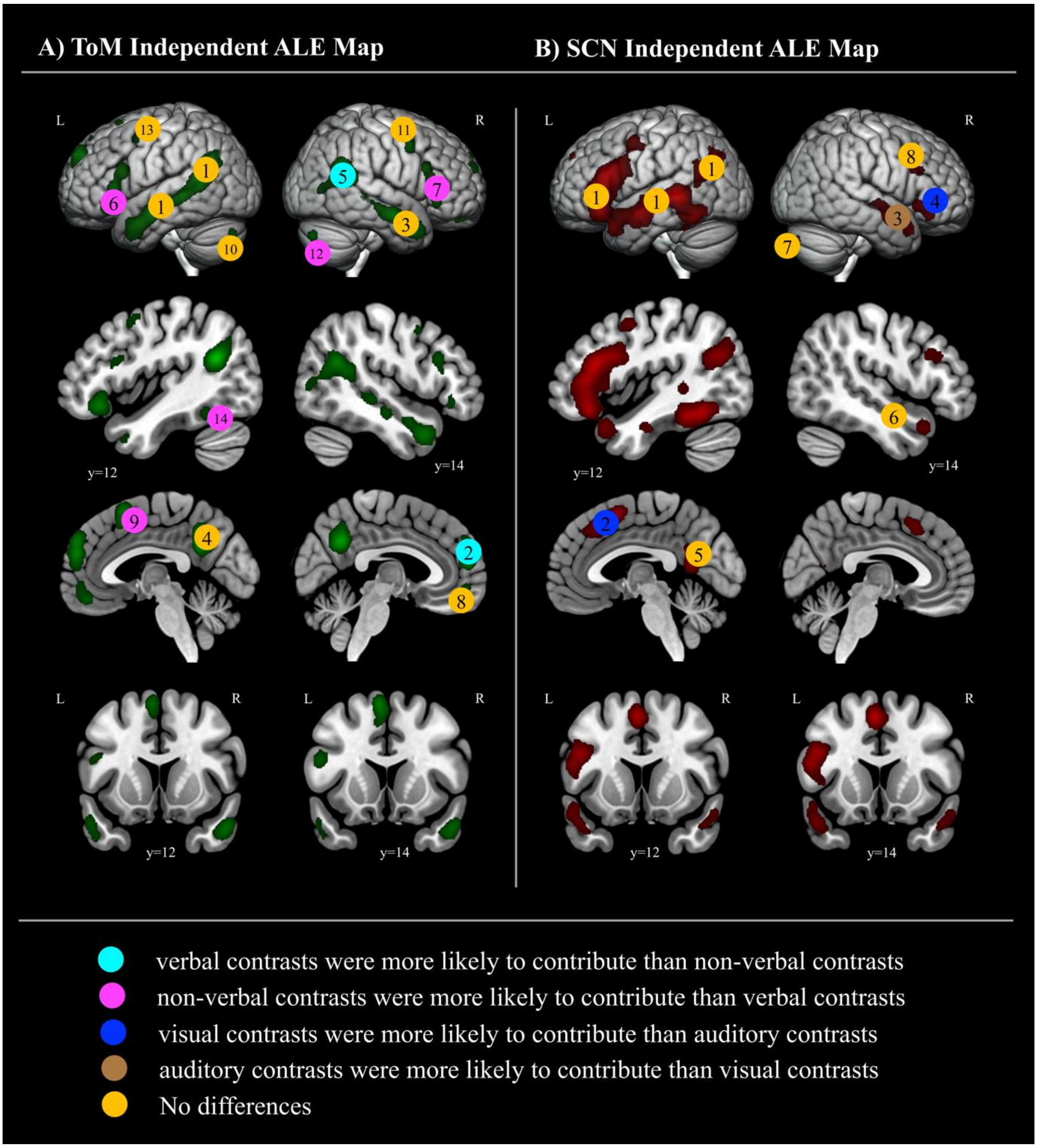

***Supplementary Figure CA1*** *The outcomes of the cluster analyses comparing the likelihood of higher / lower contribution of different experiment type (verbal, non-verbal, visual, auditory) to each cluster that was initially revealed by the main independent ALE analysis of the ToM and SC data sets.*

***Supplementary Table CA1*** *The outcomes of the cluster analyses comparing the likelihood of higher / lower contribution of different experiment type (verbal, non-verbal, visual, auditory) to each cluster that was initially revealed by the main independent ALE analysis of the ToM and SC data sets*

| **Data Set** | **Cluster** | **Cluster Location** | **Verbal vs. Non-Verbal** | | **Visual vs. Auditory** | |
|---|---|---|---|---|---|---|
| | | | **P** | **OR** | **P** | **OR** |
| **Theory of Mind** | 1 | left MTG | 0.444 | 0.645 | 0.238 | 3.013 |
| | 2 | anterior medial SFG | 0.017 | 2.745 | 1.000 | 0.777 |
| | 3 | right MTG | 0.842 | 1.124 | 1.000 | 0.904 |
| | 4 | Precunesus | 0.164 | 1.847 | 1.000 | 0.807 |
| | 5 | right posterior STG/AG | 0.043 | 2.469 | 0.649 | 2.099 |
| | 6 | left IFG | <.001 | 0.127 | 1.000 | 1.149 |
| | 7 | right IFG | 0.003 | 0.238 | 0.178 | Inf |
| | 8 | mPFC | 0.361 | 1.565 | 0.333 | Inf |
| | 9 | left posterior medial SFG | 0.003 | 0.134 | 1.000 | 1.107 |
| | 10 | left Cerebellum | 0.612 | 0.708 | 1.000 | 0.875 |
| | 11 | right Precentral | 0.755 | 1.283 | 1.000 | Inf |
| | 12 | right Cerebellum | 0.054 | 0.156 | 1.000 | Inf |
| | 13 | left Precentral | 0.708 | 0.564 | 0.317 | 0.288 |
| | 14 | left pITG | 0.013 | 0.000 | 1.000 | Inf |
| **Semantic Cognition** | 1 | left IFG-aSTG-MTG-AG | 0.079 | 0.277 | 1.000 | 0.957 |
| | 2 | posterior medial SFG | 0.827 | 0.904 | 0.033 | 2.134 |
| | 3 | right STG | 1.000 | 1.206 | 0.026 | 0.428 |
| | 4 | right IFG | 0.775 | 0.919 | 0.051 | 2.962 |
| | 5 | left Precuneus | 0.378 | 0.645 | 0.189 | 2.026 |
| | 6 | right medial TL | 0.134 | 0.411 | 0.785 | 1.330 |
| | 7 | right Cerebellum | 0.375 | Inf | 0.764 | 1.526 |
| | 8 | right IFG opercularis | 0.600 | 0.822 | 0.686 | 2.671 |

We conducted complementary cluster analyses to assess whether VERBAL, NON-VERBAL, VISUAL and AUDITORY experiments were equally likely to contribute to each activation cluster that was revealed by the primary independent ALE analyses in both the Theory of Mind (ToM) and Semantic Cognition (SC) data sets comprising of all experiment type (see Supplementary Figure 11.). We were also interested if these cluster analyses would reflect the results of our sub-analyses conducted after splitting the data into verbal, non-verbal, visual and auditory experiments. Accordingly, we conducted Fischer's exact tests of independence and then followed up the significant effects with pairwise comparisons. We report the strength of the association as given by the alpha value and the direction of the association as given by the odds ratio (OR).

In the ToM data set, the cluster analysis revealed that all experiment type were likely to contribute equally to clusters in the left MTG (cluster 1; p=0.443; OR=0.645 for verbal vs. non-verbal and p= 0.238; OR=3.013 for visual vs. auditory), right MTG (cluster 3; p=0.842; OR=1.124 for verbal vs. non-verbal and p=1; OR=0.904 for visual vs. auditory), left precuneus (cluster 4; p=0.164; OR=1.847 for verbal vs. non-verbal and p=1; OR=0.807 for visual vs. auditory), left gyrus rectus/anterior cingulum (cluster 8; p=0.361; OR=1.565 for verbal vs. non-verbal and p=0.332; OR= Inf for visual vs. auditory. Note: area corresponds to mPFC), left cerebellum (cluster 10; p=0.612; OR= 0.708 for verbal vs. non-verbal and p=1; OR=0.875 for visual vs. auditory), right precentral gyrus (cluster 11; p= 0.755; OR= 1.283for verbal vs. non-verbal and p=1; OR= Inf for visual vs. auditory), left precentral gyrus (cluster 13, p=0.708; OR= 0.564 for verbal vs. non-verbal and p=0.317; OR= 0.288 for visual vs. auditory).

However, when comparing the verbal vs. non-verbal experiments we observed significant differences in the bilateral anterior medial SFG (cluster 2; p=0.017; OR=2.745) whereby verbal experiments were 2.74 times more likely to contribute to this cluster as opposed to non-verbal experiments. There were also significant differences in the right pMTG/pSTS (cluster 5; p=0.043; OR=2.469), whereby verbal experiments were 2.47 times more likely to contribute than non-verbal experiments. Significant difference was also observed in the in the bilateral IFG (cluster 6, p=<.001; OR= 0.127and cluster 7; p= <.00023; OR= 0.238) with the verbal experiments being 80% less likely to contribute to the left IFG and 76% less likely to contribute to the right IFG cluster than the non-verbal experiments. This is in line with our sub analyses in which after splitting the ToM data to verbal and non-verbal there is no converging IFG activation for the verbal experiments. We also observed a difference in the medial SFG (cluster 9, p=<.00023; OR= 0.134) whereby verbal experiments were 87% less likely to contribute than the non- verbal experiments. There was also a difference in the right cerebellum (cluster 12, p=0.054; OR= 0.156) with the verbal experiments being 85% less likely to contribute than the non-verbal experiments. Finally, there was also a difference in the left posterior ITG (cluster 14; p=0.013; OR= 0) with the verbal experiments being 100% less likely to contribute than the non-verbal experiments. There were no differences observed for the visual and auditory experiments in neither of the clusters.

For the SC data set, the cluster analysis revealed that all experiment type were likely to contribute equally to the cluster stretching from the left IFG via the anterior STG, along the length of the MTG right back to the AG (cluster 1, p=0.079; OR= 0.277 for verbal vs. non-verbal and p=1; OR=0.957 for visual vs. auditory), left precuneus (cluster 5; p=0.378; OR= 0.645 for verbal vs. non-verbal and p=0.189; OR= 2.026 for visual vs. auditory); right medial ATL (cluster 6, p=0.134; OR=0.411 for verbal vs. non-verbal and p=0.785; OR= 1.330 for visual vs. auditory); right cerebellum (cluster 7, p= 0.375; OR= Inf for verbal vs. non-verbal and p=0.764; OR=1.526 for visual vs. auditory), right IFG pars opercularis (cluster 8, p=0.600; OR=0.822 for verbal vs. non-verbal and p=0.686; OR=2.671 for visual vs. auditory).

For the visual vs. auditory data, we observed significant differences in the medial SFG (cluster 2, p=0.033; OR= 2.134) whereby the visual experiments were 2.13 time more likely to contribute to this cluster than the auditory experiments. There were also significant differences in the right STG/MTG (cluster 3, p=0.026; OR= 0.428) with the visual experiments being 57% less likely to contribute to this cluster than the auditory experiments. This is also what we can see after the visual-auditory data split where the auditory experiments show bilateral ATL/MTG pattern and the visual are left lateralised. There were also significant differences in the right IFG (cluster 4, p=0.051; OR= 2.962) whereby there was a 2.96 times higher likelihood of the visual experiments contributing more to this cluster than the auditory experiments. There were no verbal vs. non-verbal differences observed in neither of the clusters of the SC data set.
